# Engineering *Escherichia coli* to utilize erythritol as sole carbon source

**DOI:** 10.1101/2022.10.03.510612

**Authors:** Fang Ba, Xiangyang Ji, Shuhui Huang, Yufei Zhang, Wan-Qiu Liu, Yifan Liu, Shengjie Ling, Jian Li

## Abstract

Erythritol, one of the natural sugar alcohols, is widely used as a sugar substitute sweetener in food industries. Human themselves are not able to catabolize erythritol and their gut microbes lack related catabolic pathways either to metabolize erythritol. Here, we engineer *Escherichia coli* to utilize erythritol as sole carbon source aiming for defined applications. First, we isolate the erythritol metabolic gene cluster and experimentally characterize the erythritol-binding transcriptional repressor and its DNA-binding site. Transcriptome analysis suggests that carbohydrate metabolism-related genes in the engineered *E. coli* are overall upregulated, which then guides the selection of four genes for overexpression that notably enhances cell growth. Finally, engineered *E. coli* strains can be used as a living detector to distinguish erythritol-containing soda soft drinks and can grow in the simulated intestinal fluid supplemented with erythritol. We expect our work will inspire the engineering of more hosts to respond and utilize erythritol for broad applications in metabolic engineering, synthetic biology, and biomedical engineering.

## Introduction

The global obesity epidemic and correlative diseases like diabetes, hypertension, and hyperlipidemia encourage people to change their dietary style by limiting excessive intake of carbohydrates. Yet, many people like sweet food. In this context, food industry is seeking to produce more carbohydrate/sugar-reduced foods by using alternative sweeteners such as sugar alcohols (also called polyols that contain multiple hydroxyl groups)^1, 2^. Erythritol, a C4 sugar alcohol, naturally exists in some vegetables, fruits, mushrooms, and fermented foods^3, 4^. It also has been found in human tissues such as semen, lens, cerebrospinal fluid, and serum^1, 4, 5^. Currently, erythritol is primarily produced by fermentation using engineered lactic acid bacteria and yeast^6–10^. Because erythritol is stable in a wide range of temperature and pH conditions, is not reactive in Maillard reactions, and is a non-hygroscopic compound^11^, it has been recognized as a stable, cheap, and safe sugar alcohol as well as a natural sweetener for food industry. Since human bodies are not able to catabolize erythritol, most erythritol is absorbed in the small intestine and then directly excreted in urine within 24 h^5, 11–14^. In addition, the human gut microbiota also lacks microbes to metabolize erythritol. Therefore, erythritol by food intake cannot be utilized by human bodies as a nutrient/energy source^15, 16^.

So far, two distinct erythritol degradation pathways have been identified from different species of microorganisms. The first pathway was discovered in several *Hyphomicrobiales* species, including the environmental bacterium *Ochrobactrum*^17^, the pathogenic bacterium *Brucella*^18–23^, and the nitrogen-fixing plant endosymbionts *Rhizobium*^22^ and *Sinorhizobium*^24^. These microorganisms usually uptake erythritol as an initial nutrient and yield erythrose-4-phosphate via a five-step catalysis^17^. The second pathway was characterized in *Mycolicibacterium smegmatis*^25^. This pathway contains a short three-step catalysis and the final product is also erythrose-4-phosphate. In both erythritol degradation pathways, erythrose-4-phosphate can serve as an intermediate product of the pentose-phosphate pathway, entering into other primary metabolic pathways (e.g., glycolysis) as a C4 carbon source. Without such degradation pathways, most other microorganisms are not able to metabolize erythritol. To the best of our knowledge, no attempts have been made reprogramming a model chassis microbe to live solely on erythritol as a carbon source. However, this is important because erythritol is a safe compound not used by human metabolism, but it can work as a unique molecule to support the growth of engineered microbes/probiotics, which then will be potentially used as living therapeutics for disease treatment in human gut microbiota or other focal tissues (e.g., tumors).

In metabolic engineering and synthetic biology, *Escherichia coli* has been engineered for a broad range of applications with a main focus on the production of valuable products^26–28^. However, the cultivation of *E. coli* in laboratory is predominantly based on easy-to-use carbon sources, especially, glucose (C6). Of note, a few studies recently tried to engineer *E. coli* to use C1-based carbon sources including CO_2_^29, 30^ and methanol^31, 32^. Yet, the use of C4-based carbon sources (e.g., erythritol) for *E. coli* cultivation has not been reported so far.

To address this opportunity, here we report two engineered *E. coli* strains (MG1655^33^ and Nissle 1917^34^) that can utilize erythritol as a carbon nutrient. The gene cluster of a five-step erythritol metabolic pathway is obtained from *Ochrobactrum* spp., which is isolated from the outdoor aerosol. To use the metabolic pathway, we initially characterize the erythritol-responding genetic repressor eryD and its DNA-binding site. Then, we perform mRNA transcriptional analysis to evaluate the metabolism of erythritol in *E. coli* MG1655. On the basis of the transcriptome data, we design genetic circuits to facilitate erythritol catabolism and significantly increase the growth of *E. coli* in a modified M9 medium (glucose is replaced with erythritol). Using the engineered *E. coli* MG1655, soft drinks that contain erythritol (as a sugar-free sweetener) can be distinguished from those without erythritol according to the cell growth. Moreover, the probiotic *E. coli* Nissle 1917 (EcN) is also reprogrammed with the erythritol metabolic pathway, allowing EcN to grow in simulated intestinal fluid (SIF) containing erythritol as sole carbon source. Taken together, our study shows the successful transformation of a natural metabolism of erythritol into surrogate *E. coli* hosts for defined applications. Importantly, we fill the gap that *E. coli* can utilize not only C6 (e.g., glucose) and C1-based carbon sources (CO_2_ and methanol), but also C4-based erythritol in this work. Looking forward, we anticipate that the erythritol catabolic *E. coli* strains will provide new opportunities for compelling research in different fields such as carbon cycle, synthetic biology, metabolic engineering, biomedical engineering, and living therapeutics.

## Results

### Engineering *E. coli* to utilize erythritol

To obtain the gene cluster for erythritol metabolism, we initially designed an erythritol-based selective medium derived from the M9 minimal medium, in which glucose is completely replaced with erythritol (0.4%), to isolate microbes from the outdoor aerosol that can grow in the modified medium. By doing this, we successfully separated, identified, and characterized an environmental bacterium *Ochrobactrum* spp. from the aerosol (see **Supplementary Figs. 1** and **2** for the isolation and characterization process)^35^. Then, ten erythritol catabolism-associated genes were PCR amplified from the genome of *Ochrobactrum* spp. followed by sequencing. Both DNA sequence and protein alignments suggested that all ten genes/proteins were almost the same as those of the reported *Brucella abortus* 2308 strain^36, 37^ (**Supplementary Figs. 3-5**), including five erythritol catabolic genes (*eryA*, *eryB*, *eryC*, *eryH*, and *eryI*), two putative erythritol-binding transcriptional repressor (*eryD* and *eryR*), and three erythritol ABC-transporter genes (*eryE*, *eryF*, and *eryG*) (**Fig. 1a**). Catalytic functions of the five-step erythritol catabolism proteins have been demonstrated previously *in vitro*^17^. At the end of the pathway, erythritol is eventually converted to D-erythrose 4-phosphate, which serves as one intermediate in the pentose-phosphate pathway (**Fig. 1b**). This C4 phosphate-substrate will then enter into carbohydrate utilization via glycolysis, nucleotide synthesis, and the shikimate pathway for aromatic amino acids biosynthesis (**Fig. 1b**).

**Fig. 1.**
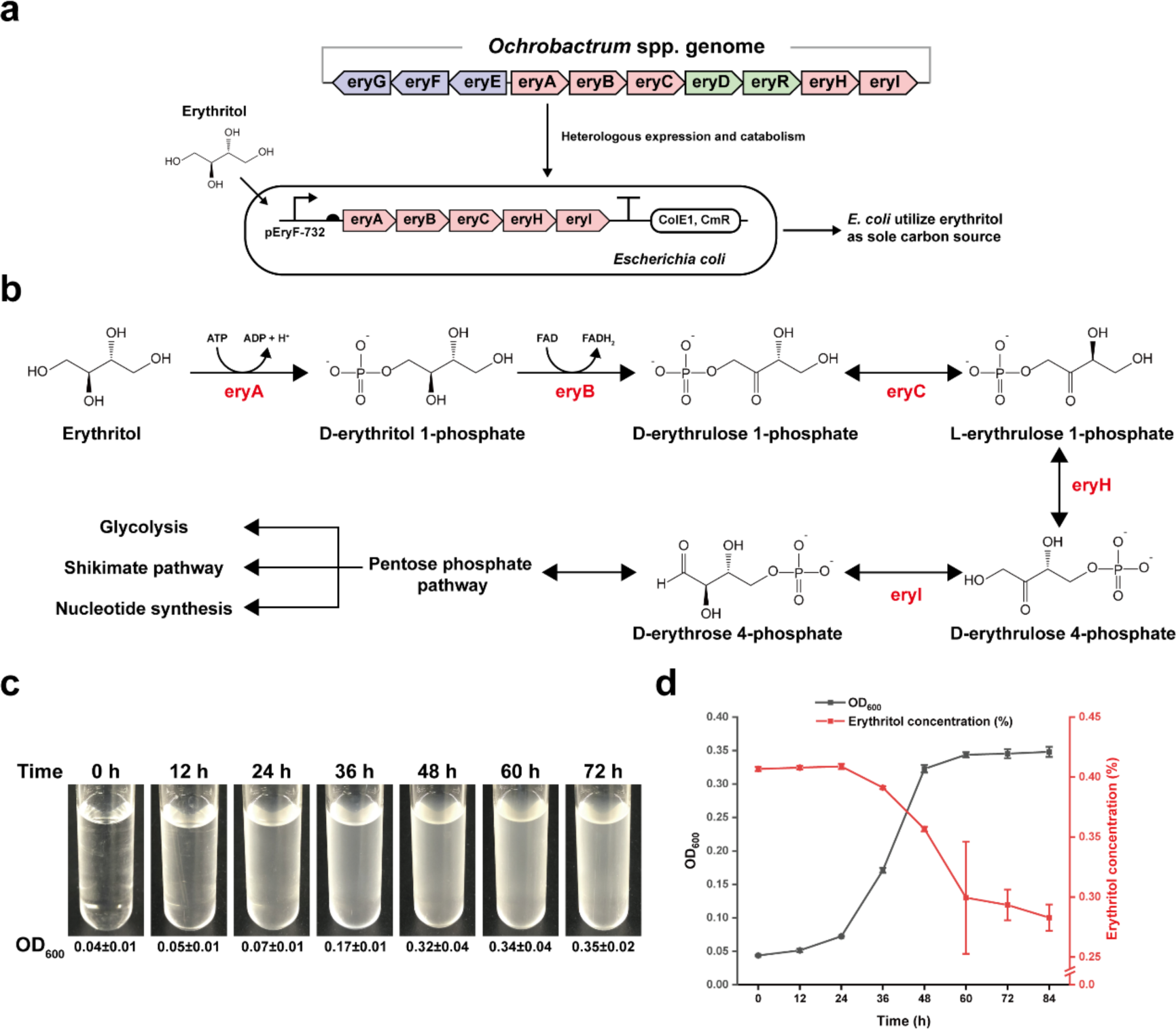
Reconstitution of the erythritol metabolic pathway in *E. coli* to support cell growth on erythritol as sole carbon source. (**a**) The erythritol metabolism gene cluster isolated from *Ochrobactrum* spp. and reconstituted in *E. coli* to utilize erythritol. (**b**) Catalytic pathway for the conversion of erythritol to D-erythrose 4-phosphate, which then enters into the downstream metabolic pathways. (**c**) The growth of *E. coli* MG1655 harboring plasmid pFB147 in liquid M9-erythritol medium. Each OD_600_ value (average ± standard deviation) was measured with three biological replicates. (**d**) Cell growth curve and the profile of erythritol utilization in M9-erythritol medium. Erythritol concentrations were measured by HPLC with three biological replicates. The error bars represent the standard deviation (s.d.).

Next, we aimed to reconstitute the gene cluster of erythritol metabolism (i.e., *eryA*, *eryB*, *eryC*, *eryH*, and *eryI*) in *E. coli* to see if cells can grow on erythritol as sole carbon source. To this end, we constructed a plasmid containing the above five genes and transformed it into *E. coli* MG1655. Then, the cells were cultivated in liquid M9-erythritol medium with shaking for 72 h (**Fig. 1c**). Clearly, *E. coli* cells were able to grow by using erythritol and the final OD_600_ value reached approximately 0.35. To determine the profile of erythritol consumption, filtered medium supernatants were analyzed by HPLC. The results indicated that along with cell growth erythritol was gradually consumed by *E. coli* with 0.28% erythritol left in the medium (**Fig. 1d**). Since the gene cluster worked to support *E. coli* growth, we were curious about the expression level of each protein *in vivo* as well as the strength of their regulatory parts (e.g., promoter and ribosomal binding site – RBS). First, all *in vivo* expressed proteins excepted two membrane proteins (eryE and eryG) were analyzed by SDS-PAGE and Western-Blot (**Extended Data Fig. 1**). The results showed that all proteins could be well expressed with a high percentage of soluble fractions. Second, the strength of the native promoters and RBSs of these genes (except *eryR*) were measured and compared with the iGEM standard regulatory parts (**Extended Data Figs. 2** and **3**)^38–40^. Overall, their strength was low as compared to the iGEM standard elements. In addition, three erythritol ABC-transporter proteins (eryE, eryF, and eryG) were predicted and, in particular, the expression, transmembrane phenotype, and localization in *E. coli* cells of the two membrane proteins (eryE and eryG) were demonstrated (**Supplementary Fig. 6** and **Extended Data Fig. 4**). However, it is worth noting that *E. coli* MG1655 could transport erythritol without this ABC-transporter system (**Fig. 1a**, **c**), suggesting that *E. coli*’s native transporter systems can also be responsible for erythritol transportation.

### eryD is an erythritol-binding transcriptional repressor

Sangari *et al*. reported that erythritol might act as an inducer and probably bind to eryD in *Brucella abortus* 2308 to inhibit the repressor activity of eryD^19^. However, a detailed biochemical characterization of eryD is still lacking. Interestingly, our isolated erythritol cluster of *Ochrobactrum* spp. contains an *eryD* gene, whose DNA sequence is totally aligned with *eryD* in *Brucella abortus* 2308 (**Supplementary Fig. 4**). In addition, DNA sequence analysis indicated that eryR, another putative DNA-binding transcriptional repressor, might also act in a similar way as eryD. Hence, characterization of these two proteins will help understand their regulation mechanism and expand erythritol-associated synthetic biology applications. To this end, we sought to determine the putative DNA-binding sites within two regions. One is the 732 bp gap between eryE and eryA and the other region is approximately 400 bp gap between eryD and eryR (**Supplementary Fig. 3**). Here, we designed an *in vivo* characterization strategy to precisely identify the DNA-binding sites. In principle, the repressor’s DNA-binding site was supposed to be located after a promoter. We individually evaluated different putative DNA sequences by using a reporter plasmid, which is derived from the iGEM standard backbone of pSB1C3 containing the reporter sfGFP. First, both directions of the 732 bp gap between eryE and eryA were presumed as two promoters called pEryF-732 (F, forward) and pEryR-732 (R, reverse), respectively (**Supplementary Figs. 7** and **8**). Meanwhile, several compatible plasmids were constructed to express eryD or eryR using a gradient strength of constitutive promoters (**Supplementary Fig. 9**). After coexpression of eryD or eryR with sfGFP, the results indicated that only eryD had a DNA-binding site located in pEryF-732 and the eryD expression level could regulate the sfGFP expression level, whereas eryR did not show an inhibition of sfGFP expression (**Supplementary Fig. 9**). With this pEryF-732 region, we then performed truncation experiments to precisely identify both 5’- and 3’-site of eryD DNA-binding site (we call it as an erythritol operator, eryO) (**Supplementary Figs. 10-15**). Finally, eryO was successfully identified as a 29 bp sequence located between the -35 and -10 regions of the promoter pEryF-732 (**Fig. 2a**). Moreover, we carried out *in vitro* electrophoretic gel mobility shift assay (EMSA) by incubation of the FAM-labeled DNA fragment (pEryF, 732 bp) with different amounts of purified eryD. The EMSA results suggested that eryD was able to effectively bind to the pEryF-732 promoter (**Fig. 2b**) and a further investigation showed that eryD could precisely bind to the 29 bp region of eryO as well (**Supplementary Fig. 16**). Using the same truncation strategy, however, we did not observe any DNA-binding sites for eryD or eryR in the 400 bp region between eryD and eryR (**Supplementary Fig. 17**).

**Fig. 2.**
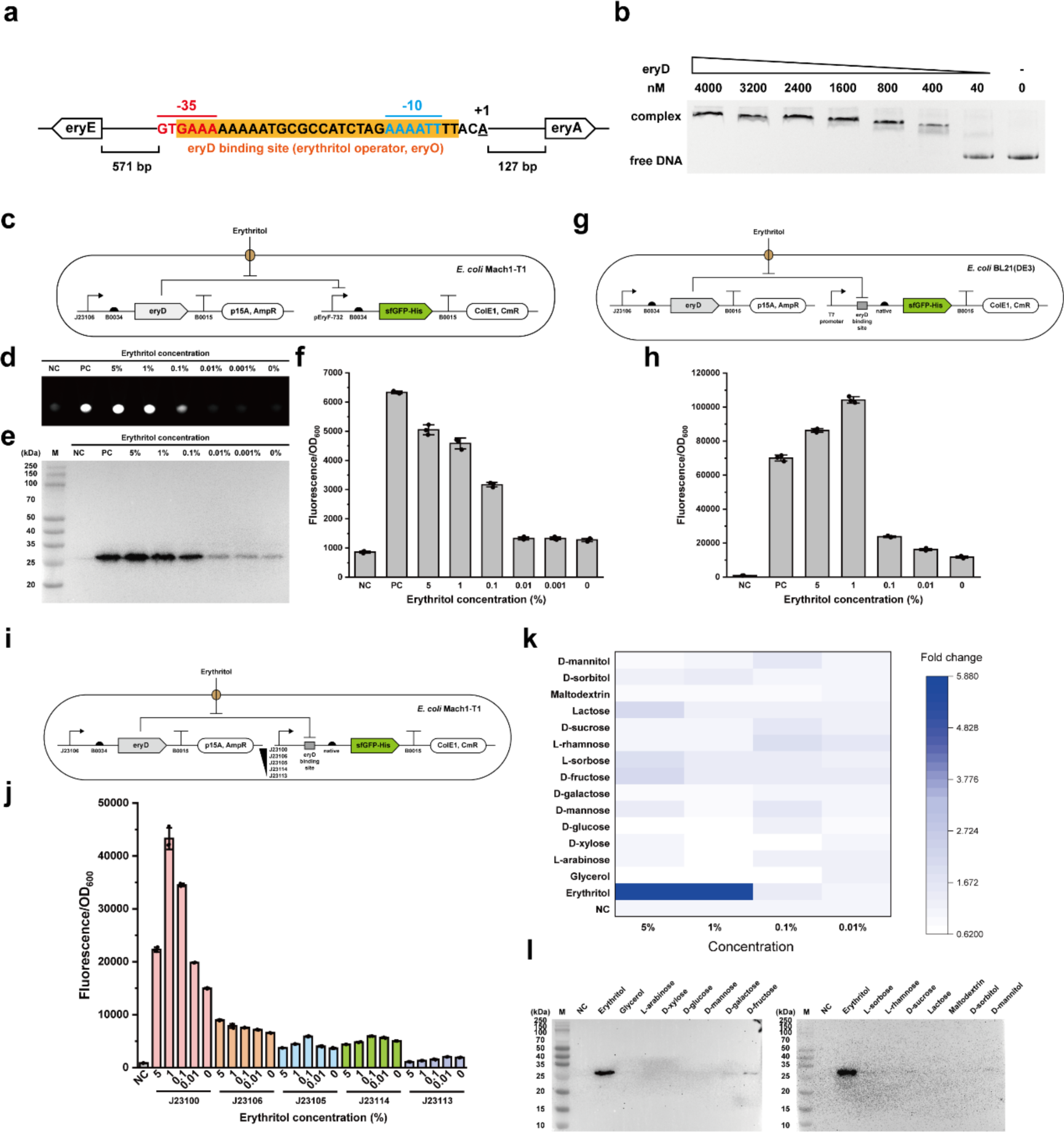
Characterization of the transcriptional repressor eryD. (**a**) DNA sequence of eryD binding site (erythritol operator, eryO, orange highlighted), promoter pEryF-732 sequence (-35, -10, and +1 regions were annotated as red, blue, and black respectively), and their genetic location in the erythritol catabolism cluster. (**b**) EMSA analysis of the interaction between eryD and pEryF-732 DNA fragment (5’FAM-labelled). (**c**) Schematic shows two compatible plasmids pFB261 and pFB186 co-existed in *E. coli* Mach1-T1. sfGFP fluorescence indicates the transcription strength of pEryF-732, which is repressed by eryD. (**d**, **f**) Relative sfGFP expression and fluorescence of *E. coli* Mach1-T1 harboring two plasmids in (**c**). Pellets were collected from 1.5 mL overnight LB culture (37°C and 250 rpm for 16 h) with supplemented erythritol. The pellets were measured as normalized fluorescence with three biological replicates. (**e**) Western-Blot analysis of sfGFP (C-terminal 6xHis tag: 27.6 kDa). (**g**, **h**) Schematic diagram of used strains and plasmids for pT7-eryO operon characterization. pFB276 and pFB261 co-existed in *E. coli* BL21(DE3) and cell pellets were collected from 1.5 mL overnight LB culture (37°C and 250 rpm for 16 h) with supplemented erythritol. The pellets were measured as normalized fluorescence with three biological replicates. (**i**, **j**) Schematic diagram of used strains and plasmids for J231xx-eryO operons characterization. pFB261 and pFB270-pFB275 co-existed in *E. coli* Mach1-T1 and cell pellets were collected from 1.5 mL overnight LB culture (37°C and 250 rpm for 16 h) with supplemented erythritol. The pellets were measured as normalized fluorescence with three biological replicates. (**k**, **l**) eryD-erythritol orthogonality test was performed by the same strains and procedure as described in (**c**). sfGFP expression was analyzed by normalized fluorescence fold change and Western-Blot. All the error bars represent the standard deviation (s.d.).

Next, we aimed to further characterize eryD. First, structure prediction by both AlphaFold and Robetta indicated that eryD monomer has an N-terminal helix-turn-helix motif responsible for DNA interaction and the C-terminal domain for erythritol regulation. SWISS-MODEL prediction showed that eryD is assembled as a homotetramer (**Supplementary Fig. 18**), which is also confirmed by size exclusion chromatography (SEC) and a native-PAGE gel analysis (the calculated molecular weight of eryD homotetramer is 138 kDa, **Extended Data Fig. 5**). Then, we evaluated the effect of erythritol concentration on the regulation of eryD and the resulting reporter gene (sfGFP) expression (**Fig. 2c**). The data indicated that eryD could be dynamically regulated by a gradient concentration of erythritol (**Fig. 2d-f** and **Supplementary Fig. 19**). To develop eryD as a potential regulatory element for genetic circuit design in synthetic biology, we further constructed two erythritol induction systems by using the T7 RNA polymerase-based promoter (**Fig. 2g**) and the *E. coli* σ70 factor-based native promoters (**Fig. 2i**), respectively. Briefly, the eryD-binding site (29 bp) was inserted between each promoter and the RBS to see if the transcription and translation of the downstream sfGFP gene is regulated by eryD and erythritol (**Fig. 2g**, **i**). Overall, their regulatory effect on sfGFP expression was observed with different constructs, albeit the expression levels were various (**Fig. 2h**, **j** and **Supplementary Fig. 20**). Finally, the orthogonality of erythritol-eryD interaction was investigated with a series of carbohydrates and polyols. We found that eryD showed the highest orthogonality with erythritol, whereas the interactions of eryD with other tested compounds were very low (**Fig. 2k**, **l**).

### mRNA transcriptional analysis of the erythritol catabolism in engineered *E. coli*

To determine the erythritol catabolism in *E. coli*, we performed mRNA transcriptional analysis by comparing the exponential growth phase cultures of *E. coli* MG1655 between M9-erythritol medium and M9-glucose medium (**Extended Data Fig. 6**). In total, 1298 genes were significantly different between these two cultivations, including 759 genes upregulated and 539 genes downregulated (**Supplementary Fig. 21**). COG annotations showed that the significant differences were found in genes related to the metabolisms of carbohydrates, amino acids, energy, and inorganic ions, as well as the transcription and ribosome-associated translation (**Extended Data Fig. 7a**). In particular, most of the genes related to carbohydrate metabolism were overexpressed, suggesting that erythritol was not efficiently utilized as a carbon source compared to glucose. Of special note, nearly all ribosome-associated genes were downregulated, which indicates that the engineered strain decreased its overall translation level to adopt to a low-carbon-source environment. KEGG and GO annotations showed a similar trend/situation of gene expression as the COG analysis (**Extended Data Fig. 7b, c**).

Using transcriptomics data, we reconstructed the carbohydrate metabolism by connecting the heterologous erythritol degradation pathway to the native pentose-phosphate pathway, glycolysis, and TCA cycle (**Fig. 3**). In the hybrid metabolic pathways, erythrose-4-phosphate, the end product of erythritol catabolism, enters into pentose-phosphate pathway and glycolysis through the catalysis of two sets of enzymes, which are transaldolase (talA and talB) and transketolase (tktA and tktB). Transcriptome analysis showed that talA, talB, tktA, and tktB were overexpressed by 5.2, 2.0, 1.3, and 5.9 times, respectively, compared to those when cells grew in glucose-based M9 medium. In addition, several other carbohydrate degradation pathways were also upregulated, including D-xylose, L-arabinose, lactose, L-rhamnose, and glycerol. By contrast, only galactose and D-mannose degradation pathways were downregulated. In general, the gene expression levels in glycolysis and TCA cycle were not significantly impacted in the two cultivation conditions.

**Fig. 3.**
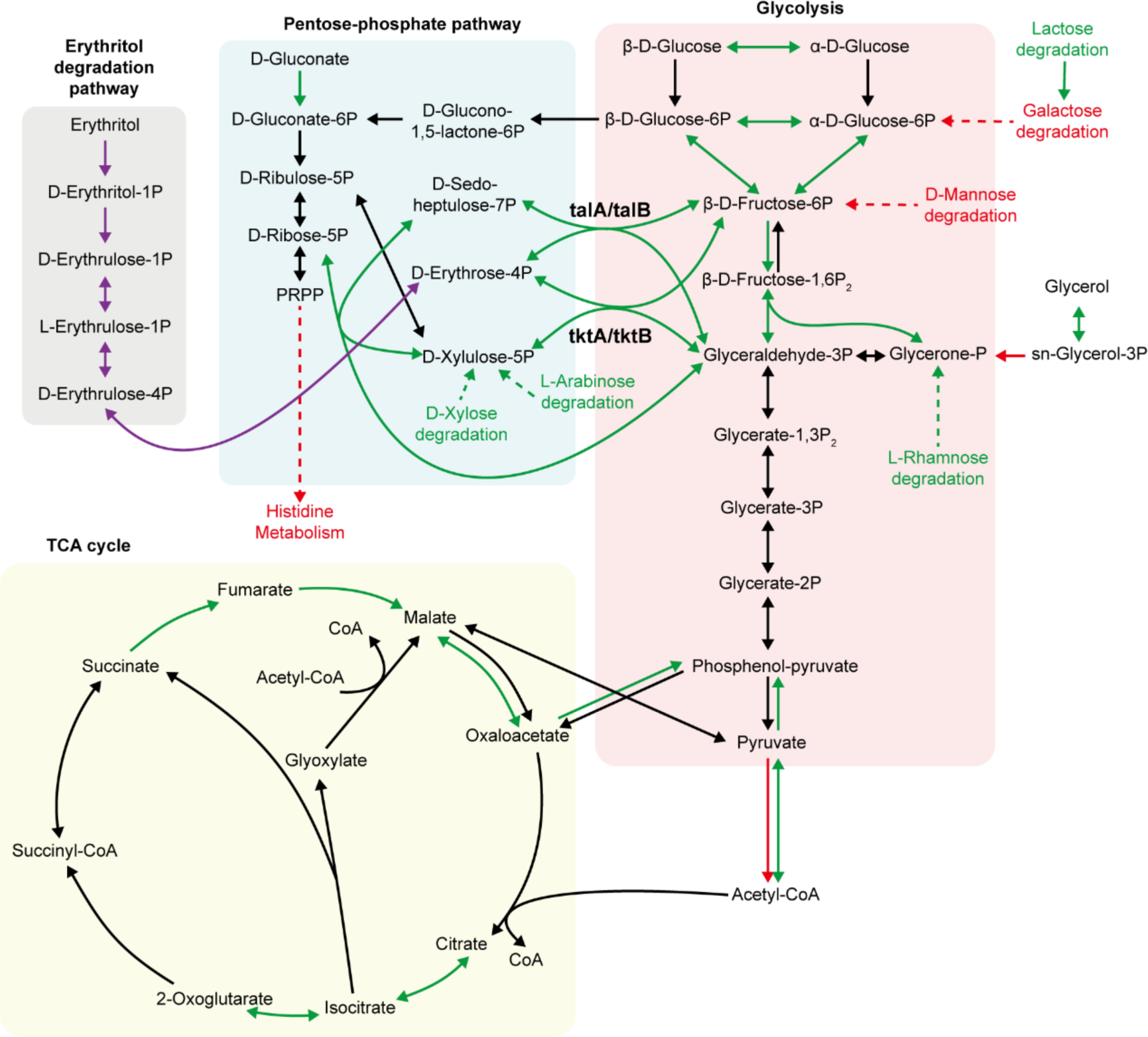
Reconstruction of the major carbohydrate metabolic pathways mapped with mRNA transcriptional analysis. Reconstructed major carbohydrate metabolic pathways of erythritol catabolic *E. coli* MG1655 (harboring pFB147) were based on the relative differential gene expression profiles between two cultures (M9-erythritol as the experimental group and M9-glucose as the control group). The genes with a significant (p < 0.05) differential expression of ≥ 1 log2(fold change) are indicated with arrows (pathway) in green (up-regulated in M9-erythritol medium) or red (down-regulated in M9-erythritol medium). The other genes without significant differential expression are indicated with black arrows (pathway). The details of mRNA transcriptional analysis are shown in the **Supplementary Information.**

By screening the gene expression level, some notably upregulated gene clusters were summarized (**Supplementary Fig. 22**). For example, the genes of glycolate utilization operon *glcCDEFGBA* were all significantly overexpressed (**Supplementary Fig. 22b**). However, no glycolate or analogs were in the M9-erythritol medium. We thus considered whether there is a transcriptional repressor in the *glc* cluster that might respond to erythritol catabolite(s). Previous work has reported that glcC is a DNA-binding transcriptional repressor that can respond to glycolate and acetate^41^. Here, our experimental data suggested that glcC could also respond to erythrose-4-phosphate or its downstream metabolite(s) (**Supplementary Fig. 23**). In addition, another gene cluster (i.e., *phn* operon^42^) involved in the phosphonate uptake and utilization was nearly all upregulated as well (**Supplementary Fig. 22e**). However, we observed that the transcriptional repressor phnP did not respond to erythritol catabolism in our experiments (data not shown). This is likely due to the requirement of more phosphonate to support erythritol phosphorylation, leading to overexpression of the *phn* operon.

### Metabolic engineering to improve cell growth based on erythritol metabolism

While we have demonstrated that engineered *E. coli* can use erythritol as sole carbon source, the utilization efficiency of erythritol was low (only 30% of added erythritol was consumed) and cells could not grow to a high density (OD_600_ reached 0.35 in 72 h) (**Fig. 1d**).

Next, we set out to increase cell growth relying on erythritol by metabolic engineering. To begin, we compared the effect of five constitutive promoters with a gradient strength and the native promoter (pEryF-732) on expression of the gene cluster and cell growth (**Fig. 4a**). We found that two medium strength promoters (J23106 and J23105) and pEryF-732 performed similar and the final OD_600_ values were comparable (**Fig. 4b**, **c**). Yet, the strongest promoter J23100 just could support cell growth to a medium level, which is lower than pEryF-732. Using the weakest promoter J23109, no cells could grow due to the low expression of the genes (**Fig. 4b**, **c**). Overall, our results suggested that a suitable expression level of the gene cluster was sufficient for erythritol catabolism and cell growth. Since the tested promoters did not show a significant increase on cell growth, we finally chose the native promoter (pEryF-732) to express the genes in our following experiments.

**Fig. 4.**
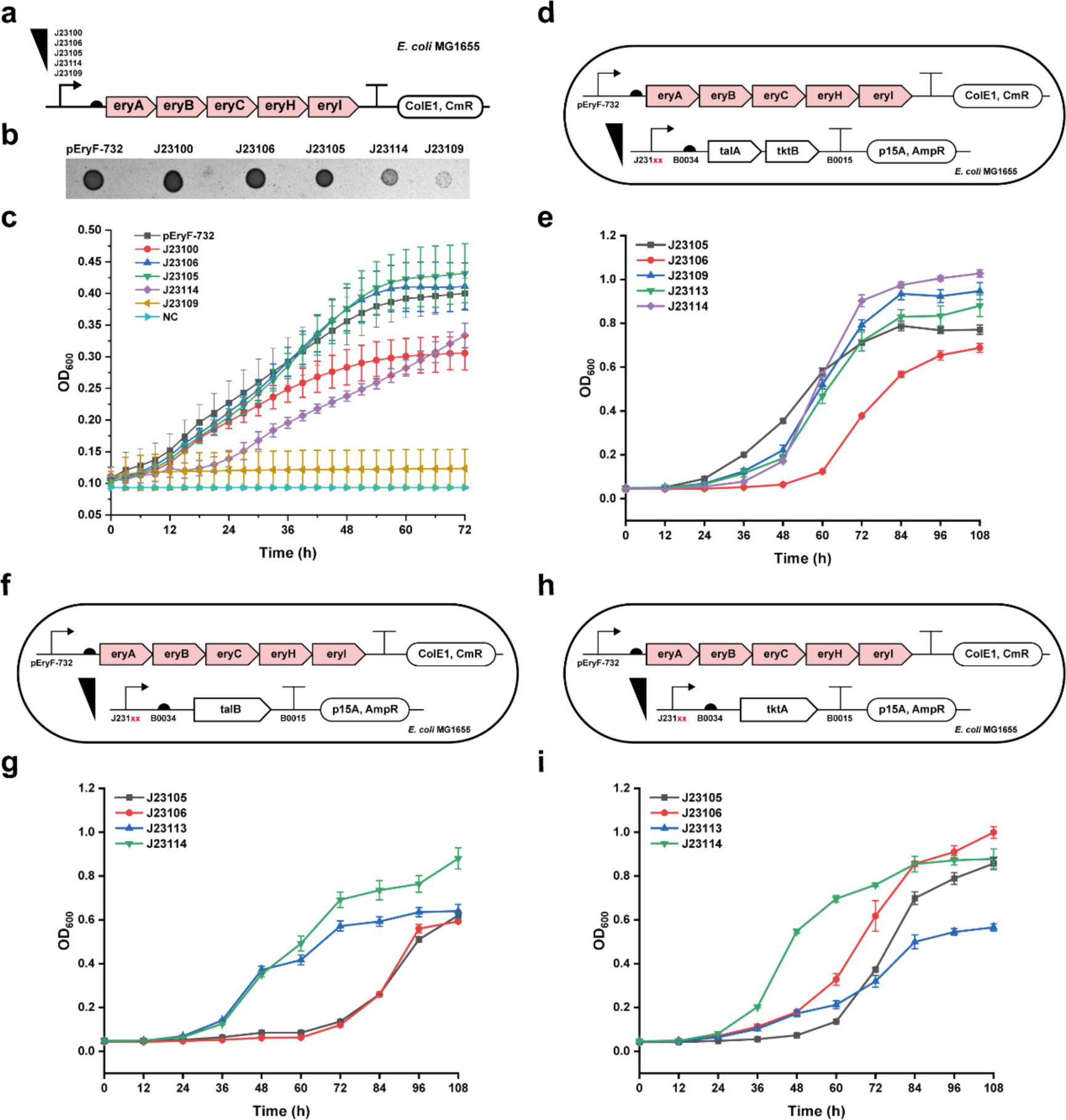
Improvement of erythritol catabolism in *E. coli* by metabolic engineering. (**a**) Schematic plasmids of pFB148 to pFB152. The native promoter pEryF-732 was replaced by a series of constitutive promoters with a gradient strength for erythritol metabolism optimization. (**b**) *E. coli* MG1655 harboring the above each plasmid grew on M9-erythritol agar plates after 72 h incubation at 37°C. (**c**) Growth curves of *E. coli* MG1655 harboring the above each plasmid in M9-erythritol liquid medium (5 mL, 37°C, 250 rpm). (**d**, **f**, **h**) Schematic shows *E. coli* MG1655 harboring the erythritol degradation plasmid pFB147 and other compatible plasmids for the coexpression of talA+tktB, talB, and tktA. (**e**, **g**, **i**) Cell growth curves by coexpression of the erythritol metabolic enzymes with talA+tktB, talB, or tktA, respectively. All the measurements were performed with three biological replicates. The error bars represent the standard deviation (s.d.).

On the basis of the transcriptome analysis, the expression of transaldolase (talA/talB) and transketolase (tktA/tktB) were upregulated to convert erythrose-4-phosphate into other metabolic pathways. We then hypothesized that overexpression of these genes might help increase the consumption of erythrose-4-phosphate to accelerate carbon flux and cell growth. To verify this, we constructed three compatible plasmids that were able to constitutively express talA+tktB, talB, and tktA, respectively (**Fig. 4d**, **f**, **h**). Note that the two genes of *talA* and *tktB* locate next to each other in the genome, thus we only constructed one plasmid to coexpress talA and tktB rather than two separate plasmids. By coexpression of the erythritol gene cluster and each of the above constructed plasmids, we obtained the profiles of cell growth after 108 h cultivation (**Fig. 4e**, **g**, **i**). In general, overexpression of talA+tktB, talB, or tktA could obviously increase cell growth with the final highest OD_600_ value of approximately 1.0 in each group, which is nearly 3-fold higher than that of the cultivation without overexpression (see **Fig. 4c** for the OD_600_ of 0.35).

### Characterization of the erythritol transport system in *E. coli*

Having demonstrated the utilization of erythritol in the engineered *E. coli* MG1655, we were curious which transport system(s) are responsible for erythritol transportation without expression of the erythritol ABC-transporter system (i.e., eryE, eryF, and eryG^22^). Since eryE and eryG are membrane proteins, we first demonstrated their transmembrane phenotype in *E. coli* Mach1-T1 cells (**Extended Data Fig. 4**). However, when the three proteins were coexpressed in other *E. coli* strains such as MG1655, we found that this ABC-transporter exhibited harmful impact on cell division and growth (**Supplementary Fig. 24b**, **d**). Therefore, we then only focused on the potential native genes in *E. coli* MG1655 that assist erythritol transportation.

First, we searched homologous genes of the erythritol ABC-transporter, including all carbohydrate or polyol transporter genes in *E. coli* MG1655 (**Extended Data Fig. 8**). After further protein-protein alignments by BLAST^43, 44^, seven permease genes that are homologous to *eryG* were screened out (**Extended Data Fig. 8c**), including the glycerol facilitator glpF, which has been reported to enable *E. coli* to transport erythritol^45^. Then, we individually knocked out the seven genes (**Extended Data Fig. 8d**) and expressed the erythritol metabolic genes in each of the seven knock-out strains. The results indicated that six transporter systems might facilitate erythritol transportation to support cell growth (**Extended Data Fig. 9**). In addition, we found 34 putative carbohydrate transport genes upregulated according to the transcriptome data, constructed 34 knock-out strains, and performed the same cultivation experiments. However, cell growth was not obviously impacted in each of the 34 knock-out strains, suggesting that these putative systems are not responsible for erythritol transportation (**Supplementary Fig. 25**). Taken together, while our preliminary data indicate that some endogenous carbohydrate transporter systems in *E. coli* might be able to transport erythritol, more characterization experiments should be performed to get more insights in the future study.

### *E. coli* as a living detector to distinguish soda drinks

In many soft drinks, some artificial and/or natural sweeteners are used as additives to reduce or replace the use of high-fructose corn syrup (including fructose and glucose) and sugar (sucrose). For instance, erythritol has been used as a natural sweetener in soda and other foods. Detection of such compound in food often relies on expensive instruments such as HPLC and GC-MS^46^. With the above engineered *E. coli* in hand, here we aim to use this strain as a living detector to detect erythritol from soda drinks. In principle, wild-type *E. coli* strains can utilize fructose and glucose as carbon sources, but not erythritol (**Supplementary Fig. 26**). Hence, the erythritol detection process is easy according to whether the cells can grow or not in the soda drinks-modified M9 medium. As shown in **Fig. 5a**, the *E. coli* strain (Erythritol Detector, ErD) containing the erythritol metabolic pathway is used for the detection. Meanwhile, a negative control (NC) strain is used for comparison, which harbors the same pathway but under the control of a weak promoter (J23109) that is not able to support cell growth on erythritol (**Fig. 4c**).

**Fig. 5.**
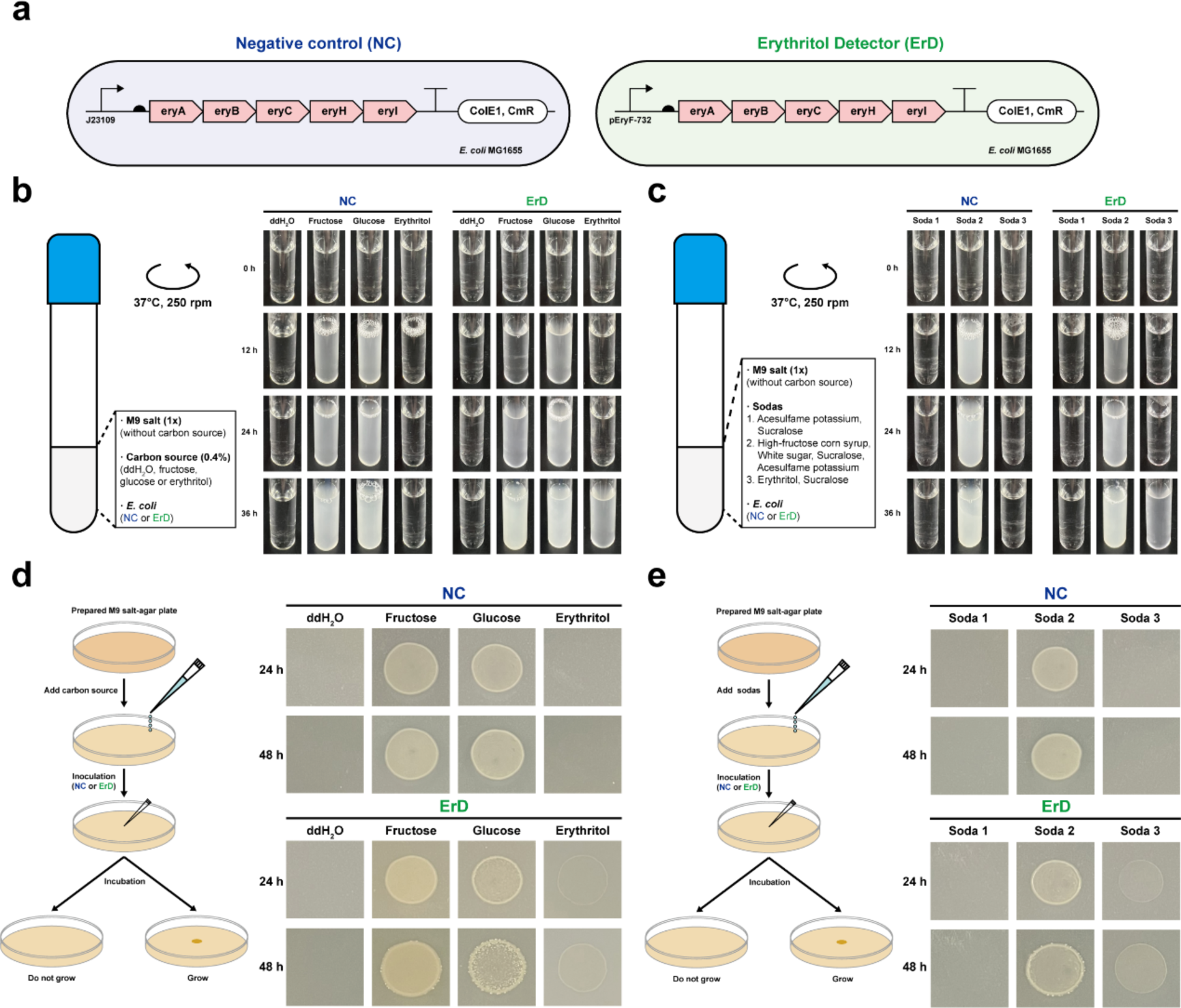
Engineered *E. coli* as a living detector to distinguish soda drinks. (**a**) Schematic of two *E. coli* MG1655 strains shows the negative control (NC, with pFB151 that cannot utilize erythritol) and the erythritol detector (ErD, with pFB147 that can utilize erythritol). (**b**, **c**) Liquid incubations were performed to distinguish synthetic M9-based carbohydrate media and soda drinks. (**d**, **e**) Solid incubations were performed for distinguishing synthetic M9-based carbohydrate media (the same as (**b**)) and soda drinks (the same as (**c**)). All samples were performed with three biological replicates showing similar results.

To prove the concept, we initially cultivated NC and ErD strains in M9 and M9-modified media, in which glucose (0.4%) is completely replaced with the same amount of ddH_2_O, fructose, or erythritol. In both groups (**Fig. 5b**), while NC and ErD cells were able to grow in fructose and glucose-based M9 media, no cell growth was observed without carbon source (ddH_2_O). Clearly, only ErD cells could grow in the erythritol-based cultivation, suggesting that our concept is feasible for detecting erythritol in soda drinks. Next, we obtained three commercial soda products (soda 1 is sugar-free and contains two artificial sweeteners: acesulfame potassium and sucralose; soda 2 contains fructose and glucose; soda 3 is sugar-free and contains erythritol and sucralose). Similarly, we modified the M9 medium by adding 10x diluted soda liquid to replace glucose. After cultivation for 36 h, we observed that the growth of ErD cells completely depended on the carbon sources from the selected soda drinks (**Fig. 5c**). There was no cell growth in soda 1 without any carbon source; the growth in soda 2 (containing fructose and glucose) was the fastest (the liquid became turbid in 12 h); by contrast, ErD could grow in soda 3 that contains erythritol but with a slow growth rate (36 h). As a result, the cell growth profile could basically tell if the soda drinks contain erythritol or not by using our ErD strain. Besides, we also cultivated cells on solid agar plates and obtained similar results (**Fig. 5d**, **e**). Of note, ErD cells/colonies became visible on the plates after 24 h incubation, a bit faster than the cultivation in liquid medium (36 h). Taken together, our living detector for erythritol is a potential method to distinguish erythritol-containing soda drinks or other foods with several properties of rapid operation, cheap detection, and easy visualization.

### Erythritol facilitates the growth of engineered *E. coli* Nissle 1917 in simulated intestinal fluid

Next, we aimed to bring the erythritol metabolic pathway to other *E. coli* strains for potential applications. In particular, we chose *E. coli* Nissle 1917 (EcN), which is a probiotic (generally recognized as safe, GRAS)^34^ and has been widely engineered and applied for diagnosis and therapy^47–50^. After transferring the erythritol gene cluster into EcN, we then sought to evaluate if the cells can grow in the simulated intestinal fluid (SIF) supplemented with erythritol. The incubation was carried out at 37°C for 4 days and at different time points the liquid cultures were spread to LB-agar plates for cell growth to count colony forming units (CFU) (**Fig. 6a**). First, we mixed SIF solution (0.4% erythritol) with different inoculated cells (10^2^, 10^4^, 10^6^, and 10^8^) and evaluated their growth. Overall, cells could grow by utilizing erythritol and the CFUs reached the highest after 2-day cultivation with the inoculation of 10^2^, 10^4^, and 10^6^ cells (**Fig. 6b-d**). Yet, the highest inoculation of 10^8^ cells led to a rapid increase of CFU in just 2-h cultivation, followed by a significant reduction of CFU within one day, which is likely due to the rapid exhaustion of nutrients by over-inoculated cells (**Fig. 6e**). Since the growth profiles between 10^4^ and 10^6^ were similar, we then decided to use a lower density of 10^4^ EcN cells to investigate the effect of erythritol concentration (0%, 0.01%, 0.1%, 1%, 4%, and 16%) on cell growth. The results showed that in general lower erythritol concentrations (0.01-4%) were beneficial to facilitate cell growth for 1-2 days (**Fig. 6g-j**). The growth of cells with the highest erythritol concentration (16%) was bad probably due to a high osmosis pressure in the culture (**Fig. 6k**). Unexpectedly, the negative control cultivation (SIF without erythritol) could also support cell growth with a maximum CFU of 2.8×10^9^ at the first day (**Fig. 6f**). This reason might be some residual erythritol existed in bile salts^51^, which is used to prepare SIF solutions. Taken together, engineered EcN cells are able to grow in SIF and supplemented erythritol can help significantly increase cell density. Yet, the inoculated cells and erythritol concentration are two factors subjected to be optimized in future potential applications.

**Fig. 6.**
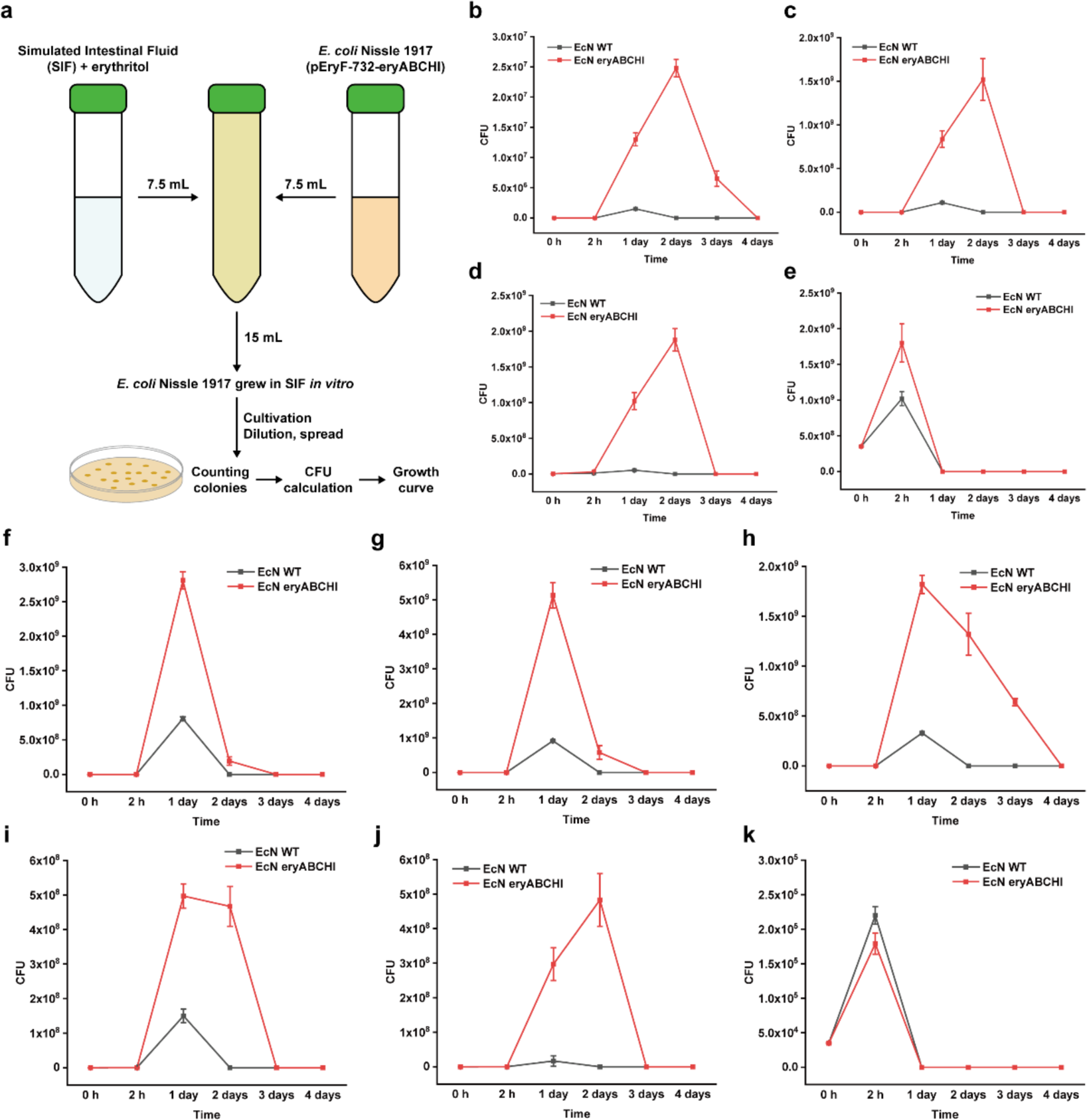
Erythritol facilitates the growth of engineered probiotic *E. coli* Nissle 1917 in SIF. (**a**) Schematic workflow of sample preparation, incubation, and measurement. The mixture was incubated at 37°C without shaking. (**b**-**e**) Growth curves with different inoculated cells of 10^2^, 10^4^, 10^6^, and 10^8^, respectively. Erythritol was added as 0.4% in SIF. (**f**-**k**) Growth curves with different erythritol concentrations of 0%, 0.01%, 0.1%, 1%, 4%, and 16% in SIF, respectively. All the initial inoculated cells were the same as 10^4^. “EcN WT” represents the wild-type *E. coli* Nissle 1917. “EcN eryABCHI” represents *E. coli* Nissle 1917 harboring pFB147 that can utilize erythritol. All the measurements were performed with three biological replicates. The error bars represent the standard deviation (s.d.).

## Discussion

Erythritol, a natural sweetener found/used in foods, cannot be metabolized by human bodies or the microbes in human gut^13, 15^, which is excreted by human as a carbon waste. Without an erythritol metabolism, most other microorganisms are not able to utilize erythritol either. In this study, we engineer *E. coli* strains by reconstitution of the erythritol metabolic pathway in a special effort to expand the carbon source scope that can be utilized by the *E. coli* chassis. The erythritol gene cluster(s) have been identified from several microbes^17, 19, 23, 25^. While *in vitro* characterization of the catalytic enzymes (eryA, eryB, eryC, eryH, and eryI) has been reported previously^17^, other regulatory proteins (eryD and eryR) and transporter proteins (eryE, eryF, and eryG) have not been investigated in detail with experimental evidence to our knowledge. Thus, we particularly investigate these proteins in this work in order to well employ the whole gene cluster for erythritol metabolism in a heterologous host (*E. coli*). A major finding of our study is the identification of the so far unknown DNA-binding site (29 bp sequence) of the transcriptional repressor eryD (**Fig. 2a**). This pair of eryD and DNA sequence (29 bp) can be further developed as an erythritol-responding repressor-operator system for metabolic engineering and synthetic biology applications. Recombinant expression of the transporter proteins in *E. coli*, however, makes cells either death (MG1655) or morphology changed (Nissle 1917) (**Supplementary Fig. 24b**, **d**). Interestingly, both *E. coli* strains are able to grow in pure erythritol-based M9 medium without coexpression of the erythritol transporter system, which suggests that *E. coli*’s native transporter(s) can work for the transportation. To figure out the potential transporter(s) in *E. coli*, we construct a total of 41 knock-out strains, yet only find six transporters showing positive effect on cell growth (**Extended Data Fig. 9**).

Importantly, different types of *E. coli* strains can be engineered to grow on erythritol as sole carbon source. This expands the scope of carbon nutrients from C6 (e.g., glucose) and C1 (e.g., CO_2_ and methanol) to C4-based erythritol that can be utilized by *E. coli*. Moreover, we find the expression of more than 750 genes is upregulated and most of the genes are related to carbohydrate transport and metabolism (**Extended Data Fig. 7a**). According to these overexpressed genes, we particularly select transaldolase (talA/talB) and transketolase (tktA/tktB), which can convert erythrose-4-phosphate into other pathways, for overexpression, leading to a significant improvement of cell growth (**Fig. 4**). This suggests that metabolic engineering strategies can further increase the efficiency of erythritol utilization and metabolism. With these engineered *E. coli* strains, we apply them to potential applications. First, *E. coli* MG1655 is used as a living erythritol detector for distinguishing soda drinks whether they contain erythritol or not. This method is cheap and easily visible compared to the instrument-based detection. Second, the probiotic *E. coli* Nissle 1917 (EcN) with erythritol metabolism is able to grow in the simulated intestinal fluid (SIF) supplemented with erythritol. This is promising and exciting because EcN might be developed as living therapeutics, which can respond to the human-safety compound erythritol, to help regulate/cure human gut microbiota-related diseases or even other focal tissues like tumors. While here two applications are showcased, we expect erythritol and its associated gene cluster to be used in more fields such as high cell-density cultivation (**Supplementary Fig. 28**) and recombinant protein expression (**Supplementary Fig. 29**), which are just two straightforward examples of using erythritol as carbon source.

Overall, we reconstitute the natural erythritol metabolic pathway in different *E. coli* hosts, which then show the capability to grow by using erythritol as sole carbon source. Looking forward, this success might be expanded to other heterologous hosts for more applications, if an erythritol-responding and/or utilizing system is needed, for instance in the areas of metabolic engineering, synthetic biology, and biomedical engineering.

## Methods

### Strains, vectors, plasmids, primers, and reagents

The details of *E. coli* strains, vectors, plasmids, and primers used in this study are listed in **Supplementary Tables 1, 4, 5,** and **6**, respectively. Some vectors and plasmids are derived from our previous work^52^. The full sequences of all plasmids are listed in **Supplementary Information** (Excel Sheet Data) and their correctness is verified by Sanger sequencing (GENEWIZ) unless otherwise noted. Q5 High-Fidelity DNA Polymerase (New England Biolabs), Phanta Super-Fidelity DNA Polymerase (Vazyme), FastPure Gel DNA Extraction Mini Kit (Vazyme), and ClonExpress Ultra One Step Cloning Kit (Vazyme) were used for molecular cloning. DreamTaq Green PCR Master Mix (Thermo Scientific) was used for colony PCR. Lysogeny Broth (LB) liquid medium contains 10 g tryptone, 5 g yeast extract, and 10 g sodium chloride in 1 L ddH_2_O. LB-agar plates were prepared by adding 15 g per liter LB. M9 minimal medium (also M9-glucose medium) contains 200 mL of 5× M9 salts (15 g/L KH_2_PO_4_, 2.5 g/L NaCl, 33.9 g/L Na2HPO4, and 5 g/L NH4Cl), 20 mL of glucose (20% w/v), 2 mL of 1 M MgSO_4_, and 100 μL of 1 M CaCl_2_ in 1 L ddH_2_O. M9-erythritol medium contains 200 mL of 5× M9 salts, 20 mL of erythritol (20% w/v), 2 mL of 1 M MgSO_4_, and 100 μL of 1 M CaCl_2_ in 1 L ddH_2_O. Antibiotic stocks (1000×) are 100 mg/mL ampicillin, 50 mg/mL kanamycin, and 34 mg/mL chloramphenicol.

### Genetic parts and gene clusters

All genetic parts (promoter, RBS, promoter-RBS, terminator, CDS, and protein binding site) and gene clusters used in this study are listed in **Supplementary Tables 2** and **3**. Genetic parts shorter than 100 bp were synthesized within the oligonucleotides (GENEWIZ) and inserted into PCR fragments during molecular cloning. Gene clusters were amplified from the isolated *Ochrobactrum* spp. genome and the referenced genome for DNA-DNA alignments was *Brucella anthropi* strain T16R-87 chromosome 2 (GenBank: CP044971.1). Gene clusters were checked for correctness with Sanger sequencing (GENEWIZ), which then were used as gene templates for PCR amplification.

### Plasmid construction

All plasmids were constructed by Gibson Assembly. In brief, all linear polymerase chain reaction (PCR) products (Q5 High-Fidelity DNA Polymerase or Phanta Super-Fidelity DNA Polymerase) were extracted by gel extraction and then assembled by using ClonExpress Ultra One Step Cloning Kit (Vazyme). After assembly, the reaction mixture was added to 50 μL competent *E. coli* Mach1-T1 cells for transformation, followed by incubation overnight on LB-agar plate. Then, DreamTaq Green PCR Master Mix (Thermo Scientific) was used for colony PCR. The PCR products were sequenced by GENEWIZ.

### Western-Blot analysis

*E. coli* pellets were collected in 1.5 mL centrifuge tubes at 5000 g and 4°C for 10 min. The supernatant was discarded and the pellets were resuspended with 1 mL 1× phosphate-buffered saline (pH 7.4) and lysed by sonication (Q125 sonicator, Qsonica, 10 s on/off, 50% of amplitude, input energy ∼600 Joules). The lysate was then centrifuged at 12000 g and 4°C for 10 min. The total (cell lysate, T), soluble (supernatant, S), and pellet (P) fractions were separated by SDS-PAGE or Native-PAGE (Omni-Easy One-Step PAGE Gel Fast Preparation Kit, EpiZyme), followed by wet transferring to PVDF membrane (Bio-Rad) with 1× transfer buffer (25 mM Tris-HCl, 192 mM glycine and 20% v/v methanol in 1 L ddH_2_O, pH 8.3). Then, the PVDF membrane was blocked (Protein Free Rapid Blocking Buffer, Epizyme) for 1 h at room temperature. After washing thrice with TBST for each 5 min, 1:10000 (TBST buffer-based) diluted His-Tag Mouse Monoclonal Antibody (Proteintech) solution was added to the membrane and incubated for 1 h at room temperature. After washing thrice with TBST for each 5 min, 1:10000 (TBST buffer-based) diluted HRP-Goat Anti-Mouse IgG (H+L) Antibody (Proteintech) solution was added to the membrane and incubated for another 1 h at room temperature. After the last washing with TBST thrice for each 5 min, the membrane was visualized using Omni ECL reagent (EpiZyme) under UVP ChemStudio (analytikjena).

### *Ochrobactrum* spp. separation, identification, and characterization

M9-erythritol medium (25 mL) was added to a 50 mL centrifuge tube and the lid was replaced by gauze to block aerosol dust and enable microorganisms to drop into the medium. This device was stationary-placed outdoors at approximately 25°C for about 7 days. On the seventh day, the medium became turbid and then the mixture was spread onto an M9-erythritol agar plate for picking single colonies at 25°C. After three days, some white colonies were formed and picked up for identification. 16s rRNA was sequenced by primer pair 27F/1492R. Cell morphology was characterized by scanning electron microscope (SEM, JSM-7800F Prime) and transmission electron microscope (TEM, JEM-2100 Plus). *Ochrobactrum* spp. genome sequence was referred to the *Brucella anthropi* strain T16R-87 chromosomes 1 and 2 (GenBank: CP044970.1 and CP044971.1).

### Incubation of erythritol catabolic *E. coli* in M9-erythritol liquid medium

All engineered *E. coli* strains that could utilize erythritol as sole carbon source were cultured and followed the same procedure. Cell pellets from overnight LB culture were collected in 1.5 mL centrifuge tubes at 5000 g and 4°C for 10 min. The supernatant was discarded and the pellets were washed thrice by M9-erythritol medium, then the pellets were resuspended with M9-erythritol and diluted to OD_600_=1.0 for standardization. Then, the resuspended mixture was inoculated into new M9-erythritol medium (100 mL medium in a 250 mL conical flask or 1 L medium in a 2 L conical flask) as 1:500 (v/v) for the following incubation. All antibiotic concentrations in M9-erythritol medium were diluted by five times to avoid growth inhibition in the minimal medium as far as possible.

### HPLC analysis of erythritol

1 mL M9-erythritol medium (with *E. coli*) cell cultures were collected in 1.5 mL centrifuge tubes at 5000 g and 4°C for 10 min. The supernatants were then passed through a 0.22 μm filter to remove cell pellets and other precipitation. The resulting samples were used for HPLC analysis (see **Supplementary Fig. 30** for a standard curve of erythritol).

The 1260 Infinity II Prime HPLC System (Aligent) with a Hi-Plex H column (Aligent) was used for erythritol analysis. The experiment parameters were set as follows: column temperature: 60°C, mobile phase: 5 mM H_2_SO_4_ solution, flow rate: 0.6 mL/min, detector: Refractive Index Detector (RID) at 40°C, sample loading volume: 2 μL, testing time: 20 min for each sample.

### Characterization of eryD binding site *in vivo*

eryD binding site (eryO) was characterized and located in pEryF-732, which was a 732 bp gap between eryE (5’ start) and eryA (3’ end). In general, the characterization was performed as an *in vivo* incremental truncation process in two steps.

First, to determine the 5’ site, a series of 100 bp-truncated variant plasmids (pFB186 to pFB191, pFB210) were constructed and characterized that eryO was located between pEryF-232 to pEryF-132. Second, a series of 10 bp-truncated variant plasmids (pFB192 to pFB198, pFB208, and pFB209) were constructed and characterized that eryO was located between pEryF-162 to pEryF-152. Third, a series of 1 bp-truncated variant plasmids (pFB202 to pFB207) were constructed and characterized that eryO 5’ site was located in pEryF-159.

To determine the 3’ site, the truncation process was the same as described above. The constructed plasmids (pFB214 to pFB238) were characterized step-by-step. Finally, eryO 3’ site was located in pEryF-131. The exact eryO site was characterized as a 29 bp DNA sequence of 5’-gaaaaaaaatgcgccatctagaaaatttt-3’.

### Electrophoretic mobility shift assay (EMSA)

DNA fragment containing the 732 bp eryE-eryA intergenic region (pEryF-732) was amplified with the 5’FAM-labeled primers from the plasmid pFB147 and then gel purified. The DNA fragment (200 nM) was incubated with different concentrations of purified eryD in the binding buffer (10 mM Tris-HCl pH 7.5, 1 mM EDTA, 100 mM NaCl, 0.1 mM DTT, 10 μg/mL BSA, 5% glycerol). The 20 μL mixtures were incubated for 30 min at room temperature and then mixed with 5 μL of the same buffer supplemented with 50% glycerol and bromophenol blue. Free DNA and eryD-DNA complexes were separated on 7.5% polyacrylamide gels in 1x TAE buffer (4.844 g/L Tris-base, 1.21 mL/L acetate acid, 0.372 g/L EDTA disodium salt dihydrate, pH 8.3). The fluorescence of FAM-labeled bands was visualized under UVP ChemStudio (analytikjena).

### Characterization of the eryD orthogonality

*E. coli* Mach1-T1 with plasmids pFB186 and pFB261 was used for the characterization. Inducers (erythritol, glycerol, L-arabinose, D-xylose, D-glucose, D-mannose, D-galactose, D-fructose, L-sorbose, L-rhamnose, D-sucrose, lactose, maltodextrin, D-sorbitol, and D-mannitol) were dissolved in ddH_2_O at different concentrations (5%, 1%, 0.1%, and 0.01%, w/v), followed by filtration through a 0.22 μm filter.

*E. coli* Mach1-T1 was initially transformed with the relevant plasmids. Starter cultures (LB containing 50 μg/mL ampicillin and 17 μg/mL chloramphenicol) were inoculated from a single colony and grew overnight at 37°C for 16 h. The next day, 10 μL starter culture was used to inoculate 5 mL LB medium containing 50 μg/mL ampicillin and 17 μg/mL chloramphenicol in test tubes. The cultures were incubated at 37°C with shaking (250 rpm) until OD_600_ reached 0.6-0.8. The cultures were then added with different inducer solutions and incubated for another 16 h at 37°C and 250 rpm.

Then, cell pellets were collected in 1.5 mL centrifuge tubes at 5000 g and 4°C for 10 min. The supernatant was discarded and the pellets were resuspended with 1 mL 1× phosphate-buffered saline (pH 7.4). The sfGFP fluorescence of resuspended mixture was measured by microplate reader (Synergy H1, BioTek) and performed with excitation and emission wavelengths at 485 and 528 nm, respectively. All measurements were performed at least in triplicate. In addition, cell pellets were lysed by sonication (Q125 sonicator, Qsonica, 10 s on/off, 50% of amplitude, input energy ∼600 Joules). The lysate was then centrifuged at 12000 g and 4°C for 10 min. The total fraction was separated and analyzed by Western-Blot.

### eryD structure modeling

*Ochrobactrum* spp. eryD amino acid sequence was the same as eryD (Uniprot ID: Q2YIQ4) from *Brucella abortus* (strain 2308). AlphaFold, Robetta, and SWISS-MODEL were used for structure modeling.

### Size exclusion chromatography

To determine the native molecular weight and assembly model of eryD, size exclusion chromatography was performed using a Superdex 200 Increase 10/300 GL column (GE Healthcare) and two reference proteins including Vlm2 (284 kDa)^53^ and T7 RNA polymerase (99 kDa) were used for comparison.

### Characterization of transmembrane phenotype of the erythritol ABC-transporter system

TMHMM-2.0 was used for transmembrane prediction. eryE was predicted as a single transmembrane protein and eryG was predicted as a ten-times transmembrane protein. To determine the transmembrane phenotypes, pFB162 and pFB164 were constructed for eryE-sfGFP and eryG-sfGFP expression. After transformation into *E. coli* Mach1-T1, starter cultures (LB containing 34 μg/mL chloramphenicol) were inoculated from a single colony and grew overnight at 37°C for 16 h. The next day, 10 μL starter culture was used to inoculate 5 mL LB medium containing 34 μg/mL chloramphenicol in test tubes. The cultures were incubated at 37°C with shaking (250 rpm) until OD_600_ reached 0.6-0.8. The cultures were then added 1% arabinose (w/v) and incubated for another 6 h at 30°C and 250 rpm. Afterward, cell pellets from 1 mL cell culture were collected in 1.5 mL centrifuge tubes at 5000 g and 4°C for 10 min. The supernatant was discarded and the pellets were resuspended with 1 mL 1× phosphate-buffered saline (pH 7.4). Then, the mixture was imaged under laser scanning confocal microscopy (FV3000, Olympus).

### Protein purification

Plasmids pFB161 and pFB167 were used to express eryD and eryR, respectively. The two proteins were expressed and purified with the same protocol. *E. coli* BL21(DE3) was transformed with the above plasmids, respectively. Starter cultures (LB containing 100 μg/mL ampicillin) were inoculated from a single colony and grew overnight at 37°C for 16 h. 5 mL starter culture was used to inoculate 1 L LB medium containing 100 μg/mL ampicillin in a 2 L conical flask. The cultures were incubated at 37°C with shaking (250 rpm) until OD_600_ reached 0.6-0.8. The cultures were then quickly cooled down on ice rapidly to 20°C, and isopropyl-β-D-thiogalactopyranoside (IPTG) was added for induction at a final concentration of 0.5 mM. The cultures then grew for another 16 h at 20°C and 220 rpm. Afterward, cells were harvested by centrifugation (Avanti JXN-26 High-Speed Centrifuge, Beckman Coulter) at 5000 g and 4°C for 10 min. The pellets were resuspended in Buffer A, containing 20 mM sodium phosphate (pH 7.4), 1 M sodium chloride, 1 mM dithiothreitol (DTT), and 50 mM imidazole. The suspension was cooled on ice and then lysed thrice at 1500 bar by ultra-high-pressure homogenization (JNBIO). The lysate was then centrifuged at 4°C and 20000 g for 30 min. The supernatant was collected and passed through a 0.22 μm filter. The filtered solution was then purified by Ni^2+^ affinity chromatography using a 1 mL HisTrap FF column (GE Healthcare). The column was equilibrated with 25 mL Buffer A at a constant flow rate of 1 mL/min. After equilibration, the filtered protein solution was loaded, followed by washing with 25 mL Buffer A. Then, bounded proteins were eluted with Buffer B (Buffer A, but with 500 mM imidazole) and collected into 1.5 mL centrifuge tubes with 0.5 mL elution. Each 0.5 mL elution was analyzed by SDS-PAGE. Eluted protein samples were combined in one tube and desalinated using an ultrafiltration tube (Amicon Ultra 3 kDa molecular weight cut-off, Merck/Millipore) with Buffer C (25 mM Tris-HCl (pH 7.5), 1 mM DTT, and 1 M sodium chloride). The desalinated and concentrated protein solution was then mixed with 40% glycerol (v/v in water) with a volume ratio of 1:1. The concentration of the final protein solution was measured at 280 nm after molar attenuation coefficient correction. Purified proteins were flash frozen by liquid nitrogen and stored at -80°C until further use.

### Protein expression and solubility characterization

Plasmids pFB158 to pFB167 were used to express ten erythritol catabolism-associated genes. *E. coli* strains were transformed with the relevant plasmids. Starter cultures (LB containing 100 μg/mL ampicillin or 34 μg/mL chloramphenicol) were inoculated from a single colony and grew overnight at 37°C for 16 h. The next day, 10 μL starter culture was used to inoculate 5 mL LB medium containing 100 μg/mL ampicillin or 34 μg/mL chloramphenicol in test tubes. The cultures were incubated at 37°C with shaking (250 rpm) until OD_600_ reached 0.6-0.8. The cultures were then added with inducers (1% arabinose (w/v) or 0.5 mM IPTG) and incubated continuously. Afterward, cell pellets were collected in 1.5 mL centrifuge tubes at 5000 g and 4°C for 10 min. The supernatant was discarded and the pellets were resuspended with 1 mL 1× phosphate-buffered saline (pH 7.4) and lysed by sonication (Q125 sonicator, Qsonica, 10 s on/off, 50% of amplitude, input energy ∼600 Joules). The lysate was then centrifuged at 12000 g and 4°C for 10 min. The total (cell lysate, T), soluble (supernatant, S), and pellet (P) fractions were separated and analyzed by SDS-PAGE and Western-Blot.

### Fluorescence measurement standardization

sfGFP (C-terminal 6×His) was used as a fluorescence reporter. pFB286 to pFB291 were performed as standardized sfGFP expression plasmids. These plasmids contained four parts within iGEM standardized vector pSB1C3: *E. coli* constitutive promoter library (J23100, J23106, J23105, J23114, J23109, and J23113, from strong to weak), strong ribosome binding site B0034, sfGFP, and a strong terminator B0015. Other plasmids containing sfGFP were compared to these six standardized plasmids to determine the sfGFP expression level. sfGFP fluorescence was measured by microplate reader (Synergy H1, BioTek) and performed with excitation and emission wavelengths at 485 and 528 nm, respectively. All measurements were performed at least in triplicate.

### Knocking out *E. coli* genes

Gene knock-out strategy was derived from Lambda-Red recombination. In brief, plasmids pFB278 to pFB284 and pFB306 to pFB339 were constructed to obtain linear PCR products. The descriptions “homoL” and “homoR” of these plasmids represented “homologous left arm” and “homologous right arm” between the target knock-out gene or gene cluster. These two DNA sequences were designed approximately longer than 300 bp for a higher recombination efficiency. The linear PCR products (homoL - FRT - KanR - FRT - homoR) were obtained and purified for use.

Wild-type *E. coli* MG1655 was firstly transformed with pKD46. On the next day, a single colony was picked up and inoculated in LB medium (with 1% arabinose, w/v) and incubated at 30°C and 250 rpm for approximately 4-6 h. When OD_600_ reached 0.6-0.8, the cell pellets were collected in 1.5 mL centrifuge tubes at 5000 g and 4°C for 10 min. The supernatant was discarded and the pellets were washed thrice with 4°C ddH_2_O. Then, 1 μg purified linear PCR products were added to the cell mixture and proceeded electro-transformation (Gene Pulser Xcell system (Bio-Rad) and Gene Pulser/Micropulser Electroporation Cuvettes, 0.1 cm gap (Bio-Rad) were used). Subsequently, colony PCR (primer pair: “homoL” forward primer / “homoR” reverse primer) was performed to test whether the gene or gene cluster deleted. Finally, the expected knock-out *E. coli* MG1655 strains were continuously incubated for days at 37°C for pKD46 (temperature-sensing plasmid) elimination.

### mRNA transcriptome analysis

To process transcriptome analysis, biological triplicate of *E. coli* MG1655 (with plasmid pFB147) in both M9-glucose medium (control group) and M9-erythritol medium (experimental group) were performed.

Cell pellets from overnight LB culture (37°C, 250 rpm for 16 h) were collected in 1.5 mL centrifuge tubes at 5000 g and 4°C for 10 min. The supernatant was discarded and the pellets were washed thrice by M9-glucose or M9-erythritol medium. Then, the pellets were resuspended and diluted to OD_600_=1.0 for standardization. Afterward, the suspensions were inoculated into new M9-glucose or M9-erythritol medium (1 L medium in a 2 L conical flask) as 1:500 v/v for the following incubation.

For M9-glucose cultivation, when OD_600_ reached 0.6 (mid-log phase, approximately 6 h after inoculation), cell pellets were collected. For M9-erythritol cultivation, when OD_600_ reached 0.15 (mid-log phase, approximately 36 h after inoculation), cell pellets were collected. Then, the cell pellets were sent to GENEWIZ for further mRNA transcriptome analysis.

### Distinguishing soda drinks

*E. coli* MG1655 harboring pFB151 and pFB147 were used as negative control (NC) and living erythritol detector (ErD), respectively. Three kinds of sweet soda drinks were bought from the market. The ingredients of soda 1 contain water, carbon dioxide, citric acid, potassium citrate, acesulfame potassium, sodium benzoate, sucralose, and sodium gluconate. Soda 2 contains water, high-fructose corn syrup (including fructose and glucose), sugar (sucrose), carbon dioxide, citric acid, potassium citrate, sodium benzoate, sucralose, and acesulfame potassium. Soda 3 contains water, erythritol, carbon dioxide, citric acid, potassium citrate, sodium bicarbonate, and sucralose. All sodas were passed through a 0.22 μm filter to remove impurities and then shaken to eliminate carbon dioxide.

For liquid incubation, the sodas were diluted by 10 times in ddH_2_O to decrease the ingredient concentration, and the diluted sodas were used as different liquid mediums. NC and ErD cell pellets from overnight LB culture (37°C, 250 rpm for 16 h) were collected in 1.5 mL centrifuge tubes at 5000 g and 4°C for 10 min. The supernatant was discarded and the pellets were washed thrice by ddH_2_O. Then, the pellets were resuspended and diluted to OD_600_=1.0 for standardization. Then, 10 μL mixtures were inoculated into 5 mL diluted sodas and incubated at 37°C and 250 rpm for 12 h, 24 h, and 36 h, respectively. The solutions were imaged at each time point.

For solid incubation, M9 agar plates were prepared (10 mL mixture in 10 mm plastic plates) for cell cultivation. 1 mL of filtered sodas (without dilution) were added and spread on the solid medium and waited for drying. Then, 10 μL NC or ErD cell mixtures were inoculated onto the solid medium and incubated at 37°C for 24 h and 48 h, respectively. The plates were imaged at each time point.

### Engineered *E. coli* Nissle 1917 grows in simulated intestinal fluid

The probiotic *E. coli* Nissle 1917 (EcN, Mutaflor) was transformed with plasmid pFB147. Then, EcN cells were diluted in ddH_2_O with a gradient density (2×10^2^, 2×10^4^, 2×10^6^, and 2×10^8^) for the following experiments. 2× simulated intestinal fluid (SIF) was prepared according to a standard protocol with additional erythritol (0%, 0.02%, 0.2%, 2%, 8%, and 32%, w/v)^54, 55^. The SIF solutions were passed through a 0.22 μM filter. Then, 7.5 mL EcN culture and 7.5 mL 2× SIF were mixed as 1:1 volume to obtain a 15 mL culture. Then, the cultures were stationary-placed for incubation for days at 37°C. At each time point, approximately 100 μL samples were taken and spread (diluted if necessary) on LB-agar plates for counting colony forming units (CFU). All experiments were performed at least in triplicate.

## Acknowledgements

This work was supported by grants from the National Natural Science Foundation of China (Nos. 31971348 and 32171427) and the Double First-Class Initiative Fund of ShanghaiTech University (No. SYLDX0292022).

## Author contributions

J. L. and F. B. designed the experiments. F. B. performed all experiments. X. J. helped perform cell cultivation. S. H. and W. Q. L. performed HPLC analysis. Y. Z. helped perform molecular cloning. F. B. analyzed the data and drafted the manuscript. J. L., S. L., and Y. L. revised and edited the manuscript. J. L. conceived and supervised the study. All authors read and approved the final manuscript.

## Competing interests

The authors declare no competing interests.

## Extended Data Figures

**Extended Data Fig. 1.**
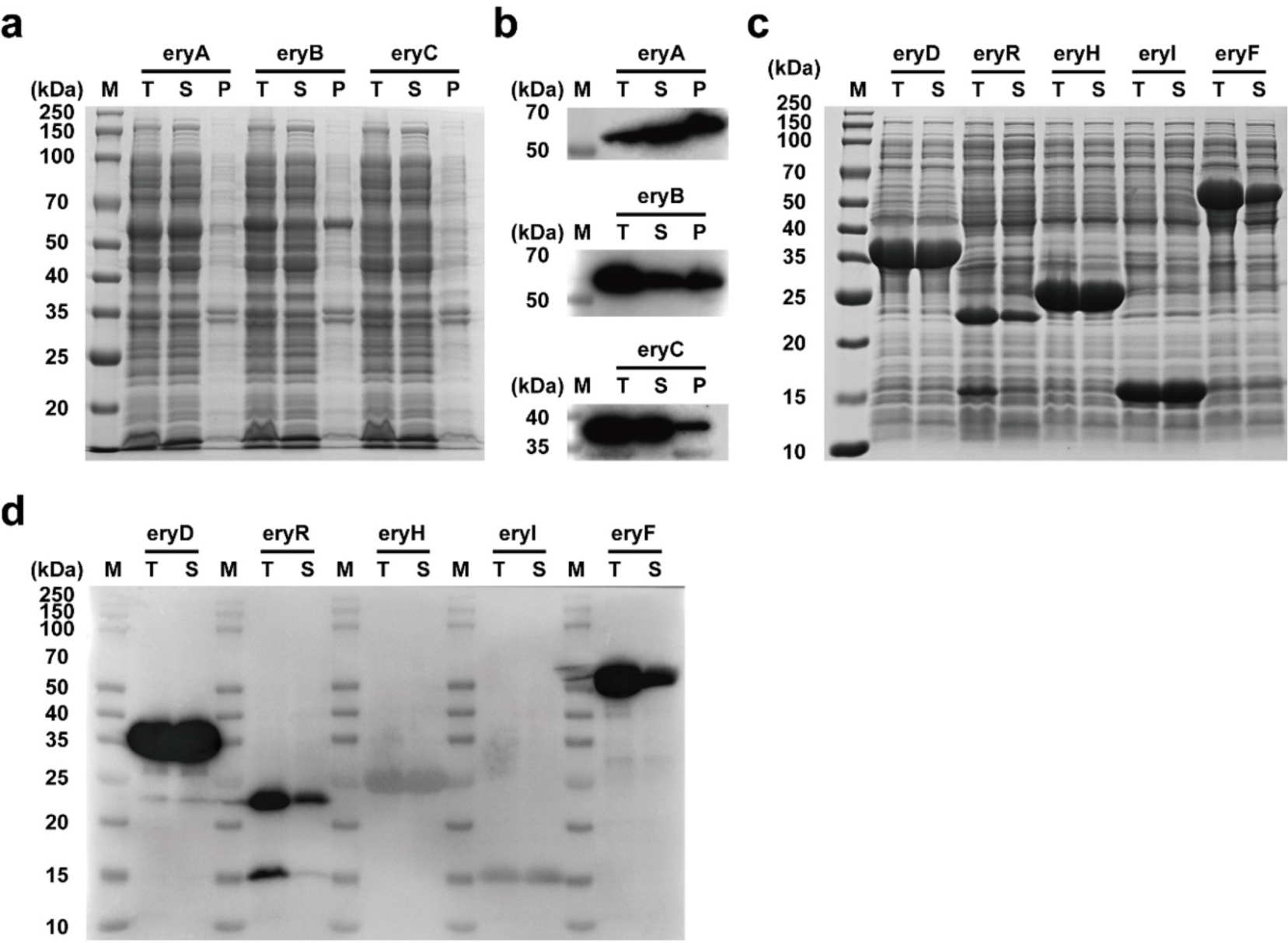
*In vivo* expression of erythritol catabolism-associated genes. (**a**) SDS-PAGE analysis of gene expression of eryA (with N-terminal 6xHis tag, 56.2 kDa), eryB (with N-terminal 6xHis tag, 57.2 kDa), and eryC (with N-terminal 6xHis tag, 35.8 kDa). All these three genes were expressed by plasmids pFB158, pFB159, and pFB160, respectively. *E. coli* Mach1-T1 harboring each above plasmid was initially inoculated with overnight culture (5 mL LB at 37°C and 250 rpm for 16 h) as a volume ratio of 1:500, and then incubated in 5 mL new LB medium at 37°C and 250 rpm. When OD_600_ reached 0.6, 1% arabinose (w/v) was added and incubated for another 16 h at 20°C and 250 rpm. Then, cell pellets were resuspended (equal volume) with 1x phosphate-buffered saline (pH 7.4) and lysed by sonication. The total cell lysate “T”, supernatant “S”, and pellets “P” were analyzed by SDS-PAGE. (**b**) Western-Blot analysis of the same samples in (**a**). (**c**) SDS-PAGE analysis of gene expression of eryD (with N-terminal 6xHis tag, 34.5 kDa), eryR (with N-terminal 6xHis tag, 24.8 kDa), eryH (with N-terminal 6xHis tag, 28.8 kDa), eryI (with N-terminal 6xHis tag, 16.6 kDa), and eryF (with N-terminal 6xHis tag, 56.2 kDa). All these five genes were expressed by plasmids pFB161, pFB167, pFB165, pFB166, and pFB163, respectively. *E. coli* BL21(DE3) harboring each above plasmid was initially inoculated with overnight culture (5 mL LB at 37°C and 250 rpm for 16 h) as a volume ratio of 1:500, and then incubated in 5 mL new LB medium at 37°C and 250 rpm. When OD_600_ reached 0.6, 0.5 mM IPTG was added and incubated for another 16 h at 20°C and 250 rpm. Then, cell pellets were resuspended (equal volume) with 1x phosphate-buffered saline (pH 7.4) and lysed by sonication. The total cell lysate “T”, supernatant “S”, and pellets “P” were analyzed by SDS-PAGE. (**d**) Western-Blot analysis of the same samples in (**c**).

**Extended Data Fig. 2.**
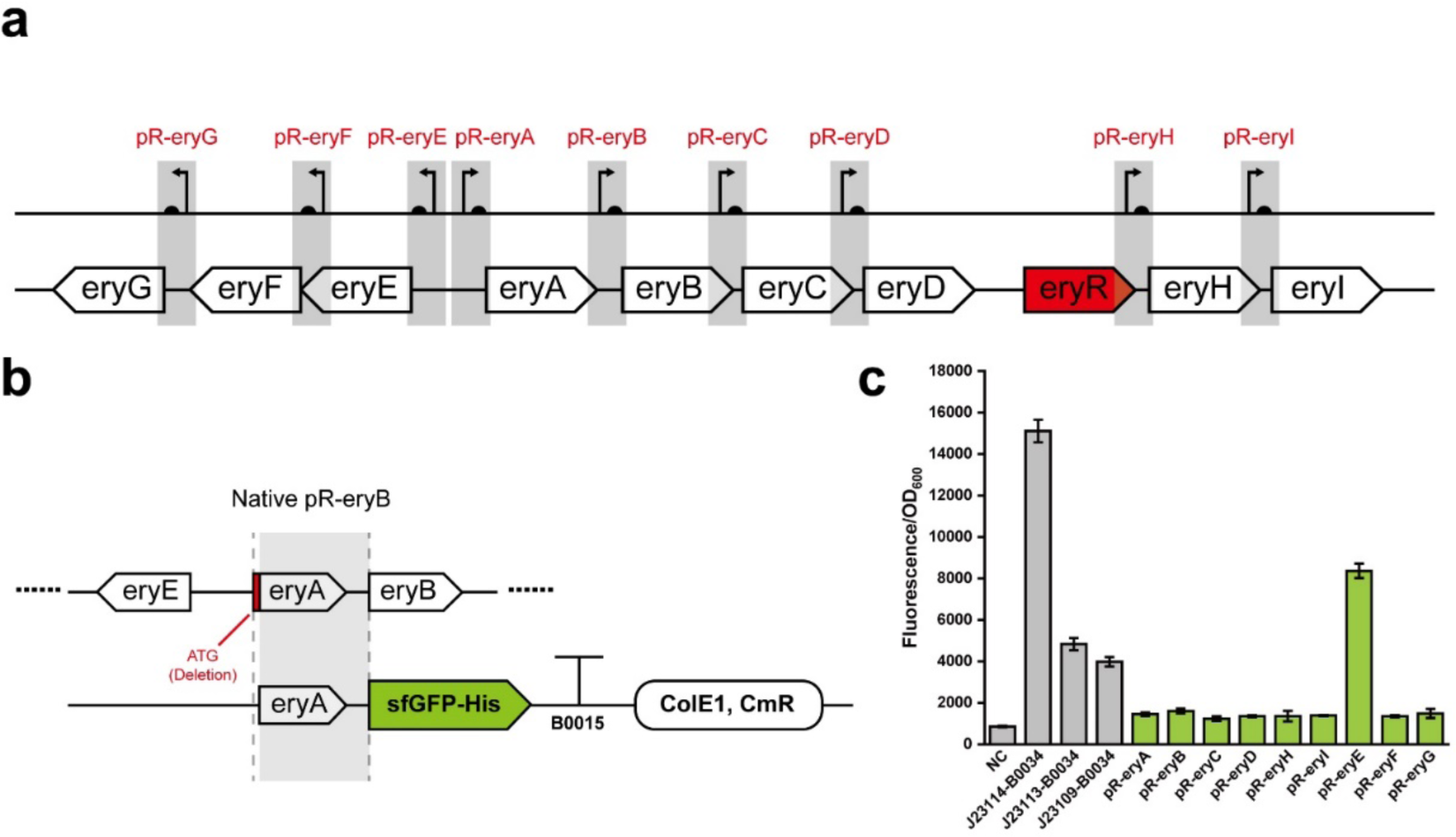
Native promoter-RBS (pR-) composite part characterization in the erythritol gene cluster. (**a**) Schematic of promoter-RBS (pR-) composite part localization. Note that the function of eryR was not successfully characterized, thus here we did not test pR-eryR. (**b**) Schematic procedure of “pR-ery” reporter plasmid construction (pFB177 to pFB185). Here we illustrate the construction of pR-eryB-sfGFP reporter plasmid as an example. The DNA sequence between “eryA CDS except the initiation codon ATG” and “the last nucleotide before eryB CDS” was considered as pR-eryB. This part was then assembled with the following sfGFP (6xHis) and a terminator B0015 in the vector pSB1C3 (pFB178). (**c**) Normalized fluorescence of all nine “pR-ery” parts. All the measurements were performed in *E. coli* Mach1-T1 with three biological replicates. “NC” represents *E. coli* Mach1-T1 without plasmids. The other three gray columns correspond to pFB291, pFB290, and pFB289, respectively, to act as positive controls (iGEM standard biological parts). The results indicate that all nine native composite parts are not strong regulatory elements compared to the reference “J23114-B0034”.

**Extended Data Fig. 3.**
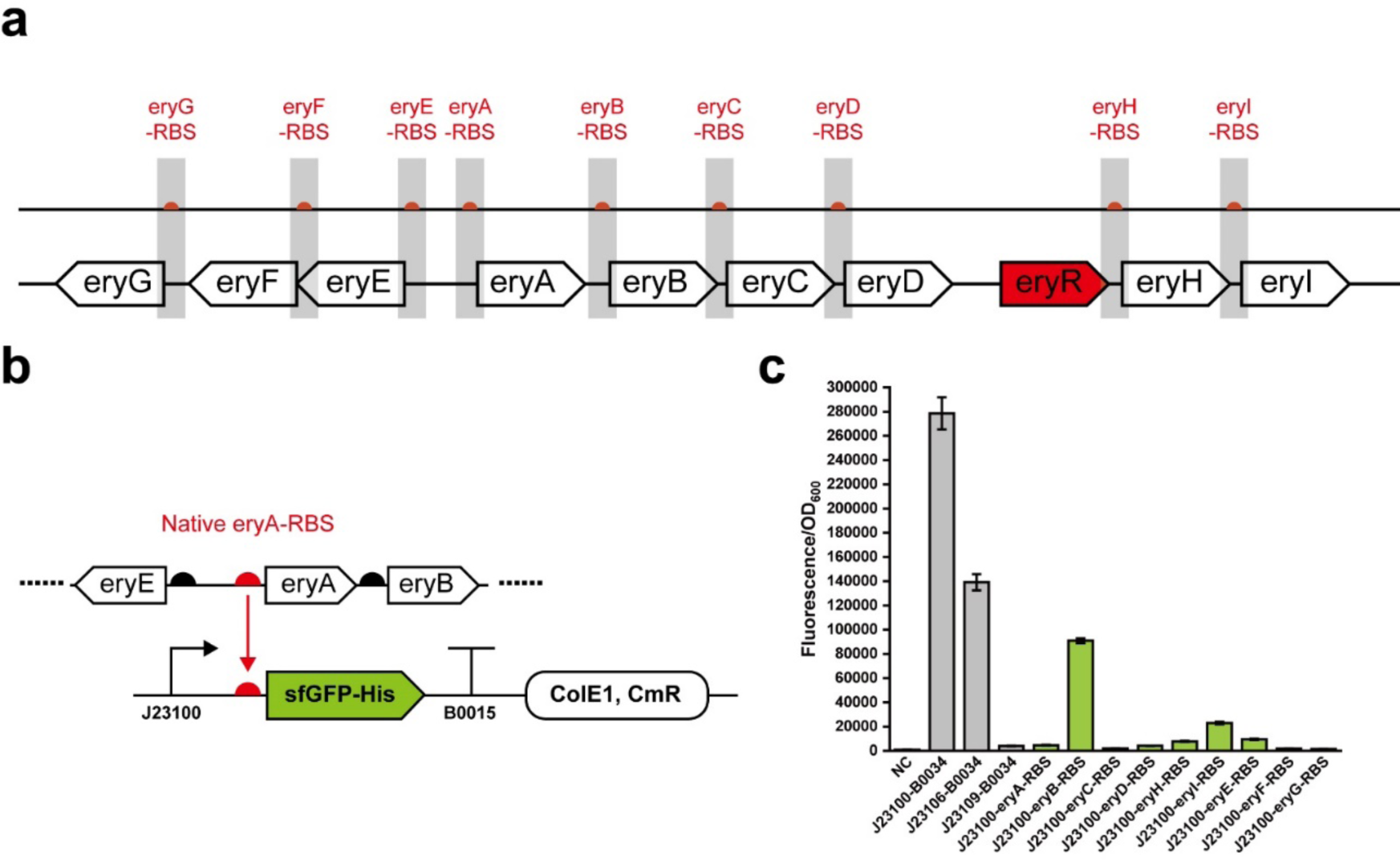
Native RBS characterization in the erythritol gene cluster. (**a**) Schematic of RBS localization. Note that the function of eryR was not successfully characterized, thus here we did not test eryR-RBS. (**b**) Schematic procedure of “ery-RBS” reporter plasmid construction (pFB168 to pFB176). Here we illustrate the construction of J23100-eryA-RBS-sfGFP reporter plasmid as an example. The putative “eryA-RBS” sequence (approximately 40 bp) was assembled with the promoter J23100, sfGFP (6xHis), and a terminator B0015 in the vector pSB1C3 (pFB168). (**c**) Normalized fluorescence of all nine RBS parts. All the measurements were performed in *E. coli* Mach1-T1 with three biological replicates. “NC” represents *E. coli* Mach1-T1 without plasmids. The other three gray columns correspond to pFB286, pFB288, and pFB289, respectively, to act as positive controls (iGEM standard biological parts). The results indicate that “eryB-RBS” is a relatively strong RBS, but the other eight RBSs are not strong.

**Extended Data Fig. 4.**
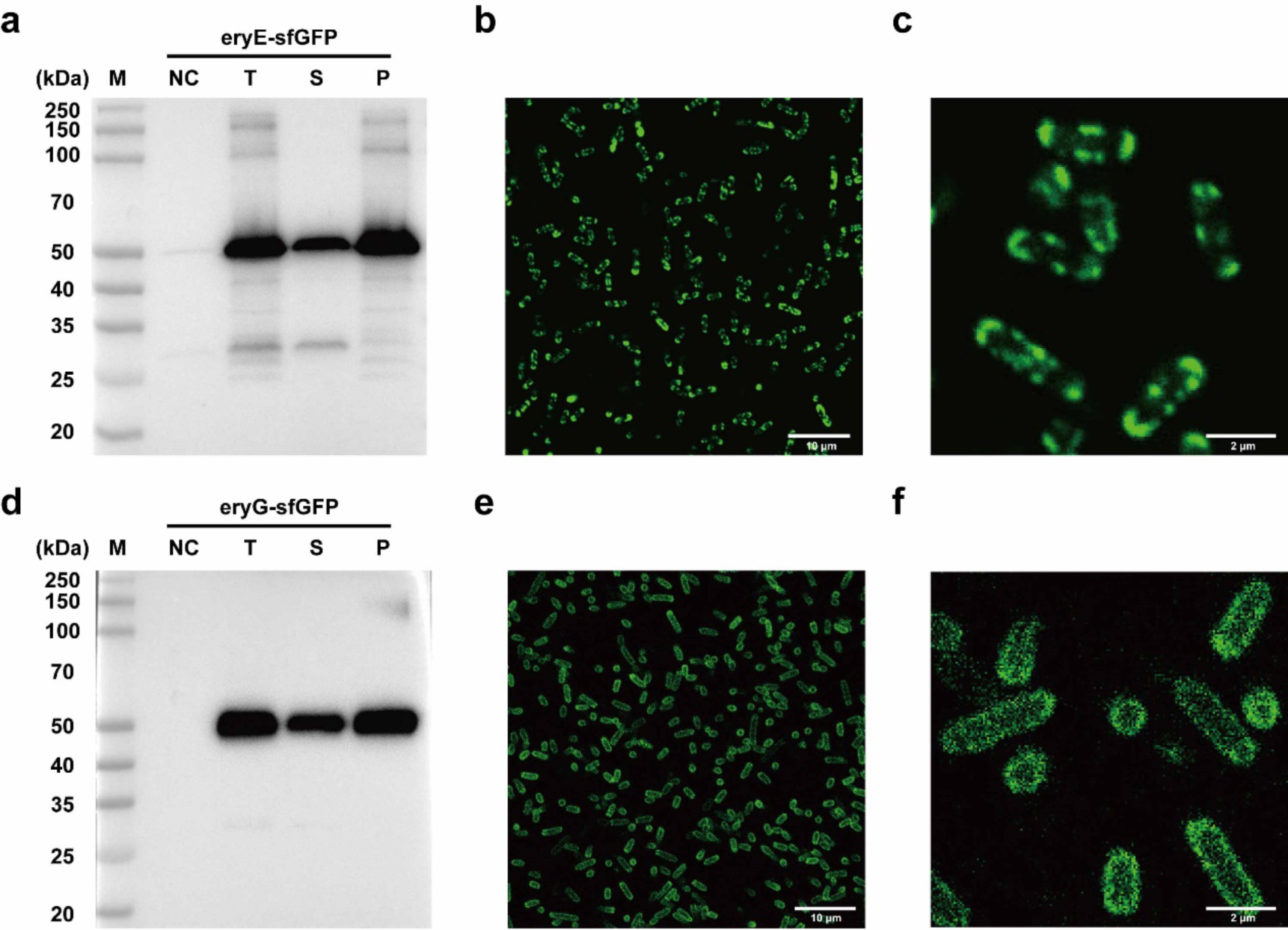
eryE-sfGFP and eryG-sfGFP characterization. (**a**) Western-Blot analysis of eryE-sfGFP(6xHis) (50.4 kDa). *E. coli* Mach1-T1 harboring the plasmid pFB162 was initially inoculated with overnight culture (5 mL LB medium at 37°C and 250 rpm for 16 h) as a volume ratio of 1:500, and then incubated in 5 mL new LB medium at 37°C and 250 rpm. When OD_600_ reached 0.6, 1% arabinose (w/v) was added and incubated for another 6 h at 30°C and 250 rpm. Then, cell pellets were resuspended (equal volume) with 1x phosphate-buffered saline (pH 7.4) and lysed by sonication. The total cell lysate “T”, supernatant “S”, and pellets “P” were analyzed by Western-Blot. “NC” represents the total cell lysate of *E. coli* Mach1-T1 harboring pFB162 without arabinose induction. (**b**) Confocal laser scanning microscopy image of *E. coli* Mach1-T1 expressing eryE-sfGFP. Green border indicates the transmembrane localization of eryE-sfGFP. (**c**) Partially enlarged view of the image in (**b**). (**d**) Western-Blot analysis of eryG-sfGFP(6xHis) (64.6 kDa). *E. coli* Mach1-T1 harboring the plasmid pFB164 was initially incubated in 5 mL LB medium at 37°C and 250 rpm. When OD_600_ reached 0.6, 1% arabinose (w/v) was added and incubated for another 6 h at 30°C and 250 rpm. Then, cell pellets were resuspended (equal volume) with 1x phosphate-buffered saline (pH 7.4) and lysed by sonication. The total cell lysate “T”, supernatant “S”, and pellets “P” were analyzed by Western-Blot. “NC” represents the total cell lysate of *E. coli* Mach1-T1 harboring plasmid pFB164 without arabinose induction. (**e**) Confocal laser scanning microscopy image of *E. coli* Mach1-T1 expressing eryG-sfGFP. Green border indicates the transmembrane localization of eryG-sfGFP. (**f**) Partially enlarged view of the image in (**e**).

**Extended Data Fig. 5.**
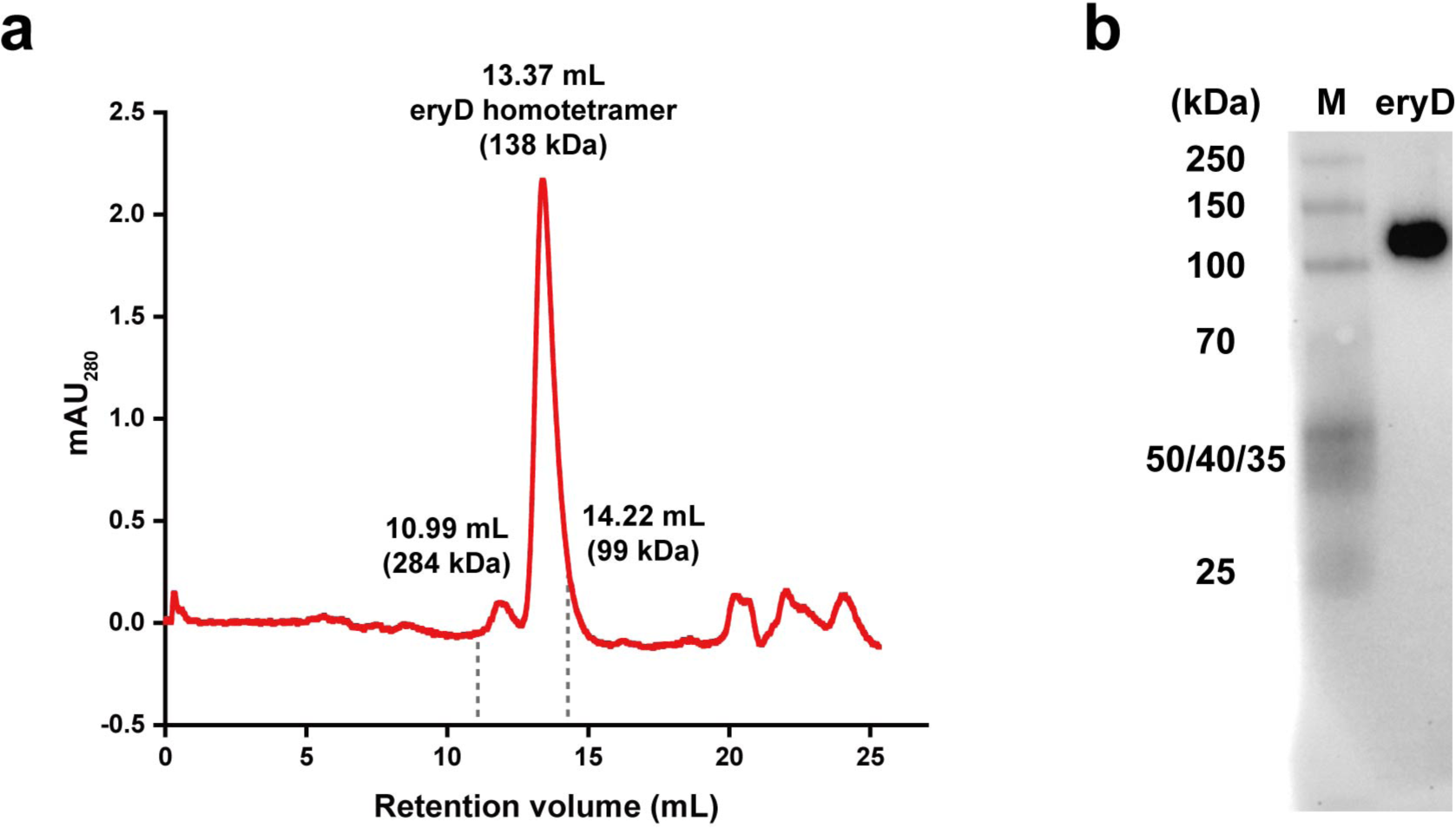
eryD assembled as a homotetramer. (**a**) Size exclusion chromatography of the purified eryD (6xHis) with Superdex 200 increase 10/300 GL column. eryD homotetramer was eluted at 13.37 mL (red peak). The other two purified proteins were used as standards: Vlm2 (284 kDa) was eluted at 10.99 mL and T7 RNA polymerase (99 kDa) was eluted at 14.22 mL. (**b**) Native-PAGE based Western-Blot analysis of the purified eryD (6xHis). The expected homotetramer band (approximately 138 kDa) was shown between the two protein markers of 100 kDa and 150 kDa.

**Extended Data Fig. 6.**
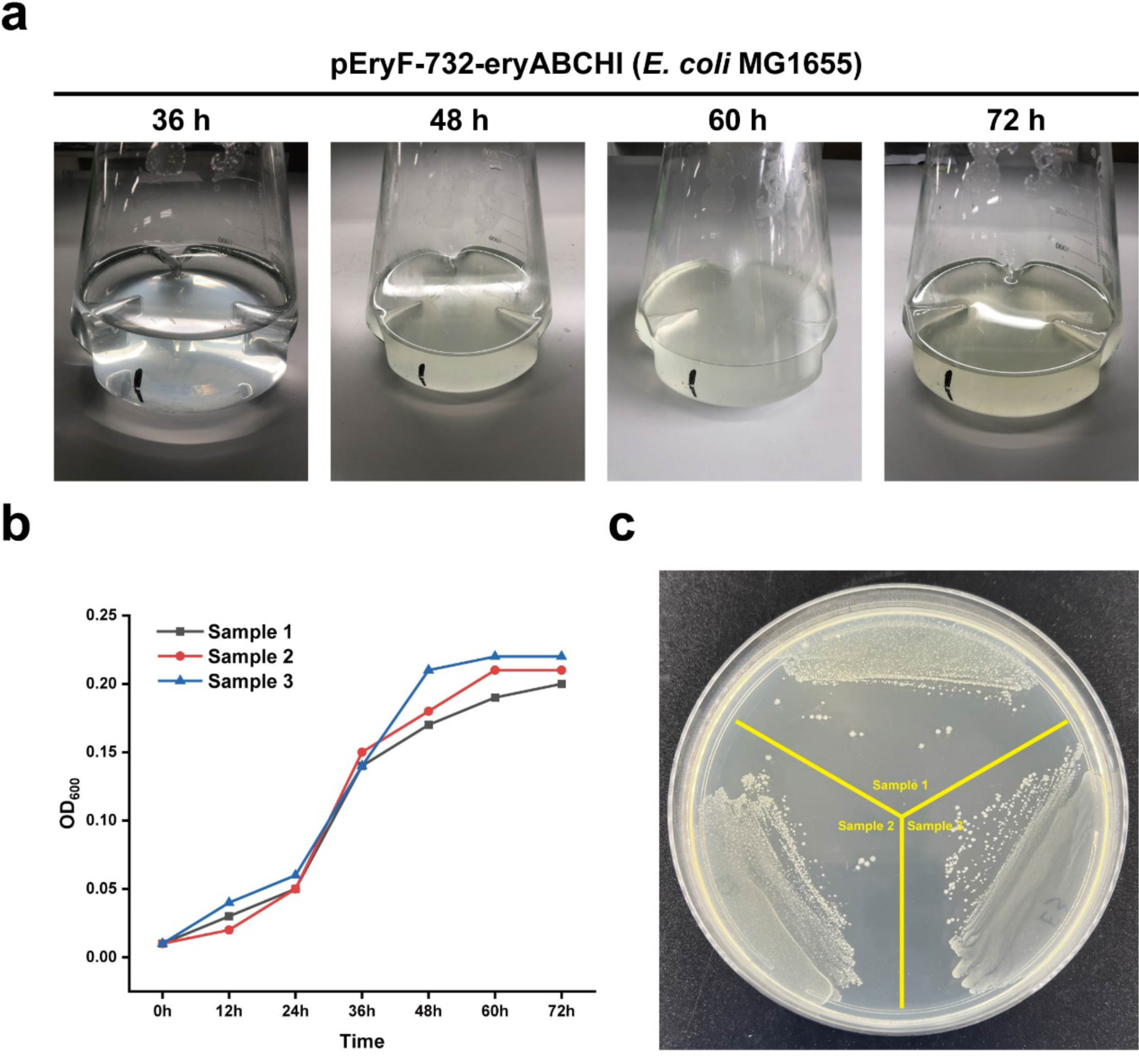
Sample preparation for mRNA transcriptional analysis. Wild-type *E. coli* MG1655 harboring plasmid pFB147 was incubated in 5 mL LB medium (with 34 μg/mL chloramphenicol) for overnight cultivation (37°C and 250 rpm for 16 h). On the second day, 5 mL cell culture was washed with 1x phosphate-buffered saline (pH 7.4) by three times and then resuspended by 5 mL M9-erythritol liquid medium. Then, the resuspended cell mixture was inoculated into 1 L M9-erythritol liquid medium (with 6 μg/mL chloramphenicol) as 1:200 v/v in 2 L shaking flask. After that, the cells were incubated for 72 h (37°C and 250 rpm). Each cultivation was repeated with three biological replicates. The cells after 36 h cultivation were collected to prepare RNA-seq samples. (**a**) The cell cultures in pre-experiments reached stationary phase at 72 h. (**b**) Cell growth curves of pre-experiment samples. The middle logarithmic phase reached at 36 h and the stationary phase reached at 72 h. (**c**) The cell cultures (samples in pre-experiment at 72 h) were picked and spread on LB-agar plates (with 34 μg/mL chloramphenicol) to test the cell viability (overnight cultivation on the plate for 16 h at 37°C). The colonies were normal and did not lose the plasmid pFB147.

**Extended Data Fig. 7.**
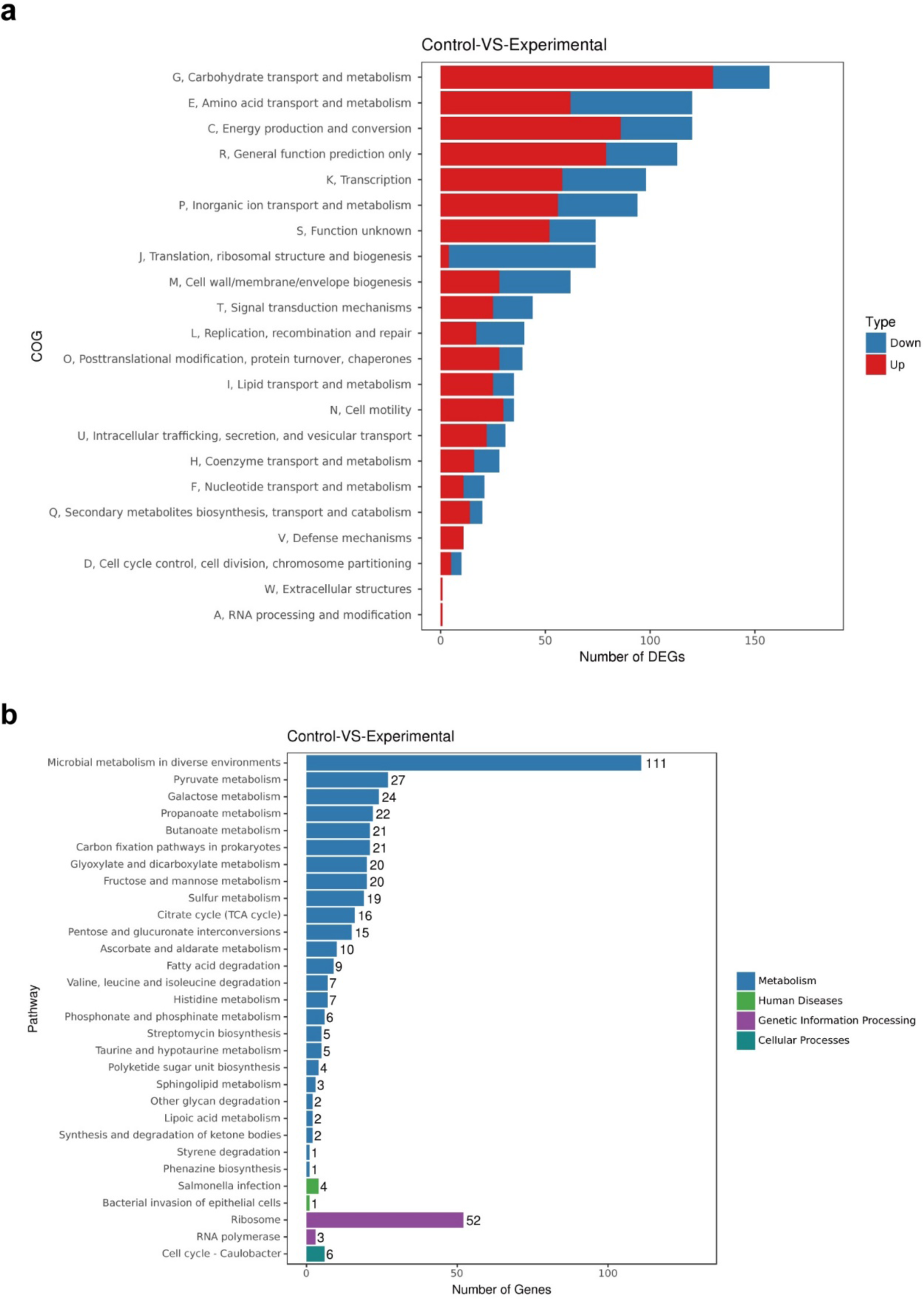
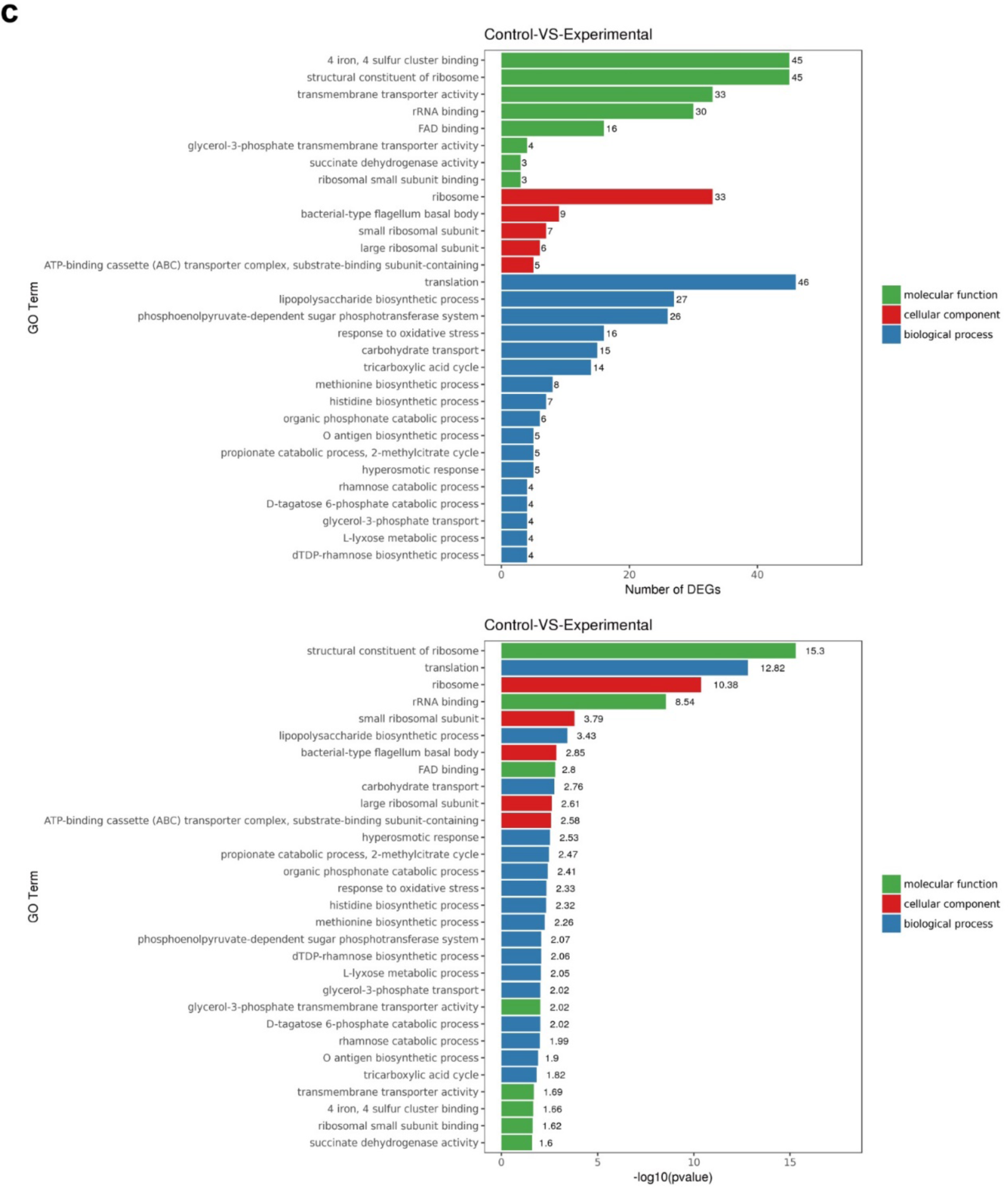
mRNA transcriptional analysis. Transcriptome results were analyzed via (**a**) COG, (**b**) KEGG, and (**c**) GO databases. The shared different genes were associated with carbohydrate metabolism, amino acid metabolism, and ribosome. The genes of carbohydrate metabolism were up-regulated; however, ribosome and translation related genes were down-regulated.

**Extended Data Fig. 8.**
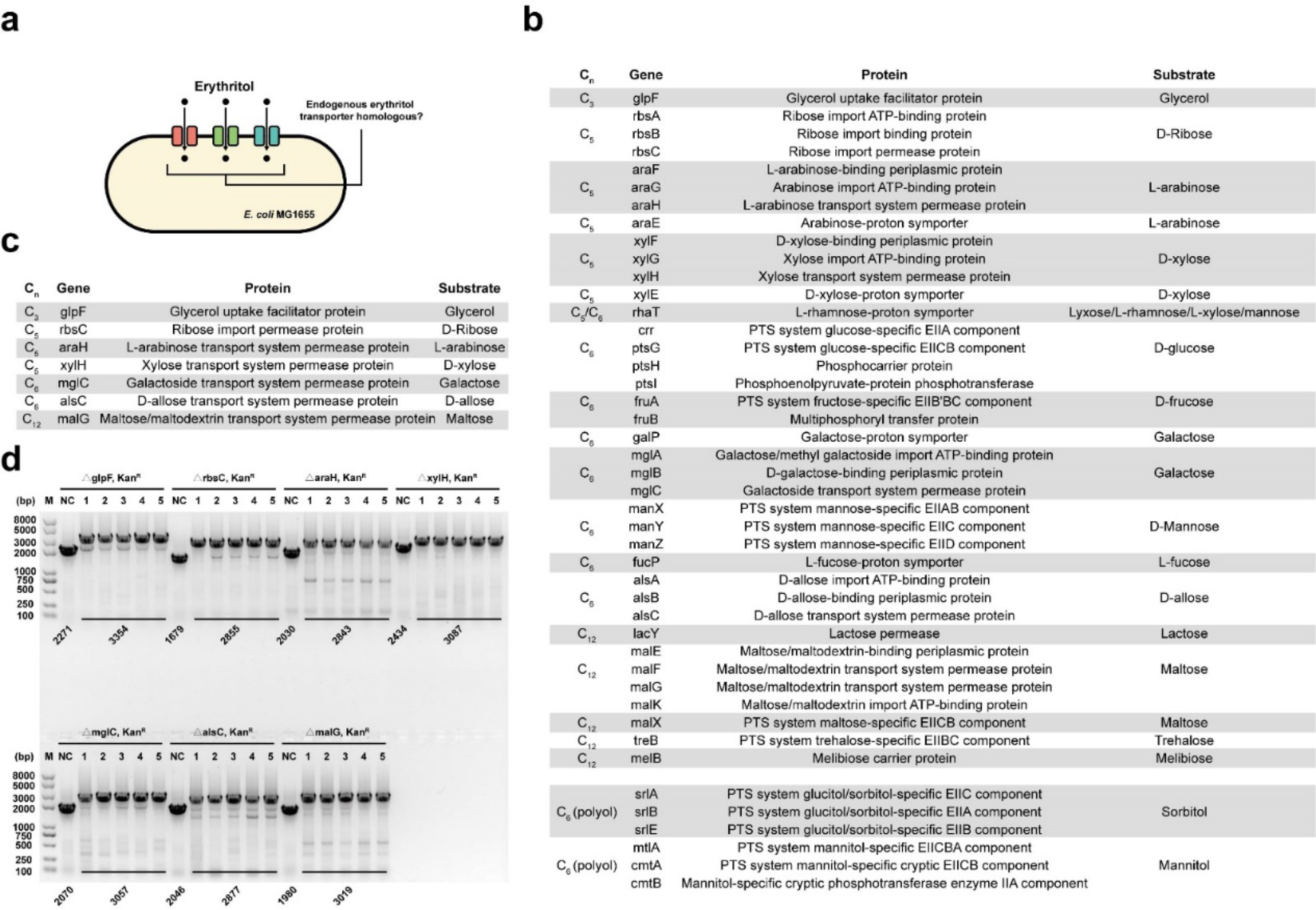
Potential endogenous erythritol transporter homologous screening. (**a**) Schematic diagram of screening *E. coli* MG1655 native transporters that might facilitate erythritol transportation. (**b**) List of all carbohydrate transportation genes and proteins in *E. coli* MG1655. (**c**) Screened seven carbohydrate transporter-associated genes. glpF has been reported that is able to transport erythritol. The other six genes are somewhat homologous to erythritol ABC transporter gene eryG by BLAST protein-protein alignment analysis. (**d**) Construction of mutated *E. coli* MG1655 strains by Lambda-Red recombination. In each of the seven mutated strains, “NC” represents the wild-type *E. coli* MG1655 genome PCR product by the primer pair “forward left-homologous-arm” and “reverse right-homologous-arm”. The other five PCR products are the desired mutation strains by the same primer pair. The KanR genes are not deleted by pCP20.

**Extended Data Fig. 9.**
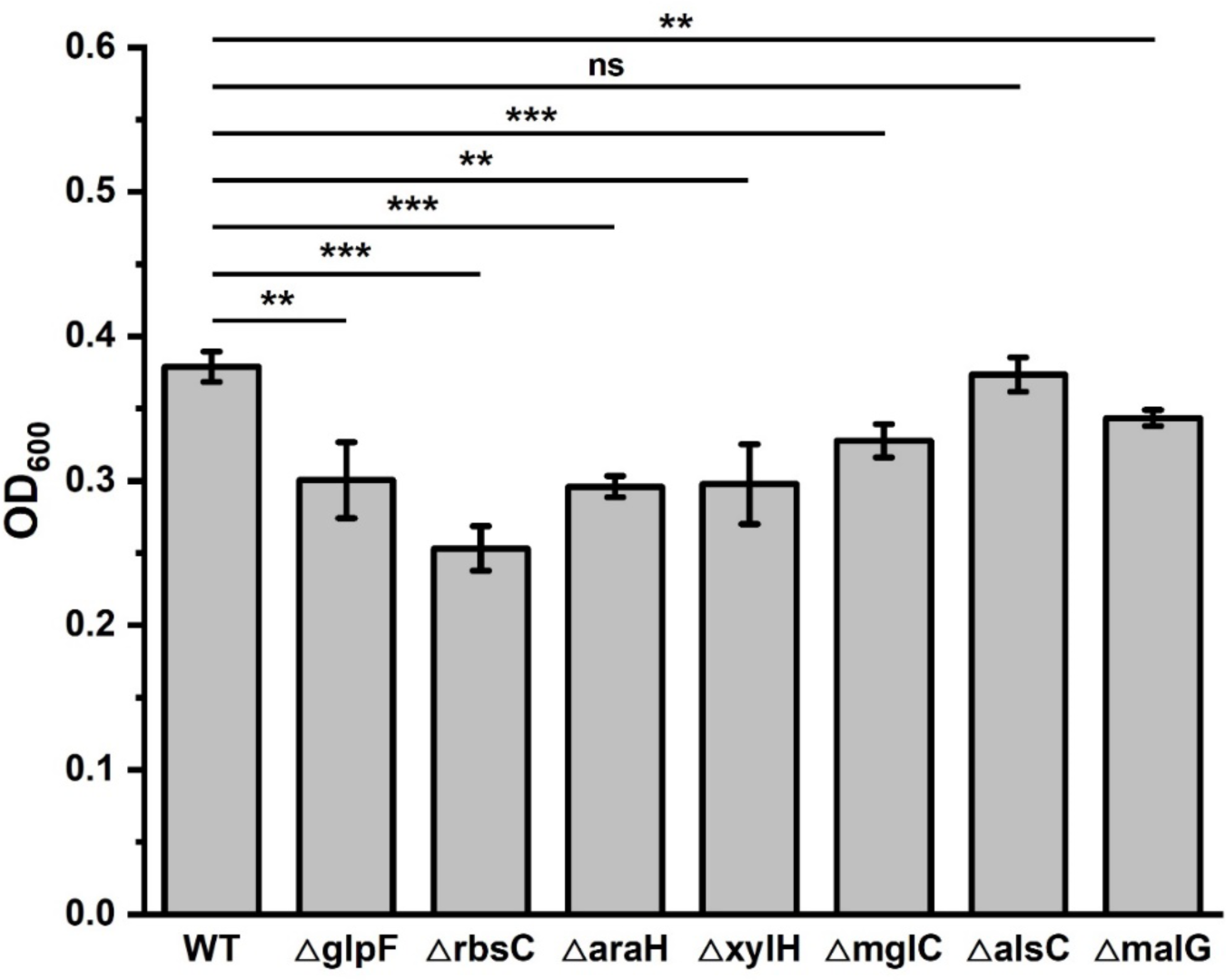
Several eryG homologous permeases in *E. coli* may facilitate erythritol catabolism. BLAST protein-protein alignment results showed that seven *E. coli* native carbohydrate ABC-transporter permeases are homologous to erythritol permease eryG. We then constructed 7 different knock-out *E. coli* MG1655 strains by Lambda-Red recombination. All these 7 strains and the WT (wild-type *E. coli* MG1655) strain were transformed with pFB147 to utilize erythritol. Then, all strains were incubated (M9-erythritol medium, 37°C, no shaking) in 96-well plates for 72 h (stationary phase). OD_600_ values were measured at each 12 h. The final OD_600_ values at 72 h were shown in the graph. All values were measured as three biological replicates. The error bars represent the standard deviation (s.d.). Student’s *t*-tests are used for statistical analysis and *p* < 0.05 indicates statistical significance (\*\**p* < 0.01 and \*\*\**p* < 0.001, ns, no significance).

## Notes

### Competing Interest Statement

The authors have declared no competing interest.

### Summary of Updates

A revised manuscript as version 2.

## References

1. Grembecka, M. Sugar alcohols-their role in the modern world of sweeteners: a review. Eur. Food Res. Technol. 241, 15–16 (2015).

2. Shankar, P., Ahuja, S. & Sriram, K. Non-nutritive sweeteners: review and update. Nutrition 29, 1293–1299 (2013).

3. Bernt, W. O., Borzelleca, J. F., Flamm, G., & Munro, I. C. Erythritol: a review of biological and toxicological studies. Regul. Toxicol. Pharmacol. 24, S191–S197 (1996).

4. Munro, I. C., Berndt, W. O., Borzelleca, J. F., Flamm, G., Lynch, B. S., Kennepohl, E., Bär, E. A., & Modderman, J. Erythritol: an interpretive summary of biochemical, metabolic, toxicological and clinical data. Food Chem. Toxicol. 36, 1139–1174 (1998).

5. Oku, T., & Okazaki M. Laxative threshold of sugar alcohol erythritol in human subjects. Nutr. Res. 16, 577–589 (1996).

6. Veiga-da-Cunha, M., Santos, H., & Van Schaftingen, E. Pathway and regulation of erythritol formation in *Leuconostoc oenos*. J Bacteriol. 175, 3941–3948 (1993).

7. Richter, H., Vlad, D., & Unden, G. Significance of pantothenate for glucose fermentation by *Oenococcus oeni* and for suppression of the erythritol and acetate production. Arch. Microbiol. 175, 26–31 (2001).

8. Mirończuk, A., Dobrowolski, A., Rakicka, M., Rywińska, A., & Rymowicz, W. Newly isolated mutant of *Yarrowia lipolytica* MK1 as a proper host for efficient erythritol biosynthesis from glycerol. Process Biochem. 50, 61–68 (2015).

9. Rymowicz, W., Rywińska, A., & Marcinkiewicz, M. High-yield production of erythritol from raw glycerol in fed-batch cultures of *Yarrowia lipolytica*. Biotechnol. Lett. 31, 377–380 (2009).

10. Qiu, X., Xu, P., Zhao, X., Du, G., Zhang, J., & Li, J. Combining genetically-encoded biosensors with high throughput strain screening to maximize erythritol production in *Yarrowia lipolytica*. Metab. Eng. 60, 66–76 (2020).

11. de Cock, P. Erythritol. in Sweeteners and Sugar Alternatives in Food Technology (eds. O’Donnell, K. & Kearsley, M. W.) 213–241 (Wiley Online Library, 2012).

12. Livesey G. Health potential of polyols as sugar replacers, with emphasis on low glycaemic properties. Nutr. Res. Rev. 16, 163–191 (2003).

13. Arrigoni, E., Brouns, F., & Amadò, R. Human gut microbiota does not ferment erythritol. Br. J. Nutr. 94, 643–646 (2005).

14. den Hartog, G. J., Boots, A. W., Adam-Perrot, A., Brouns, F., Verkooijen, I. W., Weseler, A. R., Haenen, G. R., & Bast, A. Erythritol is a sweet antioxidant. Nutrition 26, 449–458 (2010).

15. Ruiz-Ojeda, F. J., Plaza-Díaz, J., Sáez-Lara, M. J., & Gil, A. Effects of sweeteners on the gut microbiota: a review of experimental studies and clinical trials. Adv Nutr. 10, S31–S48 (2019).

16. Carocho, M., Morales, P., & Ferreira, I. Sweeteners as food additives in the XXI century: a review of what is known, and what is to come. Food Chem. Toxicol. 107, 302–317 (2017).

17. Barbier, T., Collard, F., Zúñiga-Ripa, A., Moriyón, I., Godard, T., Becker, J., Wittmann, C., Van Schaftingen, E., & Letesson, J. J. Erythritol feeds the pentose phosphate pathway via three new isomerases leading to D-erythrose-4-phosphate in *Brucella*. Proc. Natl. Acad. Sci. U. S. A. 111, 17815–17820 (2014).

18. Lillo, A. M., Tetzlaff, C. N., Sangari, F. J., & Cane, D. E. Functional expression and characterization of EryA, the erythritol kinase of *Brucella abortus*, and enzymatic synthesis of L-erythritol-4-phosphate. Bioorg. Med. Chem. Lett. 13, 737–739 (2003).

19. Sangari, F. J., Agüero, J., & Garcı A-Lobo, J. M. The genes for erythritol catabolism are organized as an inducible operon in *Brucella abortus*. Microbiology 146, 487–495 (2000).

20. Sperry, J. F., & Robertson, D. C. Erythritol catabolism by *Brucella abortus*. J. Bacteriol. 121, 619–630 (1975).

21. Rodríguez, M. C., Viadas, C., Seoane, A., Sangari, F. J., López-Goñi, I., & García-Lobo, J. M. Evaluation of the effects of erythritol on gene expression in *Brucella abortus*. PLoS O 7, e50876 (2012).

22. Geddes, B. A., Hausner, G., & Oresnik, I. J. Phylogenetic analysis of erythritol catabolic loci within the *Rhizobiales* and proteobacteria. BMC Microbiol. 13, 46 (2013).

23. Barbier, T. et al. *Brucella* central carbon metabolism: an update. Crit. Rev. Microbiol. 44, 182–211 (2018).

24. Geddes, B. A., & Oresnik, I. J. Genetic characterization of a complex locus necessary for the transport and catabolism of erythritol, adonitol and L-arabitol in *Sinorhizobium meliloti*. Microbiology 158, 2180–2191 (2012).

25. Huang, H., Carter, M. S., Vetting, M. W., Al-Obaidi, N., Patskovsky, Y., Almo, S. C., & Gerlt, J. A. A general strategy for the discovery of metabolic pathways: D-threitol, L-threitol, and erythritol utilization in *Mycobacterium smegmatis*. J. Am. Chem. Soc. 137, 14570–14573 (2015).

26. Park, S. Y., Eun, H., Lee, M. H., & Lee, S. Y. Metabolic engineering of *Escherichia coli* with electron channelling for the production of natural products. Nat. Catal. 5, 726–737 (2022).

27. Ayikpoe, R. S. et al. A scalable platform to discover antimicrobials of ribosomal origin. Nat. Commun. 13, 6135 (2022).

28. Li, J., & Neubauer, P. *Escherichia coli* as a cell factory for heterologous production of nonribosomal peptides and polyketides. New Biotechnol. 31, 579–585 (2014).

29. Bang, J., & Lee, S. Y. Assimilation of formic acid and CO_2_ by engineered *Escherichia coli* equipped with reconstructed one-carbon assimilation pathways. Proc. Natl. Acad. Sci. U. S. A. 115, E9271–E9279 (2018).

30. Gleizer, S. et al. Conversion of *Escherichia coli* to generate all biomass carbon from CO_2_. Cell 179, 1255–1263 (2019).

31. Chen, F. Y., Jung, H. W., Tsuei, C. Y., & Liao, J. C. Converting *Escherichia coli* to a synthetic methylotroph growing solely on methanol. Cell 182, 933–946 (2020).

32. Keller, P., Reiter, M. A., Kiefer, P., Gassler, T., Hemmerle, L., Christen, P., Noor, E., & Vorholt, J. A. Generation of an *Escherichia coli* strain growing on methanol via the ribulose monophosphate cycle. Nat. Commun. 13, 5243 (2022).

33. Adams B. L. The next generation of synthetic biology chassis: moving synthetic biology from the laboratory to the field. ACS Synth. Biol. 5, 1328–1330 (2016).

34. Reister, M., Hoffmeier, K., Krezdorn, N., Rotter, B., Liang, C., Rund, S., Dandekar, T., Sonnenborn, U., & Oelschlaeger, T. A. Complete genome sequence of the gram-negative probiotic *Escherichia coli* strain Nissle 1917. J. Biotechnol. 187, 106–107 (2014).

35. Bathe, S., Achouak, W., Hartmann, A., Heulin, T., Schloter, M., & Lebuhn, M. Genetic and phenotypic microdiversity of *Ochrobactrum* spp. FEMS Microbiol. Ecol. 56, 272–280 (2006).

36. Chain, P. S., Comerci, D. J., Tolmasky, M. E., Larimer, F. W., Malfatti, S. A., Vergez, L. M., Aguero, F., Land, M. L., Ugalde, R. A., & Garcia, E. Whole-genome analyses of speciation events in pathogenic Brucellae. Infect. Immun. 73, 8353–8361 (2005).

37. Lamontagne, J., Béland, M., Forest, A., Côté-Martin, A., Nassif, N., Tomaki, F., Moriyón, I., Moreno, E., & Paramithiotis, E. Proteomics-based confirmation of protein expression and correction of annotation errors in the *Brucella abortus* genome. BMC Genomics 11, 300 (2010).

38. Kelly, J. R., Rubin, A. J., Davis, J. H., Ajo-Franklin, C. M., Cumbers, J., Czar, M. J., de Mora, K., Glieberman, A. L., Monie, D. D., & Endy, D. Measuring the activity of BioBrick promoters using an *in vivo* reference standard. J. Biol. Eng. 3, 4 (2009).

39. Beal, J., Haddock-Angelli, T., Baldwin, G., Gershater, M., Dwijayanti, A., Storch, M., de Mora, K., Lizarazo, M., Rettberg, R., & with the iGEM Interlab Study Contributors. Quantification of bacterial fluorescence using independent calibrants. PLoS One 13, e0199432 (2018).

40. Beal, J. et al. Comparative analysis of three studies measuring fluorescence from engineered bacterial genetic constructs. PLoS One 16, e0252263 (2021).

41. Pellicer, M. T., Fernandez, C., Badía, J., Aguilar, J., Lin, E. C., & Baldom, L. Cross-induction of *glc* and *ace* operons of *Escherichia coli* attributable to pathway intersection. Characterization of the *glc* promoter. J. Biol. Chem. 274, 1745–1752 (1999).

42. Seweryn, P., Van, L. B., Kjeldgaard, M., Russo, C. J., Passmore, L. A., Hove-Jensen, B., Jochimsen, B., & Brodersen, D. E. Structural insights into the bacterial carbon-phosphorus lyase machinery. Nature 525, 68–72 (2015).

43. Altschul, S. F., Madden, T. L., Schäffer, A. A., Zhang, J., Zhang, Z., Miller, W., & Lipman, D. J. Gapped BLAST and PSI-BLAST: a new generation of protein database search programs. Nucleic Acids Res. 25, 3389–3402 (1997).

44. Johnson, M., Zaretskaya, I., Raytselis, Y., Merezhuk, Y., McGinnis, S., & Madden, T. L. NCBI BLAST: a better web interface. Nucleic Acids Res. 36, W5–W9 (2008).

45. Heller, K. B., Lin, E. C., & Wilson, T. H. Substrate specificity and transport properties of the glycerol facilitator of *Escherichia coli*. J. Bacteriol. 144, 274–278 (1980).

46. Shindou, T., Sasaki, Y., Eguchi, T., Euguchi, T., Hagiwara, K., & Ichikawa, T. Identification of erythritol by HPLC and GC-MS and quantitative measurement in pulps of various fruits. J. Agric. Food Chem. 37, 1474–1476 (1989).

47. Sonnenborn U. *Escherichia coli* strain Nissle 1917-from bench to bedside and back: history of a special *Escherichia coli* strain with probiotic properties. FEMS Microbiol. Lett. 363, fnw212 (2016).

48. Pedrolli, D. B., Ribeiro, N. V., Squizato, P. N., de Jesus, V. N., Cozetto, D. A., & Team AQA Unesp at iGEM 2017. Engineering microbial living therapeutics: the synthetic biology toolbox. Trends Biotechnol. 37, 100–115 (2019).

49. Kelly, V. W., Liang, B. K., & Sirk, S. J. Living therapeutics: the next frontier of precision medicine. ACS Synth. Biol. 9, 3184–3201 (2020).

50. Cubillos-Ruiz, A., Guo, T., Sokolovska, A., Miller, P. F., Collins, J. J., Lu, T. K., & Lora, J. M. Engineering living therapeutics with synthetic biology. Nat. Rev. Drug Discov. 20, 941–960 (2021).

51. Forker, E. L. Two sites of bile formation as determined by mannitol and erythritol clearance in the guinea pig. J. Clin. Invest. 46, 1189–1195 (1967).

52. Ba, F., Liu, Y., Liu, W. Q., Tian, X., & Li, J. SYMBIOSIS: synthetic manipulable biobricks via orthogonal serine integrase systems. Nucleic Acids Res. 50, 2973–2985 (2022).

53. Zhuang, L., Huang, S., Liu, W. Q., Karim, A. S., Jewett, M. C., & Li, J. Total *in vitro* biosynthesis of the nonribosomal macrolactone peptide valinomycin. Metab. Eng. 60, 37–44 (2020).

54. Brodkorb, A. et al. INFOGEST static *in vitro* simulation of gastrointestinal food digestion. Nat. Protoc. 14, 991–1014 (2019).

55. Minekus, M. et al. A standardised static *in vitro* digestion method suitable for food - an international consensus. Food Funct. 5, 1113–1124 (2014).

