## Supplementary Information for "Engineering *Escherichia coli* to utilize erythritol as sole carbon source"

I. Supplementary Tables

I. Supplementary Tables

### Supplementary Table 1. *E. coli* strains.

| Strain name | Purpose | Genotype | Origin |
|---|---|---|---|
| Mach1-T1 | Molecular cloning, promoter characterization, ribosome binding site characterization | $F^-$ φ80(*lac*Z)Δ*M15* Δ*lac*X74 *hsd*R($r_K^-$ $m_K^+$) Δ*rec*A1398 *end*A1 *ton*A | TransGen Biotech |
| MG1655 | Erythritol catabolism characterization, host strain of RNA-Seq | K12 $F^-$ *lambda- ilvG- rfb-50 rph-1* | Shanghai Weidi Biotechnology |
| BL21(DE3) | Promoter characterization, ribosome binding site characterization | $F^-$ *omp*T *hsd*$S_B$($r_B^-$ $m_B^-$) *gal dcm*(DE3) | TransGen Biotech |
| DH5α λpir | Cloning R6K-ori plasmids | $F^-$ φ80 *lac*ZΔM15 Δ(*lac*ZYA-*arg* F) *LAMpir* U169 *end*A1 *rec*A1 *hsd*R17($r_k^-$,$m_k^+$) *sup*E44λ- *thi* -1 *gyr*A96 *rel*A1 *pho*A | Shanghai Weidi Biotechnology |
| Nissle 1917 (EcN) | Host strain of simulated intestinal fluid experiment | | Mutaflor |

All *E. coli* strains used in this study are listed above.

### Supplementary Table 2. Genetic parts.

| Part name | Type | DNA sequence | Reference |
|---|---|---|---|
| J23100 | promoter | ttgacggctagctcagtcctaggtacagtgctagc | 1, BBa_J23100 |
| J23105 | promoter | tttacggctagctcagtcctaggtactatgctagc | 1, BBa_J23105 |
| J23106 | promoter | tttacggctagctcagtcctaggtatagtgctagc | 1, BBa_J23106 |
| J23109 | promoter | tttacagctagctcagtcctagggactgtgctagc | 1, BBa_J23109 |
| J23113 | promoter | ctgatggctagctcagtcctagggattatgctagc | 1, BBa_J23113 |
| J23114 | promoter | tttatggctagctcagtcctaggtacaatgctagc | 1, BBa_J23114 |
| pEryF-732 | promoter | tatcaaacgccttcagcaattctgccgtattgtcgttatggaaacggatgatgcggctgaattagaccgccc<br>ggagattaccctcgaccattgtcatcacgcttaccggataggcaaaagcctccgccaaagcggtgcagg<br>acgcgacagacgagatcatgatagcctccacatgaattccagggccccgctcctccgtgcatcctggaa<br>acagatgcgccgcaatcaaaacctatgatagcggcgtgttattaactccaataatgatgttataaagctcg<br>aaagctgtcaatcggatgttttcgaaacaccggtgaaatagcacaatttcgcgccgaatctcgatcacgat<br>ctagagaaatccactacataattcgttatcaatcgcactattcacagcaccgtcttttctcgcaaagccagcc<br>acaaagatactcacaaacttctatattttcttttattttcaatgaattaaacatatttaaaaccgaagaaacatta<br>agctattgatggaaaacagtgggaaaaccgcactcgaaggcgcggagatttcccctacattcactctttgg<br>tgaaaaaaaatgcgccatctagaaaattttacagacaacgtgatagcgttatgttatctccagcatacgcc<br>atcgcccgatttcattataaacagggctgcgagcgacgtattttcggtcgcgggaggaatccaaatgtcaa<br>tgaaagcacatagccgcc | This work |
| pEryF-632 | promoter | gcttaccggataggcaaaagcctccgccaaagcggtgcaggacgcgacagacgagatcatgatagcc<br>tccacatgaattccagggccccgctcctccgtgcatcctggaaacagatgcgccgcaatcaaaacctatg<br>atagcggcgtgttattaactccaataatgatgttataaagctcgaaagctgtcaatcggatgttttcgaaaca<br>ccggtgaaatagcacaatttcgcgccgaatctcgatcacgatctagagaaatccactacataattcgttatc<br>aatcgcactattcacagcaccgtcttttctcgcaaagccagccacaaagatactcacaaacttctatattttct<br>tttattttcaatgaattaaacatatttaaaaccgaagaaacattaagctattgatggaaaacagtgggaaaa<br>ccgcactcgaaggcgcggagatttcccctacattcactctttggtgaaaaaaaatgcgccatctagaaaat<br>tttacagacaacgtgatagcgttatgttatctccagcatacgccatcgcccgatttcattataaacagggctg<br>cgagcgacgtattttcggtcgcgggaggaatccaaatgtcaatgaaagcacatagccgcc | This work |
| pEryF-532 | promoter | catcctggaaacagatgcgccgcaatcaaaacctatgatagcggcgtgttattaactccaataatgatgtt<br>ataaagctcgaaagctgtcaatcggatgttttcgaaacaccggtgaaatagcacaatttcgcgccgaatct<br>cgatcacgatctagagaaatccactacataattcgttatcaatcgcactattcacagcaccgtcttttctcgca<br>aagccagccacaaagatactcacaaacttctatattttcttttattttcaatgaattaaacatatttaaaaccga<br>agaaacattaagctattgatggaaaacagtgggaaaaccgcactcgaaggcgcggagatttcccctac<br>attcactctttggtgaaaaaaaatgcgccatctagaaaattttacagacaacgtgatagcgttatgttatctcc<br>agcatacgccatcgcccgatttcattataaacagggctgcgagcgacgtattttcggtcgcgggaggaatc<br>caaatgtcaatgaaagcacatagccgcc | This work |
| pEryF-432 | promoter | tttcgaaacaccggtgaaatagcacaatttcgcgccgaatctcgatcacgatctagagaaatccactacat<br>aattcgttatcaatcgcactattcacagcaccgtcttttctcgcaaagccagccacaaagatactcacaaac<br>ttctatattttcttttattttcaatgaattaaacatatttaaaaccgaagaaacattaagctattgatggaaaaca<br>gtgggaaaaccgcactcgaaggcgcggagatttcccctacattcactctttggtgaaaaaaaatgcgcca<br>tctagaaaattttacagacaacgtgatagcgttatgttatctccagcatacgccatcgcccgatttcattataa<br>acagggctgcgagcgacgtattttcggtcgcgggaggaatccaaatgtcaatgaaagcacatagccgcc | This work |
| pEryF-332 | promoter | accgtcttttctcgcaaagccagccacaaagatactcacaaacttctatattttcttttattttcaatgaattaaa<br>catatttaaaaccgaagaaacattaagctattgatggaaaacagtgggaaaaccgcactcgaaggcgc<br>ggagatttcccctacattcactctttggtgaaaaaaaatgcgccatctagaaaattttacagacaacgtgat | This work |

| | | | |
|---|---|---|---|
| | | agcgttatgttatctccagcatacgccatcgcccgatttcattataaacagggctgcgagcgacgtattttcg gtcgcgggaggaatccaaatgtcaatgaaagcacatagccgcc | |
| pEryF-232 | promoter | aagctattgatggaaaacagtgggaaaaccgcactcgaaggcgcggagatttcccctacattcactctttg gtgaaaaaaaatgcgccatctagaaaattttacagacaacgtgatagcgttatgttatctccagcatacgc catcgcccgatttcattataaacagggctgcgagcgacgtattttcggtcgcgggaggaatccaaatgtca atgaaagcacatagccgcc | This work |
| pEryF-222 | promoter | tggaaaacagtgggaaaaccgcactcgaaggcgcggagatttcccctacattcactctttggtgaaaaa aaatgcgccatctagaaaattttacagacaacgtgatagcgttatgttatctccagcatacgccatcgcccg atttcattataaacagggctgcgagcgacgtattttcggtcgcgggaggaatccaaatgtcaatgaaagca catagccgcc | This work |
| pEryF-212 | promoter | tgggaaaaccgcactcgaaggcgcggagatttcccctacattcactctttggtgaaaaaaaatgcgccat ctagaaaattttacagacaacgtgatagcgttatgttatctccagcatacgccatcgcccgatttcattataaa cagggctgcgagcgacgtattttcggtcgcgggaggaatccaaatgtcaatgaaagcacatagccgcc | This work |
| pEryF-202 | promoter | gcactcgaaggcgcggagatttcccctacattcactctttggtgaaaaaaaatgcgccatctagaaaatttt acagacaacgtgatagcgttatgttatctccagcatacgccatcgcccgatttcattataaacagggctgcg agcgacgtattttcggtcgcgggaggaatccaaatgtcaatgaaagcacatagccgcc | This work |
| pEryF-192 | promoter | gcgcggagatttcccctacattcactctttggtgaaaaaaaatgcgccatctagaaaattttacagacaacg tgatagcgttatgttatctccagcatacgccatcgcccgatttcattataaacagggctgcgagcgacgtattt tcggtcgcgggaggaatccaaatgtcaatgaaagcacatagccgcc | This work |
| pEryF-182 | promoter | ttcccctacattcactctttggtgaaaaaaaatgcgccatctagaaaattttacagacaacgtgatagcgtta tgttatctccagcatacgccatcgcccgatttcattataaacagggctgcgagcgacgtattttcggtcgcgg gaggaatccaaatgtcaatgaaagcacatagccgcc | This work |
| pEryF-172 | promoter | ttcactctttggtgaaaaaaaatgcgccatctagaaaattttacagacaacgtgatagcgttatgttatctcca gcatacgccatcgcccgatttcattataaacagggctgcgagcgacgtattttcggtcgcgggaggaatcc aaatgtcaatgaaagcacatagccgcc | This work |
| pEryF-162 | promoter | ggtgaaaaaaaatgcgccatctagaaaattttacagacaacgtgatagcgttatgttatctccagcatacg ccatcgcccgatttcattataaacagggctgcgagcgacgtattttcggtcgcgggaggaatccaaatgtc aatgaaagcacatagccgcc | This work |
| pEryF-161 | promoter | cgtgaaaaaaaatgcgccatctagaaaattttacagacaacgtgatagcgttatgttatctccagcatacg ccatcgcccgatttcattataaacagggctgcgagcgacgtattttcggtcgcgggaggaatccaaatgtc aatgaaagcacatagccgcc | This work |
| pEryF-160 | promoter | cctgaaaaaaaatgcgccatctagaaaattttacagacaacgtgatagcgttatgttatctccagcatacg ccatcgcccgatttcattataaacagggctgcgagcgacgtattttcggtcgcgggaggaatccaaatgtc aatgaaagcacatagccgcc | This work |
| pEryF-159 | promoter | cccgaaaaaaaatgcgccatctagaaaattttacagacaacgtgatagcgttatgttatctccagcatacg ccatcgcccgatttcattataaacagggctgcgagcgacgtattttcggtcgcgggaggaatccaaatgtc aatgaaagcacatagccgcc | This work |
| pEryF-158 | promoter | ccctaaaaaaaatgcgccatctagaaaattttacagacaacgtgatagcgttatgttatctccagcatacgc catcgcccgatttcattataaacagggctgcgagcgacgtattttcggtcgcgggaggaatccaaatgtca atgaaagcacatagccgcc | This work |
| pEryF-157 | promoter | cccttaaaaaaatgcgccatctagaaaattttacagacaacgtgatagcgttatgttatctccagcatacgc catcgcccgatttcattataaacagggctgcgagcgacgtattttcggtcgcgggaggaatccaaatgtca atgaaagcacatagccgcc | This work |
| pEryF-156 | promoter | ccctttaaaaaatgcgccatctagaaaattttacagacaacgtgatagcgttatgttatctccagcatacgcc atcgcccgatttcattataaacagggctgcgagcgacgtattttcggtcgcgggaggaatccaaatgtcaa tgaaagcacatagccgcc | This work |

| | | | |
|---|---|---|---|
| pEryF-155 | promoter | ccctttcaaaaatgcgccatctagaaaattttacagacaacgtgatagcgttatgttatctccagcatacgcc<br>atcgcccgatttcattataaacagggctgcgagcgacgtattttcggtcgcgggaggaatccaaatgtcaa<br>tgaaagcacatagccgcc | This work |
| pEryF-154 | promoter | ccctttccaaaatgcgccatctagaaaattttacagacaacgtgatagcgttatgttatctccagcatacgcc<br>atcgcccgatttcattataaacagggctgcgagcgacgtattttcggtcgcgggaggaatccaaatgtcaa<br>tgaaagcacatagccgcc | This work |
| pEryF-153 | promoter | ccctttcccaaatgcgccatctagaaaattttacagacaacgtgatagcgttatgttatctccagcatacgcc<br>atcgcccgatttcattataaacagggctgcgagcgacgtattttcggtcgcgggaggaatccaaatgtcaa<br>tgaaagcacatagccgcc | This work |
| pEryF-152 | promoter | aatgcgccatctagaaaattttacagacaacgtgatagcgttatgttatctccagcatacgccatcgcccga<br>tttcattataaacagggctgcgagcgacgtattttcggtcgcgggaggaatccaaatgtcaatgaaagcac<br>atagccgcc | This work |
| pEryF-142 | promoter | ctagaaaattttacagacaacgtgatagcgttatgttatctccagcatacgccatcgcccgatttcattataaa<br>cagggctgcgagcgacgtattttcggtcgcgggaggaatccaaatgtcaatgaaagcacatagccgcc | This work |
| pEryF-132 | promoter | ttacagacaacgtgatagcgttatgttatctccagcatacgccatcgcccgatttcattataaacagggctgc<br>gagcgacgtattttcggtcgcgggaggaatccaaatgtcaatgaaagcacatagccgcc | This work |
| pEryF-122 | promoter | cgtgatagcgttatgttatctccagcatacgccatcgcccgatttcattataaacagggctgcgagcgacgt<br>attttcggtcgcgggaggaatccaaatgtcaatgaaagcacatagccgcc | This work |
| pEryF-112 | promoter | ttatgttatctccagcatacgccatcgcccgatttcattataaacagggctgcgagcgacgtattttcggtcgc<br>gggaggaatccaaatgtcaatgaaagcacatagccgcc | This work |
| pEryF-102 | promoter | tccagcatacgccatcgcccgatttcattataaacagggctgcgagcgacgtattttcggtcgcgggagga<br>atccaaatgtcaatgaaagcacatagccgcc | This work |
| pEryF-159-10 | promoter | tatcaaacgccttcagcaattctgccgtattgtcgttatggaaacggatgatgcggctgaattagaccgccc<br>ggagattaccctcgaccattgtcatcacgcttaccggataggcaaaagcctccgccaaagcggtgcagg<br>acgcgacagacgagatcatgatagcctccacatgaattccagggccccgctcctccgtgcatcctggaa<br>acagatgcgccgcaatcaaaacctatgatagcggcgtgttattaactccaataatgatgttataaagctcg<br>aaagctgtcaatcggatgttttcgaaacaccggtgaaatagcacaatttcgcgccgaatctcgatcacgat<br>ctagagaaatccactacataattcgttatcaatcgcactattcacagcaccgtcttttctcgcaaagccagcc<br>acaaagatactcacaaacttctatatttcttttattttcaatgaattaaacatatttaaaaccgaagaaacatta<br>agctattgatggaaaacagtgggaaaaccgcactcgaaggcgcggagatttcccctacattcactctttgg<br>tgaaaaaaaatgcgccatctagaaaattttacagacaacgtgatagcgttatgttatctccagcatacgcc<br>atcgcccgatttcattataaacagggctgcgagcgacgtattttcggtcgcgggaggaatccaaatgtcaa<br>tgaaagca | This work |
| pEryF-159-20 | promoter | tatcaaacgccttcagcaattctgccgtattgtcgttatggaaacggatgatgcggctgaattagaccgccc<br>ggagattaccctcgaccattgtcatcacgcttaccggataggcaaaagcctccgccaaagcggtgcagg<br>acgcgacagacgagatcatgatagcctccacatgaattccagggccccgctcctccgtgcatcctggaa<br>acagatgcgccgcaatcaaaacctatgatagcggcgtgttattaactccaataatgatgttataaagctcg<br>aaagctgtcaatcggatgttttcgaaacaccggtgaaatagcacaatttcgcgccgaatctcgatcacgat<br>ctagagaaatccactacataattcgttatcaatcgcactattcacagcaccgtcttttctcgcaaagccagcc<br>acaaagatactcacaaacttctatatttcttttattttcaatgaattaaacatatttaaaaccgaagaaacatta<br>agctattgatggaaaacagtgggaaaaccgcactcgaaggcgcggagatttcccctacattcactctttgg<br>tgaaaaaaaatgcgccatctagaaaattttacagacaacgtgatagcgttatgttatctccagcatacgcc<br>atcgcccgatttcattataaacagggctgcgagcgacgtattttcggtcgcgggaggaatccaaatgtc | This work |
| pEryF-159-30 | promoter | tatcaaacgccttcagcaattctgccgtattgtcgttatggaaacggatgatgcggctgaattagaccgccc<br>ggagattaccctcgaccattgtcatcacgcttaccggataggcaaaagcctccgccaaagcggtgcagg<br>acgcgacagacgagatcatgatagcctccacatgaattccagggccccgctcctccgtgcatcctggaa | This work |

| | | | |
|---|---|---|---|
| | | acagatgcgccgcaatcaaaacctatgatagcggcgtgttattaactccaataatgatgttataaagctcg<br>aaagctgtcaatcggatgttttcgaaacaccggtgaaatagcacaatttcgcgccgaatctcgatcacgat<br>ctagagaaatccactacataattcgttatcaatcgcactattcacagcaccgtcttttctcgcaaagccagcc<br>acaaagatactcacaaacttctatatttcttttattttcaatgaattaaacatatttaaaaccgaagaaacatta<br>agctattgatggaaaacagtgggaaaaccgcactcgaaggcgcggagatttcccctacattcactctttgg<br>tgaaaaaaaatgcgccatctagaaaattttacagacaacgtgatagcgttatgttatctccagcatacgcc<br>atcgcccgatttcattataaacagggctgcgagcgacgtattttcggtcgcgggaggaa | |
| pEryF-159-40 | promoter | tatcaaacgccttcagcaattctgccgtattgtcgttatggaaacggatgatgcggctgaattagaccgccc<br>ggagattaccctcgaccattgtcatcacgcttaccggataggcaaaagcctccgccaaagcggtgcagg<br>acgcgacagacgagatcatgatagcctccacatgaattccagggccccgctcctccgtgcatcctggaa<br>acagatgcgccgcaatcaaaacctatgatagcggcgtgttattaactccaataatgatgttataaagctcg<br>aaagctgtcaatcggatgttttcgaaacaccggtgaaatagcacaatttcgcgccgaatctcgatcacgat<br>ctagagaaatccactacataattcgttatcaatcgcactattcacagcaccgtcttttctcgcaaagccagcc<br>acaaagatactcacaaacttctatatttcttttattttcaatgaattaaacatatttaaaaccgaagaaacatta<br>agctattgatggaaaacagtgggaaaaccgcactcgaaggcgcggagatttcccctacattcactctttgg<br>tgaaaaaaaatgcgccatctagaaaattttacagacaacgtgatagcgttatgttatctccagcatacgcc<br>atcgcccgatttcattataaacagggctgcgagcgacgtattttcggtc | This work |
| pEryF-159-50 | promoter | tatcaaacgccttcagcaattctgccgtattgtcgttatggaaacggatgatgcggctgaattagaccgccc<br>ggagattaccctcgaccattgtcatcacgcttaccggataggcaaaagcctccgccaaagcggtgcagg<br>acgcgacagacgagatcatgatagcctccacatgaattccagggccccgctcctccgtgcatcctggaa<br>acagatgcgccgcaatcaaaacctatgatagcggcgtgttattaactccaataatgatgttataaagctcg<br>aaagctgtcaatcggatgttttcgaaacaccggtgaaatagcacaatttcgcgccgaatctcgatcacgat<br>ctagagaaatccactacataattcgttatcaatcgcactattcacagcaccgtcttttctcgcaaagccagcc<br>acaaagatactcacaaacttctatatttcttttattttcaatgaattaaacatatttaaaaccgaagaaacatta<br>agctattgatggaaaacagtgggaaaaccgcactcgaaggcgcggagatttcccctacattcactctttgg<br>tgaaaaaaaatgcgccatctagaaaattttacagacaacgtgatagcgttatgttatctccagcatacgcc<br>atcgcccgatttcattataaacagggctgcgagcgacgt | This work |
| pEryF-159-60 | promoter | tatcaaacgccttcagcaattctgccgtattgtcgttatggaaacggatgatgcggctgaattagaccgccc<br>ggagattaccctcgaccattgtcatcacgcttaccggataggcaaaagcctccgccaaagcggtgcagg<br>acgcgacagacgagatcatgatagcctccacatgaattccagggccccgctcctccgtgcatcctggaa<br>acagatgcgccgcaatcaaaacctatgatagcggcgtgttattaactccaataatgatgttataaagctcg<br>aaagctgtcaatcggatgttttcgaaacaccggtgaaatagcacaatttcgcgccgaatctcgatcacgat<br>ctagagaaatccactacataattcgttatcaatcgcactattcacagcaccgtcttttctcgcaaagccagcc<br>acaaagatactcacaaacttctatatttcttttattttcaatgaattaaacatatttaaaaccgaagaaacatta<br>agctattgatggaaaacagtgggaaaaccgcactcgaaggcgcggagatttcccctacattcactctttgg<br>tgaaaaaaaatgcgccatctagaaaattttacagacaacgtgatagcgttatgttatctccagcatacgcc<br>atcgcccgatttcattataaacagggctg | This work |
| pEryF-159-70 | promoter | tatcaaacgccttcagcaattctgccgtattgtcgttatggaaacggatgatgcggctgaattagaccgccc<br>ggagattaccctcgaccattgtcatcacgcttaccggataggcaaaagcctccgccaaagcggtgcagg<br>acgcgacagacgagatcatgatagcctccacatgaattccagggccccgctcctccgtgcatcctggaa<br>acagatgcgccgcaatcaaaacctatgatagcggcgtgttattaactccaataatgatgttataaagctcg<br>aaagctgtcaatcggatgttttcgaaacaccggtgaaatagcacaatttcgcgccgaatctcgatcacgat<br>ctagagaaatccactacataattcgttatcaatcgcactattcacagcaccgtcttttctcgcaaagccagcc<br>acaaagatactcacaaacttctatatttcttttattttcaatgaattaaacatatttaaaaccgaagaaacatta<br>agctattgatggaaaacagtgggaaaaccgcactcgaaggcgcggagatttcccctacattcactctttgg<br>tgaaaaaaaatgcgccatctagaaaattttacagacaacgtgatagcgttatgttatctccagcatacgcc | This work |

| | | | |
|---|---|---|---|
| | | atcgcccgatttcattata | |
| pEryF-159-80 | promoter | tatcaaacgccttcagcaattctgccgtattgtcgttatggaaacggatgatgcggctgaattagaccgccc<br>ggagattaccctcgaccattgtcatcacgcttaccggataggcaaaagcctccgccaaagcggtgcagg<br>acgcgacagacgagatcatgatagcctccacatgaattccagggccccgctcctccgtgcatcctggaa<br>acagatgcgccgcaatcaaaacctatgatagcggcgtgttattaactccaataatgatgttataaagctcg<br>aaagctgtcaatcggatgttttcgaaacaccggtgaaatagcacaatttcgcgccgaatctcgatcacgat<br>ctagagaaatccactacataattcgttatcaatcgcactattcacagcaccgtcttttctcgcaaagccagcc<br>acaaagatactcacaaacttctatattttcttttattttcaatgaattaaacatatttaaaaccgaagaaacatta<br>agctattgatggaaaacagtgggaaaaccgcactcgaaggcgcggagatttcccctacattcactctttgg<br>tgaaaaaaaatgcgccatctagaaaattttacagacaacgtgatagcgttatgttatctccagcatacgcc<br>atcgcccga | This work |
| pEryF-159-90 | promoter | tatcaaacgccttcagcaattctgccgtattgtcgttatggaaacggatgatgcggctgaattagaccgccc<br>ggagattaccctcgaccattgtcatcacgcttaccggataggcaaaagcctccgccaaagcggtgcagg<br>acgcgacagacgagatcatgatagcctccacatgaattccagggccccgctcctccgtgcatcctggaa<br>acagatgcgccgcaatcaaaacctatgatagcggcgtgttattaactccaataatgatgttataaagctcg<br>aaagctgtcaatcggatgttttcgaaacaccggtgaaatagcacaatttcgcgccgaatctcgatcacgat<br>ctagagaaatccactacataattcgttatcaatcgcactattcacagcaccgtcttttctcgcaaagccagcc<br>acaaagatactcacaaacttctatattttcttttattttcaatgaattaaacatatttaaaaccgaagaaacatta<br>agctattgatggaaaacagtgggaaaaccgcactcgaaggcgcggagatttcccctacattcactctttgg<br>tgaaaaaaaatgcgccatctagaaaattttacagacaacgtgatagcgttatgttatctccagcatacgc | This work |
| pEryF-159-100 | promoter | tatcaaacgccttcagcaattctgccgtattgtcgttatggaaacggatgatgcggctgaattagaccgccc<br>ggagattaccctcgaccattgtcatcacgcttaccggataggcaaaagcctccgccaaagcggtgcagg<br>acgcgacagacgagatcatgatagcctccacatgaattccagggccccgctcctccgtgcatcctggaa<br>acagatgcgccgcaatcaaaacctatgatagcggcgtgttattaactccaataatgatgttataaagctcg<br>aaagctgtcaatcggatgttttcgaaacaccggtgaaatagcacaatttcgcgccgaatctcgatcacgat<br>ctagagaaatccactacataattcgttatcaatcgcactattcacagcaccgtcttttctcgcaaagccagcc<br>acaaagatactcacaaacttctatattttcttttattttcaatgaattaaacatatttaaaaccgaagaaacatta<br>agctattgatggaaaacagtgggaaaaccgcactcgaaggcgcggagatttcccctacattcactctttgg<br>tgaaaaaaaatgcgccatctagaaaattttacagacaacgtgatagcgttatgttatctc | This work |
| pEryF-159-110 | promoter | tatcaaacgccttcagcaattctgccgtattgtcgttatggaaacggatgatgcggctgaattagaccgccc<br>ggagattaccctcgaccattgtcatcacgcttaccggataggcaaaagcctccgccaaagcggtgcagg<br>acgcgacagacgagatcatgatagcctccacatgaattccagggccccgctcctccgtgcatcctggaa<br>acagatgcgccgcaatcaaaacctatgatagcggcgtgttattaactccaataatgatgttataaagctcg<br>aaagctgtcaatcggatgttttcgaaacaccggtgaaatagcacaatttcgcgccgaatctcgatcacgat<br>ctagagaaatccactacataattcgttatcaatcgcactattcacagcaccgtcttttctcgcaaagccagcc<br>acaaagatactcacaaacttctatattttcttttattttcaatgaattaaacatatttaaaaccgaagaaacatta<br>agctattgatggaaaacagtgggaaaaccgcactcgaaggcgcggagatttcccctacattcactctttgg<br>tgaaaaaaaatgcgccatctagaaaattttacagacaacgtgatagcgtt | This work |
| pEryF-159-120 | promoter | tatcaaacgccttcagcaattctgccgtattgtcgttatggaaacggatgatgcggctgaattagaccgccc<br>ggagattaccctcgaccattgtcatcacgcttaccggataggcaaaagcctccgccaaagcggtgcagg<br>acgcgacagacgagatcatgatagcctccacatgaattccagggccccgctcctccgtgcatcctggaa<br>acagatgcgccgcaatcaaaacctatgatagcggcgtgttattaactccaataatgatgttataaagctcg<br>aaagctgtcaatcggatgttttcgaaacaccggtgaaatagcacaatttcgcgccgaatctcgatcacgat<br>ctagagaaatccactacataattcgttatcaatcgcactattcacagcaccgtcttttctcgcaaagccagcc<br>acaaagatactcacaaacttctatattttcttttattttcaatgaattaaacatatttaaaaccgaagaaacatta<br>agctattgatggaaaacagtgggaaaaccgcactcgaaggcgcggagatttcccctacattcactctttgg | This work |

| | | | |
|---|---|---|---|
| | | tgaaaaaaaatgcgccatctagaaaattttacagacaacg | |
| pEryF-159-130 | promoter | tatcaaacgccttcagcaattctgccgtattgtcgttatggaaacggatgatgcggctgaattagaccgccc<br>ggagattaccctcgaccattgtcatcacgcttaccggataggcaaaagcctccgccaaagcggtgcagg<br>acgcgacagacgagatcatgatagcctccacatgaattccagggccccgctcctccgtgcatcctggaa<br>acagatgcgccgcaatcaaaacctatgatagcggcgtgttattaactccaataatgatgttataaagctcg<br>aaagctgtcaatcggatgttttcgaaacaccggtgaaatagcacaatttcgcgccgaatctcgatcacgat<br>ctagagaaatccactacataattcgttatcaatcgcactattcacagcaccgtcttttctcgcaaagccagcc<br>acaaagatactcacaaacttctatatttcttttattttcaatgaattaaacatatttaaaaccgaagaaacatta<br>agctattgatggaaaacagtgggaaaaccgcactcgaaggcgcggagatttcccctacattcactctttgg<br>tgaaaaaaaatgcgccatctagaaaatttt | This work |
| pEryF-159-140 | promoter | tatcaaacgccttcagcaattctgccgtattgtcgttatggaaacggatgatgcggctgaattagaccgccc<br>ggagattaccctcgaccattgtcatcacgcttaccggataggcaaaagcctccgccaaagcggtgcagg<br>acgcgacagacgagatcatgatagcctccacatgaattccagggccccgctcctccgtgcatcctggaa<br>acagatgcgccgcaatcaaaacctatgatagcggcgtgttattaactccaataatgatgttataaagctcg<br>aaagctgtcaatcggatgttttcgaaacaccggtgaaatagcacaatttcgcgccgaatctcgatcacgat<br>ctagagaaatccactacataattcgttatcaatcgcactattcacagcaccgtcttttctcgcaaagccagcc<br>acaaagatactcacaaacttctatatttcttttattttcaatgaattaaacatatttaaaaccgaagaaacatta<br>agctattgatggaaaacagtgggaaaaccgcactcgaaggcgcggagatttcccctacattcactctttgg<br>tgaaaaaaaatgcgccatct | This work |
| pEryF-159-150 | promoter | tatcaaacgccttcagcaattctgccgtattgtcgttatggaaacggatgatgcggctgaattagaccgccc<br>ggagattaccctcgaccattgtcatcacgcttaccggataggcaaaagcctccgccaaagcggtgcagg<br>acgcgacagacgagatcatgatagcctccacatgaattccagggccccgctcctccgtgcatcctggaa<br>acagatgcgccgcaatcaaaacctatgatagcggcgtgttattaactccaataatgatgttataaagctcg<br>aaagctgtcaatcggatgttttcgaaacaccggtgaaatagcacaatttcgcgccgaatctcgatcacgat<br>ctagagaaatccactacataattcgttatcaatcgcactattcacagcaccgtcttttctcgcaaagccagcc<br>acaaagatactcacaaacttctatatttcttttattttcaatgaattaaacatatttaaaaccgaagaaacatta<br>agctattgatggaaaacagtgggaaaaccgcactcgaaggcgcggagatttcccctacattcactctttgg<br>tgaaaaaaaa | This work |
| pEryF-37 | promoter | tatcaaacgccttcagcaattctgccgtattgtcgttatggaaacggatgatgcggctgaattagaccgccc<br>ggagattaccctcgaccattgtcatcacgcttaccggataggcaaaagcctccgccaaagcggtgcagg<br>acgcgacagacgagatcatgatagcctccacatgaattccagggccccgctcctccgtgcatcctggaa<br>acagatgcgccgcaatcaaaacctatgatagcggcgtgttattaactccaataatgatgttataaagctcg<br>aaagctgtcaatcggatgttttcgaaacaccggtgaaatagcacaatttcgcgccgaatctcgatcacgat<br>ctagagaaatccactacataattcgttatcaatcgcactattcacagcaccgtcttttctcgcaaagccagcc<br>acaaagatactcacaaacttctatatttcttttattttcaatgaattaaacatatttaaaaccgaagaaacatta<br>agctattgatggaaaacagtgggaaaaccgcactcgaaggcgcggagatttcccctacattcactctttgg<br>tgaaaaaaaatgcgccatctagaaaattttacagacaagctgatagcgttatgttatctc | This work |
| pEryF-35 | promoter | tatcaaacgccttcagcaattctgccgtattgtcgttatggaaacggatgatgcggctgaattagaccgccc<br>ggagattaccctcgaccattgtcatcacgcttaccggataggcaaaagcctccgccaaagcggtgcagg<br>acgcgacagacgagatcatgatagcctccacatgaattccagggccccgctcctccgtgcatcctggaa<br>acagatgcgccgcaatcaaaacctatgatagcggcgtgttattaactccaataatgatgttataaagctcg<br>aaagctgtcaatcggatgttttcgaaacaccggtgaaatagcacaatttcgcgccgaatctcgatcacgat<br>ctagagaaatccactacataattcgttatcaatcgcactattcacagcaccgtcttttctcgcaaagccagcc<br>acaaagatactcacaaacttctatatttcttttattttcaatgaattaaacatatttaaaaccgaagaaacatta<br>agctattgatggaaaacagtgggaaaaccgcactcgaaggcgcggagatttcccctacattcactctttgg<br>tgaaaaaaaatgcgccatctagaaaattttacagacttgctgatagcgttatgttatctc | This work |

| | | | |
|---|---|---|---|
| pEryF-33 | promoter | tatcaaacgccttcagcaattctgccgtattgtcgttatggaaacggatgatgcggctgaattagaccgccc<br>ggagattaccctcgaccattgtcatcacgcttaccggataggcaaaagcctccgccaaagcggtgcagg<br>acgcgacagacgagatcatgatagcctccacatgaattccagggccccgctcctccgtgcatcctggaa<br>acagatgcgccgcaatcaaaacctatgatagcggcgtgttattaactccaataatgatgttataaagctcg<br>aaagctgtcaatcggatgttttcgaaacaccggtgaaatagcacaatttcgcgccgaatctcgatcacgat<br>ctagagaaatccactacataattcgttatcaatcgcactattcacagcaccgtcttttctcgcaaagccagcc<br>acaaagatactcacaaacttctatattttcttttattttcaatgaattaaacatatttaaaaccgaagaaacatta<br>agctattgatggaaaacagtgggaaaaccgcactcgaaggcgcggagatttcccctacattcactctttgg<br>tgaaaaaaaatgcgccatctagaaaattttacagtgttgctgatagcgttatgttatctc | This work |
| pEryF-31 | promoter | tatcaaacgccttcagcaattctgccgtattgtcgttatggaaacggatgatgcggctgaattagaccgccc<br>ggagattaccctcgaccattgtcatcacgcttaccggataggcaaaagcctccgccaaagcggtgcagg<br>acgcgacagacgagatcatgatagcctccacatgaattccagggccccgctcctccgtgcatcctggaa<br>acagatgcgccgcaatcaaaacctatgatagcggcgtgttattaactccaataatgatgttataaagctcg<br>aaagctgtcaatcggatgttttcgaaacaccggtgaaatagcacaatttcgcgccgaatctcgatcacgat<br>ctagagaaatccactacataattcgttatcaatcgcactattcacagcaccgtcttttctcgcaaagccagcc<br>acaaagatactcacaaacttctatattttcttttattttcaatgaattaaacatatttaaaaccgaagaaacatta<br>agctattgatggaaaacagtgggaaaaccgcactcgaaggcgcggagatttcccctacattcactctttgg<br>tgaaaaaaaatgcgccatctagaaaattttactctgttgctgatagcgttatgttatctc | This work |
| pEryF-29 | promoter | tatcaaacgccttcagcaattctgccgtattgtcgttatggaaacggatgatgcggctgaattagaccgccc<br>ggagattaccctcgaccattgtcatcacgcttaccggataggcaaaagcctccgccaaagcggtgcagg<br>acgcgacagacgagatcatgatagcctccacatgaattccagggccccgctcctccgtgcatcctggaa<br>acagatgcgccgcaatcaaaacctatgatagcggcgtgttattaactccaataatgatgttataaagctcg<br>aaagctgtcaatcggatgttttcgaaacaccggtgaaatagcacaatttcgcgccgaatctcgatcacgat<br>ctagagaaatccactacataattcgttatcaatcgcactattcacagcaccgtcttttctcgcaaagccagcc<br>acaaagatactcacaaacttctatattttcttttattttcaatgaattaaacatatttaaaaccgaagaaacatta<br>agctattgatggaaaacagtgggaaaaccgcactcgaaggcgcggagatttcccctacattcactctttgg<br>tgaaaaaaaatgcgccatctagaaaattttgtctgttgctgatagcgttatgttatctc | This work |
| pEryF-28 | promoter | tatcaaacgccttcagcaattctgccgtattgtcgttatggaaacggatgatgcggctgaattagaccgccc<br>ggagattaccctcgaccattgtcatcacgcttaccggataggcaaaagcctccgccaaagcggtgcagg<br>acgcgacagacgagatcatgatagcctccacatgaattccagggccccgctcctccgtgcatcctggaa<br>acagatgcgccgcaatcaaaacctatgatagcggcgtgttattaactccaataatgatgttataaagctcg<br>aaagctgtcaatcggatgttttcgaaacaccggtgaaatagcacaatttcgcgccgaatctcgatcacgat<br>ctagagaaatccactacataattcgttatcaatcgcactattcacagcaccgtcttttctcgcaaagccagcc<br>acaaagatactcacaaacttctatattttcttttattttcaatgaattaaacatatttaaaaccgaagaaacatta<br>agctattgatggaaaacagtgggaaaaccgcactcgaaggcgcggagatttcccctacattcactctttgg<br>tgaaaaaaaatgcgccatctagaaaatttATGTCTGTTGCtgatagcgttatgttatctc | This work |
| pEryF-27 | promoter | tatcaaacgccttcagcaattctgccgtattgtcgttatggaaacggatgatgcggctgaattagaccgccc<br>ggagattaccctcgaccattgtcatcacgcttaccggataggcaaaagcctccgccaaagcggtgcagg<br>acgcgacagacgagatcatgatagcctccacatgaattccagggccccgctcctccgtgcatcctggaa<br>acagatgcgccgcaatcaaaacctatgatagcggcgtgttattaactccaataatgatgttataaagctcg<br>aaagctgtcaatcggatgttttcgaaacaccggtgaaatagcacaatttcgcgccgaatctcgatcacgat<br>ctagagaaatccactacataattcgttatcaatcgcactattcacagcaccgtcttttctcgcaaagccagcc<br>acaaagatactcacaaacttctatattttcttttattttcaatgaattaaacatatttaaaaccgaagaaacatta<br>agctattgatggaaaacagtgggaaaaccgcactcgaaggcgcggagatttcccctacattcactctttgg<br>tgaaaaaaaatgcgccatctagaaaattaatgtctgttgctgatagcgttatgttatctc | This work |
| pEryF-25 | promoter | tatcaaacgccttcagcaattctgccgtattgtcgttatggaaacggatgatgcggctgaattagaccgccc | This work |

| | | | |
|---|---|---|---|
| | | ggagattaccctcgaccattgtcatcacgcttaccggataggcaaaagcctccgccaaagcggtgcagg<br>acgcgacagacgagatcatgatagcctccacatgaattccagggccccgctcctccgtgcatcctggaa<br>acagatgcgccgcaatcaaaacctatgatagcggcgtgttattaactccaataatgatgttataaagctcg<br>aaagctgtcaatcggatgttttcgaaacaccggtgaaatagcacaatttcgcgccgaatctcgatcacgat<br>ctagagaaatccactacataattcgttatcaatcgcactattcacagcaccgtcttttctcgcaaagccagcc<br>acaaagatactcacaaacttctatattttcttttattttcaatgaattaaacatatttaaaaccgaagaaacatta<br>agctattgatggaaaacagtgggaaaaccgcactcgaaggcgcggagatttcccctacattcactctttgg<br>tgaaaaaaaatgcgccatctagaaaaaaaatgtctgttgctgatagcgttatgttatctc | |
| pEryF-23 | promoter | tatcaaacgccttcagcaattctgccgtattgtcgttatggaaacggatgatgcggctgaattagaccgccc<br>ggagattaccctcgaccattgtcatcacgcttaccggataggcaaaagcctccgccaaagcggtgcagg<br>acgcgacagacgagatcatgatagcctccacatgaattccagggccccgctcctccgtgcatcctggaa<br>acagatgcgccgcaatcaaaacctatgatagcggcgtgttattaactccaataatgatgttataaagctcg<br>aaagctgtcaatcggatgttttcgaaacaccggtgaaatagcacaatttcgcgccgaatctcgatcacgat<br>ctagagaaatccactacataattcgttatcaatcgcactattcacagcaccgtcttttctcgcaaagccagcc<br>acaaagatactcacaaacttctatattttcttttattttcaatgaattaaacatatttaaaaccgaagaaacatta<br>agctattgatggaaaacagtgggaaaaccgcactcgaaggcgcggagatttcccctacattcactctttgg<br>tgaaaaaaaatgcgccatctagaattaaaatgtctgttgctgatagcgttatgttatctc | This work |
| pEryF-21 | promoter | tatcaaacgccttcagcaattctgccgtattgtcgttatggaaacggatgatgcggctgaattagaccgccc<br>ggagattaccctcgaccattgtcatcacgcttaccggataggcaaaagcctccgccaaagcggtgcagg<br>acgcgacagacgagatcatgatagcctccacatgaattccagggccccgctcctccgtgcatcctggaa<br>acagatgcgccgcaatcaaaacctatgatagcggcgtgttattaactccaataatgatgttataaagctcg<br>aaagctgtcaatcggatgttttcgaaacaccggtgaaatagcacaatttcgcgccgaatctcgatcacgat<br>ctagagaaatccactacataattcgttatcaatcgcactattcacagcaccgtcttttctcgcaaagccagcc<br>acaaagatactcacaaacttctatattttcttttattttcaatgaattaaacatatttaaaaccgaagaaacatta<br>agctattgatggaaaacagtgggaaaaccgcactcgaaggcgcggagatttcccctacattcactctttgg<br>tgaaaaaaaatgcgccatctagttttaaaatgtctgttgctgatagcgttatgttatctc | This work |
| pEryR-732 | promoter | ggcggctatgtgctttcattgacatttggattcctcccgcgaccgaaaatacgtcgctcgcagccctgtttata<br>atgaaatcgggcgatggcgtatgctggagataacataacgctatcacgttgtctgtaaaattttctagatggc<br>gcatttttttcaccaaagagtgaatgtaggggaaatctccgcgccttcgagtgcggttttcccactgttttccat<br>caatagcttaatgtttcttcggttttaaatatgtttaattcattgaaaataaaagaaaatatagaagtttgtgagt<br>atctttgtggctggctttgcgagaaaagacggtgctgtgaatagtgcgattgataacgaattatgtagtggatt<br>tctctagatcgtgatcgagattcggcgcgaaattgtgctatttcaccggtgtttcgaaaacatccgattgaca<br>gctttcgagctttataacatcattattggagttaataacacgccgctatcataggttttgattgcggcgcatctgt<br>ttccaggatgcacggaggagcggggccctggaattcatgtggaggctatcatgatctcgtctgtcgcgtcct<br>gcaccgctttggcggaggcttttgcctatccggtaagcgtgatgacaatggtcgagggtaatctccgggcg<br>gtctaattcagccgcatcatccgtttccataacgacaatacggcagaattgctgaaggcgtttgata | This work |
| pEryR-632 | promoter | gagataacataacgctatcacgttgtctgtaaaattttctagatggcgcatttttttcaccaaagagtgaatgt<br>aggggaaatctccgcgccttcgagtgcggttttcccactgttttccatcaatagcttaatgtttcttcggttttaaa<br>tatgtttaattcattgaaaataaaagaaaatatagaagtttgtgagtatctttgtggctggctttgcgagaaaa<br>gacggtgctgtgaatagtgcgattgataacgaattatgtagtggatttctctagatcgtgatcgagattcggc<br>gcgaaattgtgctatttcaccggtgtttcgaaaacatccgattgacagctttcgagctttataacatcattattg<br>gagttaataacacgccgctatcataggttttgattgcggcgcatctgtttccaggatgcacggaggagcgg<br>ggccctggaattcatgtggaggctatcatgatctcgtctgtcgcgtcctgcaccgctttggcggaggcttttgc<br>ctatccggtaagcgtgatgacaatggtcgagggtaatctccgggcggtctaattcagccgcatcatccgttt<br>ccataacgacaatacggcagaattgctgaaggcgtttgata | This work |
| pEryR-532 | promoter | gcggttttcccactgttttccatcaatagcttaatgtttcttcggttttaaatatgtttaattcattgaaaataaaaga | This work |

| | | | |
|---|---|---|---|
| | | aaatatagaagtttgtgagtatctttgtggctggctttgcgagaaaagacggtgctgtgaatagtgcgattgataacgaattatgtagtggatttctctagatcgtgatcgagattcggcgcgaaattgtgctatttcaccggtgtttcgaaaacatccgattgacagctttcgagctttataacatcattattggagttaataacacgccgctatcataggttttgattgcggcgcatctgtttccaggatgcacggaggagcggggccctggaattcatgtggaggctatcatgatctcgtctgtcgcgtcctgcaccgctttggcggaggcttttgcctatccggtaagcgtgatgacaatggtcgagggtaatctccgggcggtctaattcagccgcatcatccgtttccataacgacaatacggcagaattgctgaaggcgtttgata | |
| pEryR-432 | promoter | tctttgtggctggctttgcgagaaaagacggtgctgtgaatagtgcgattgataacgaattatgtagtggatttctctagatcgtgatcgagattcggcgcgaaattgtgctatttcaccggtgtttcgaaaacatccgattgacagctttcgagctttataacatcattattggagttaataacacgccgctatcataggttttgattgcggcgcatctgtttccaggatgcacggaggagcggggccctggaattcatgtggaggctatcatgatctcgtctgtcgcgtcctgcaccgctttggcggaggcttttgcctatccggtaagcgtgatgacaatggtcgagggtaatctccgggcggtctaattcagccgcatcatccgtttccataacgacaatacggcagaattgctgaaggcgtttgata | This work |
| pEryR-332 | promoter | cgaaattgtgctatttcaccggtgtttcgaaaacatccgattgacagctttcgagctttataacatcattattggagttaataacacgccgctatcataggttttgattgcggcgcatctgtttccaggatgcacggaggagcggggccctggaattcatgtggaggctatcatgatctcgtctgtcgcgtcctgcaccgctttggcggaggcttttgcctatccggtaagcgtgatgacaatggtcgagggtaatctccgggcggtctaattcagccgcatcatccgtttccataacgacaatacggcagaattgctgaaggcgtttgata | This work |
| pEryR-322 | promoter | ctatttcaccggtgtttcgaaaacatccgattgacagctttcgagctttataacatcattattggagttaataacacgccgctatcataggttttgattgcggcgcatctgtttccaggatgcacggaggagcggggccctggaattcatgtggaggctatcatgatctcgtctgtcgcgtcctgcaccgctttggcggaggcttttgcctatccggtaagcgtgatgacaatggtcgagggtaatctccgggcggtctaattcagccgcatcatccgtttccataacgacaatacggcagaattgctgaaggcgtttgata | This work |
| pEryR-312 | promoter | ggtgtttcgaaaacatccgattgacagctttcgagctttataacatcattattggagttaataacacgccgctatcataggttttgattgcggcgcatctgtttccaggatgcacggaggagcggggccctggaattcatgtggaggctatcatgatctcgtctgtcgcgtcctgcaccgctttggcggaggcttttgcctatccggtaagcgtgatgacaatggtcgagggtaatctccgggcggtctaattcagccgcatcatccgtttccataacgacaatacggcagaattgctgaaggcgtttgata | This work |
| pEryR-302 | promoter | aaacatccgattgacagctttcgagctttataacatcattattggagttaataacacgccgctatcataggttttgattgcggcgcatctgtttccaggatgcacggaggagcggggccctggaattcatgtggaggctatcatgatctcgtctgtcgcgtcctgcaccgctttggcggaggcttttgcctatccggtaagcgtgatgacaatggtcgagggtaatctccgggcggtctaattcagccgcatcatccgtttccataacgacaatacggcagaattgctgaaggcgtttgata | This work |
| pEryR-292 | promoter | ttgacagctttcgagctttataacatcattattggagttaataacacgccgctatcataggttttgattgcggcgcatctgtttccaggatgcacggaggagcggggccctggaattcatgtggaggctatcatgatctcgtctgtcgcgtcctgcaccgctttggcggaggcttttgcctatccggtaagcgtgatgacaatggtcgagggtaatctccgggcggtctaattcagccgcatcatccgtttccataacgacaatacggcagaattgctgaaggcgtttgata | This work |
| pEryR-282 | promoter | tcgagctttataacatcattattggagttaataacacgccgctatcataggttttgattgcggcgcatctgtttccaggatgcacggaggagcggggccctggaattcatgtggaggctatcatgatctcgtctgtcgcgtcctgcaccgctttggcggaggcttttgcctatccggtaagcgtgatgacaatggtcgagggtaatctccgggcggtctaattcagccgcatcatccgtttccataacgacaatacggcagaattgctgaaggcgtttgata | This work |
| pEryR-272 | promoter | taacatcattattggagttaataacacgccgctatcataggttttgattgcggcgcatctgtttccaggatgcacggaggagcggggccctggaattcatgtggaggctatcatgatctcgtctgtcgcgtcctgcaccgctttggcggaggcttttgcctatccggtaagcgtgatgacaatggtcgagggtaatctccgggcggtctaattcagccgcatcatccgtttccataacgacaatacggcagaattgctgaaggcgtttgata | This work |

| | | | |
|---|---|---|---|
| pEryR-262 | promoter | attggagttaataacacgccgctatcataggttttgattgcggcgcatctgtttccaggatgcacggaggagcggggccctggaattcatgtggaggctatcatgatctcgtctgtcgcgtcctgcaccgctttggcggaggcttttgcctatccggtaagcgtgatgacaatggtcgagggtaatctccgggcggtctaattcagccgcatcatccgtttccataacgacaatacggcagaattgctgaaggcgtttgata | This work |
| pEryR-252 | promoter | ataacacgccgctatcataggttttgattgcggcgcatctgtttccaggatgcacggaggagcggggccctggaattcatgtggaggctatcatgatctcgtctgtcgcgtcctgcaccgctttggcggaggcttttgcctatccggtaagcgtgatgacaatggtcgagggtaatctccgggcggtctaattcagccgcatcatccgtttccataacgacaatacggcagaattgctgaaggcgtttgata | This work |
| pEryR-242 | promoter | gctatcataggttttgattgcggcgcatctgtttccaggatgcacggaggagcggggccctggaattcatgtggaggctatcatgatctcgtctgtcgcgtcctgcaccgctttggcggaggcttttgcctatccggtaagcgtgatgacaatggtcgagggtaatctccgggcggtctaattcagccgcatcatccgtttccataacgacaatacggcagaattgctgaaggcgtttgata | This work |
| pEryR-232 | promoter | gttttgattgcggcgcatctgtttccaggatgcacggaggagcggggccctggaattcatgtggaggctatcatgatctcgtctgtcgcgtcctgcaccgctttggcggaggcttttgcctatccggtaagcgtgatgacaatggtcgagggtaatctccgggcggtctaattcagccgcatcatccgtttccataacgacaatacggcagaattgctgaaggcgtttgata | This work |
| pEryR-132 | promoter | ctttggcggaggcttttgcctatccggtaagcgtgatgacaatggtcgagggtaatctccgggcggtctaattcagccgcatcatccgtttccataacgacaatacggcagaattgctgaaggcgtttgata | This work |
| pEryR-32 | promoter | caatacggcagaattgctgaaggcgtttgata | This work |
| pEry2-400 | promoter | acgccaatggccagcgccttgaaaccgcgctgaccgcccgcaccatcgctgcatcagtcgaaaacgcggatatgagccgaatcgtgggccttgccggtggaatttccaaggcggaagccatccgcgctgtgctgaaaagcggtcgtctttacggtctcatcaccgatgaacgcacggcaaagacgctcgttgagcagccgaacggaaaataacataataacatttttgcaatggattattttgtgattttgtgatctaacttcgcgcagaaaatgttccgccaataagatcggacgagagcgtgaagcgcgacgacagaagacaggcgattataaaccttctgatcgaaaatcaggccgtcgacctcgacgatctggcggatcggtttgccgtatcaaaa | This work |
| pEry2-300 | promoter | aatttccaaggcggaagccatccgcgctgtgctgaaaagcggtcgtctttacggtctcatcaccgatgaacgcacggcaaagacgctcgttgagcagccgaacggaaaataacataataacatttttgcaatggattattttgtgattttgtgatctaacttcgcgcagaaaatgttccgccaataagatcggacgagagcgtgaagcgcgacgacagaagacaggcgattataaaccttctgatcgaaaatcaggccgtcgacctcgacgatctggcggatcggtttgccgtatcaaaa | This work |
| pEry2-200 | promoter | aacggaaaataacataataacatttttgcaatggattattttgtgattttgtgatctaacttcgcgcagaaaatgttccgccaataagatcggacgagagcgtgaagcgcgacgacagaagacaggcgattataaaccttctgatcgaaaatcaggccgtcgacctcgacgatctggcggatcggtttgccgtatcaaaa | This work |
| pEry2-100 | promoter | cgtgaagcgcgacgacagaagacaggcgattataaaccttctgatcgaaaatcaggccgtcgacctcgacgatctggcggatcggtttgccgtatcaaaa | This work |
| B0034 | RBS | aaagaggagaaa | 1, BBa_B0034 |
| eryA-RBS | RBS | ggaatccaaatgtcaatgaaagcacatagccgcc | This work |
| eryB-RBS | RBS | agactatgggatgcaattgcatccgaaggagcatgaa | This work |
| eryC-RBS | RBS | ggaatggttcgacaacgctaaaatggtctcttgaggacact | This work |
| eryD-RBS | RBS | gaagatataaacgatttcaggaaccggaccgctttca | This work |
| eryE-RBS | RBS | aacgacaatacggcagaattgctgaaggcgtttgata | This work |
| eryF-RBS | RBS | gaagaactggctcgtcacgcctgtggggctcgaagtgaa | This work |
| eryG-RBS | RBS | aaaaggatagagcataagacccgacaaagtcgggttgttaa | This work |
| eryH-RBS | RBS | gctgacgactgcgcataaacctagaggacgaaaatcgagc | This work |
| eryI-RBS | RBS | ggccaagtgtgccgccaagactcgagctaattaaggagcaat | This work |

| | | | |
|---|---|---|---|
| pR-eryA | promoter-RBS | tatcaaacgccttcagcaattctgccgtattgtcgttatggaaacggatgatgcggctgaattagaccgcccggagattaccctcgaccattgtcatcacgcttaccggataggcaaaagcctccgccaaagcggtgcaggacgcgacagacgagatcatgatagcctccacatgaattccagggccccgctcctccgtgcatcctggaaacagatgcgccgcaatcaaaacctatgatagcggcgtgttattaactccaataatgatgttataaagctcgaaagctgtcaatcggatgttttcgaaacaccggtgaaatagcacaatttcgcgccgaatctcgatcacgatctagagaaatccactacataattcgttatcaatcgcactattcacagcaccgtcttttctcgcaaagccagccacaaagatactcacaaacttctatattttcttttattttcaatgaattaaacatatttaaaaccgaagaaacattaagctattgatggaaaacagtgggaaaaccgcactcgaaggcgcggagatttcccctacattcactctttggtgaaaaaaaatgcgccatctagaaaatttacagacaacgtgatagcgttatgttatctccagcatacgccatcgcccgatttcattataaacagggctgcgagcgacgtattttcggtcgcgggaggaatccaaatgtcaatgaaagcacatagccgcc | This work |
| pR-eryB | promoter-RBS | cgtgagaaaggtgacatcatcatcgggatcgacgcgggcacgtctgtgctcaaggcagtcgctttcgaccttagcggacgccagatcgaatctgctgccgttcgcaataaatacgataccggcgaacatggcgctgtcacacagtcgctttcccagacgtggcaggactgcgcgcgcgccttgcgcgacctcggtgcaaagatgcccggccttgccgaacgcactgcggcaatcgccgttaccggtcagggtgacggaacctggcttgttggccgcgacaaccagccggtcggcgatgcctggctgtggctcgatgcccgcgcggcaacgacggtgacgcgcctcgccgctggccccgtgaaccgcgcgcgctttgaagcgaccggcactgggttgaacacctgccagcaaggctcgcagatggcgcatatggacagcatcgcgcccgagctgctggacaatgccgaagtcgcgctccactgcaaagactggctctatctcaatctcaccggcgttcgcgccaccgacccttcggaagcaagcttcaccttcggcaatttccgcaatcgccagtatgacgatgtcgtcatcgacgcgctcggtctgaggaaacggcgtggattgctgccggaaatcatcgacggcaccgaagtgcagcatccgctatcccccgaagcggcagctgcaacgggtttgcgggcagggacgccggtgggtctcgcctatgtcgatatggccatgacagcacttggtgcgggcgtgcgcggcggcacagcaggcgccgggtgctcgaccattggttcgacaggcgtccatatgcgcgccaagccggttgccgatatccacctgaacaaggaaggcaccggttacgtcatcgcgctgcccattcccggcatcgtcacccaggtgcagaccaatatgggggcaacgatcaacatcgactggatattgcaggttgctgccgatctcatgtcgacgccagacaagcctgtttcgctcagtgacctcattccccgccttgatgactggtttaatgccagccgtccgggtgcaatcctctatcatccgtatatttcggaagcgggcgaacgcggtcccttcgtcaatgcctatgcccggccggtttcgtcggtctttccagccgcgaccgtttcccggaaatggtgcgctcggttgtcgagggcctcggaatggcgacgcgcgattgctacacggccatgggtgaaatgcccgcagaattgcgcatcaccggcggtgcagcgcgttcaaaagcattgcgcagcacccttgccgcagcggtgaatgcgcccgtgcgcgtttcctcgcgtgaagaggcgggagccgcaggtgctgccatgatggctgcggtggccattggcgcttatccaagcatggatgcatgtattgcggaatgggtcgagccgctgcttggcccttgcgaggccccggatggtccgcgcgcaaagcactatgaagaactcttcgttgcctatcaggaagcccggctggcactgacgcccgtctgggacaagttggcttccgatagagcatttccagcaaaagcgtgaagcggttttgcgttcggaaatgcggtgaacaaaacgaataaggccggaccttccttccggcacatatagactatgggatgcaattgcatccgaaggagcatgaa | This work |
| pR-eryC | promoter-RBS | gctgaaccggaaacctgcgatctctttgtcatcggcggtggtatcaacggcgcgggcgtggcccgtgacgccgccgggcgcggcctcaaggtggtgctggcggaaaaagacgatctggcgcagggaacttcgtcacgctccggcaagctggtacatggtggtctgcgttatctcgaatattatgagtttcgccttgtgcgcgaagcgctgatcgagcgcgaagtgctgttgaacgccgctccccatatcatctggccaatgcgtttcgttctgccgcacagcccgcaggaccgtcccgcctggctggtgcggcttggcctgtttctctatgaccacctcggcggccgcaagaagcttcccggcacacgcacgctcgacctgcggcgcgatccggaaggcacgccgatcctcgaccaatataccaagggcttcgaatattccgactgttgggtcgatgatgcccgtctcgttgcattgaatgcggttggcgctgctgaaaaaggcgcgaccattctgacccgcacgcctgtcgtctccgctcgtcgcgaaaaaggcggctggatcgtggaaacgaaaaaccgcgacacgggcgaaacccgcacattccgcgcgcgctgcatcgtcaattgcgccggaccatgggtcacggatgtcatccacaatgtcgccggctccaactcgtcgcgcaatgtgcgtctcgtcaagggcagccacatcatcgttccgaaattctggtcgggcgcaaatgcctatctcgtccagaaccacgaca | This work |

| | | | |
|---|---|---|---|
| | | agcgcgttatctttatcaatccctacgagggcgacaaggcgctgatcggcaccaccgacatcgcctatga<br>aggccgtgccgaagatgttgcagcagatgagaaggaaatcgaatatctgctgactgcagtgaaccgcta<br>tttcaaggaaaagctcaggcgtgaagacgtgctgcattccttctccggtgtgcgtccgctgttcgacgacgg<br>caagggcaacccttccgccgtcacccgcgactatgtcttcgacctcgatgaaaccaacgacgcgccgct<br>gctcaacgtctttggcggcaagatcaccaccttccgcgagctggccgagcgcggcatgcatcgcctcaa<br>gcacattttcccgaagatgggcggcgactggactcatggcgcaccgcttccgggcggcgaaatcgccaa<br>tgccgattatgaaaccttcgccaactccttgcgcgacacctatccgtggatgccgcgcgcgctcgtccaac<br>actacggtcgtctctacggcgcacgcacgaaagacgttgttgcaggcgcacagaacctcgaagggctc<br>ggccgtcatttcggcggcaatttccatgaggcggaagtgcgctatctggtggccagggaatgggcaatga<br>cggcggaagacattctctatcgccgcaccaagcactatctgcacctgaccgaagccgaacgcgccgctt<br>tcgtggaatggttcgacaacgctaaaatggtctcttgaggacact | |
| pR-eryD | promoter-RBS | gcccttacgctttctctgaacaccaatccacttgtgaaccgctttgccgaaccggatgacctgattgaaacg<br>gttgcgcgcgatctgcgcctgcgcgatctccagctcacacacgagttcatcaatccaagctggcaggctc<br>caaccatccgccgcctgacgcgcgacatggacaaggccttgcagcgtaccggtgttcgcgtcacttcgg<br>gcatgaccggcccctatggccgcctcaaccatttcggccatccggacgcagaagtgcgccgttactatgt<br>cgactggttcaagaccttcgccgacattatcggcgatcttggcggcaagtcggtcggcacgcagtttgcga<br>tcttcacctataaggatttcgatgacgctgcgcgcccgcgaagatctgatcaagatcgccatcgattgctggg<br>cggaagtcgccgaacatgcgagccgtgccggtctcgactatgtgttctgggaaccgatgagcatcgggc<br>gtgaatttggcgagacgattgccgaatgcatgaagcttcaggatcgcctgaccgccgccgacatggctatt<br>ccgatgtggatgatggccgatatcgaccacggcgatgtgacttccgccaatccggacgatttcgatcccta<br>cgcctgggcacgcgccgtgccgaaagtctcgccgatcattcacatcaagcagagcctgatggacaagg<br>gcggacatcgtcctttcacagccgcgttcaatgccaagggacgcatccagccggaaccgcttctgaaag<br>cctttgccgaaggcggcgctgtggacaatgaaatctgcctcgaactgtcgttcaaggagcgcgagccga<br>atgaccgtgaagtcattccgcagattgcggaaagtgtggctttctgggctccgcacattgacaccggcgct<br>aaggacttgaagatataaacgatttcaggaaccggaccgctttca | This work |
| pR-eryE | promoter-RBS | ggcggctatgtgctttcattgacatttggattcctcccgcgaccgaaaatacgtcgctcgcagccctgtttata<br>atgaaatcgggcgatggcgtatgctggagataacataacgctatcacgttgtctgtaaaattttctagatggc<br>gcatttttttcaccaaagagtgaatgtaggggaaatctccgcgccttcgagtgcggttttcccactgttttccat<br>caatagcttaatgtttcttcggttttaaatatgtttaattcattgaaaataaaagaaaatatagaagtttgtgagt<br>atctttgtggctggctttgcgagaaaagacggtgctgtgaatagtgcgattgataacgaattatgtagtggatt<br>tctctagatcgtgatcgagattcggcgcgaaattgtgctatttcaccggtgtttcgaaaacatccgattgaca<br>gctttcgagctttataacatcattattggagttaataacacgccgctatcataggttttgattgcggcgcatctgt<br>ttccaggatgcacggaggagcggggccctggaattcatgtggaggctatcatgatctcgtctgtcgcgtcct<br>gcaccgctttggcggaggcttttgcctatccggtaagcgtgatgacaatggtcgagggtaatctccgggcg<br>gtctaattcagccgcatcatccgtttccataacgacaatacggcagaattgctgaaggcgtttgata | This work |
| pR-eryF | promoter-RBS | agtgcacaaaccagtcaagtttcacaaaaacccgccaagctgagccgcgccgtcgtctggagcgtgatc<br>gctgcgatcgtggttgtcggtgcgattgccttcgacacgaaggtcgtgaagatcgggtctgattccgatgtg<br>cgccagcaggctttctcccccgatgcctacggtgcttccgaattccccaaggtgaaggcgagcgttgagc<br>agcgcgccgtggatgcagtggaagttggaacggctctcgccgccgacaaggccgccgccggaaaga<br>aatatggcgtcggcgacgtcaatccggttgtcccggtcaagtttacaggcaccgttgaggagcgcaaatc<br>caactacaatgtcgtgaaggtcgacggcctgccggaaggcgttgccatccgcgttcagaccggccctgc<br>cgtcaacggcaccgatctgcgcgatgcgaccggtgaaatccagttcggccagttcaagaaccagatcg<br>aatatcaaaatgccggttctgccctgaacaacgagatgaagaagcaagtgctttccggcgtcgatgtcga<br>gaatctcgtcggcaagaccgtgacggtggttggcgtgttcaaggtcgtcaatccgaagaactggctcgtca<br>cgcctgtggggctcgaagtgaa | This work |
| pR-eryG | promoter-RBS | agcaccgcttcgaaaatcgaaggcaaaaacggtgatgtcgttcttgccgcaaacaacgttgccaaatcct | This work |

| | | | |
|---|---|---|---|
| | | atggtcgcattcatgccctgaagggcgtgaatttcgagattcgccgtggtcaggtcacgacgcttttcggcg<br>aaaacggcgctggcaaatcaacgttgatgaaagtgctttcgggcgtcatccagccgacttcggggacgat<br>cattctcgatggcgaacctgtgacgttcaattcgtccaccgaagcgcgcgacctcggcatctcgatcatcc<br>atcaggaattgagcctcgcgcccaacatgaacgtgcgcgacaacatcttcatggggcgcgaaatccgc<br>accgccaccggcgttgattttgccgaggaagaacgcctgacgcgcgcgcttctgaaggaactggaaga<br>agacatcgacccgctgacgccggtcgaggaacttcgtctcggccagcagcaggttgttgaaatcgcccg<br>cgccctttcggtcaattcgcgcattctcatcatggatgagccgacttcggcgctcagcgcctcggaagtgga<br>agtcctgttcaaggtcattcgcgacctgacggcgcgcggtgtcgccatcgtctacatttcgcatcatctggaa<br>gaagcgttgcagatcaccaatcatgcggtggtgctgcgcgacggaaccatgacggcctatgcgccgcgt<br>gaggaaattgatctggaatggatcgtgcgcaacatggtcggcgagaacttcgatctcggctcgcctccaa<br>ccggatatgactggggcgatgtggcgctctctgttgagaacctgaccgttcccgacccgggtggcgcggg<br>cttctcgctggtcgaccgcatgtcgctcaatgtacgcgccggtgaaatcgtctgcatttatggtcttatgggcg<br>ccggtcgcacagaactgctggaaaccgttgccggtcgtctcaaggcaagcggcggacgtgttctcctcaa<br>agggcaggacgtttccggcctcaccatcgcacaacgtatcgagaagggtcttgtgctggtgccggaagat<br>cgccagcgcgacggtctcgtccagacgatgacggtcggcaagaacctgtcgctcgccagcatcgccga<br>aatgaccaagggtctgttcacatcgcgcaagcgtgaaaagcagattgtcgatcagtcgatcaagaatgtg<br>cacatcaagacggatggcggcgaagcagcaatcggttcgctttccggtggtaaccagcagaaggtcgtg<br>atcggcaagatgctggcgaccgaaccggaagtcatcctgcttgatgagccgagccgcgggatcgacat<br>cggggcgaaggcggaagtgttcaagcttctggctgaaaaggcgaagcagggtctggctgtcgtctacac<br>gacttcggaagtcggcgaatgcctcagcatcgctcatcgtatcatcgtcatgcaccgtggacgcatctctgc<br>cgaattcggatcggacgtgtccaaggagaagatcatggccgcctcgggcgaagccatggtcggtcacta<br>aaacataatcccgaaaagttgcagacttttcggacaagattatgcgttaaaaacaaaaggatagagcat<br>aagacccgacaaagtcgggttgttaa | |
| pR-eryH | promoter-RBS | acgatccatcgcgatctggattcgctggaacataccggcgtgctgcgcaaagtacgcggcggcgcgac<br>aatcgataccggcacacaatttgaaagcgatttccgcttccgcgaacggcaggacggcgaagcgaagc<br>agaaaatggcccgcgccgcgcttgagcttgttgaacccggcatgacggtgatgattaatgacggctccat<br>ggcggctgtccttggcggcatgcttctggaaaagcgcccgctgacagtcatcaccaacaatgccgtcatc<br>atcgatgaactgaagggcgagaacgggatcaatctgattgcgctcggcggaacctattcagccaagttc<br>aatgcgtttttcggcatcctgacggaaggggccctgtcgcatctgagcgccgacatcgccttcatttcctcgc<br>ctgcagtcaatggcaagctcgtctatcacatggatgagaatgttgttcgcaccaagcgcgcgatgacggc<br>atccgccgcccgcacctgccttctggtcaatcaccagcggttcggacgccctgccctgcatgtcatggctg<br>atctttccgatttcgacgcgatcataaccgatgcctcccccgacgcgaccgtactggccgatcttgaacag<br>gcgggcattgcactgaccatcgctgacgactgcgcataaacctagaggacgaaaatcgagc | This work |
| pR-eryI | promoter-RBS | acaaaattctggattggcaccagctggaagatgaacaagacgctggccgaggcccgcattttcgccgaa<br>gccttgaaagctgccgatgcaagccgctcgaccgacattcagcgtttcgtcatcccgccctttaccgccgtg<br>cgcgaagtgaaggaaatcctgttcggcacctccgtcaaggtcggcgcgcagaacatgcattgggccgat<br>cagggagcatggaccggcgagatttcgccgctgatgctcaaggactgcaatctcgatatcgtcgaactcg<br>gtcattccgagcgccgtgaacatttcggtgaaaccaacgaaacggtcggcctcaaggtcgaagctgcgg<br>tgcgccacggcctgatcccactcatctgcatcggtgagacgctggaagaccgcgaaagcggacgcgcc<br>gcggaagttcttgaggaagaagtgcgcggtgcactttccaagctttccggtgaccagaagcaggcggaa<br>atcctgtttgcctatgagccggtctgggccatcggcgaaaatggtatcccggcatcggcggaatatgccga<br>tgcgcgccaggcggaaatcatcgcggttgcagaaagcgtactcggtcgtcgtgtgccttgcctctatggcg<br>gctcggtcaatccgggcaattgcgaagagctgatcgcttgcccgcacattgacgggcttttcatcggtcgct<br>cggcatggaacgtcgaaggttatctcgacattctggccaagtgtgccgccaagactcgagctaattaagg<br>agcaat | This work |
| B0015 | terminator | ccaggcatcaaataaaacgaaaggctcagtcgaaagactgggcctttcgttttatctgttgtttgtcggtgaa | 1, BBa_B0015 |

| | | | |
|---|---|---|---|
| | | cgctctctactagagtcacactggctcaccttcgggtgggcctttctgcgtttata | |
| His operon terminator | terminator | tccggcaaaaaagggcaaggtgtcaccaccctgccctttttctttaaaaccgaaaaga | 1, pSB1C3 |
| Bacterial terminator | terminator | gaaatcatccttagcgaaagctaaggatttttttatctgaaat | 1, pSB1C3 |
| eryA | CDS | atgcgtgagaaaggtgacatcatcatcgggatcgacgcgggcacgtctgtgctcaaggcagtcgctttcg<br>accttagcggacgccagatcgaatctgctgccgttcgcaataaatacgataccggcgaacatggcgctgt<br>cacacagtcgctttcccagacgtggcaggactgcgcgcgcgccttgcgcgacctcggtgcaaagatgcc<br>cggccttgccgaacgcactgcggcaatcgccgttaccggtcagggtgacggaacctggcttgttggccgc<br>gacaaccagccggtcggcgatgcctggctgtggctcgatgcccgcgcggcaacgacggtgacgcgcct<br>cgccgctggccccgtgaaccgcgcgcgctttgaagcgaccggcactgggttgaacacctgccagcaag<br>gctcgcagatggcgcatatggacagcatcgcgcccgagctgctggacaatgccgaagtcgcgctccact<br>gcaaagactggctctatctcaatctcaccggcgttcgcgccaccgacccttcggaagcaagcttcaccttc<br>ggcaatttccgcaatcgccagtatgacgatgtcgtcatcgacgcgctcggtctgaggaaacggcgtggatt<br>gctgccggaaatcatcgacggcaccgaagtgcagcatccgctatcccccgaagcggcagctgcaacg<br>ggtttgcgggcagggacgccggtgggtctcgcctatgtcgatatggccatgacagcacttggtgcgggcgt<br>gcgcggcggcacagcaggcgccgggtgctcgaccattggttcgacaggcgtccatatgcgcgccaagc<br>cggttgccgatatccacctgaacaaggaaggcaccggttacgtcatcgcgctgcccattcccggcatcgt<br>cacccaggtgcagaccaatatgggggcaacgatcaacatcgactggatattgcaggttgctgccgatctc<br>atgtcgacgccagacaagcctgtttcgctcagtgacctcattccccgccttgatgactggtttaatgccagcc<br>gtccgggtgcaatcctctatcatccgtatatttcggaagcgggcgaacgcggtcccttcgtcaatgcctatgc<br>ccgggccggtttcgtcggtctttccagccgcgaccgtttcccggaaatggtgcgctcggttgtcgagggcct<br>cggaatggcgacgcgcgattgctacacggccatgggtgaaatgcccgcagaattgcgcatcaccggcg<br>gtgcagcgcgttcaaaagcattgcgcagcacccttgccgcagcggtgaatgcgcccgtgcgcgtttcctc<br>gcgtgaagaggcgggagccgcaggtgctgccatgatggctgcggtggccattggcgcttatccaagcat<br>ggatgcatgtattgcggaatgggtcgagccgctgcttggcccttgcgaggccccggatggtccgcgcgca<br>aagcactatgaagaactcttcgttgcctatcaggaagcccggctggcactgacgcccgtctgggacaagt<br>tggcttccgatagagcatttccagcaaaagcgtga | This work |
| eryB | CDS | atggctgaaccggaaacctgcgatctctttgtcatcggcggtggtatcaacggcgcgggcgtggcccgtg<br>acgccgccgggcgcggcctcaaggtggtgctggcggaaaaagacgatctggcgcagggaacttcgtc<br>acgctccggcaagctggtacatggtggtctgcgttatctcgaatattatgagtttcgccttgtgcgcgaagcg<br>ctgatcgagcgcgaagtgctgttgaacgccgctccccatatcatctggccaatgcgtttcgttctgccgcac<br>agcccgcaggaccgtcccgcctggctggtgcggcttggcctgtttctctatgaccacctcggcggccgcaa<br>gaagcttcccggcacacgcacgctcgacctgcggcgcgatccggaaggcacgccgatcctcgaccaat<br>ataccaagggcttcgaatattccgactgttgggtcgatgatgcccgtctcgttgcattgaatgcggttggcgct<br>gctgaaaaaggcgcgaccattctgacccgcacgcctgtcgtctccgctcgtcgcgaaaaaggcggctgg<br>atcgtggaaacgaaaaaccgcgacacgggcgaaacccgcacattccgcgcgcgctgcatcgtcaattg<br>cgccggaccatgggtcacggatgtcatccacaatgtcgccggctccaactcgtcgcgcaatgtgcgtctc<br>gtcaagggcagccacatcatcgttccgaaattctggtcgggcgcaaatgcctatctcgtccagaaccacg<br>acaagcgcgttatctttatcaatccctacgagggcgacaaggcgctgatcggcaccaccgacatcgccta<br>tgaaggccgtgccgaagatgttgcagcagatgagaaggaaatcgaatatctgctgactgcagtgaaccg<br>ctatttcaaggaaaagctcaggcgtgaagacgtgctgcattccttctccggtgtgcgtccgctgttcgacga<br>cggcaagggcaacccttccgccgtcacccgcgactatgtcttcgacctcgatgaaaccaacgacgcgcc<br>gctgctcaacgtctttggcggcaagatcaccaccttccgcgagctggccgagcgcggcatgcatcgcctc<br>aagcacattttcccgaagatgggcggcgactggactcatggcgcaccgcttccgggcggcgaaatcgcc<br>aatgccgattatgaaaccttcgccaactccttgcgcgacacctatccgtggatgccgcgcgcgctcgtcca | This work |

| | | | |
|---|---|---|---|
| | | acactacggtcgtctctacggcgcacgcacgaaagacgttgttgcaggcgcacagaacctcgaagggc<br>tcggccgtcatttcggcggcaatttccatgaggcggaagtgcgctatctggtggccagggaatgggcaat<br>gacggcggaagacattctctatcgccgcaccaagcactatctgcacctgaccgaagccgaacgcgccg<br>ctttcgtggaatggttcgacaacgctaaaatggtctcttga | |
| eryC | CDS | atggcccttacgctttctctgaacaccaatccacttgtgaaccgctttgccgaaccggatgacctgattgaaa<br>cggttgcgcgcgatctgcgcctgcgcgatctccagctcacacacgagttcatcaatccaagctggcaggc<br>tccaaccatccgccgcctgacgcgcgacatggacaaggccttgcagcgtaccggtgttcgcgtcacttcg<br>ggcatgaccggcccctatggccgcctcaaccatttcggccatccggacgcagaagtgcgccgttactatg<br>tcgactggttcaagaccttcgccgacattatcggcgatcttggcggcaagtcggtcggcacgcagtttgcg<br>atcttcacctataaggatttcgatgacgctgcgcgccgcgaagatctgatcaagatcgccatcgattgctgg<br>gcggaagtcgccgaacatgcgagccgtgccggtctcgactatgtgttctgggaaccgatgagcatcggg<br>cgtgaatttggcgagacgattgccgaatgcatgaagcttcaggatcgcctgaccgccgccgacatggcta<br>ttccgatgtggatgatggccgatatcgaccacggcgatgtgacttccgccaatccggacgatttcgatccct<br>acgcctgggcacgcgccgtgccgaaagtctcgccgatcattcacatcaagcagagcctgatggacaag<br>ggcggacatcgtcctttcacagccgcgttcaatgccaagggacgcatccagccggaaccgcttctgaaa<br>gcctttgccgaaggcggcgctgtggacaatgaaatctgcctcgaactgtcgttcaaggagcgcgagccg<br>aatgaccgtgaagtcattccgcagattgcggaaagtgtggctttctgggctccgcacattgacaccggcgc<br>taaggacttgaagatataa | This work |
| eryD | CDS | atggcagatgctgacgattcactggcgctgcgcgccgcatggcttcacttcgtggcgggcatgacccagtc<br>agccgttgcgaagcgccttggtctgccctccgtgaaagcgcatcgccttattgccaaagccgtcgcggac<br>ggtgcggtgaaggtcagtatcgacggcgacattaccgaatgtatcgacctcgaaaatcgtctcgccgatat<br>gtacggcctcgactattgcgaagtcgttcccgacatcggtgaggaaggtctgccgctcatggcgcttggcc<br>atgcgggtgcagattttctgcgccgcgagatcgagcatggcgaccatgaagtcatcggcatcggccatgg<br>ccgcacgctttcggctgcagtcaactatatgccgcgcgtcgtcgccaatgatctgcgcttcgtgtcgctgctc<br>ggcggcctgacgcgcaatttcgccgccaacccccatgacgttatgcaccgtatcgccgaaaagaccggt<br>atgcctgcttatgtgatgccggtgcccttcttcgccaacacggcggaagaccgcgaagtgctgctcgcaca<br>gcgcggcgtcaccaccgttttcgacatgggttgccaagccgaactgaagatcgtcggcatcggcactgtc<br>gatgcgcaggcgcagcttgtcacatccggcatgatagaactggccgaggtcgaagagatcgccagcct<br>cggcggtgttggcgaaatgctcggccatttctttgacgccaatggccagcgccttgaaaccgcgctgaccg<br>cccgcaccatcgctgcatcagtcgaaaacgcggatatgagccgaatcgtgggccttgccggtggaatttc<br>caaggcggaagccatccgcgctgtgctgaaaagcggtcgtctttacggtctcatcaccgatgaacgcac<br>ggcaaagacgctcgttgagcagccgaacggaaaataa | This work |
| eryE | CDS | atgagtgcacaaaccagtcaagtttcacaaaaacccgccaagctgagccgcgccgtcgtctggagcgt<br>gatcgctgcgatcgtggttgtcggtgcgattgccttcgacacgaaggtcgtgaagatcgggtctgattccga<br>tgtgcgccagcaggctttctcccccgatgcctacggtgcttccgaattccccaaggtgaaggcgagcgttg<br>agcagcgcgccgtggatgcagtggaagttggaacggctctcgccgccgacaaggccgccgccggaa<br>agaaatatggcgtcggcgacgtcaatccggttgtcccggtcaagtttacaggcaccgttgaggagcgcaa<br>atccaactacaatgtcgtgaaggtcgacggcctgccggaaggcgttgccatccgcgttcagaccggccct<br>gccgtcaacggcaccgatctgcgcgatgcgaccggtgaaatccagttcggccagttcaagaaccagatc<br>gaatatcaaaatgccggttctgccctgaacaacgagatgaagaagcaagtgctttccggcgtcgatgtcg<br>agaatctcgtcggcaagaccgtgacggtggttggcgtgttcaaggtcgtcaatccgaagaactggctcgtc<br>acgcctgtggggctcgaagtgaaatga | This work |
| eryF | CDS | atgagcaccgcttcgaaaatcgaaggcaaaaacggtgatgtcgttcttgccgcaaacaacgttgccaaat<br>cctatggtcgcattcatgccctgaagggcgtgaatttcgagattcgccgtggtcaggtcacgacgcttttcgg<br>cgaaaacggcgctggcaaatcaacgttgatgaaagtgctttcgggcgtcatccagccgacttcggggac<br>gatcattctcgatggcgaacctgtgacgttcaattcgtccaccgaagcgcgcgacctcggcatctcgatcat | This work |

| | | | |
|---|---|---|---|
| | | ccatcaggaattgagcctcgcgcccaacatgaacgtgcgcgacaacatcttcatggggcgcgaaatccgcaccgccaccggcgttgattttgccgaggaagaacgcctgacgcgcgcgcttctgaaggaactggaagaagacatcgacccgctgacgccggtcgaggaacttcgtctcggccagcagcaggttgttgaaatcgcccgcgccctttcggtcaattcgcgcattctcatcatggatgagccgacttcggcgctcagcgcctcggaagtggaagtcctgttcaaggtcattcgcgacctgacggcgcgcggtgtcgccatcgtctacatttcgcatcatctggaagaagcgttgcagatcaccaatcatgcggtggtgctgcgcgacggaaccatgacggcctatgcgccgcgtgaggaaattgatctggaatggatcgtgcgcaacatggtcggcgagaacttcgatctcggctcgcctccaaccggatatgactggggcgatgtggcgctctctgttgagaacctgaccgttcccgacccgggtggcgcgggcttctcgctggtcgaccgcatgtcgctcaatgtacgcgccggtgaaatcgtctgcatttatggtcttatgggcgccggtcgcacagaactgctggaaaccgttgccggtcgtctcaaggcaagcggcggacgtgttctcctcaaagggcaggacgtttccggcctcaccatcgcacaacgtatcgagaagggtcttgtgctggtgccggaagatcgccagcgcgacggtctcgtccagacgatgacggtcggcaagaacctgtcgctcgccagcatcgccgaaatgaccaagggtctgttcacatcgcgcaagcgtgaaaagcagattgtcgatcagtcgatcaagaatgtgcacatcaagacggatggcggcgaagcagcaatcggttcgctttccggtggtaaccagcagaaggtcgtgatcggcaagatgctggcgaccgaaccggaagtcatcctgcttgatgagccgagccgcgggatcgacatcggggcgaaggcggaagtgttcaagcttctggctgaaaaggcgaagcagggtctggctgtcgtctacacgacttcggaagtcggcgaatgcctcagcatcgctcatcgtatcatcgtcatgcaccgtggacgcatctctgccgaattcggatcggacgtgtccaaggagaagatcatggccgcctcgggcgaagccatggtcggtcactaa | |
| eryG | CDS | atgctctggttatggagcactgatagaatgtcagtcacgagcaccaacaaagaaacagcggctccgaacggcgcgaagcggaaacccggcatcgtgcatctgctgctcgaaggtcgtgcatttttttgcactgatcgcgatcattgcagttttctcgttcctttcgccttattatttctcggtcgataacttcctcatcatgtcgtcgcacgtggccattttcggcctgcttgccatcggcatgctgctcgtcatcctcaatggcggcatcgatctttccgttggctccacattggggctggccggtgtggtcgccgggttcctgatgcagggcgtgacgctgcaggaattcggcatcattctctatttcccggtctgggccgtggttctcatcacctgtgcgctcggcgcattcgtcggcgcggtcaatggtgtgctcatcgcctatctgcgcgttccggccttcgttgcgacgctcggcgttctctattgcgcgcgtggcgtggcgcttctgatgaccaatggcctgacctacaacaatctcggcggacgtccggagcttggcaatacaggctttgactggctcggtttcaacaagctgttcggcgtgccgatcggcgttctggttctggcggttctcgccatcatctgcgccatcgttctcaaccgcaccgcatttggtcgctggctctatgcgtcgggtggcaacgaacgtgcggctgaactgtccggcgttccggtcaagcgcgtcaaggtgtcggtctatgtcatttccggcatctgcgctgccattgcaggtctggttctttcttcgcagctgacgtcggccggtccgacggcaggcaccacctacgaactgacggcaatcgctgccgtggttatcggtggcgctgcgctcactgggggggcgcggtactatccagggcacacttctcggcgctttcgtgatcggtttcctttccgacggcctcgtgattatcggcgtgtcctcctactggcagaccgtctttaccggcgcggtaatcgtgctcgcggttctgctcaacagcattcaatactcccggcgatga | This work |
| eryH | CDS | atgacaaaattctggattggcaccagctggaagatgaacaagacgctggccgaggcccgcattttcgccgaagccttgaaagctgccgatgcaagccgctcgaccgacattcagcgtttcgtcatcccgccctttaccgccgtgcgcgaagtgaaggaaatcctgttcggcacctccgtcaaggtcggcgcgcagaacatgcattgggccgatcagggagcatggaccggcgagatttcgccgctgatgctcaaggactgcaatctcgatatcgtcgaactcggtcattccgagcgccgtgaacatttcggtgaaaccaacgaaacggtcggcctcaaggtcgaagctgcggtgcgccacggcctgatcccactcatctgcatcggtgagacgctggaagaccgcgaaagcggacgcgccgcggaagttcttgaggaagaagtgcgcggtgcactttccaagctttccggtgaccagaagcaggcggaaatcctgtttgcctatgagccggtctgggccatcggcgaaaatggtatcccggcatcggcggaatatgccgatgcgcgccaggcggaaatcatcgcggttgcagaaagcgtactcggtcgtcgtgtgccttgcctctatggcggctcggtcaatccgggcaattgcgaagagctgatcgcttgcccgcacattgacgggcttttcatcggtcgctcggcatggaacgtcgaaggttatctcgacattctggccaagtgtgccgccaagactcgagctaattaa | This work |

| | | | |
|---|---|---|---|
| eryI | CDS | atgatgaaagtagcagtagcaggcgacagcgccggcgaaggtctggccaaggttctcgccgatcacct<br>caaggatcgcttcgaggtttcggaaatctcgcgtacagacgctggcgcagatacattctacgccaacctct<br>ccgaccgcgtggcttccgccgttctcgacggcacctatgaccgcgccatcctcgtttgcggcaccggcatc<br>ggcgtatgcattgcagccaataaggttccgggcatccgcgccgcgctgacgcacgacacctattcggca<br>gagcgcgcagcgctttccaacaatgcccagatcatcaccatgggcgcccgcgtcatcggctcggaagta<br>gccaagaccattgccgacgccttccttgctgagacctttgacgaaaacggccgttcggctggcaatgtcaa<br>cgcgatcaacgaagttgacgcgaagtacaacaagggctga | This work |
| eryR | CDS | atgacgatccatcgcgatctggattcgctggaacataccggcgtgctgcgcaaagtacgcggcggcgcg<br>acaatcgataccggcacacaatttgaaagcgatttccgcttccgcgaacggcaggacggcgaagcgaa<br>gcagaaaatggcccgcgccgcgcttgagcttgttgaacccggcatgacggtgatgattaatgacggctcc<br>atggcggctgtccttggcggcatgcttctggaaaagcgcccgctgacagtcatcaccaacaatgccgtcat<br>catcgatgaactgaagggcgagaacgggatcaatctgattgcgctcggcggaacctattcagccaagttc<br>aatgcgtttttcggcatcctgacggaagggggccctgtcgcatctgagcgccgacatcgccttcatttcctcgc<br>ctgcagtcaatggcaagctcgtctatcacatggatgagaatgttgttcgcaccaagcgcgcgatgacggc<br>atccgccgcccgcacctgccttctggtcaatcaccagcggttcggacgccctgccctgcatgtcatggctg<br>atctttccgatttcgacgcgatcataaccgatgcctcccccgacgcgaccgtactggccgatcttgaacag<br>gcgggcattgcactgaccatcgctgacgactgcgcataa | This work |
| sfGFP | CDS | atgcgtaaaggcgaagagctgttcactggtgtcgtccctattctggtggaactggatggtgatgtcaacggt<br>cataagttttccgtgcgtggcgagggtgaaggtgacgcaactaatggtaaactgacgctgaagttcatctgt<br>actactggtaaactgccggtaccttggccgactctggtaacgacgctgacttatggtgttcagtgctttgctcg<br>ttatccggaccatatgaagcagcatgacttcttcaagtccgccatgccggaaggctatgtgcaggaacgc<br>acgatttcctttaaggatgacggcacgtacaaaacgcgtgcggaagtgaaatttgaaggcgataccctgg<br>taaaccgcattgagctgaaaggcattgactttaaagaagacggcaatatcctgggccataagctggaata<br>caattttaacagccacaatgtttacatcaccgccgataaacaaaaaaatggcattaaagcgaattttaaaa<br>ttcgccacaacgtggaggatggcagcgtgcagctggctgatcactaccagcaaaacactccaatcggtg<br>atggtcctgttctgctgccagacaatcactatctgagctaccaaagcgttctgtctaaagatccgaacgaga<br>aacgcgatcatatggttctgctggagttcgtaaccgcagcgggcatcacgcatggtatggatgaactgtac<br>taa | 2 |
| eryE-sfGFP-6xHis | CDS | atgagtgcacaaaccagtcaagtttcacaaaaacccgccaagctgagccgcgccgtcgtctggagcgt<br>gatcgctgcgatcgtggttgtcggtgcgattgccttcgacacgaaggtcgtgaagatcgggtctgattccga<br>tgtgcgccagcaggctttctcccccgatgcctacggtgcttccgaattccccaaggtgaaggcgagcgttg<br>agcagcgcgccgtggatgcagtggaagttggaacggctctcgccgccgacaaggccgccgccggaa<br>agaaatatggcgtcggcgacgtcaatccggttgtcccggtcaagtttacaggcaccgttgaggagcgcaa<br>atccaactacaatgtcgtgaaggtcgacggcctgccggaaggcgttgccatccgcgttcagaccggccct<br>gccgtcaacggcaccgatctgcgcgatgcgaccggtgaaatccagttcggccagttcaagaaccagatc<br>gaatatcaaaatgccggttctgccctgaacaacgagatgaagaagcaagtgctttccggcgtcgatgtcg<br>agaatctcgtcggcaagaccgtgacggtggttggcgtgttcaaggtcgtcaatccgaagaactggctcgtc<br>acgcctgtggggctcgaagtgaaaatgcgtaaaggcgaagagctgttcactggtgtcgtccctattctggt<br>ggaactggatggtgatgtcaacggtcataagttttccgtgcgtggcgagggtgaaggtgacgcaactaat<br>ggtaaactgacgctgaagttcatctgtactactggtaaactgccggtaccttggccgactctggtaacgacg<br>ctgacttatggtgttcagtgctttgctcgttatccggaccatatgaagcagcatgacttcttcaagtccgccatg<br>ccggaaggctatgtgcaggaacgcacgatttcctttaaggatgacggcacgtacaaaacgcgtgcgga<br>agtgaaatttgaaggcgataccctggtaaaccgcattgagctgaaaggcattgactttaaagaagacggc<br>aatatcctgggccataagctggaatacaattttaacagccacaatgtttacatcaccgccgataaacaaaa<br>aaatggcattaaagcgaattttaaaattcgccacaacgtggaggatggcagcgtgcagctggctgatcac<br>taccagcaaaacactccaatcggtgatggtcctgttctgctgccagacaatcactatctgagctaccaaag | This work |

| | | | |
|---|---|---|---|
| | | cgttctgtctaaagatccgaacgagaaacgcgatcatatggttctgctggagttcgtaaccgcagcgggca<br>tcacgcatggtatggatgaactgtaccatcatcatcatcatcactaataa | |
| eryG-sfGFP-6xHis | CDS | atgctctggttatggagcactgatagaatgtcagtcacgagcaccaacaaagaaacagcggctccgaac<br>ggcgcgaagcggaaacccggcatcgtgcatctgctgctcgaaggtcgtgcatttttttgcactgatcgcgat<br>cattgcagttttctcgttcctttcgccttattatttctcggtcgataacttcctcatcatgtcgtcgcacgtggccattt<br>tcggcctgcttgccatcggcatgctgctcgtcatcctcaatggcggcatcgatctttccgttggctccacattg<br>gggctggccggtgtggtcgccgggttcctgatgcagggcgtgacgctgcaggaattcggcatcattctctat<br>ttcccggtctgggccgtggttctcatcacctgtgcgctcggcgcattcgtcggcgcggtcaatggtgtgctcat<br>cgcctatctgcgcgttccggccttcgttgcgacgctcggcgttctctattgcgcgcgtggcgtggcgcttctga<br>tgaccaatggcctgacctacaacaatctcggcggacgtccggagcttggcaatacaggctttgactggct<br>cggtttcaacaagctgttcggcgtgccgatcggcgttctggttctggcggttctcgccatcatctgcgccatcg<br>ttctcaaccgcaccgcatttggtcgctggctctatgcgtcgggtggcaacgaacgtgcggctgaactgtccg<br>gcgttccggtcaagcgcgtcaaggtgtcggtctatgtcatttccggcatctgcgctgccattgcaggtctggtt<br>ctttcttcgcagctgacgtcggccggtccgacggcaggcaccacctacgaactgacggcaatcgctgcc<br>gtggttatcggtggcgctgcgctcactgggggcgcggtactatccagggcacacttctcggcgctttcgtg<br>atcggtttcctttccgacggcctcgtgattatcggcgtgtcctcctactggcagaccgtctttaccggcgcggt<br>aatcgtgctcgcggttctgctcaacagcattcaatactcccggcgaatgcgtaaaggcgaagagctgttca<br>ctggtgtcgtccctattctggtggaactggatggtgatgtcaacggtcataagttttccgtgcgtggcgagggt<br>gaaggtgacgcaactaatggtaaactgacgctgaagttcatctgtactactggtaaactgccggtaccttg<br>gccgactctggtaacgacgctgacttatggtgttcagtgctttgctcgttatccggaccatatgaagcagcat<br>gacttcttcaagtccgccatgccggaaggctatgtgcaggaacgcacgatttcctttaaggatgacggcac<br>gtacaaaacgcgtgcggaagtgaaatttgaaggcgataccctggtaaaccgcattgagctgaaaggcat<br>tgactttaaagaagacggcaatatcctgggccataagctggaatacaattttaacagccacaatgtttacat<br>caccgccgataaacaaaaaaatggcattaaagcgaattttaaaattcgccacaacgtggaggatggca<br>gcgtgcagctggctgatcactaccagcaaaacactccaatcggtgatggtcctgttctgctgccagacaat<br>cactatctgagctaccaaagcgttctgtctaaagatccgaacgagaaacgcgatcatatggttctgctgga<br>gttcgtaaccgcagcgggcatcacgcatggtatggatgaactgtaccatcatcatcatcatcactaataa | This work |
| eryD binding site (eryO) | protein binding site | gaaaaaaaatgcgccatctagaaaatttt | This work |

**Supplementary Table 3. Gene clusters.**

| Cluster name | Annotation | DNA sequence | Reference |
|---|---|---|---|
| original erythritol cluster | including all erythritol-associated promoters, RBSs, CDSs (eryA/B/C/D/E/F/G/H/I/R) and terminators | tcatcgccgggagtattgaatgctgttgagcagaaccgcgagcacgattaccgcgccggtaaagacggtctgccagtagg<br>aggacacgccgataatcacgaggccgtcggaaaggaaaccgatcacgaaagcgccgagaagtgtgccctggatagta<br>ccgcgccccccagtgagcgcagcgccaccgataaccacggcagcgattgccgtcagttcgtaggtggtgcctgccgtcg<br>gaccggccgacgtcagctgcgaagaaagaaccagacctgcaatggcagcgcagatgccggaaatgacatagaccga<br>caccttgacgcgcttgaccggaacgccggacagttcagccgcacgttcgttgccacccgacgcatagagccagcgacca<br>aatgcggtgcggttgagaacgatggcgcagatgatggcgagaaccgccagaaccagaacgccgatcggcacgccga<br>acagcttgttgaaaccgagccagtcaaagcctgtattgccaagctccggacgtccgccgagattgttgtaggtcaggccatt<br>ggtcatcagaagcgccacgccacgcgcgcaatagagaacgccgagcgtcgcaacgaaggccggaacgcgcagata<br>ggcgatgagcacaccattgaccgcgccgacgaatgcgccgagcgcacaggtgatgagaaccacggcccagaccggg<br>aaatagagaatgatgccgaattcctgcagcgtcacgccctgcatcaggaacccggcgaccacaccggccagccccaat<br>gtggagccaacggaaagatcgatgccgccattgaggatgacgagcagcatgccgatggcaagcaggccgaaaatggc<br>cacgtgcgacgacatgatgaggaagttatcgaccgagaaataataaggcgaaaggaacgagaaaactgcaatgatcg<br>cgatcagtgcaaaaaatgcacgaccttcgagcagcagatgcacgatgccgggtttccgcttcgcgccgttcggagccgct<br>gtttctttgttggtgctcgtgactgacattctatcagtgctccataaccagagcatttaacaacccgactttgtcgggtcttatgctct<br>atccttttgtttttaacgcataatcttgtccgaaaagtctgcaacttttcgggattatgttttagtgaccgaccatggcttcgcccga<br>ggcggccatgatcttctccttggacacgtccgatccgaattcggcagagatgcgtccacggtgcatgacgatgatacgatga<br>gcgatgctgaggcattcgccgacttccgaagtcgtgtagacgacagccagaccctgcttcgccttttcagccagaagcttga<br>acacttccgccttcgccccgatgtcgatcccgcggctcggctcatcaagcaggatgacttccggttcggtcgccagcatcttg<br>ccgatcacgaccttctgctggttaccaccggaaagcgaaccgattgctgcttcgccgccatccgtcttgatgtgcacattcttg<br>atcgactgatcgacaatctgcttttcacgcttgcgcgatgtgaacagacccttggtcatttcggcgatgctggcgagcgacag<br>gttcttgccgaccgtcatcgtctggacgagaccgtcgcgctggcgatcttccggcaccagcacaagacccttctcgatacgtt<br>gtgcgatggtgaggccggaaacgtcctgccctttgaggagaacacgtccgccgcttgccttgagacgaccggcaacggttt<br>ccagcagttctgtgcgaccggcgcccataagaccataaatgcagacgatttcaccggcgcgtacattgagcgacatgcgg<br>tcgaccagcgagaagcccgcgccacccgggtcgggaacggtcaggttctcaacagagagcgccacatcgccccagtc<br>atatccggttggaggcgagccgagatcgaagttctcgccgaccatgttgcgcacgatccattccagatcaatttcctcacgc<br>ggcgcataggccgtcatggttccgtcgcgcagcaccaccgcatgattggtgatctgcaacgcttcttccagatgatgcgaaa<br>tgtagacgatggcgacaccgcgcgccgtcaggtcgcgaatgaccttgaacaggacttccacttccgaggcgctgagcgc<br>cgaagtcggctcatccatgatgagaatgcgcgaattgaccgaaagggcgcgggcgatttcaacaacctgctgctggccg<br>agacgaagttcctcgaccggcgtcagcgggtcgatgtcttcttccagttccttcagaagcgcgcgcgtcaggcgttcttcctcg<br>gcaaaatcaacgccggtggcggtgcggatttcgcgccccatgaagatgttgtcgcgcacgttcatgttgggcgcgaggctc<br>aattcctgatggatgatcgagatgccgaggtcgcgcgcttcggtggacgaattgaacgtcacaggttcgccatcgagaatg<br>atcgtccccgaagtcggctggatgacgcccgaaagcactttcatcaacgttgatttgccagcgccgttttcgccgaaaagcg<br>tcgtgacctgaccacggcgaatctcgaaattcacgcccttcagggcatgaatgcgaccataggatttggcaacgttgtttgcg<br>gcaagaacgacatcaccgtttttgccttcgattttcgaagcggtgctcatttcacttcgagccccacaggcgtgacgagccag<br>ttcttcggattgacgaccttgaacacgccaaccaccgtcacggtcttgccgacgagattctcgacatcgacgccggaaagc<br>acttgcttcttcatctcgttgttcagggcagaaccggcattttgatattcgatctggttcttgaactggccgaactggatttcaccg<br>gtcgcatcgcgcagatcggtgccgttgacggcagggccggtctgaacgcggatggcaacgccttccggcaggccgtcga<br>ccttcacgacattgtagttggatttgcgctcctcaacggtgcctgtaaacttgaccgggacaaccggattgacgtcgccgacg<br>ccatatttctttccggcggcggccttgtcggcggcgagagccgttccaacttccactgcatccacggcgcgctgctcaacgct<br>cgccttcaccttggggaattcggaagcaccgtaggcatcgggggagaaagcctgctggcgcacatcggaatcagacccg<br>atcttcacgaccttcgtgtcgaaggcaatcgcaccgacaaccacgatcgcagcgatcacgctccagacgacggcgcggc<br>tcagcttggcgggtttttgtgaaacttgactggtttgtgcactcattatcaaacgccttcagcaattctgccgtattgtcgttatgga<br>aacggatgatgcggctgaattagaccgcccggagattaccctcgaccattgtcatcacgcttaccggataggcaaaagcct | This work |

| | | |
|---|---|---|
| | | ccgccaaagcggtgcaggacgcgacagacgagatcatgatagcctccacatgaattccagggccccgctcctccgtgca<br>tcctggaaacagatgcgccgcaatcaaaacctatgatagcggcgtgttattaactccaataatgatgttataaagctcgaaa<br>gctgtcaatcggatgttttcgaaacaccggtgaaatagcacaatttcgcgccgaatctcgatcacgatctagagaaatccac<br>tacataattcgttatcaatcgcactattcacagcaccgtcttttctcgcaaagccagccacaaagatactcacaaacttctatat<br>tttcttttattttcaatgaattaaacatatttaaaaccgaagaaacattaagctattgatggaaaacagtgggaaaaccgcactc<br>gaaggcgcggagatttcccctacattcactctttggtgaaaaaaaatgcgccatctagaaaattttacagacaacgtgatag<br>cgttatgttatctccagcatacgccatcgcccgatttcattataaacagggctgcgagcgacgtattttcggtcgcgggagga<br>atccaaatgtcaatgaaagcacatagccgccatgcgtgagaaaggtgacatcatcatcgggatcgacgcgggcacgtct<br>gtgctcaaggcagtcgctttcgaccttagcggacgccagatcgaatctgctgccgttcgcaataaatacgataccggcgaa<br>catggcgctgtcacacagtcgctttcccagacgtggcaggactgcgcgcgcgccttgcgcgacctcggtgcaaagatgcc<br>cggccttgccgaacgcactgcggcaatcgccgttaccggtcagggtgacggaacctggcttgttggccgcgacaaccagc<br>cggtcggcgatgcctggctgtggctcgatgcccgcgcggcaacgacggtgacgcgcctcgccgctggccccgtgaaccg<br>cgcgcgctttgaagcgaccggcactgggttgaacacctgccagcaaggctcgcagatggcgcatatggacagcatcgcg<br>cccgagctgctggacaatgccgaagtcgcgctccactgcaaagactggctctatctcaatctcaccggcgttcgcgccacc<br>gacccttcggaagcaagcttcaccttcggcaatttccgcaatcgccagtatgacgatgtcgtcatcgacgcgctcggtctga<br>ggaaacggcgtggattgctgccggaaatcatcgacggcaccgaagtgcagcatccgctatcccccgaagcggcagctg<br>caacgggtttgcgggcagggacgccggtgggtctcgcctatgtcgatatggccatgacagcacttggtgcgggcgtgcgc<br>ggcggcacagcaggcgccgggtgctcgaccattggttcgacaggcgtccatatgcgcgccaagccggttgccgatatcc<br>acctgaacaaggaaggcaccggttacgtcatcgcgctgcccattcccggcatcgtcacccaggtgcagaccaatatggg<br>ggcaacgatcaacatcgactggatattgcaggttgctgccgatctcatgtcgacgccagacaagcctgtttcgctcagtgac<br>ctcattccccgccttgatgactggtttaatgccagccgtccgggtgcaatcctctatcatccgtatatttcggaagcgggcgaa<br>cgcggtcccttcgtcaatgcctatgcccgggccggtttcgtcggtctttccagccgcgaccgtttcccggaaatggtgcgctcg<br>gttgtcgagggcctcggaatggcgacgcgcgattgctacacggccatgggtgaaatgcccgcagaattgcgcatcaccgg<br>cggtgcagcgcgttcaaaagcattgcgcagcacccttgccgcagcggtgaatgcgcccgtgcgcgtttcctcgcgtgaag<br>aggcgggagccgcaggtgctgccatgatggctgcggtggccattggcgcttatccaagcatggatgcatgtattgcggaat<br>gggtcgagccgctgcttggcccttgcgaggccccggatggtccgcgcgcaaagcactatgaagaactcttcgttgcctatc<br>aggaagcccggctggcactgacgcccgtctgggacaagttggcttccgatagagcatttccagcaaaagcgtgaagcgg<br>ttttgcgttcggaaatgcggtgaacaaaacgaataaggccggaccttccttccggcacatatagactatgggatgcaattgc<br>atccgaaggagcatgaaatggctgaaccggaaacctgcgatctctttgtcatcggcggtggtatcaacggcgcgggcgtg<br>gcccgtgacgccgccgggcgcggcctcaaggtggtgctggcggaaaaagacgatctggcgcagggaacttcgtcacgc<br>tccggcaagctggtacatggtggtctgcgttatctcgaatattatgagtttcgccttgtgcgcgaagcgctgatcgagcgcgaa<br>gtgctgttgaacgccgctccccatatcatctggccaatgcgtttcgttctgccgcacagcccgcaggaccgtcccgcctggct<br>ggtgcggcttggcctgtttctctatgaccacctcggcggccgcaagaagcttcccggcacacgcacgctcgacctgcggcg<br>cgatccggaaggcacgccgatcctcgaccaatataccaagggcttcgaatattccgactgttgggtcgatgatgcccgtctc<br>gttgcattgaatgcggttggcgctgctgaaaaaggcgcgaccattctgacccgcacgcctgtcgtctccgctcgtcgcgaaa<br>aaggcggctggatcgtggaaacgaaaaaccgcgacacgggcgaaacccgcacattccgcgcgcgctgcatcgtcaatt<br>gcgccggaccatgggtcacggatgtcatccacaatgtcgccggctccaactcgtcgcgcaatgtgcgtctcgtcaagggca<br>gccacatcatcgttccgaaattctggtcgggcgcaaatgcctatctcgtccagaaccacgacaagcgcgttatctttatcaat<br>ccctacgagggcgacaaggcgctgatcggcaccaccgacatcgcctatgaaggccgtgccgaagatgttgcagcagat<br>gagaaggaaatcgaatatctgctgactgcagtgaaccgctatttcaaggaaaagctcaggcgtgaagacgtgctgcattcc<br>ttctccggtgtgcgtccgctgttcgacgacggcaagggcaacccttccgccgtcacccgcgactatgtcttcgacctcgatga<br>aaccaacgacgcgccgctgctcaacgtctttggcggcaagatcaccaccttccgcgagctggccgagcgcggcatgcat<br>cgcctcaagcacattttcccgaagatgggcggcgactggactcatggcgcaccgcttccgggcggcgaaatcgccaatgc<br>cgattatgaaaccttcgccaactccttgcgcgacacctatccgtggatgccgcgcgcgctcgtccaacactacggtcgtctct<br>acggcgcacgcacgaaagacgttgttgcaggcgcacagaacctcgaagggctcggccgtcatttcggcggcaatttccat<br>gaggcggaagtgcgctatctggtggccagggaatgggcaatgacggcggaagacattctctatcgccgcaccaagcact |

| | | |
|---|---|---|
| | | atctgcacctgaccgaagccgaacgcgccgctttcgtggaatggttcgacaacgctaaaatggtctcttgaggacactatgg<br>cccttacgctttctctgaacaccaatccacttgtgaaccgctttgccgaaccggatgacctgattgaaacggttgcgcgcgatc<br>tgcgcctgcgcgatctccagctcacacacgagttcatcaatccaagctggcaggctccaaccatccgccgcctgacgcgc<br>gacatggacaaggccttgcagcgtaccggtgttcgcgtcacttcgggcatgaccggcccctatggccgcctcaaccatttcg<br>gccatccggacgcagaagtgcgccgttactatgtcgactggttcaagaccttcgccgacattatcggcgatcttggcggcaa<br>gtcggtcggcacgcagtttgcgatcttcacctataaggatttcgatgacgctgcgcgccgcgaagatctgatcaagatcgcc<br>atcgattgctgggcggaagtcgccgaacatgcgagccgtgccggtctcgactatgtgttctgggaaccgatgagcatcggg<br>cgtgaatttggcgagacgattgccgaatgcatgaagcttcaggatcgcctgaccgccgccgacatggctattccgatgtgg<br>atgatggccgatatcgaccacggcgatgtgacttccgccaatccggacgatttcgatccctacgcctgggcacgcgccgtg<br>ccgaaagtctcgccgatcattcacatcaagcagagcctgatggacaagggcggacatcgtcctttcacagccgcgttcaat<br>gccaagggacgcatccagccggaaccgcttctgaaagcctttgccgaaggcggcgctgtggacaatgaaatctgcctcg<br>aactgtcgttcaaggagcgcgagccgaatgaccgtgaagtcattccgcagattgcggaaagtgtggctttctgggctccgc<br>acattgacaccggcgctaaggacttgaagatataaacgatttcaggaaccggaccgctttcaatggcagatgctgacgatt<br>cactggcgctgcgcgccgcatggcttcacttcgtggcgggcatgacccagtcagccgttgcgaagcgccttggtctgccctc<br>cgtgaaagcgcatcgccttattgccaaagccgtcgcggacggtgcggtgaaggtcagtatcgacggcgacattaccgaat<br>gtatcgacctcgaaaatcgtctcgccgatatgtacggcctcgactattgcgaagtcgttcccgacatcggtgaggaaggtctg<br>ccgctcatggcgcttggccatgcgggtgcagattttctgcgccgcgagatcgagcatggcgaccatgaagtcatcggcatc<br>ggccatggccgcacgctttcggctgcagtcaactatatgccgcgcgtcgtcgccaatgatctgcgcttcgtgtcgctgctcgg<br>cggcctgacgcgcaatttcgccgccaacccccatgacgttatgcaccgtatcgccgaaaagaccggtatgcctgcttatgtg<br>atgccggtgcccttcttcgccaacacggcggaagaccgcgaagtgctgctcgcacagcgcggcgtcaccaccgttttcga<br>catgggttgccaagccgaactgaagatcgtcggcatcggcactgtcgatgcgcaggcgcagcttgtcacatccggcatgat<br>agaactggccgaggtcgaagagatcgccagcctcggcggtgttggcgaaatgctcggccatttctttgacgccaatggcca<br>gcgccttgaaaccgcgctgaccgcccgcaccatcgctgcatcagtcgaaaacgcggatatgagccgaatcgtgggccttg<br>ccggtggaatttccaaggcggaagccatccgcgctgtgctgaaaagcggtcgtctttacggtctcatcaccgatgaacgca<br>cggcaaagacgctcgttgagcagccgaacggaaaataacataataacatttttgcaatggattattttgtgattttgtgatctaa<br>cttcgcgcagaaaatgttccgccaataagatcggacgagagcgtgaagcgcgacgacagaagacaggcgattataaac<br>cttctgatcgaaaatcaggccgtcgacctcgacgatctggcggatcggtttgccgtatcaaaaatgacgatccatcgcgatct<br>ggattcgctggaacataccggcgtgctgcgcaaagtacgcggcggcgcgacaatcgataccggcacacaatttgaaagc<br>gatttccgcttccgcgaacggcaggacggcgaagcgaagcagaaaatggcccgcgccgcgcttgagcttgttgaacccg<br>gcatgacggtgatgattaatgacggctccatggcggctgtccttggcggcatgcttctggaaaagcgcccgctgacagtcat<br>caccaacaatgccgtcatcatcgatgaactgaagggcgagaacgggatcaatctgattgcgctcggcggaacctattcag<br>ccaagttcaatgcgtttttcggcatcctgacggaaggggccctgtcgcatctgagcgccgacatcgccttcatttcctcgcctg<br>cagtcaatggcaagctcgtctatcacatggatgagaatgttgttcgcaccaagcgcgcgatgacggcatccgccgcccgc<br>acctgccttctggtcaatcaccagcggttcggacgccctgccctgcatgtcatggctgatctttccgatttcgacgcgatcataa<br>ccgatgcctcccccgacgcgaccgtactggccgatcttgaacaggcgggcattgcactgaccatcgctgacgactgcgca<br>taaacctagaggacgaaaatcgagcatgacaaaattctggattggcaccagctggaagatgaacaagacgctggccga<br>ggcccgcattttcgccgaagccttgaaagctgccgatgcaagccgctcgaccgacattcagcgtttcgtcatcccgcccttta<br>ccgccgtgcgcgaagtgaaggaaatcctgttcggcacctccgtcaaggtcggcgcgcagaacatgcattgggccgatca<br>gggagcatggaccggcgagatttcgccgctgatgctcaaggactgcaatctcgatatcgtcgaactcggtcattccgagcg<br>ccgtgaacatttcggtgaaaccaacgaaacggtcggcctcaaggtcgaagctgcggtgcgccacggcctgatcccactc<br>atctgcatcggtgagacgctggaagaccgcgaaagcggacgcgccgcggaagttcttgaggaagaagtgcgcggtgca<br>ctttccaagctttccggtgaccagaagcaggcggaaatcctgtttgcctatgagccggtctgggccatcggcgaaaatggta<br>tcccggcatcggcggaatatgccgatgcgcgccaggcggaaatcatcgcggttgcagaaagcgtactcggtcgtcgtgtg<br>ccttgcctctatggcggctcggtcaatccgggcaattgcgaagagctgatcgcttgcccgcacattgacgggcttttcatcggt<br>cgctcggcatggaacgtcgaaggttatctcgacattctggccaagtgtgccgccaagactcgagctaattaaggagcaata<br>tgatgaaagtagcagtagcaggcgacagcgccggcgaaggtctggccaaggttctcgccgatcacctcaaggatcgcttc |

| | | | |
|---|---|---|---|
| | | gaggtttcggaaatctcgcgtacagacgctggcgcagatacattctacgccaacctctccgaccgcgtggcttccgccgttctcgacggcacctatgaccgcgccatcctcgtttgcggcaccggcatcggcgtatgcattgcagccaataaggttccgggcatccgcgccgcgctgacgcacgacacctattcggcagagcgcgcagcgctttccaacaatgcccagatcatcaccatgggcgcccgcgtcatcggctcggaagtagccaagaccattgccgacgccttccttgctgagacctttgacgaaaacggccgttcggctggcaatgtcaacgcgatcaacgaagttgacgcgaagtacaacaagggctga | |
| programmed erythritol catabolism cluster | including all erythritol catabolism-associated promoters, RBSs, CDSs (eryA/B/C/H/I) and terminators | tatcaaacgccttcagcaattctgccgtattgtcgttatggaaacggatgatgcggctgaattagaccgcccggagattaccctcgaccattgtcatcacgcttaccggataggcaaaagcctccgccaaagcggtgcaggacgcgacagacgagatcatgatagcctccacatgaattccagggccccgctcctccgtgcatcctggaaacagatgcgccgcaatcaaaacctatgatagcggcgtgttattaactccaataatgatgttataaagctcgaaagctgtcaatcggatgttttcgaaacaccggtgaaatagcacaatttcgcgccgaatctcgatcacgatctagagaaatccactacataattcgttatcaatcgcactattcacagcaccgtcttttctcgcaaagccagccacaaagatactcacaaacttctatattttcttttattttcaatgaattaaacatatttaaaaccgaagaaacattaagctattgatggaaaacagtgggaaaaccgcactcgaaggcgcggagatttcccctacattcactctttggtgaaaaaaaatgcgccatctagaaaattttacagacaacgtgatagcgttatgttatctccagcatacgccatcgcccgatttcattataaacagggctgcgagcgacgtattttcggtcgcgggaggaatccaaatgtcaatgaaagcacatagccgccatgcgtgagaaaggtgacatcatcatcgggatcgacgcgggcacgtctgtgctcaaggcagtcgctttcgaccttagcggacgccagatcgaatctgctgccgttcgcaataaatacgataccggcgaacatggcgctgtcacacagtcgctttcccagacgtggcaggactgcgcgcgcgccttgcgcgacctcggtgcaaagatgcccggccttgccgaacgcactgcggcaatcgccgttaccggtcagggtgacggaacctggcttgttggccgcgacaaccagccggtcggcgatgcctggctgtggctcgatgcccgcgcggcaacgacggtgacgcgcctcgccgctggccccgtgaaccgcgcgcgctttgaagcgaccggcactgggttgaacacctgccagcaaggctcgcagatggcgcatatggacagcatcgcgcccgagctgctggacaatgccgaagtcgcgctccactgcaaagactggctctatctcaatctcaccggcgttcgcgccaccgacccttcggaagcaagcttcaccttcggcaatttccgcaatcgccagtatgacgatgtcgtcatcgacgcgctcggtctgaggaaacggcgtggattgctgccggaaatcatcgacggcaccgaagtgcagcatccgctatcccccgaagcggcagctgcaacgggtttgcgggcagggacgccggtgggtctcgcctatgtcgatatggccatgacagcacttggtgcgggcgtgcgcggcggcacagcaggcgccgggtgctcgaccattggttcgacaggcgtccatatgcgcgccaagccggttgccgatatccacctgaacaaggaaggcaccggttacgtcatcgcgctgcccattcccggcatcgtcacccaggtgcagaccaatatgggggcaacgatcaacatcgactggatattgcaggttgctgccgatctcatgtcgacgccagacaagcctgtttcgctcagtgacctcattccccgccttgatgactggtttaatgccagccgtccgggtgcaatcctctatcatccgtatatttcggaagcgggcgaacgcggtcccttcgtcaatgcctatgcccgggccggtttcgtcggtctttccagccgcgaccgtttcccggaaatggtgcgctcggttgtcgagggcctcggaatggcgacgcgcgattgctacacggccatgggtgaaatgcccgcagaattgcgcatcaccggcggtgcagcgcgttcaaaagcattgcgcagcacccttgccgcagcggtgaatgcgcccgtgcgcgtttcctcgcgtgaagaggcgggagccgcaggtgctgccatgatggctgcggtggccattggcgcttatccaagcatggatgcatgtattgcggaatgggtcgagccgctgcttggcccttgcgaggccccggatggtccgcgcgcaaagcactatgaagaactcttcgttgcctatcaggaagcccggctggcactgacgcccgtctgggacaagttggcttccgatagagcatttccagcaaaagcgtgaagcggttttgcgttcggaaatgcggtgaacaaaacgaataaggccggaccttccttccggcacatatagactatgggatgcaattgcatccgaaggagcatgaaatggctgaaccggaaacctgcgatctctttgtcatcggcggtggtatcaacggcgcgggcgtggcccgtgacgccgccgggcgcggcctcaaggtggtgctggcggaaaaagacgatctggcgcagggaacttcgtcacgctccggcaagctggtacatggtggtctgcgttatctcgaatattatgagtttcgccttgtgcgcgaagcgctgatcgagcgcgaagtgctgttgaacgccgctccccatatcatctggccaatgcgtttcgttctgccgcacagcccgcaggaccgtcccgcctggctggtgcggcttggcctgtttctctatgaccacctcggcggccgcaagaagcttcccggcacacgcacgctcgacctgcggcgcgatccggaaggcacgccgatcctcgaccaatataccaagggcttcgaatattccgactgttgggtcgatgatgcccgtctcgttgcattgaatgcggttggcgctgctgaaaaaggcgcgaccattctgacccgcacgcctgtcgtctccgctcgtcgcgaaaaaggcggctggatcgtggaaacgaaaaaccgcgacacgggcgaaacccgcacattccgcgcgcgctgcatcgtcaattgcgccggaccatgggtcacggatgtcatccacaatgtcgccggctccaactcgtcgcgcaatgtgcgtctcgtcaagggcagccacatcatcgttccgaaattctggtcgggcgcaaatgcctatctcgtccagaaccacgacaagcgcgttatctttatcaatccctacgagggcgacaaggcgctgatcggcaccaccgacatcgcctatgaaggccgtgccgaagatgttgcagcagatgagaaggaaatcgaatatctgctgactgcagtgaaccgctatttcaa | This work |

| | | | |
|---|---|---|---|
| | | ggaaaagctcaggcgtgaagacgtgctgcattccttctccggtgtgcgtccgctgttcgacgacggcaagggcaacccttc<br>cgccgtcacccgcgactatgtcttcgacctcgatgaaaccaacgacgcgccgctgctcaacgtctttggcggcaagatcac<br>caccttccgcgagctggccgagcgcggcatgcatcgcctcaagcacattttcccgaagatgggcggcgactggactcatg<br>gcgcaccgcttccgggcggcgaaatcgccaatgccgattatgaaaccttcgccaactccttgcgcgacacctatccgtgga<br>tgccgcgcgcgctcgtccaacactacggtcgtctctacggcgcacgcacgaaagacgttgttgcaggcgcacagaacctc<br>gaagggctcggccgtcatttcggcggcaatttccatgaggcggaagtgcgctatctggtggccagggaatgggcaatgac<br>ggcggaagacattctctatcgccgcaccaagcactatctgcacctgaccgaagccgaacgcgccgctttcgtggaatggtt<br>cgacaacgctaaaatggtctcttgaggacactatggcccttacgctttctctgaacaccaatccacttgtgaaccgctttgccg<br>aaccggatgacctgattgaaacggttgcgcgcgatctgcgcctgcgcgatctccagctcacacacgagttcatcaatccaa<br>gctggcaggctccaaccatccgccgcctgacgcgcgacatggacaaggccttgcagcgtaccggtgttcgcgtcacttcg<br>ggcatgaccggcccctatggccgcctcaaccatttcggccatccggacgcagaagtgcgccgttactatgtcgactggttca<br>agaccttcgccgacattatcggcgatcttggcggcaagtcggtcggcacgcagtttgcgatcttcacctataaggatttcgatg<br>acgctgcgcgccgcgaagatctgatcaagatcgccatcgattgctgggcggaagtcgccgaacatgcgagccgtgccgg<br>tctcgactatgtgttctgggaaccgatgagcatcgggcgtgaatttggcgagacgattgccgaatgcatgaagcttcaggatc<br>gcctgaccgccgccgacatggctattccgatgtggatgatggccgatatcgaccacggcgatgtgacttccgccaatccgg<br>acgatttcgatccctacgcctgggcacgcgccgtgccgaaagtctcgccgatcattcacatcaagcagagcctgatggaca<br>agggcggacatcgtcctttcacagccgcgttcaatgccaagggacgcatccagccggaaccgcttctgaaagcctttgccg<br>aaggcggcgctgtggacaatgaaatctgcctcgaactgtcgttcaaggagcgcgagccgaatgaccgtgaagtcattccg<br>cagattgcggaaagtgtggctttctgggctccgcacattgacaccggcgctaaggacttgaagatataaacgatccatcgc<br>gatctggattcgctggaacataccggcgtgctgcgcaaagtacgcggcggcgcgacaatcgataccggcacacaatttga<br>aagcgatttccgcttccgcgaacggcaggacggcgaagcgaagcagaaaatggcccgcgccgcgcttgagcttgttgaa<br>cccggcatgacggtgatgattaatgacggctccatggcggctgtccttggcggcatgcttctggaaaagcgcccgctgaca<br>gtcatcaccaacaatgccgtcatcatcgatgaactgaagggcgagaacgggatcaatctgattgcgctcggcggaaccta<br>ttcagccaagttcaatgcgtttttcggcatcctgacggaaggggccctgtcgcatctgagcgccgacatcgccttcatttcctcg<br>cctgcagtcaatggcaagctcgtctatcacatggatgagaatgttgttcgcaccaagcgcgcgatgacggcatccgccgcc<br>cgcacctgccttctggtcaatcaccagcggttcggacgccctgccctgcatgtcatggctgatctttccgatttcgacgcgatc<br>ataaccgatgcctcccccgacgcgaccgtactggccgatcttgaacaggcgggcattgcactgaccatcgctgacgactg<br>cgcataaacctagaggacgaaaatcgagcatgacaaaattctggattggcaccagctggaagatgaacaagacgctgg<br>ccgaggcccgcattttcgccgaagccttgaaagctgccgatgcaagccgctcgaccgacattcagcgtttcgtcatcccgc<br>cctttaccgccgtgcgcgaagtgaaggaaatcctgttcggcacctccgtcaaggtcggcgcgcagaacatgcattgggcc<br>gatcagggagcatggaccggcgagatttcgccgctgatgctcaaggactgcaatctcgatatcgtcgaactcggtcattccg<br>agcgccgtgaacatttcggtgaaaccaacgaaacggtcggcctcaaggtcgaagctgcggtgcgccacggcctgatccc<br>actcatctgcatcggtgagacgctggaagaccgcgaaagcggacgcgccgcggaagttcttgaggaagaagtgcgcgg<br>tgcactttccaagctttccggtgaccagaagcaggcggaaatcctgtttgcctatgagccggtctgggccatcggcgaaaat<br>ggtatcccggcatcggcggaatatgccgatgcgcgccaggcggaaatcatcgcggttgcagaaagcgtactcggtcgtcg<br>tgtgccttgcctctatggcggctcggtcaatccgggcaattgcgaagagctgatcgcttgcccgcacattgacgggcttttcat<br>cggtcgctcggcatggaacgtcgaaggttatctcgacattctggccaagtgtgccgccaagactcgagctaattaaggagc<br>aatatgatgaaagtagcagtagcaggcgacagcgccggcgaaggtctggccaaggttctcgccgatcacctcaaggatc<br>gcttcgaggtttcggaaatctcgcgtacagacgctggcgcagatacattctacgccaacctctccgaccgcgtggcttccgc<br>cgttctcgacggcacctatgaccgcgccatcctcgtttgcggcaccggcatcggcgtatgcattgcagccaataaggttccg<br>ggcatccgcgccgcgctgacgcacgacacctattcggcagagcgcgcagcgctttccaacaatgcccagatcatcacca<br>tgggcgcccgcgtcatcggctcggaagtagccaagaccattgccgacgccttccttgctgagacctttgacgaaaacggcc<br>gttcggctggcaatgtcaacgcgatcaacgaagttgacgcgaagtacaacaagggctga | |
| programmed erythritol transport | including all erythritol transport- | ggcggctatgtgctttcattgacatttggattcctcccgcgaccgaaaatacgtcgctcgcagccctgtttataatgaaatcggg<br>cgatggcgtatgctggagataacataacgctatcacgttgtctgtaaaattttctagatggcgcatttttttcaccaaagagtga<br>atgtaggggaaatctccgcgccttcgagtgcggttttcccactgttttccatcaatagcttaatgtttcttcggttttaaatatgtttaa | This work |

| | | |
|---|---|---|
| cluster | associated promoters, RBSs, CDSs (eryE/F/G) and terminators | ttcattgaaaataaaagaaaatatagaagtttgtgagtatctttgtggctggctttgcgagaaaagacggtgctgtgaatagtg<br>cgattgataacgaattatgtagtggatttctctagatcgtgatcgagattcggcgcgaaattgtgctatttcaccggtgtttcgaa<br>aacatccgattgacagctttcgagctttataacatcattattggagttaataacacgccgctatcataggttttgattgcggcgca<br>tctgtttccaggatgcacggaggagcggggccctggaattcatgtggaggctatcatgatctcgtctgtcgcgtcctgcaccg<br>ctttggcggaggcttttgcctatccggtaagcgtgatgacaatggtcgagggtaatctccgggcggtctaattcagccgcatc<br>atccgtttccataacgacaatacggcagaattgctgaaggcgtttgataatgagtgcacaaaccagtcaagtttcacaaaaa<br>cccgccaagctgagccgcgccgtcgtctggagcgtgatcgctgcgatcgtggttgtcggtgcgattgccttcgacacgaag<br>gtcgtgaagatcgggtctgattccgatgtgcgccagcaggctttctcccccgatgcctacggtgcttccgaattccccaaggt<br>gaaggcgagcgttgagcagcgcgccgtggatgcagtggaagttggaacggctctcgccgccgacaaggccgccgccg<br>gaaagaaatatggcgtcggcgacgtcaatccggttgtcccggtcaagtttacaggcaccgttgaggagcgcaaatccaac<br>tacaatgtcgtgaaggtcgacggcctgccggaaggcgttgccatccgcgttcagaccggccctgccgtcaacggcaccga<br>tctgcgcgatgcgaccggtgaaatccagttcggccagttcaagaaccagatcgaatatcaaaatgccggttctgccctgaa<br>caacgagatgaagaagcaagtgctttccggcgtcgatgtcgagaatctcgtcggcaagaccgtgacggtggttggcgtgtt<br>caaggtcgtcaatccgaagaactggctcgtcacgcctgtggggctcgaagtgaaatgagcaccgcttcgaaaatcgaag<br>gcaaaaacggtgatgtcgttcttgccgcaaacaacgttgccaaatcctatggtcgcattcatgccctgaagggcgtgaatttc<br>gagattcgccgtggtcaggtcacgacgcttttcggcgaaaacggcgctggcaaatcaacgttgatgaaagtgctttcgggc<br>gtcatccagccgacttcggggacgatcattctcgatggcgaacctgtgacgttcaattcgtccaccgaagcgcgcgacctcg<br>gcatctcgatcatccatcaggaattgagcctcgcgcccaacatgaacgtgcgcgacaacatcttcatggggcgcgaaatc<br>cgcaccgccaccggcgttgattttgccgaggaagaacgcctgacgcgcgcgcttctgaaggaactggaagaagacatcg<br>acccgctgacgccggtcgaggaacttcgtctcggccagcagcaggttgttgaaatcgcccgcgccctttcggtcaattcgcg<br>cattctcatcatggatgagccgacttcggcgctcagcgcctcggaagtggaagtcctgttcaaggtcattcgcgacctgacg<br>gcgcgcggtgtcgccatcgtctacatttcgcatcatctggaagaagcgttgcagatcaccaatcatgcggtggtgctgcgcg<br>acggaaccatgacggcctatgcgccgcgtgaggaaattgatctggaatggatcgtgcgcaacatggtcggcgagaacttc<br>gatctcggctcgcctccaaccggatatgactggggcgatgtggcgctctctgttgagaacctgaccgttcccgacccgggtg<br>gcgcgggcttctcgctggtcgaccgcatgtcgctcaatgtacgcgccggtgaaatcgtctgcatttatggtcttatgggcgccg<br>gtcgcacagaactgctggaaaccgttgccggtcgtctcaaggcaagcggcggacgtgttctcctcaaagggcaggacgttt<br>ccggcctcaccatcgcacaacgtatcgagaagggtcttgtgctggtgccggaagatcgccagcgcgacggtctcgtccag<br>acgatgacggtcggcaagaacctgtcgctcgccagcatcgccgaaatgaccaagggtctgttcacatcgcgcaagcgtg<br>aaaagcagattgtcgatcagtcgatcaagaatgtgcacatcaagacggatggcggcgaagcagcaatcggttcgctttcc<br>ggtggtaaccagcagaaggtcgtgatcggcaagatgctggcgaccgaaccggaagtcatcctgcttgatgagccgagcc<br>gcgggatcgacatcggggcgaaggcggaagtgttcaagcttctggctgaaaaggcgaagcagggtctggctgtcgtctac<br>acgacttcggaagtcggcgaatgcctcagcatcgctcatcgtatcatcgtcatgcaccgtggacgcatctctgccgaattcg<br>gatcggacgtgtccaaggagaagatcatggccgcctcgggcgaagccatggtcggtcactaaaacataatcccgaaaa<br>gttgcagacttttcggacaagattatgcgttaaaaacaaaaggatagagcataagacccgacaaagtcgggttgttaaatg<br>ctctggttatggagcactgatagaatgtcagtcacgagcaccaacaaagaaacagcggctccgaacggcgcgaagcgg<br>aaacccggcatcgtgcatctgctgctcgaaggtcgtgcatttttttgcactgatcgcgatcattgcagttttctcgttcctttcgcctt<br>attatttctcggtcgataacttcctcatcatgtcgtcgcacgtggccattttcggcctgcttgccatcggcatgctgctcgtcatcct<br>caatggcggcatcgatctttccgttggctccacattggggctggccggtgtggtcgccgggttcctgatgcagggcgtgacgc<br>tgcaggaattcggcatcattctctatttcccggtctgggccgtggttctcatcacctgtgcgctcggcgcattcgtcggcgcggtc<br>aatggtgtgctcatcgcctatctgcgcgttccggccttcgttgcgacgctcggcgttctctattgcgcgcgtggcgtggcgcttct<br>gatgaccaatggcctgacctacaacaatctcggcggacgtccggagcttggcaatacaggctttgactggctcggtttcaac<br>aagctgttcggcgtgccgatcggcgttctggttctggcggttctcgccatcatctgcgccatcgttctcaaccgcaccgcatttg<br>gtcgctggctctatgcgtcgggtggcaacgaacgtgcggctgaactgtccggcgttccggtcaagcgcgtcaaggtgtcggt<br>ctatgtcatttccggcatctgcgctgccattgcaggtctggttctttcttcgcagctgacgtcggccggtccgacggcaggcacc<br>acctacgaactgacggcaatcgctgccgtggttatcggtggcgctgcgctcactgggggcgcggtactatccagggcac<br>acttctcggcgctttcgtgatcggtttcctttccgacggcctcgtgattatcggcgtgtcctcctactggcagaccgtctttaccgg |

| | | |
|---|---|---|
| | | cgcggtaatcgtgctcgcggttctgctcaacagcattcaatactcccggcgatga |

### Supplementary Table 4. Vectors.

| Vector name | Antibiotic resistance gene | Origin of replication | Reference |
|---|---|---|---|
| pSB1C3 | CmR | ColE1 | 1, pSB1C3 |
| pET22b | AmpR | ColE1+rop | This work |
| pFB1 | AmpR | p15A | 2 |
| pKD4 | AmpR | R6K | 3 |

Vector pSB1C3 was obtained from "iGEM *Registry of Standard Biological Parts*" registry distribution kits (Spring 2016 Distribution and Spring 2017 Distribution).

### Supplementary Table 5. Plasmids.

| Plasmid name | Biobricks | Vector | Antibiotic resistance gene | Origin of replication |
|---|---|---|---|---|
| pFB147 | pEryF-732 - eryA - eryB - eryC - eryH - eryI | pSB1C3 | CmR | ColE1 |
| pFB148 | J23100 - eryA - eryB - eryC - eryH - eryI | pSB1C3 | CmR | ColE1 |
| pFB149 | J23105 - eryA - eryB - eryC - eryH - eryI | pSB1C3 | CmR | ColE1 |
| pFB150 | J23106 - eryA - eryB - eryC - eryH - eryI | pSB1C3 | CmR | ColE1 |
| pFB151 | J23109 - eryA - eryB - eryC - eryH - eryI | pSB1C3 | CmR | ColE1 |
| pFB152 | J23114 - eryA - eryB - eryC - eryH - eryI | pSB1C3 | CmR | ColE1 |
| pFB153 | pEryF-732 - eryA - eryB - eryC - eryH | pSB1C3 | CmR | ColE1 |
| pFB154 | pEryF-732 - eryA - eryB - eryC | pSB1C3 | CmR | ColE1 |
| pFB155 | pEryF-732 - eryA - eryB | pSB1C3 | CmR | ColE1 |
| pFB156 | pEryF-732 - eryA | pSB1C3 | CmR | ColE1 |
| pFB157 | pEryR-732 - eryE - eryF - eryG | pFB1 | AmpR | p15A |
| pFB158 | araC - paraBAD - B0034 - (6xHis) - eryA - B0015 | pSB1C3 | CmR | ColE1 |
| pFB159 | araC - paraBAD - B0034 - (6xHis) - eryB - B0015 | pSB1C3 | CmR | ColE1 |
| pFB160 | araC - paraBAD - B0034 - (6xHis) - eryC - B0015 | pSB1C3 | CmR | ColE1 |
| pFB161 | pT7 - lacO - RBS - (6xHis) - eryD - T7 terminator | pET22b | AmpR | ColE1 + rop |
| pFB162 | araC - paraBAD - B0034 - eryE - sfGFP - (6xHis) - B0015 | pSB1C3 | CmR | ColE1 |
| pFB163 | pT7 - lacO - RBS - (6xHis) - eryF - T7 terminator | pET22b | AmpR | ColE1 + rop |
| pFB164 | araC - paraBAD - B0034 - eryG - sfGFP - (6xHis) - B0015 | pSB1C3 | CmR | ColE1 |
| pFB165 | pT7 - lacO - RBS - (6xHis) - eryH - T7 terminator | pET22b | AmpR | ColE1 + rop |
| pFB166 | pT7 - lacO - RBS - (6xHis) - eryI - T7 terminator | pET22b | AmpR | ColE1 + rop |
| pFB167 | pT7 - lacO - RBS - (6xHis) - eryR - T7 terminator | pET22b | AmpR | ColE1 + rop |
| pFB168 | J23100 - eryA-RBS - sfGFP - (6xHis) - B0015 | pSB1C3 | CmR | ColE1 |
| pFB169 | J23100 - eryB-RBS - sfGFP - (6xHis) - B0015 | pSB1C3 | CmR | ColE1 |
| pFB170 | J23100 - eryC-RBS - sfGFP - (6xHis) - B0015 | pSB1C3 | CmR | ColE1 |
| pFB171 | J23100 - eryD-RBS - sfGFP - (6xHis) - B0015 | pSB1C3 | CmR | ColE1 |
| pFB172 | J23100 - eryE-RBS - sfGFP - (6xHis) - B0015 | pSB1C3 | CmR | ColE1 |

| | | | | |
|---|---|---|---|---|
| pFB173 | J23100 - eryF-RBS - sfGFP - (6xHis) - B0015 | pSB1C3 | CmR | ColE1 |
| pFB174 | J23100 - eryG-RBS - sfGFP - (6xHis) - B0015 | pSB1C3 | CmR | ColE1 |
| pFB175 | J23100 - eryH-RBS - sfGFP - (6xHis) - B0015 | pSB1C3 | CmR | ColE1 |
| pFB176 | J23100 - eryI-RBS - sfGFP - (6xHis) - B0015 | pSB1C3 | CmR | ColE1 |
| pFB177 | pR-eryA - sfGFP - (6xHis) - B0015 | pSB1C3 | CmR | ColE1 |
| pFB178 | pR-eryB - sfGFP - (6xHis) - B0015 | pSB1C3 | CmR | ColE1 |
| pFB179 | pR-eryC - sfGFP - (6xHis) - B0015 | pSB1C3 | CmR | ColE1 |
| pFB180 | pR-eryD - sfGFP - (6xHis) - B0015 | pSB1C3 | CmR | ColE1 |
| pFB181 | pR-eryE - sfGFP - (6xHis) - B0015 | pSB1C3 | CmR | ColE1 |
| pFB182 | pR-eryF - sfGFP - (6xHis) - B0015 | pSB1C3 | CmR | ColE1 |
| pFB183 | pR-eryG - sfGFP - (6xHis) - B0015 | pSB1C3 | CmR | ColE1 |
| pFB184 | pR-eryH - sfGFP - (6xHis) - B0015 | pSB1C3 | CmR | ColE1 |
| pFB185 | pR-eryI - sfGFP - (6xHis) - B0015 | pSB1C3 | CmR | ColE1 |
| pFB186 | pEryF-732 - B0034 - sfGFP - (6xHis) - B0015 | pSB1C3 | CmR | ColE1 |
| pFB187 | pEryF-632 - B0034 - sfGFP - (6xHis) - B0015 | pSB1C3 | CmR | ColE1 |
| pFB188 | pEryF-532 - B0034 - sfGFP - (6xHis) - B0015 | pSB1C3 | CmR | ColE1 |
| pFB189 | pEryF-432 - B0034 - sfGFP - (6xHis) - B0015 | pSB1C3 | CmR | ColE1 |
| pFB190 | pEryF-332 - B0034 - sfGFP - (6xHis) - B0015 | pSB1C3 | CmR | ColE1 |
| pFB191 | pEryF-232 - B0034 - sfGFP - (6xHis) - B0015 | pSB1C3 | CmR | ColE1 |
| pFB192 | pEryF-222 - B0034 - sfGFP - (6xHis) - B0015 | pSB1C3 | CmR | ColE1 |
| pFB193 | pEryF-212 - B0034 - sfGFP - (6xHis) - B0015 | pSB1C3 | CmR | ColE1 |
| pFB194 | pEryF-202 - B0034 - sfGFP - (6xHis) - B0015 | pSB1C3 | CmR | ColE1 |
| pFB195 | pEryF-192 - B0034 - sfGFP - (6xHis) - B0015 | pSB1C3 | CmR | ColE1 |
| pFB196 | pEryF-182 - B0034 - sfGFP - (6xHis) - B0015 | pSB1C3 | CmR | ColE1 |
| pFB197 | pEryF-172 - B0034 - sfGFP - (6xHis) - B0015 | pSB1C3 | CmR | ColE1 |
| pFB198 | pEryF-162 - B0034 - sfGFP - (6xHis) - B0015 | pSB1C3 | CmR | ColE1 |
| pFB199 | pEryF-161 - B0034 - sfGFP - (6xHis) - B0015 | pSB1C3 | CmR | ColE1 |
| pFB200 | pEryF-160 - B0034 - sfGFP - (6xHis) - B0015 | pSB1C3 | CmR | ColE1 |
| pFB201 | pEryF-159 - B0034 - sfGFP - (6xHis) - B0015 | pSB1C3 | CmR | ColE1 |
| pFB202 | pEryF-158 - B0034 - sfGFP - (6xHis) - B0015 | pSB1C3 | CmR | ColE1 |
| pFB203 | pEryF-157 - B0034 - sfGFP - (6xHis) - B0015 | pSB1C3 | CmR | ColE1 |
| pFB204 | pEryF-156 - B0034 - sfGFP - (6xHis) - B0015 | pSB1C3 | CmR | ColE1 |
| pFB205 | pEryF-155 - B0034 - sfGFP - (6xHis) - B0015 | pSB1C3 | CmR | ColE1 |
| pFB206 | pEryF-154 - B0034 - sfGFP - (6xHis) - B0015 | pSB1C3 | CmR | ColE1 |
| pFB207 | pEryF-153 - B0034 - sfGFP - (6xHis) - B0015 | pSB1C3 | CmR | ColE1 |
| pFB208 | pEryF-152 - B0034 - sfGFP - (6xHis) - B0015 | pSB1C3 | CmR | ColE1 |
| pFB209 | pEryF-142 - B0034 - sfGFP - (6xHis) - B0015 | pSB1C3 | CmR | ColE1 |
| pFB210 | pEryF-132 - B0034 - sfGFP - (6xHis) - B0015 | pSB1C3 | CmR | ColE1 |
| pFB211 | pEryF-122 - B0034 - sfGFP - (6xHis) - B0015 | pSB1C3 | CmR | ColE1 |
| pFB212 | pEryF-112 - B0034 - sfGFP - (6xHis) - B0015 | pSB1C3 | CmR | ColE1 |
| pFB213 | pEryF-102 - B0034 - sfGFP - (6xHis) - B0015 | pSB1C3 | CmR | ColE1 |
| pFB214 | pEryF-159-10 - B0034 - sfGFP - (6xHis) - B0015 | pSB1C3 | CmR | ColE1 |
| pFB215 | pEryF-159-20 - B0034 - sfGFP - (6xHis) - B0015 | pSB1C3 | CmR | ColE1 |
| pFB216 | pEryF-159-30 - B0034 - sfGFP - (6xHis) - B0015 | pSB1C3 | CmR | ColE1 |

| | | | | |
|---|---|---|---|---|
| pFB217 | pEryF-159-40 - B0034 - sfGFP - (6xHis) - B0015 | pSB1C3 | CmR | ColE1 |
| pFB218 | pEryF-159-50 - B0034 - sfGFP - (6xHis) - B0015 | pSB1C3 | CmR | ColE1 |
| pFB219 | pEryF-159-60 - B0034 - sfGFP - (6xHis) - B0015 | pSB1C3 | CmR | ColE1 |
| pFB220 | pEryF-159-70 - B0034 - sfGFP - (6xHis) - B0015 | pSB1C3 | CmR | ColE1 |
| pFB221 | pEryF-159-80 - B0034 - sfGFP - (6xHis) - B0015 | pSB1C3 | CmR | ColE1 |
| pFB222 | pEryF-159-90 - B0034 - sfGFP - (6xHis) - B0015 | pSB1C3 | CmR | ColE1 |
| pFB223 | pEryF-159-100 - B0034 - sfGFP - (6xHis) - B0015 | pSB1C3 | CmR | ColE1 |
| pFB224 | pEryF-159-110 - B0034 - sfGFP - (6xHis) - B0015 | pSB1C3 | CmR | ColE1 |
| pFB225 | pEryF-159-120 - B0034 - sfGFP - (6xHis) - B0015 | pSB1C3 | CmR | ColE1 |
| pFB226 | pEryF-159-130 - B0034 - sfGFP - (6xHis) - B0015 | pSB1C3 | CmR | ColE1 |
| pFB227 | pEryF-159-140 - B0034 - sfGFP - (6xHis) - B0015 | pSB1C3 | CmR | ColE1 |
| pFB228 | pEryF-159-150 - B0034 - sfGFP - (6xHis) - B0015 | pSB1C3 | CmR | ColE1 |
| pFB229 | pEryF-37 - B0034 - sfGFP - (6xHis) - B0015 | pSB1C3 | CmR | ColE1 |
| pFB230 | pEryF-35 - B0034 - sfGFP - (6xHis) - B0015 | pSB1C3 | CmR | ColE1 |
| pFB231 | pEryF-33 - B0034 - sfGFP - (6xHis) - B0015 | pSB1C3 | CmR | ColE1 |
| pFB232 | pEryF-31 - B0034 - sfGFP - (6xHis) - B0015 | pSB1C3 | CmR | ColE1 |
| pFB233 | pEryF-29 - B0034 - sfGFP - (6xHis) - B0015 | pSB1C3 | CmR | ColE1 |
| pFB234 | pEryF-28 - B0034 - sfGFP - (6xHis) - B0015 | pSB1C3 | CmR | ColE1 |
| pFB235 | pEryF-27 - B0034 - sfGFP - (6xHis) - B0015 | pSB1C3 | CmR | ColE1 |
| pFB236 | pEryF-25 - B0034 - sfGFP - (6xHis) - B0015 | pSB1C3 | CmR | ColE1 |
| pFB237 | pEryF-23 - B0034 - sfGFP - (6xHis) - B0015 | pSB1C3 | CmR | ColE1 |
| pFB238 | pEryF-21 - B0034 - sfGFP - (6xHis) - B0015 | pSB1C3 | CmR | ColE1 |
| pFB239 | pEryR-732 - B0034 - sfGFP - (6xHis) - B0015 | pSB1C3 | CmR | ColE1 |
| pFB240 | pEryR-632 - B0034 - sfGFP - (6xHis) - B0015 | pSB1C3 | CmR | ColE1 |
| pFB241 | pEryR-532 - B0034 - sfGFP - (6xHis) - B0015 | pSB1C3 | CmR | ColE1 |
| pFB242 | pEryR-432 - B0034 - sfGFP - (6xHis) - B0015 | pSB1C3 | CmR | ColE1 |
| pFB243 | pEryR-332 - B0034 - sfGFP - (6xHis) - B0015 | pSB1C3 | CmR | ColE1 |
| pFB244 | pEryR-322 - B0034 - sfGFP - (6xHis) - B0015 | pSB1C3 | CmR | ColE1 |
| pFB245 | pEryR-312 - B0034 - sfGFP - (6xHis) - B0015 | pSB1C3 | CmR | ColE1 |
| pFB246 | pEryR-302 - B0034 - sfGFP - (6xHis) - B0015 | pSB1C3 | CmR | ColE1 |
| pFB247 | pEryR-292 - B0034 - sfGFP - (6xHis) - B0015 | pSB1C3 | CmR | ColE1 |
| pFB248 | pEryR-282 - B0034 - sfGFP - (6xHis) - B0015 | pSB1C3 | CmR | ColE1 |
| pFB249 | pEryR-272 - B0034 - sfGFP - (6xHis) - B0015 | pSB1C3 | CmR | ColE1 |
| pFB250 | pEryR-262 - B0034 - sfGFP - (6xHis) - B0015 | pSB1C3 | CmR | ColE1 |
| pFB251 | pEryR-252 - B0034 - sfGFP - (6xHis) - B0015 | pSB1C3 | CmR | ColE1 |
| pFB252 | pEryR-242 - B0034 - sfGFP - (6xHis) - B0015 | pSB1C3 | CmR | ColE1 |
| pFB253 | pEryR-232 - B0034 - sfGFP - (6xHis) - B0015 | pSB1C3 | CmR | ColE1 |
| pFB254 | pEryR-132 - B0034 - sfGFP - (6xHis) - B0015 | pSB1C3 | CmR | ColE1 |
| pFB255 | pEryR-32 - B0034 - sfGFP - (6xHis) - B0015 | pSB1C3 | CmR | ColE1 |
| pFB256 | pEry2-400 - B0034 - sfGFP - (6xHis) - B0015 | pSB1C3 | CmR | ColE1 |
| pFB257 | pEry2-300 - B0034 - sfGFP - (6xHis) - B0015 | pSB1C3 | CmR | ColE1 |
| pFB258 | pEry2-200 - B0034 - sfGFP - (6xHis) - B0015 | pSB1C3 | CmR | ColE1 |
| pFB259 | pEry2-100 - B0034 - sfGFP - (6xHis) - B0015 | pSB1C3 | CmR | ColE1 |
| pFB260 | J23105 - B0034 - eryD | pFB1 | AmpR | p15A |

| | | | | |
|---|---|---|---|---|
| pFB261 | J23106 - B0034 - eryD | pFB1 | AmpR | p15A |
| pFB262 | J23109 - B0034 - eryD | pFB1 | AmpR | p15A |
| pFB263 | J23113 - B0034 - eryD | pFB1 | AmpR | p15A |
| pFB264 | J23114 - B0034 - eryD | pFB1 | AmpR | p15A |
| pFB265 | J23105 - B0034 - eryR | pFB1 | AmpR | p15A |
| pFB266 | J23106 - B0034 - eryR | pFB1 | AmpR | p15A |
| pFB267 | J23109 - B0034 - eryR | pFB1 | AmpR | p15A |
| pFB268 | J23113 - B0034 - eryR | pFB1 | AmpR | p15A |
| pFB269 | J23114 - B0034 - eryR | pFB1 | AmpR | p15A |
| pFB270 | J23100 - eryO - B0034 - sfGFP - (6xHis) - B0015 | pSB1C3 | CmR | ColE1 |
| pFB271 | J23105 - eryO - B0034 - sfGFP - (6xHis) - B0015 | pSB1C3 | CmR | ColE1 |
| pFB272 | J23106 - eryO - B0034 - sfGFP - (6xHis) - B0015 | pSB1C3 | CmR | ColE1 |
| pFB273 | J23109 - eryO - B0034 - sfGFP - (6xHis) - B0015 | pSB1C3 | CmR | ColE1 |
| pFB274 | J23113 - eryO - B0034 - sfGFP - (6xHis) - B0015 | pSB1C3 | CmR | ColE1 |
| pFB275 | J23114 - eryO - B0034 - sfGFP - (6xHis) - B0015 | pSB1C3 | CmR | ColE1 |
| pFB276 | pT7 - eryO - RBS - sfGFP - (6xHis) - B0015 | pSB1C3 | CmR | ColE1 |
| pFB277 | pT7 - RBS - sfGFP - (6xHis) - B0015 | pSB1C3 | CmR | ColE1 |
| pFB278 | alsC-homoL - FRT - KanR - FRT - alsC-homoR | pKD4 | AmpR / KanR | R6K |
| pFB279 | araH-homoL - FRT - KanR - FRT - araH-homoR | pKD4 | AmpR / KanR | R6K |
| pFB280 | glpF-homoL - FRT - KanR - FRT - glpF-homoR | pKD4 | AmpR / KanR | R6K |
| pFB281 | malG-homoL - FRT - KanR - FRT - malG-homoR | pKD4 | AmpR / KanR | R6K |
| pFB282 | mglC-homoL - FRT - KanR - FRT - mglC-homoR | pKD4 | AmpR / KanR | R6K |
| pFB283 | rbsC-homoL - FRT - KanR - FRT - rbsC-homoR | pKD4 | AmpR / KanR | R6K |
| pFB284 | xylH-homoL - FRT - KanR - FRT - xylH-homoR | pKD4 | AmpR / KanR | R6K |
| pFB285 | J23100 - B0034 - sfGFP - (6xHis) - B0015 | pFB1 | AmpR | p15A |
| pFB286 | J23100 - B0034 - sfGFP - (6xHis) - B0015 | pSB1C3 | CmR | ColE1 |
| pFB287 | J23105 - B0034 - sfGFP - (6xHis) - B0015 | pSB1C3 | CmR | ColE1 |
| pFB288 | J23106 - B0034 - sfGFP - (6xHis) - B0015 | pSB1C3 | CmR | ColE1 |
| pFB289 | J23109 - B0034 - sfGFP - (6xHis) - B0015 | pSB1C3 | CmR | ColE1 |
| pFB290 | J23113 - B0034 - sfGFP - (6xHis) - B0015 | pSB1C3 | CmR | ColE1 |
| pFB291 | J23114 - B0034 - sfGFP - (6xHis) - B0015 | pSB1C3 | CmR | ColE1 |
| pFB292 | pEryF-32 - B0034 - sfGFP - (6xHis) - B0015 | pSB1C3 | CmR | ColE1 |
| pFB293 | J23105 - B0034 - talA - tktB | pFB1 | AmpR | p15A |
| pFB294 | J23106 - B0034 - talA - tktB | pFB1 | AmpR | p15A |
| pFB295 | J23109 - B0034 - talA - tktB | pFB1 | AmpR | p15A |
| pFB296 | J23113 - B0034 - talA - tktB | pFB1 | AmpR | p15A |
| pFB297 | J23114 - B0034 - talA - tktB | pFB1 | AmpR | p15A |
| pFB298 | J23105 - B0034 - talB | pFB1 | AmpR | p15A |
| pFB299 | J23106 - B0034 - talB | pFB1 | AmpR | p15A |
| pFB300 | J23113 - B0034 - talB | pFB1 | AmpR | p15A |
| pFB301 | J23114 - B0034 - talB | pFB1 | AmpR | p15A |
| pFB302 | J23105 - B0034 - tktA | pFB1 | AmpR | p15A |
| pFB303 | J23106 - B0034 - tktA | pFB1 | AmpR | p15A |
| pFB304 | J23113 - B0034 - tktA | pFB1 | AmpR | p15A |

| | | | | |
|---|---|---|---|---|
| pFB305 | J23114 - B0034 - tktA | pFB1 | AmpR | p15A |
| pFB306 | yiaMNO-homoL - FRT - KanR - FRT - yiaMNO-homoR | pKD4 | AmpR / KanR | R6K |
| pFB307 | fryBCA-homoL - FRT - KanR - FRT - fryBCA-homoR | pKD4 | AmpR / KanR | R6K |
| pFB308 | garP-homoL - FRT - KanR - FRT - garP-homoR | pKD4 | AmpR / KanR | R6K |
| pFB309 | gntT-homoL - FRT - KanR - FRT - gntT-homoR | pKD4 | AmpR / KanR | R6K |
| pFB310 | ulaABC-homoL - FRT - KanR - FRT - ulaABC-homoR | pKD4 | AmpR / KanR | R6K |
| pFB311 | agaBCD-homoL - FRT - KanR - FRT - agaBCD-homoR | pKD4 | AmpR / KanR | R6K |
| pFB312 | agaVW-homoL - FRT - KanR - FRT - agaVW-homoR | pKD4 | AmpR / KanR | R6K |
| pFB313 | ydjK-homoL - FRT - KanR - FRT - ydjK-homoR | pKD4 | AmpR / KanR | R6K |
| pFB314 | yjiJ-homoL - FRT - KanR - FRT - yjiJ-homoR | pKD4 | AmpR / KanR | R6K |
| pFB315 | yagG-homoL - FRT - KanR - FRT - yagG-homoR | pKD4 | AmpR / KanR | R6K |
| pFB316 | yaaU-homoL - FRT - KanR - FRT - yaaU-homoR | pKD4 | AmpR / KanR | R6K |
| pFB317 | yicJ-homoL - FRT - KanR - FRT - yicJ-homoR | pKD4 | AmpR / KanR | R6K |
| pFB318 | frvAB-homoL - FRT - KanR - FRT - frvAB-homoR | pKD4 | AmpR / KanR | R6K |
| pFB319 | frwCB-homoL - FRT - KanR - FRT - frwCB-homoR | pKD4 | AmpR / KanR | R6K |
| pFB320 | nanT-homoL - FRT - KanR - FRT - nanT-homoR | pKD4 | AmpR / KanR | R6K |
| pFB321 | yfaV-homoL - FRT - KanR - FRT - yfaV-homoR | pKD4 | AmpR / KanR | R6K |
| pFB322 | sgcBCA-homoL - FRT - KanR - FRT - sgcBCA-homoR | pKD4 | AmpR / KanR | R6K |
| pFB323 | dgoT-homoL - FRT - KanR - FRT - dgoT-homoR | pKD4 | AmpR / KanR | R6K |
| pFB324 | mngA-homoL - FRT - KanR - FRT - mngA-homoR | pKD4 | AmpR / KanR | R6K |
| pFB325 | ugpBAEC-homoL - FRT - KanR - FRT - ugpBAEC-homoR | pKD4 | AmpR / KanR | R6K |
| pFB326 | ycjNOP-homoL - FRT - KanR - FRT - ycjNOP-homoR | pKD4 | AmpR / KanR | R6K |
| pFB327 | uhpT-homoL - FRT - KanR - FRT - uhpT-homoR | pKD4 | AmpR / KanR | R6K |
| pFB328 | ydjE-homoL - FRT - KanR - FRT - ydjE-homoR | pKD4 | AmpR / KanR | R6K |
| pFB329 | yphFED-homoL - FRT - KanR - FRT - yphFED-homoR | pKD4 | AmpR / KanR | R6K |
| pFB330 | yhjDE-homoL - FRT - KanR - FRT - yhjDE-homoR | pKD4 | AmpR / KanR | R6K |
| pFB331 | ytfT-homoL - FRT - KanR - FRT - ytfT-homoR | pKD4 | AmpR / KanR | R6K |
| pFB332 | srlAEB-homoL - FRT - KanR - FRT - srlAEB-homoR | pKD4 | AmpR / KanR | R6K |
| pFB333 | yfcJ-homoL - FRT - KanR - FRT - yfcJ-homoR | pKD4 | AmpR / KanR | R6K |
| pFB334 | lsrACDB-homoL - FRT - KanR - FRT - lsrACDB-homoR | pKD4 | AmpR / KanR | R6K |
| pFB335 | melB-homoL - FRT - KanR - FRT - melB-homoR | pKD4 | AmpR / KanR | R6K |
| pFB336 | mhpT-homoL - FRT - KanR - FRT - mhpT-homoR | pKD4 | AmpR / KanR | R6K |
| pFB337 | yjhF-homoL - FRT - KanR - FRT - yjhF-homoR | pKD4 | AmpR / KanR | R6K |
| pFB338 | glvCB-homoL - FRT - KanR - FRT - glvCB-homoR | pKD4 | AmpR / KanR | R6K |
| pFB339 | yidK-homoL - FRT - KanR - FRT - yidK-homoR | pKD4 | AmpR / KanR | R6K |
| pFB340 | glcC - pglcD - B0034 - sfGFP - (6xHis) - B0015 | pFB1 | AmpR | p15A |

Unless otherwise mentioned, all plasmids above were constructed by Gibson Assembly and conducted in the strain Mach1-T1. All plasmids with R6K origin of replication were conducted in the strain DH5α λpir.

All constructed plasmids were sequenced correctly.

All “6xHis” represents a “six-repeat-histidine” amino acid sequence located in *N*-terminus or *C*-terminus of each protein.

**Supplementary Table 6. Primers.**

| Primer name | Sequence (5' to 3') | Purpose | Reference |
|---|---|---|---|
| VF2 | tgccacctgacgtctaagaa | pSB1C3 and pFB1 derived plasmids sequencing, colony PCR | 1 |
| VR | attaccgcctttgagtgagc | pSB1C3 and pFB1 derived plasmids sequencing, colony PCR | 1 |
| 27F | agagtttgatcctggctcag | Prokaryotic organism 16s rRNA sequence amplification | This work |
| 1492R | ggttaccttgttacgactt | Prokaryotic organism 16s rRNA sequence amplification | This work |

All primers above were synthesized by GENEWIZ.

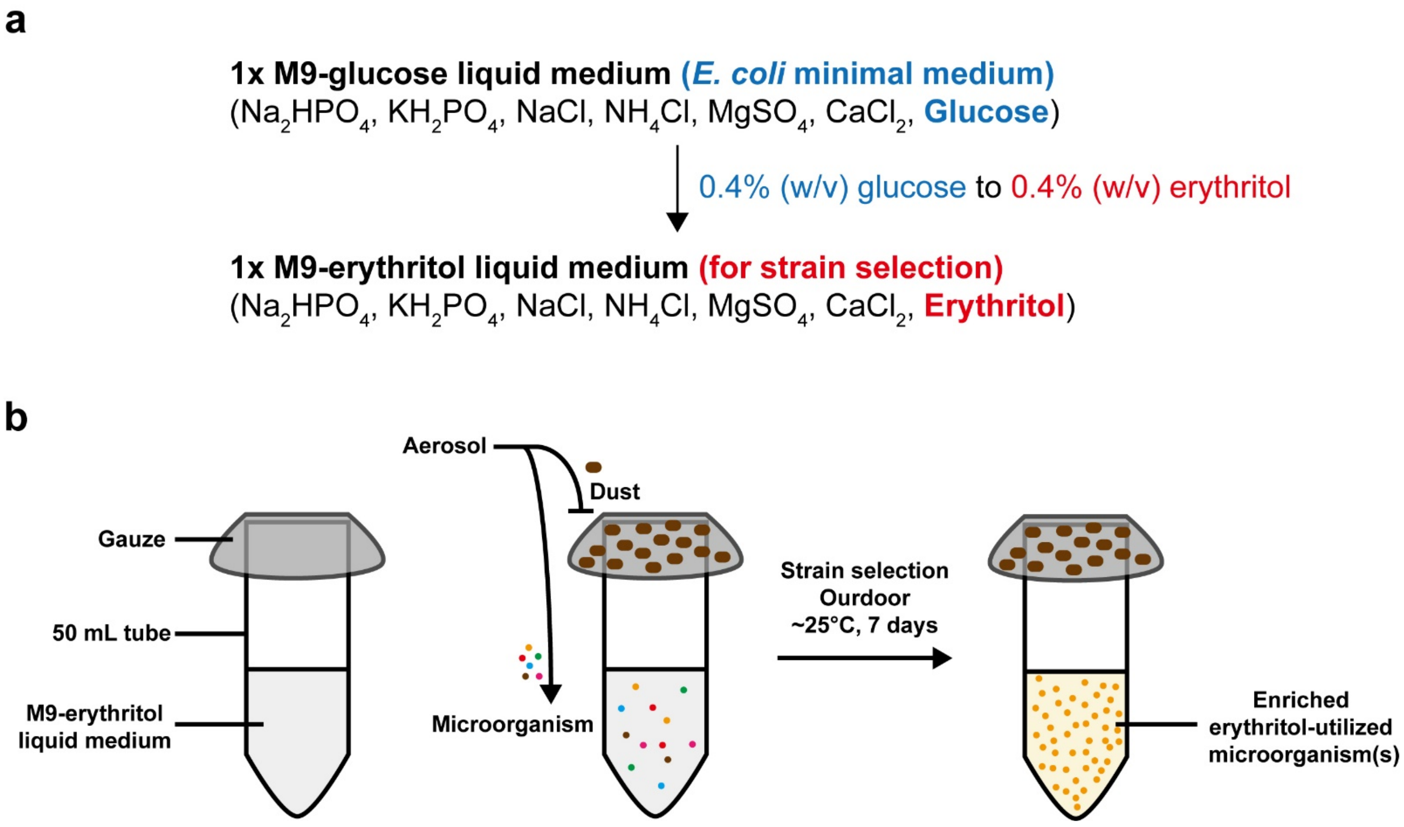

**Supplementary Figure 1. Isolation of microorganisms growing in the erythritol-based M9 medium.**

(**a**) 1x M9-erythritol liquid medium is derived from 1x M9-glucose liquid medium by changing 0.4% (w/v) glucose into 0.4% (w/v) erythritol and other components are the same. 1 L 1x M9-erythritol liquid medium consists of 12.8 g $Na_2HPO_4 \bullet 7H_2O$, 3 g $KH_2PO_4$, 0.5 g NaCl, 1 g $NH_4Cl$, 200 µL 1 M $MgSO_4$, 10 µL 1 M $CaCl_2$, and 4 g erythritol.

(**b**) Schematic of screening erythritol-utilized microorganisms. A 50 mL centrifuge tube (without lid) contains approximately 25 mL 1x M9-erythritol liquid medium. The covered gauze can block aerosol dust, but aerosol microorganisms can fall into the liquid medium. After 7 days of outdoor stationary culture (approximately 25°C), the microorganisms which can utilize erythritol as sole carbon source were screened out and enriched, then the liquid culture was separated on LB-agar plate for picking up single colony.

**a**

**16s rRNA BLAST alignments (primer pair: 27F / 1492R)**

| Strain | Query coverage | Percent identity | Accession |
|---|---|---|---|
| Ochrobactrum sp. strain SNH-K10 | 100% | 100% | MN493878.1 |
| Ochrobactrum anthropi strain SCA-1 | 100% | 100% | MH448104.1 |
| Ochrobactrum sp. TE-101 | 100% | 100% | KC998944.1 |
| Ochrobactrum sp. C14 | 100% | 100% | KT361086.1 |
| Ochrobactrum anthropi strain LSP34 | 100% | 100% | MW485664.1 |
| Ochrobactrum sp. strain LSC9 | 100% | 100% | MW485524.1 |
| Ochrobactrum anthropi strain DP5 | 100% | 99.92% | MT534544.1 |
| Ochrobactrum sp. strain JWDC2 | 100% | 99.92% | MT448953.1 |
| Ochrobactrum sp. strain JWJC3 | 100% | 99.92% | MT448947.1 |
| Ochrobactrum anthropi strain L1-W | 100% | 99.92% | MT093466.1 |
| ...... | ...... | ...... | ...... |

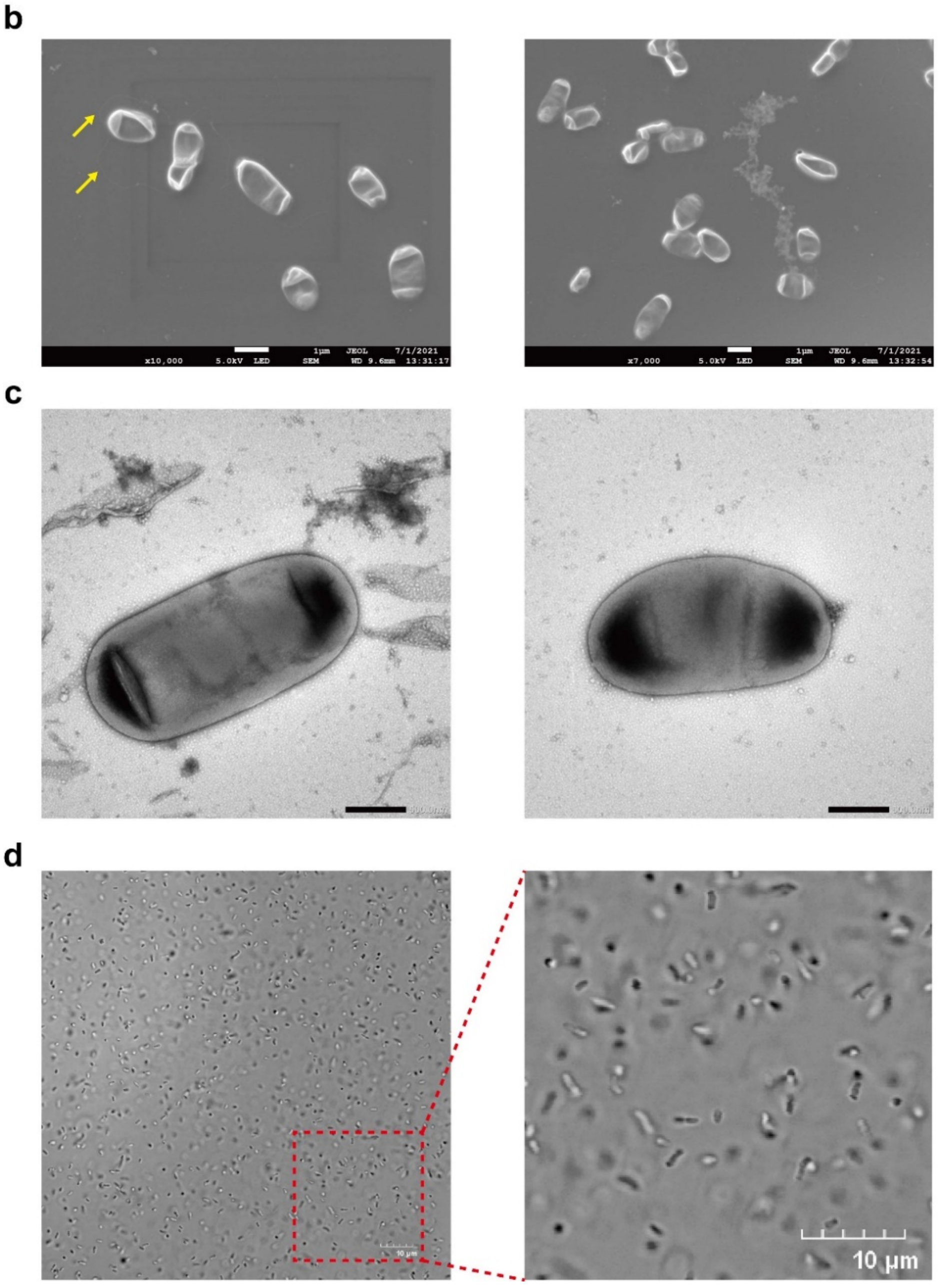

**Supplementary Figure 2. Characterization of the isolated *Ochrobactrum* spp. strain.**

(**a**) BLAST[4,5] alignments of screened-out *Ochrobactrum* spp. 16s rRNA sequence.

(**b**) Scanning electron microscopy images of *Ochrobactrum* spp.. Yellow arrows indicate peritrichous flagella.

(**c**) Transmission electron microscopy images of *Ochrobactrum* spp..

(**d**) Confocal laser scanning microscopy images of *Ochrobactrum* spp..

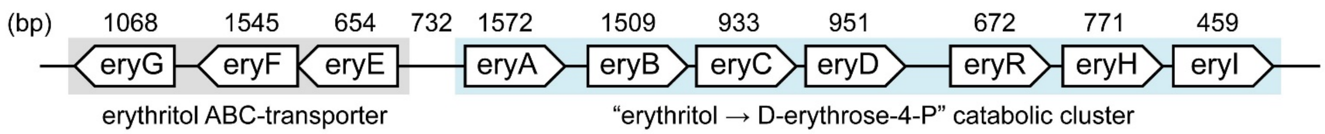

**eryE:** DUF (Domain of Unknown Function) 2291 family protein (peripheral erythritol-binding protein)
**eryF:** Sugar ABC transporter ATP-binding protein (erythritol ABC transporter ATP-binding protein)
**eryG:** ABC transporter permease (erythritol ABC transporter permease)

**eryA:** Carbohydrate kinase (erythritol kinase)
**eryB:** Glycerol-3-phosphate dehydrogenase (erythritol-4-phosphate dehydrogenase)
**eryC:** TIM barrel protein (L-3-tetrulose-4-phosphate to D-3-tetrulose-4-phosphate isomerase)
**eryD:** Sugar-binding transcriptional regulator (erythritol-binding transcriptional regulator)
**eryR:** DeoR/GlpR family DNA-binding transcription regulator (Unknown)
**eryH:** Triose-phosphate isomerase (D-3-tetrulose-4-phosphate to D-erythrulose-4-phosphate isomerase)
**eryI:** RpiB/LacA/LacB family sugar-phosphate isomerase
(D-erythrulose-4-phosphate to D-erythrose-4-phosphate isomerase)

**Supplementary Figure 3. Annotation of erythritol catabolism-associated genes.**
Schematic of erythritol catabolism-associated cluster is shown with ten different genes. Black annotations represent “protein BLAST alignment” results of each gene, red annotations represent each gene’s detailed function.

**eryA nucleotide BLAST alignments**

| Strain | Query coverage | Percent identity | Accession |
|---|---|---|---|
| Brucella anthropi strain T16R-87 | 100% | 99.81% | CP044971.1 |
| Ochrobactrum sp. MT180101 | 100% | 98.28% | CP061773.1 |
| Ochrobactrum anthropi | 100% | 98.03% | LT671862.1 |
| Brucella anthropi strain PBO | 100% | 98.03% | CP064062.1 |
| Ochrobactrum anthropi strain OAB | 100% | 97.96% | CP008819.1 |
| Brucella anthropi strain FDAARGOS_1039 | 100% | 97.96% | CP066053.1 |
| Ochrobactrum anthropi ATCC 49188 | 100% | 97.96% | CP000759.1 |
| Ochrobactrum sp. WY7 | 100% | 97.77% | CP049797.1 |
| Ochrobactrum sp. EEELCW01 | 100% | 97.65% | CP047599.1 |
| Brucella intermedia strain ZJ499 | 98% | 86.87% | CP061040.1 |

**eryB nucleotide BLAST alignments**

| Strain | Query coverage | Percent identity | Accession |
|---|---|---|---|
| Brucella anthropi strain T16R-87 | 100% | 99.93% | CP044971.1 |
| Ochrobactrum sp. EEELCW01 | 100% | 98.54% | CP047599.1 |
| Ochrobactrum sp. MT180101 | 100% | 98.54% | CP061773.1 |
| Ochrobactrum anthropi | 100% | 98.41% | LT671862.1 |
| Brucella anthropi strain PBO | 100% | 98.34% | CP064062.1 |
| Ochrobactrum anthropi strain OAB | 100% | 96.95% | CP008819.1 |
| Brucella anthropi strain FDAARGOS_1039 | 100% | 96.95% | CP066053.1 |
| Ochrobactrum anthropi ATCC 49188 | 100% | 96.95% | CP000759.1 |
| Ochrobactrum sp. WY7 | 100% | 95.96% | CP049797.1 |
| Brucella intermedia strain ZJ499 | 100% | 92.58% | CP061040.1 |

**eryC nucleotide BLAST alignments**

| Strain | Query coverage | Percent identity | Accession |
|---|---|---|---|
| Brucella anthropi strain T16R-87 | 100% | 100.00% | CP044971.1 |
| Ochrobactrum anthropi | 100% | 99.25% | LT671862.1 |
| Brucella anthropi strain PBO | 100% | 99.25% | CP064062.1 |
| Ochrobactrum sp. MT180101 | 100% | 98.50% | CP061773.1 |
| Ochrobactrum sp. EEELCW01 | 99% | 97.53% | CP047599.1 |
| Ochrobactrum anthropi strain OAB | 100% | 97.21% | CP008819.1 |
| Brucella anthropi strain FDAARGOS_1039 | 100% | 97.21% | CP066053.1 |
| Ochrobactrum anthropi ATCC 49188 | 100% | 97.21% | CP000759.1 |
| Ochrobactrum sp. WY7 | 100% | 97.11% | CP049797.1 |
| Brucella sp. 09RB8471 | 99% | 90.57% | CP019346.1 |

**eryD nucleotide BLAST alignments**

| Strain | Query coverage | Percent identity | Accession |
|---|---|---|---|
| Brucella anthropi strain T16R-87 | 100% | 100.00% | CP044971.1 |
| Ochrobactrum anthropi | 100% | 97.58% | LT671862.1 |
| Brucella anthropi strain PBO | 100% | 97.48% | CP064062.1 |
| Ochrobactrum anthropi strain OAB | 100% | 97.16% | CP008819.1 |
| Brucella anthropi strain FDAARGOS_1039 | 100% | 97.16% | CP066053.1 |
| Ochrobactrum anthropi ATCC 49188 | 100% | 97.16% | CP000759.1 |
| Ochrobactrum sp. WY7 | 100% | 96.85% | CP049797.1 |
| Ochrobactrum sp. EEELCW01 | 100% | 96.42% | CP047599.1 |
| Ochrobactrum sp. MT180101 | 100% | 96.42% | CP061773.1 |
| Brucella intermedia strain ZJ499 | 96% | 87.92% | CP061040.1 |

**eryE nucleotide BLAST alignments**

| Strain | Query coverage | Percent identity | Accession |
|---|---|---|---|
| Brucella anthropi strain T16R-87 | 100% | 100.00% | CP044971.1 |
| Ochrobactrum sp. MT180101 | 100% | 100.00% | CP061773.1 |
| Ochrobactrum sp. WY7 | 100% | 98.78% | CP049797.1 |
| Ochrobactrum anthropi strain OAB | 100% | 98.62% | CP008819.1 |
| Brucella anthropi strain FDAARGOS_1039 | 100% | 98.62% | CP066053.1 |
| Ochrobactrum anthropi ATCC 49188 | 100% | 98.62% | CP000759.1 |
| Brucella anthropi strain PBO | 100% | 98.01% | CP064062.1 |
| Ochrobactrum anthropi | 100% | 97.71% | LT671862.1 |
| Ochrobactrum sp. EEELCW01 | 100% | 96.94% | CP047599.1 |
| Brucella intermedia strain ZJ499 | 100% | 90.38% | CP061040.1 |

**eryF nucleotide BLAST alignments**

| Strain | Query coverage | Percent identity | Accession |
|---|---|---|---|
| Brucella anthropi strain T16R-87 | 100% | 99.87% | CP044971.1 |
| Ochrobactrum anthropi | 100% | 98.77% | LT671862.1 |
| Ochrobactrum sp. MT180101 | 100% | 98.77% | CP061773.1 |
| Ochrobactrum anthropi strain OAB | 100% | 98.45% | CP008819.1 |
| Brucella anthropi strain FDAARGOS_1039 | 100% | 98.45% | CP066053.1 |
| Brucella anthropi strain PBO chromosome 1 | 100% | 98.45% | CP064062.1 |
| Ochrobactrum anthropi ATCC 49188 | 100% | 98.45% | CP000759.1 |
| Ochrobactrum sp. EEELCW01 | 100% | 98.38% | CP047599.1 |
| Ochrobactrum sp. WY7 | 100% | 98.25% | CP049797.1 |
| Brucella intermedia strain ZJ499 | 99% | 93.13% | CP061040.1 |

**eryG nucleotide BLAST alignments**

| Strain | Query coverage | Percent identity | Accession |
|---|---|---|---|
| Ochrobactrum sp. MT180101 | 100% | 100.00% | CP061773.1 |
| Brucella anthropi strain T16R-87 | 100% | 99.91% | CP044971.1 |
| Brucella anthropi strain PBO | 100% | 98.78% | CP064062.1 |
| Ochrobactrum anthropi | 100% | 98.13% | LT671862.1 |
| Ochrobactrum anthropi strain OAB | 100% | 98.13% | CP008819.1 |
| Ochrobactrum sp. WY7 | 100% | 98.13% | CP049797.1 |
| Brucella anthropi strain FDAARGOS_1039 | 100% | 98.13% | CP066053.1 |
| Ochrobactrum anthropi ATCC 49188 | 100% | 98.13% | CP000759.1 |
| Ochrobactrum sp. EEELCW01 | 100% | 97.85% | CP047599.1 |
| Brucella intermedia strain ZJ499 | 99% | 91.68% | CP061040.1 |

**eryH nucleotide BLAST alignments**

| Strain | Query coverage | Percent identity | Accession |
|---|---|---|---|
| Brucella anthropi strain T16R-87 | 100% | 100.00% | CP044971.1 |
| Ochrobactrum anthropi | 100% | 100.00% | LT671862.1 |
| Ochrobactrum sp. EEELCW01 | 100% | 99.22% | CP047599.1 |
| Ochrobactrum sp. MT180101 | 100% | 97.67% | CP061773.1 |
| Ochrobactrum anthropi strain OAB | 100% | 96.89% | CP008819.1 |
| Ochrobactrum sp. WY7 | 100% | 96.89% | CP049797.1 |
| Brucella anthropi strain FDAARGOS_1039 | 100% | 96.89% | CP066053.1 |
| Ochrobactrum anthropi ATCC 49188 | 100% | 96.89% | CP000759.1 |
| Brucella anthropi strain PBO | 100% | 96.24% | CP064062.1 |
| Brucella sp. BO3 | 100% | 87.81% | CP047233.1 |

**eryI nucleotide BLAST alignments**

| Strain | Query coverage | Percent identity | Accession |
|---|---|---|---|
| Brucella anthropi strain T16R-87 | 100% | 100.00% | CP044971.1 |
| Ochrobactrum anthropi genome assembly | 100% | 100.00% | LT671862.1 |
| Ochrobactrum sp. MT180101 | 100% | 99.56% | CP061773.1 |
| Ochrobactrum sp. WY7 | 100% | 99.13% | CP049797.1 |
| Ochrobactrum anthropi strain OAB | 100% | 98.91% | CP008819.1 |
| Brucella anthropi strain FDAARGOS_1039 | 100% | 98.91% | CP066053.1 |
| Brucella anthropi strain PBO chromosome 1 | 100% | 98.91% | CP064062.1 |
| Ochrobactrum anthropi ATCC 49188 | 100% | 98.91% | CP000759.1 |
| Ochrobactrum sp. EEELCW01 | 100% | 97.39% | CP047599.1 |
| Ochrobactrum sp. PW1 | 99% | 93.86% | LC171366.1 |

**eryR nucleotide BLAST alignments**

| Strain | Query coverage | Percent identity | Accession |
|---|---|---|---|
| Brucella anthropi strain T16R-87 | 100% | 100.00% | CP044971.1 |
| Ochrobactrum anthropi | 100% | 100.00% | LT671862.1 |
| Ochrobactrum sp. EEELCW01 | 100% | 100.00% | CP047599.1 |
| Brucella anthropi strain PBO | 99% | 98.66% | CP064062.1 |
| Ochrobactrum sp. MT180101 | 100% | 98.51% | CP061773.1 |
| Ochrobactrum anthropi strain OAB | 100% | 98.36% | CP008819.1 |
| Ochrobactrum sp. WY7 | 100% | 98.36% | CP049797.1 |
| Brucella anthropi strain FDAARGOS_1039 | 100% | 98.36% | CP066053.1 |
| Ochrobactrum anthropi ATCC 49188 | 100% | 98.36% | CP000759.1 |
| Ochrobactrum sp. PW1 | 94% | 88.89% | LC171366.1 |

**Supplementary Figure 4. Nucleotide BLAST alignments of erythritol catabolism-associated genes.**
Top ten nucleotide alignments of every erythritol catabolism-associated genes (eryA/B/C/D/E/F/G/H/I/R) in *Ochrobactrum* spp..

**eryA protein BLAST alignments**

| Strain | Query coverage | Percent identity | Accession |
|---|---|---|---|
| Brucella anthropi | 99% | 99.81% | WP_151615846.1 |
| Brucella/Ochrobactrum group | 99% | 99.62% | WP_036587141.1 |
| Brucella anthropi | 99% | 99.43% | WP_011982654.1 |
| Ochrobactrum sp. UNC390CL2Tsu3S39 | 99% | 99.24% | WP_029928109.1 |
| Brucella/Ochrobactrum group | 99% | 99.24% | WP_209418660.1 |
| Brucella anthropi | 99% | 99.04% | WP_151607003.1 |
| Brucella anthropi | 99% | 98.85% | WP_151612751.1 |
| Brucella/Ochrobactrum group | 99% | 99.04% | WP_094515042.1 |
| Brucella/Ochrobactrum group | 99% | 99.04% | WP_010660871.1 |
| Brucella tritici | 99% | 97.90% | WP_151557001.1 |

**eryB protein BLAST alignments**

| Strain | Query coverage | Percent identity | Accession |
|---|---|---|---|
| Brucella anthropi | 99% | 99.80% | WP_036587140.1 |
| Brucella/Ochrobactrum group | 99% | 100.00% | WP_043061612.1 |
| Brucella anthropi | 99% | 99.60% | WP_151663531.1 |
| Brucella/Ochrobactrum group | 99% | 99.60% | WP_094515041.1 |
| Brucella anthropi | 99% | 99.40% | MBE0559574.1 |
| Ochrobactrum sp. EEELCW01 | 99% | 99.40% | WP_209418661.1 |
| Brucella anthropi | 99% | 99.20% | WP_011982653.1 |
| Ochrobactrum sp. UNC390CL2Tsu3S39 | 99% | 99.20% | WP_029928112.1 |
| Brucella anthropi | 99% | 99.40% | WP_151614594.1 |
| Brucella anthropi | 99% | 99.00% | WP_061346705.1 |

**eryC protein BLAST alignments**

| Strain | Query coverage | Percent identity | Accession |
|---|---|---|---|
| Brucella/Ochrobactrum group | 99% | 100.00% | WP_036587138.1 |
| Brucella/Ochrobactrum group | 99% | 99.68% | WP_010660869.1 |
| Brucella anthropi | 99% | 99.35% | WP_213506816.1 |
| Brucella anthropi | 99% | 99.35% | WP_125300786.1 |
| Ochrobactrum sp. EEELCW01 | 99% | 99.03% | WP_209418662.1 |
| Ochrobactrum sp. UNC390CL2Tsu3S39 | 99% | 98.71% | WP_029928115.1 |
| Brucella/Ochrobactrum group | 99% | 99.03% | WP_094515040.1 |
| Brucella anthropi | 99% | 99.03% | WP_011982652.1 |
| Brucella anthropi | 99% | 98.71% | WP_151607006.1 |
| Brucella anthropi | 99% | 98.39% | WP_061346706.1 |

**eryD protein BLAST alignments**

| Strain | Query coverage | Percent identity | Accession |
|---|---|---|---|
| Brucella/Ochrobactrum group | 94% | 100.00% | WP_011982651.1 |
| Brucella/Ochrobactrum group | 94% | 99.67% | WP_010660868.1 |
| Brucella/Ochrobactrum group | 94% | 99.67% | WP_094515039.1 |
| Brucella anthropi | 94% | 99.34% | WP_194531703.1 |
| Brucella/Ochrobactrum group | 94% | 99.34% | WP_151607008.1 |
| Brucella anthropi | 94% | 99.67% | WP_125300787.1 |
| Brucella tritici | 94% | 99.34% | WP_151557007.1 |
| Brucella tritici | 94% | 99.34% | WP_151676965.1 |
| Brucella anthropi | 94% | 99.34% | WP_061346707.1 |
| Brucella/Ochrobactrum group | 94% | 99.00% | WP_109986727.1 |

**eryE protein BLAST alignments**

| Strain | Query coverage | Percent identity | Accession |
|---|---|---|---|
| Brucella/Ochrobactrum group | 99% | 100.00% | WP_011982655.1 |
| Brucella/Ochrobactrum group | 99% | 99.54% | WP_010660872.1 |
| Brucella/Ochrobactrum group | 99% | 97.24% | WP_121986228.1 |
| Brucella anthropi | 99% | 99.54% | WP_125334792.1 |
| Brucella/Ochrobactrum group | 99% | 99.08% | WP_105529314.1 |
| Brucella anthropi | 99% | 99.08% | WP_194531702.1 |
| unclassified Ochrobactrum | 99% | 96.77% | WP_114215989.1 |
| Brucella cytisi | 99% | 98.62% | WP_071630412.1 |
| Brucella oryzae | 99% | 97.24% | WP_211693538.1 |
| Brucella oryzae | 99% | 96.31% | WP_104755620.1 |

**eryF protein BLAST alignments**

| Strain | Query coverage | Percent identity | Accession |
|---|---|---|---|
| Brucella/Ochrobactrum group | 99% | 100.00% | WP_036587143.1 |
| Brucella anthropi | 99% | 99.81% | WP_011982656.1 |
| Brucella/Ochrobactrum group | 99% | 99.61% | WP_010660873.1 |
| Brucella/Ochrobactrum group | 99% | 99.61% | WP_151607001.1 |
| Brucella/Ochrobactrum group | 99% | 99.42% | WP_094515043.1 |
| Ochrobactrum sp. MYb49 | 99% | 99.42% | WP_105529313.1 |
| Brucella anthropi | 99% | 99.42% | WP_061346701.1 |
| Ochrobactrum sp. UNC390CL2Tsu3S39 | 99% | 99.42% | WP_029928107.1 |
| Brucella cytisi | 99% | 99.03% | WP_071630413.1 |
| Brucella cytisi | 99% | 98.83% | NKC51647.1 |

**eryG protein BLAST alignments**

| Strain | Query coverage | Percent identity | Accession |
|---|---|---|---|
| Brucella/Ochrobactrum group | 97% | 100.00% | WP_010660874.1 |
| Brucella/Ochrobactrum group | 97% | 99.71% | WP_151663530.1 |
| Brucella pecoris | 97% | 99.71% | WP_140020911.1 |
| Brucella/Ochrobactrum group | 97% | 99.42% | WP_114215987.1 |
| Brucella/Ochrobactrum group | 97% | 99.42% | WP_061346700.1 |
| Brucella oryzae | 97% | 99.42% | WP_104755618.1 |
| Brucella daejeonensis | 97% | 98.55% | WP_183651231.1 |
| Brucella vulpis | 97% | 98.27% | CUW45266.1 |
| Ochrobactrum sp. CGA5 | 97% | 98.27% | WP_139975251.1 |
| Brucella | 97% | 98.55% | WP_008511001.1 |

**eryH protein BLAST alignments**

| Strain | Query coverage | Percent identity | Accession |
|---|---|---|---|
| Brucella/Ochrobactrum group | 99% | 100.00% | WP_036587137.1 |
| Brucella/Ochrobactrum group | 99% | 99.22% | WP_029928118.1 |
| Brucella/Ochrobactrum group | 99% | 99.22% | WP_151607009.1 |
| Brucella anthropi | 99% | 98.83% | WP_151663532.1 |
| Brucella lupini | 99% | 98.83% | WP_094515038.1 |
| Brucella/Ochrobactrum group | 99% | 98.44% | WP_036582067.1 |
| Brucella anthropi | 99% | 98.05% | WP_125300788.1 |
| Brucella anthropi | 99% | 98.05% | WP_061346709.1 |
| Brucella anthropi | 99% | 98.05% | WP_010660866.1 |
| Brucella anthropi | 99% | 97.66% | WP_011982649.1 |

**eryI protein BLAST alignments**

| Strain | Query coverage | Percent identity | Accession |
|---|---|---|---|
| Brucella anthropi ATCC 49188 | 99% | 100.00% | ABS15613.1 |
| Brucella/Ochrobactrum group | 98% | 100.00% | WP_029928121.1 |
| Brucella cytisi | 98% | 99.34% | WP_071630405.1 |
| Brucella tritici | 98% | 99.34% | WP_151557013.1 |
| Brucella | 98% | 99.34% | WP_010660865.1 |
| Brucella anthropi | 98% | 99.34% | WP_125300863.1 |
| Brucella/Ochrobactrum group | 98% | 98.68% | WP_151607012.1 |
| unclassified Ochrobactrum | 98% | 98.67% | WP_114215994.1 |
| Brucella melitensis bv. 1 str. 16M | 98% | 97.35% | 5IFZ_A |
| Brucella abortus str. 2308 | 98% | 97.35% | EEP61887.1 |

**eryR protein BLAST alignments**

| Strain | Query coverage | Percent identity | Accession |
|---|---|---|---|
| Brucella anthropi | 99% | 100.00% | WP_210342020.1 |
| Brucella anthropi | 99% | 100.00% | WP_210271813.1 |
| Brucella anthropi | 99% | 100.00% | HBQ33941.1 |
| Brucella/Ochrobactrum group | 99% | 100.00% | WP_010660867.1 |
| Ochrobactrum sp. P20RRXII | 99% | 100.00% | WP_011982650.1 |
| Brucella/Ochrobactrum group | 99% | 100.00% | WP_029928117.1 |
| Brucella | 99% | 99.55% | WP_061346708.1 |
| Brucella anthropi | 99% | 99.55% | WP_194531704.1 |
| Brucella tritici | 99% | 98.65% | WP_151557009.1 |
| Brucella/Ochrobactrum group | 99% | 97.76% | WP_109986726.1 |

**Supplementary Figure 5. Protein BLAST alignments of erythritol catabolism-associated genes.**

Top ten protein alignments of every erythritol catabolism-associated genes (eryA/B/C/D/E/F/G/H/I/R) in *Ochrobactrum* spp..

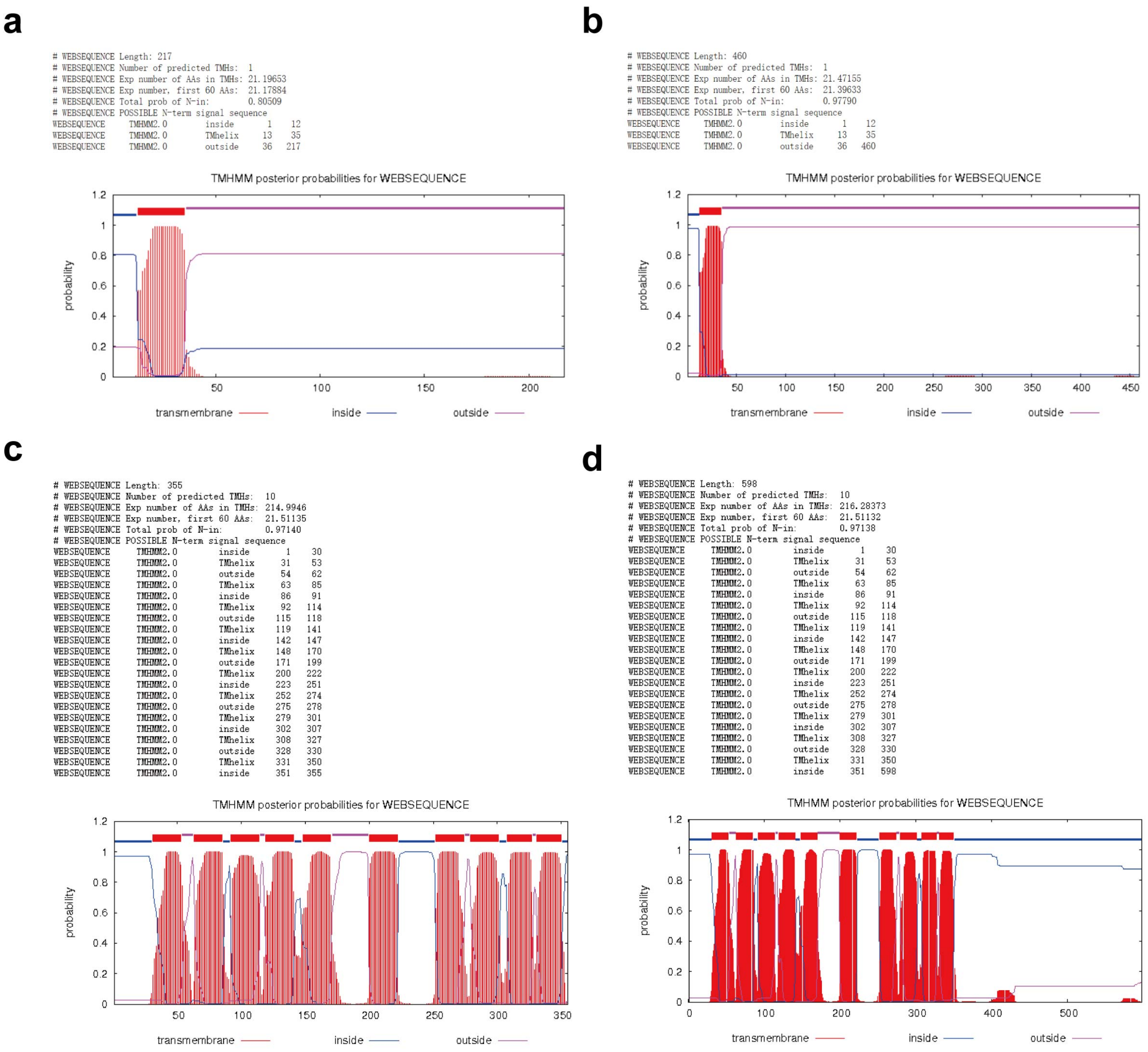

**Supplementary Figure 6. Prediction of eryE and eryG transmembrane topology by TMHMM - 2.0[6].**

(**a**) eryE transmembrane topology prediction indicates that it is a single-pass membrane protein.

(**b**) eryE-sfGFP fusion protein prediction indicates the same topology as eryE.

(**c**) eryG transmembrane topology prediction indicates that it is a ten times transmembrane protein.

(**d**) eryG-sfGFP fusion protein prediction indicates the same topology as eryG.

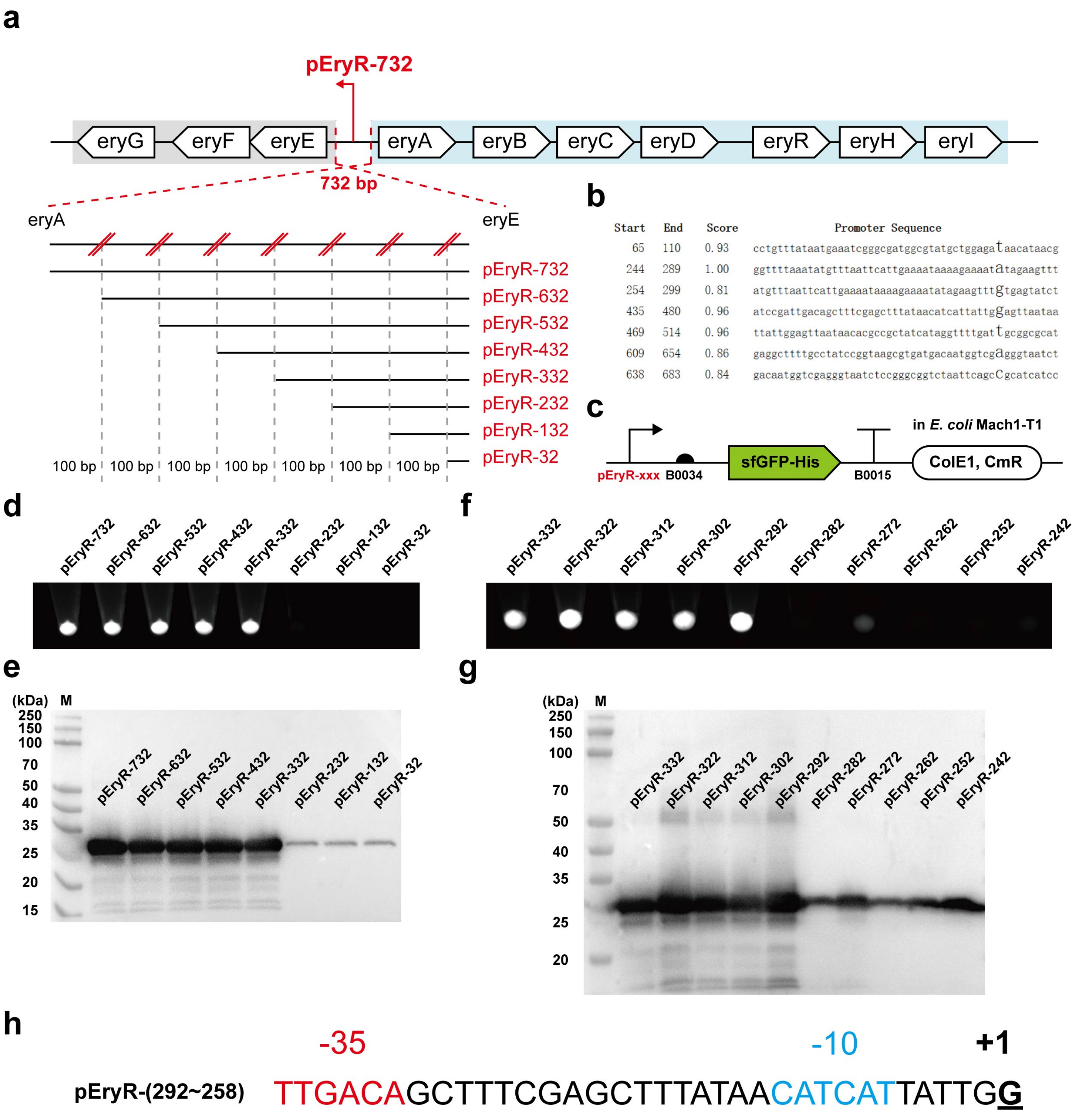

**Supplementary Figure 7. pEryR characterization.**

(**a**) Schematic characterization process of pEryR. The 732 bp DNA gap between eryA and eryE was initially acquired as pEryR-732, and then 100 bp-gradient truncated pEryR variants were constructed into B0034-sfGFP (6xHis)-based reporter plasmids (pFB239 to pFB243, pFB253 to pFB255).

(**b**) Promoter prediction result of pEryR-732 by "BDGP: Neural Network Promoter Prediction"[7].

(**c**) Schematic B0034-sfGFP (6xHis)-based reporter plasmid design. All the following characterization is in *E. coli* strain Mach1-T1.

(**d**) *E. coli* Mach1-T1 pellets were collected from 1.5 mL overnight LB culture (37°C, 250 rpm for 16 h). Brighter pellets indicated more sfGFP expression. The result showed that the core promoter region might be located between pEryR-332 and pEryR-232. Fluorescence images were generated by UVP ChemStudio (analytikjena).

(**e**) Western-Blot analysis of sfGFP (6xHis) relative expression level, the samples were the same as (**d**), the cell pellets were resuspended (equal volume) with 1x phosphate-buffered saline (pH 7.4) and lysed by sonication, the total cell lysate was analyzed.
(**f**) 10 bp-gradient truncated pEryR variants characterization. The process was the same as (**d**), and the result showed that the core promoter region might be located between pEryR-292 and pEryR-282.
(**g**) Western-Blot analysis of sfGFP (6xHis) relative expression level, the samples were the same as (**f**).
(**h**) Schematic of pEryR core region sequence after the truncated characterization. -35, -10, and +1 regions were shown as red, blue, and underlined bold black, respectively.

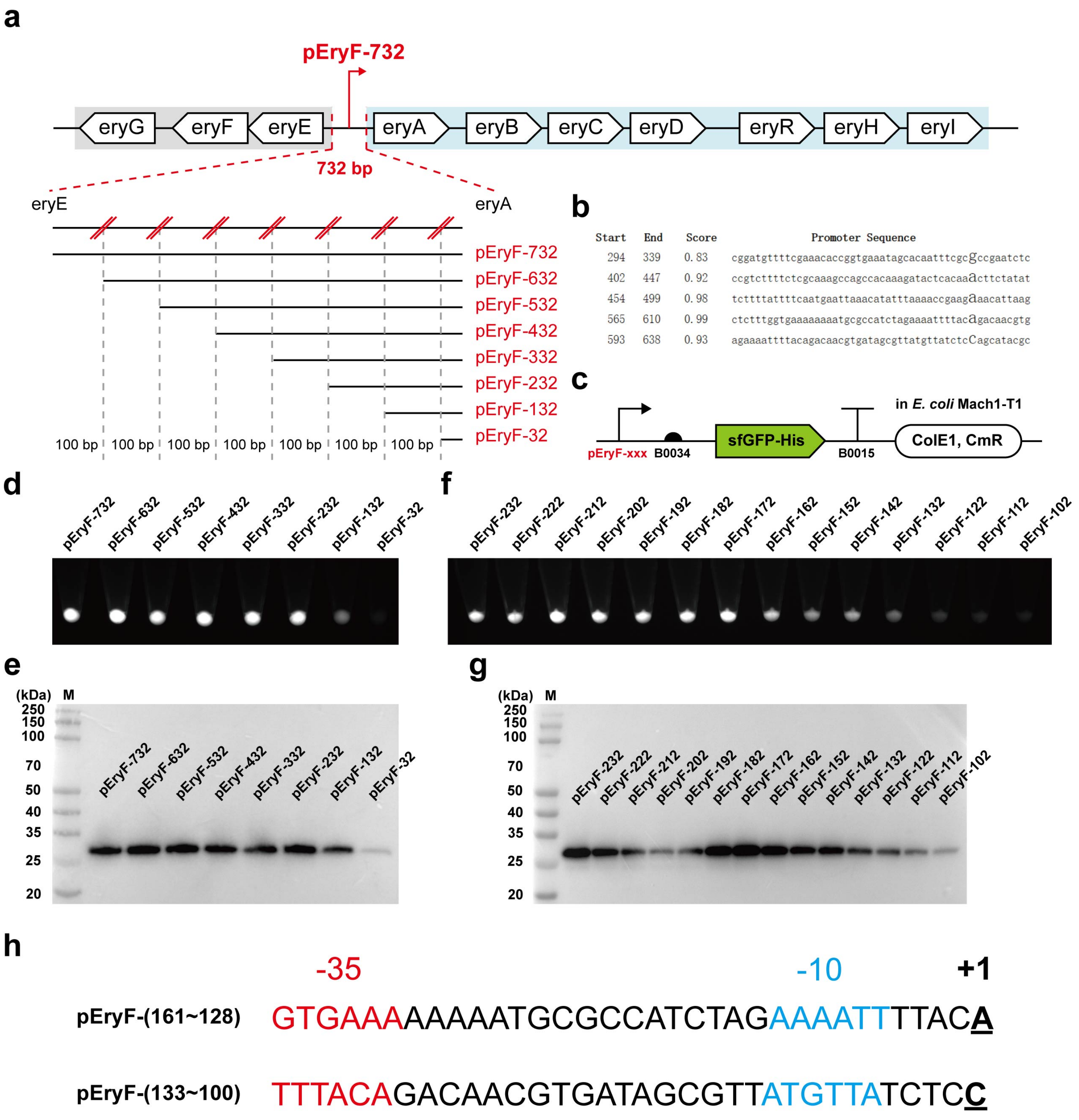

**Supplementary Figure 8. pEryF characterization.**

(**a**) Schematic characterization process of pEryF. The 732 bp DNA gap between eryE and eryA was initially acquired as pEryF-732, and then 100 bp-gradient truncated pEryF variants were constructed into B0034-sfGFP (6xHis)-based reporter plasmids (pFB186 to pFB191, pFB210, and pFB292).

(**b**) Promoter prediction result of pEryF-732 by "BDGP: Neural Network Promoter Prediction"[7].

(**c**) Schematic B0034-sfGFP (6xHis)-based reporter plasmid design. All the following characterization is in *E. coli* strain Mach1-T1.

(**d**) *E. coli* Mach1-T1 pellets were collected from 1.5 mL overnight LB culture (37°C, 250 rpm for 16 h). Brighter pellets indicated more sfGFP expression. The result showed that the core promoter region might be located between pEryF-232 and pEryF-32. Fluorescence images were generated by UVP ChemStudio (analytikjena).
(**e**) Western-Blot analysis of sfGFP (6xHis) relative expression level, the samples were the same as (**d**), the cell pellets were resuspended (equal volume) with 1x phosphate-buffered saline (pH 7.4) and lysed by sonication, the total cell lysate was analyzed.
(**f**) 10 bp-gradient truncated pEryF variants characterization. The process was the same as (**d**), the result showed that the core promoter region might be located between pEryF-172 and pEryF-102.
(**g**) Western-Blot analysis of sfGFP (6xHis) relative expression level, the samples were the same as (**f**).
(**h**) Schematic of two pEryF core region sequences after the truncated characterization. -35, -10, and +1 regions were shown as red, blue, and underlined bold black, respectively.

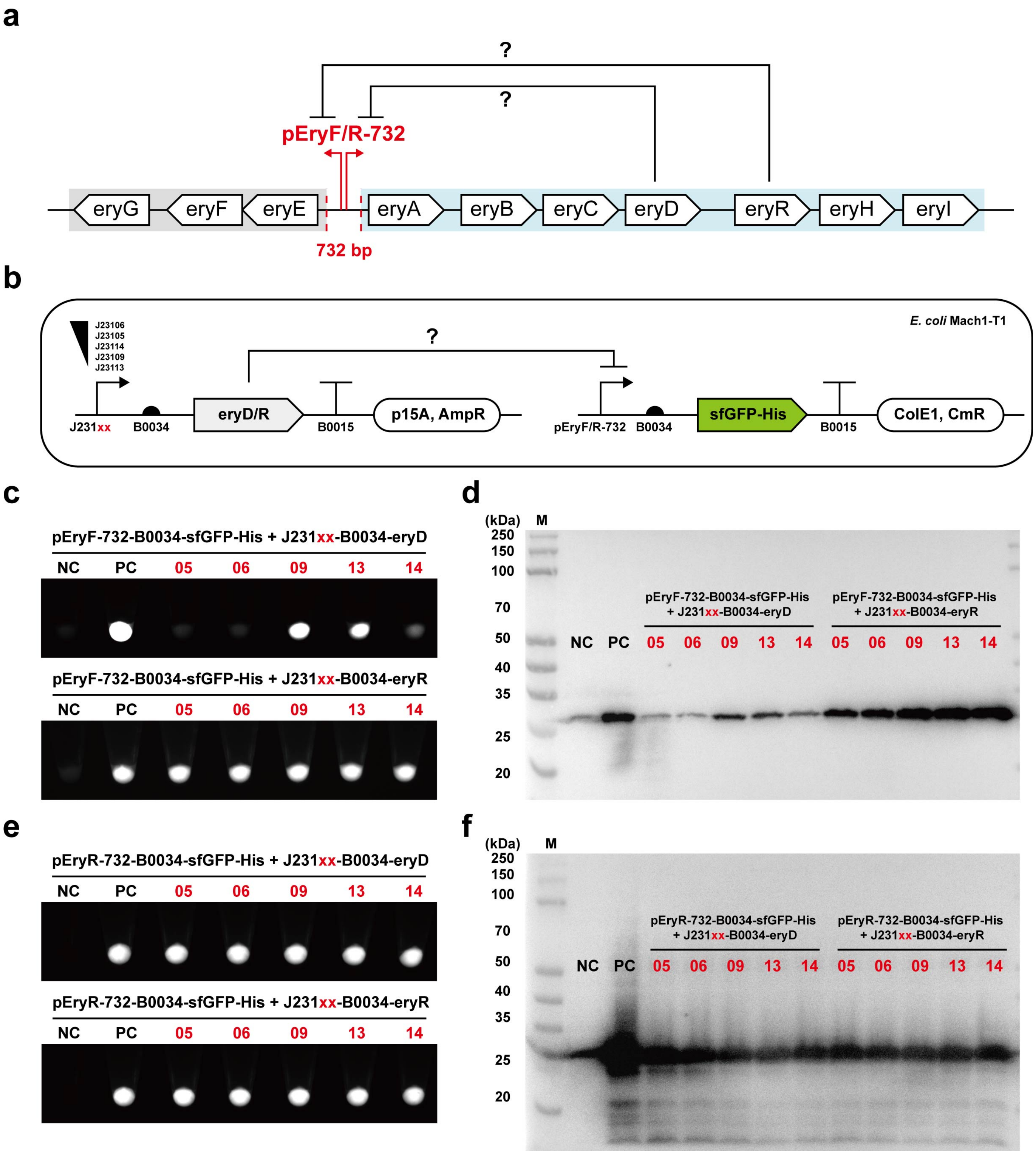

**Supplementary Figure 9. eryD and eryR characterization.**

(**a**) Schematic characterization process of eryD and eryR. We assumed that eryD/eryR might regulate erythritol cluster, and the long DNA gap between eryE and eryA might locate eryD/eryR binding sites in both forward and reverse directions.

(**b**) Schematic compatible plasmids design in *E. coli* Mach1-T1. Strength-gradient promoters were organized with RBS B0034, eryD/eryR coding sequences, and terminator B0015 (pFB260 to pFB269). The other two compatible plasmids (pFB186 and pFB239) were used as reporters.

(**c**) *E. coli* Mach1-T1 pellets were collected from 1.5 mL overnight LB culture (37°C, 250 rpm for 16 h). Brighter pellets indicated more sfGFP expression. The result showed that there was a putative eryD DNA-binding site on pEryF-732, but eryR not. Furthermore, a higher expression level of eryD caused lower sfGFP expression. “NC” represents *E. coli* Mach1-T1 without plasmids, and “PC” represents *E. coli* Mach1-T1 with plasmid pFB186. Fluorescence images were generated by UVP ChemStudio (analytikjena).
(**d**) Western-Blot analysis of sfGFP (6xHis) relative expression level, the samples were the same as (**c**), the cell pellets were resuspended (equal volume) with 1x phosphate-buffered saline (pH 7.4) and lysed by sonication, the total cell lysate was analyzed.
(**e**) *E. coli* Mach1-T1 pellets were collected from 1.5 mL overnight LB culture (37°C, 250 rpm for 16 h). Brighter pellets indicated more sfGFP expression. The result showed that there were not any eryD or eryR DNA-binding sites on pEryR-732. “NC” represents *E. coli* Mach1-T1 without plasmids, and “PC” represents *E. coli* Mach1-T1 with plasmid pFB239. Fluorescence images were generated by UVP ChemStudio (analytikjena).
(**f**) Western-Blot analysis of sfGFP (6xHis) relative expression level, the samples were the same as (**e**), the cell pellets were resuspended (equal volume) with 1x phosphate-buffered saline (pH 7.4) and lysed by sonication, the total cell lysate was analyzed.

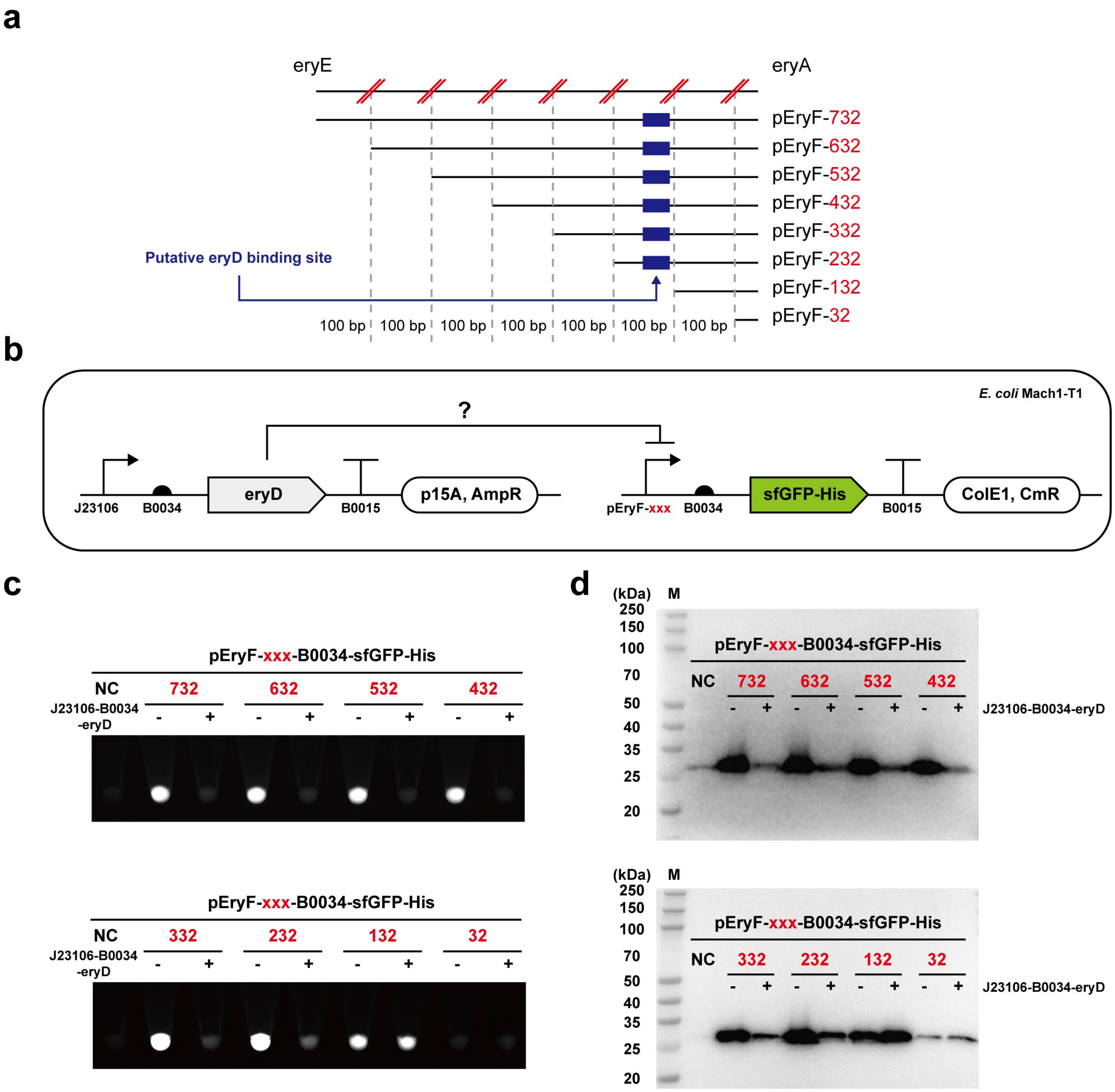

**Supplementary Figure 10. The first step for characterization of eryD binding site prefix.**

(**a**, **b**) Schematic characterization process of putative eryD binding site (eryO).

(**c**) *E. coli* Mach1-T1 pellets were collected from 1.5 mL overnight LB culture (37°C, 250 rpm for 16 h). Brighter pellets indicated more sfGFP expression. “NC” represents *E. coli* Mach1-T1 without plasmids, and the “-/+” means the *E. coli* strains without or with plasmid pFB261. The result showed that the eryO region might be located between pEryF-232 and pEryF-132. Fluorescence images were generated by UVP ChemStudio (analytikjena).

(**d**) Western-Blot analysis of sfGFP (6xHis) relative expression level, the samples were the same as (**c**), the cell pellets were resuspended (equal volume) with 1x phosphate-buffered saline (pH 7.4) and lysed by sonication, the total cell lysate was analyzed.

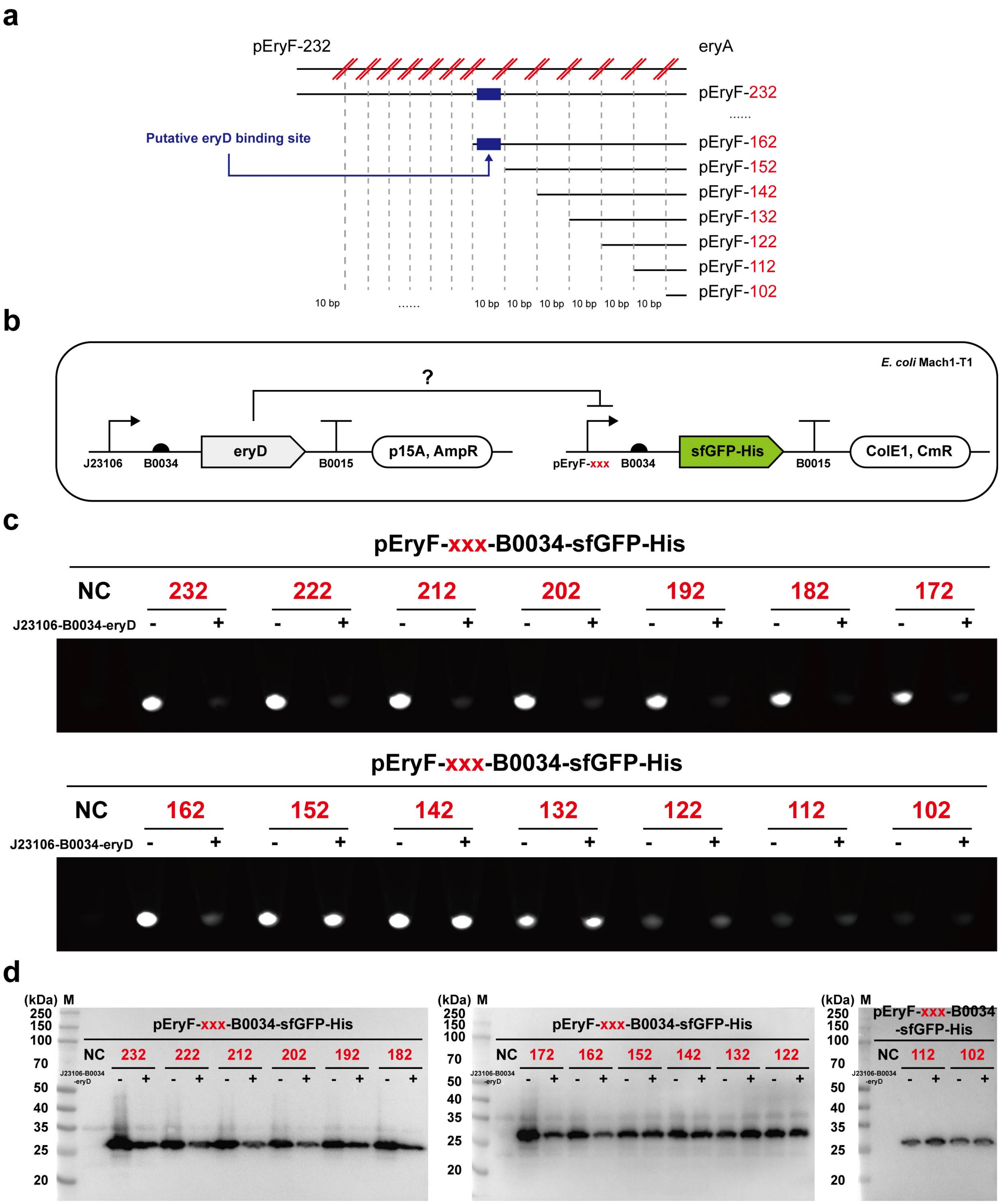

**Supplementary Figure 11. The second step for characterization of eryD binding site prefix.**

(**a**, **b**) Schematic characterization process of putative eryD binding site (eryO).

(**c**) *E. coli* Mach1-T1 pellets were collected from 1.5 mL overnight LB culture (37°C, 250 rpm for 16 h). Brighter pellets indicated more sfGFP expression. "NC" represents *E. coli* Mach1-T1 without plasmids, and the "-/+" means the *E. coli* strains without or with plasmid pFB261. The result showed that the prefix of eryO region might be located between pEryF-162 and pEryF-152. Fluorescence images were generated by UVP ChemStudio (analytikjena).

(**d**) Western-Blot analysis of sfGFP (6xHis) relative expression level, the samples were the same as (**c**), the cell pellets were resuspended (equal volume) with 1x phosphate-buffered saline (pH 7.4) and lysed by sonication, the total cell lysate was analyzed.

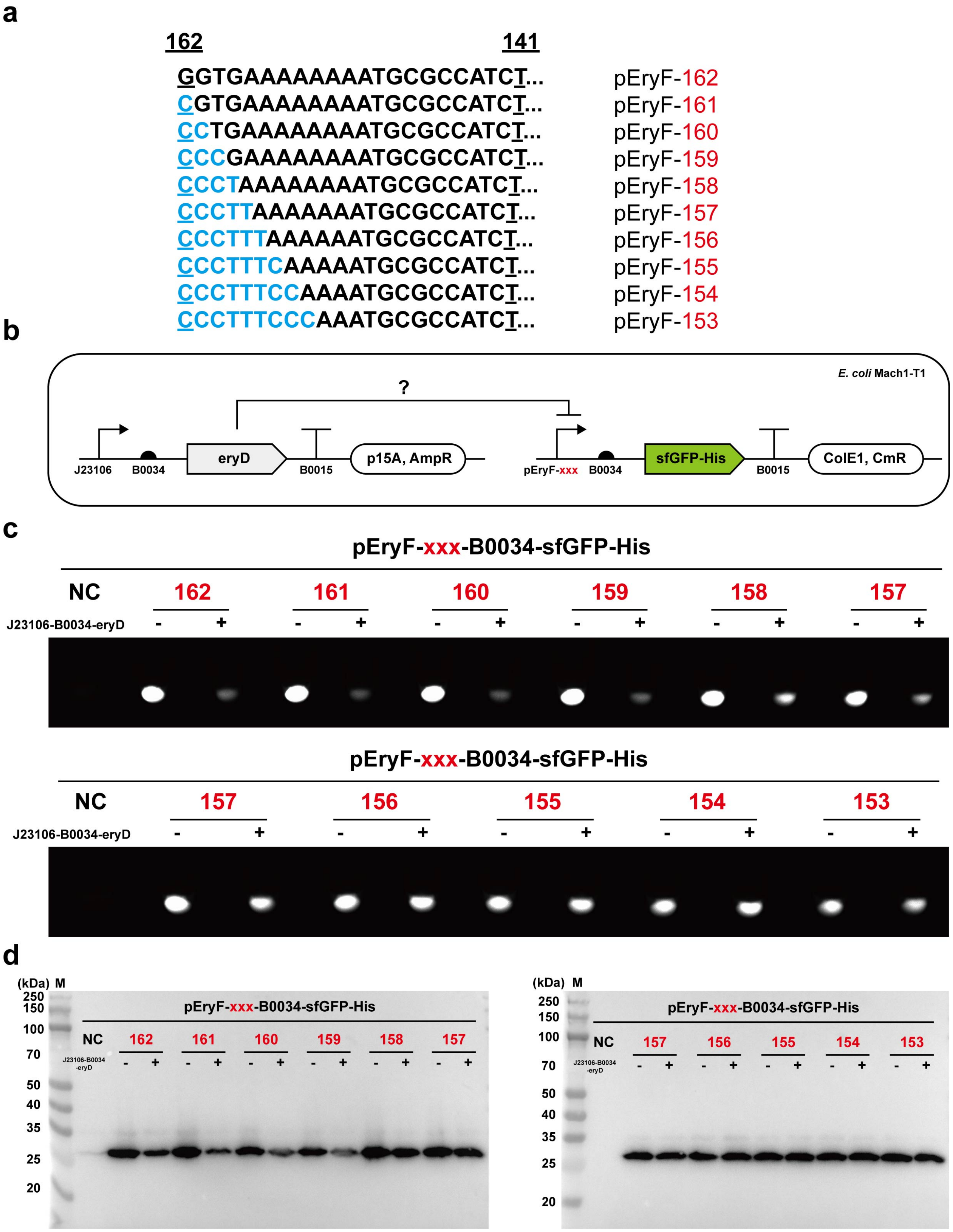
a
162
141
GGTGAAAAAAAATGCGCCATCT...
CGTGAAAAAAAATGCGCCATCT...
CCTGAAAAAAAATGCGCCATCT...
CCCGAAAAAAAATGCGCCATCT...
CCCTAAAAAAAATGCGCCATCT...
CCCTTAAAAAAATGCGCCATCT...
CCCTTTAAAAAATGCGCCATCT...
CCCTTTCAAAAATGCGCCATCT...
CCCTTTCCAAAATGCGCCATCT...
CCCTTTCCCAAATGCGCCATCT...
pEryF-162
pEryF-161
pEryF-160
pEryF-159
pEryF-158
pEryF-157
pEryF-156
pEryF-155
pEryF-154
pEryF-153
b
E. coli Mach1-T1
?
J23106
B0034
eryD
B0015
p15A, AmpR
pEryF-xxx
B0034
sfGFP-His
B0015
ColE1, CmR
c
pEryF-xxx-B0034-sfGFP-His
NC
162
161
160
159
158
157
J23106-B0034-eryD
- + - + - + - + - + - +
pEryF-xxx-B0034-sfGFP-His
NC
157
156
155
154
153
J23106-B0034-eryD
- + - + - + - + - +
d
(kDa) M
250
150
100
70
50
40
35
25
20
pEryF-xxx-B0034-sfGFP-His
NC
162
161
160
159
158
157
J23106-B0034-eryD
- + - + - + - + - + - +
(kDa) M
250
150
100
70
50
40
35
25
20
pEryF-xxx-B0034-sfGFP-His
NC
157
156
155
154
153
J23106-B0034-eryD
- + - + - + - + - +

**Supplementary Figure 12. The third step for characterization of eryD binding site prefix.**
(**a**, **b**) Schematic characterization process of putative eryD binding site (eryO).
(**c**) *E. coli* Mach1-T1 pellets were collected from 1.5 mL overnight LB culture (37°C, 250 rpm for 16 h). Brighter pellets indicated more sfGFP expression. “NC” represents *E. coli* Mach1-T1 without plasmids, and the “-/+” means the *E. coli* strains without or with plasmid pFB261. The result showed that the prefix of eryO was located in pEryF-159 (5’-GAAAAAAAAT…-3’). Fluorescence images were generated by UVP ChemStudio (analytikjena).
(**d**) Western-Blot analysis of sfGFP (6xHis) relative expression level, the samples were the same as (**c**), the cell pellets were resuspended (equal volume) with 1x phosphate-buffered saline (pH 7.4) and lysed by sonication, the total cell lysate was analyzed.

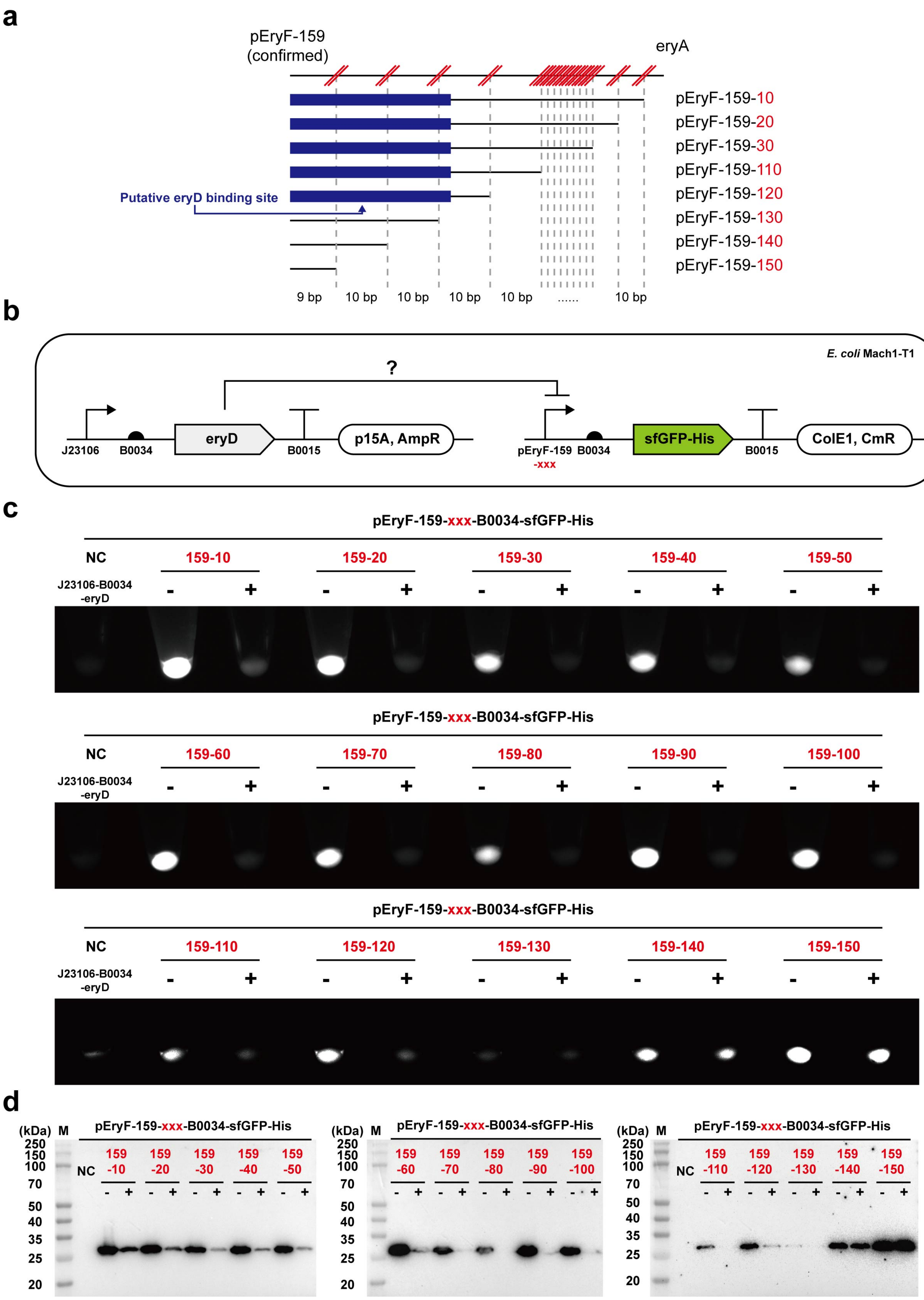

a
pEryF-159
(confirmed)
eryA
pEryF-159-10
pEryF-159-20
pEryF-159-30
pEryF-159-110
pEryF-159-120
pEryF-159-130
pEryF-159-140
pEryF-159-150
Putative eryD binding site
9 bp
10 bp
10 bp
10 bp
10 bp
......
10 bp
b
E. coli Mach1-T1
?
J23106
B0034
eryD
B0015
p15A, AmpR
pEryF-159
-xxx
B0034
sfGFP-His
B0015
ColE1, CmR
c
pEryF-159-xxx-B0034-sfGFP-His
NC
159-10
159-20
159-30
159-40
159-50
J23106-B0034
-eryD
pEryF-159-xxx-B0034-sfGFP-His
NC
159-60
159-70
159-80
159-90
159-100
J23106-B0034
-eryD
pEryF-159-xxx-B0034-sfGFP-His
NC
159-110
159-120
159-130
159-140
159-150
J23106-B0034
-eryD
d
(kDa)
M
pEryF-159-xxx-B0034-sfGFP-His
250
150
100
70
50
40
35
25
20
NC
159 -10
159 -20
159 -30
159 -40
159 -50
159 -60
159 -70
159 -80
159 -90
159 -100
159 -110
159 -120
159 -130
159 -140
159 -150

**Supplementary Figure 13. The first step for characterization of eryD binding site suffix.**
(**a**, **b**) Schematic characterization process of putative eryD binding site (eryO).
(**c**) *E. coli* Mach1-T1 pellets were collected from 1.5 mL overnight LB culture (37°C, 250 rpm for 16 h). Brighter pellets indicated more sfGFP expression. “NC” represents *E. coli* Mach1-T1 without plasmids, and the “-/+” means the *E. coli* strain without or with plasmid pFB261. The result showed that the suffix of eryO was located between pEryF-159-120 and pEryF-159-140. Fluorescence images were generated by UVP ChemStudio (analytikjena).
(**d**) Western-Blot analysis of sfGFP (6xHis) relative expression level, the samples were the same as (**c**), the cell pellets were resuspended (equal volume) with 1x phosphate-buffered saline (pH 7.4) and lysed by sonication, the total cell lysate was analyzed.

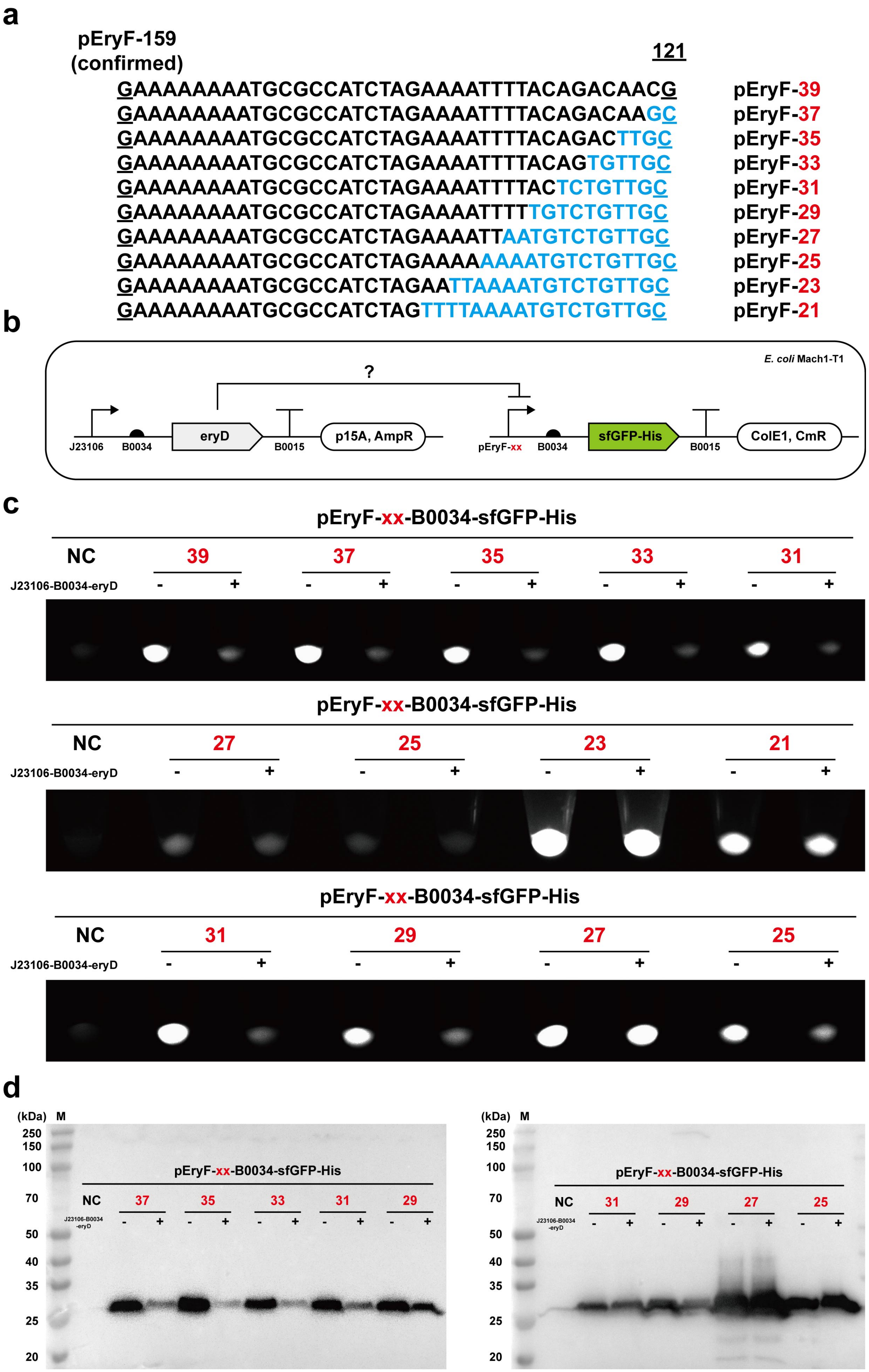

a
pEryF-159
(confirmed)
121
GAAAAAAAATGCGCCATCTAGAAAATTTTACAGACAACG pEryF-39
GAAAAAAAATGCGCCATCTAGAAAATTTTACAGACAAGC pEryF-37
GAAAAAAAATGCGCCATCTAGAAAATTTTACAGACTTGC pEryF-35
GAAAAAAAATGCGCCATCTAGAAAATTTTACAGTGTTGC pEryF-33
GAAAAAAAATGCGCCATCTAGAAAATTTTACTCTGTTGC pEryF-31
GAAAAAAAATGCGCCATCTAGAAAATTTTTGTCTGTTGC pEryF-29
GAAAAAAAATGCGCCATCTAGAAAATTAATGTCTGTTGC pEryF-27
GAAAAAAAATGCGCCATCTAGAAAAAAAATGTCTGTTGC pEryF-25
GAAAAAAAATGCGCCATCTAGAATTAAAATGTCTGTTGC pEryF-23
GAAAAAAAATGCGCCATCTAGTTTTAAAATGTCTGTTGC pEryF-21
b
E. coli Mach1-T1
?
J23106
B0034
eryD
B0015
p15A, AmpR
pEryF-xx
B0034
sfGFP-His
B0015
ColE1, CmR
c
pEryF-xx-B0034-sfGFP-His
NC
39
37
35
33
31
J23106-B0034-eryD
- + - + - + - + - +
pEryF-xx-B0034-sfGFP-His
NC
27
25
23
21
J23106-B0034-eryD
- + - + - + - +
pEryF-xx-B0034-sfGFP-His
NC
31
29
27
25
J23106-B0034-eryD
- + - + - + - +
d
(kDa) M
250
150
100
70
50
40
35
25
20
pEryF-xx-B0034-sfGFP-His
NC
37
35
33
31
29
J23106-B0034-eryD
- + - + - + - + - +
(kDa) M
250
150
100
70
50
40
35
25
20
pEryF-xx-B0034-sfGFP-His
NC
31
29
27
25
J23106-B0034-eryD
- + - + - + - +

**Supplementary Figure 14. The second step for characterization of eryD binding site suffix.**

(**a**, **b**) Schematic characterization process of putative eryD binding site (eryO).

(**c**) *E. coli* Mach1-T1 pellets were collected from 1.5 mL overnight LB culture (37°C, 250 rpm for 16 h). Brighter pellets indicated more sfGFP expression. “NC” represents *E. coli* Mach1-T1 without plasmids, and the “-/+” means the *E. coli* strains without or with plasmid pFB261. The result showed that the suffix of eryO was located between pEryF-29 and pEryF-27. Fluorescence images were generated by UVP ChemStudio (analytikjena).

(**d**) Western-Blot analysis of sfGFP (6xHis) relative expression level, the samples were the same as (**c**), the cell pellets were resuspended (equal volume) with 1x phosphate-buffered saline (pH 7.4) and lysed by sonication, the total cell lysate was analyzed.

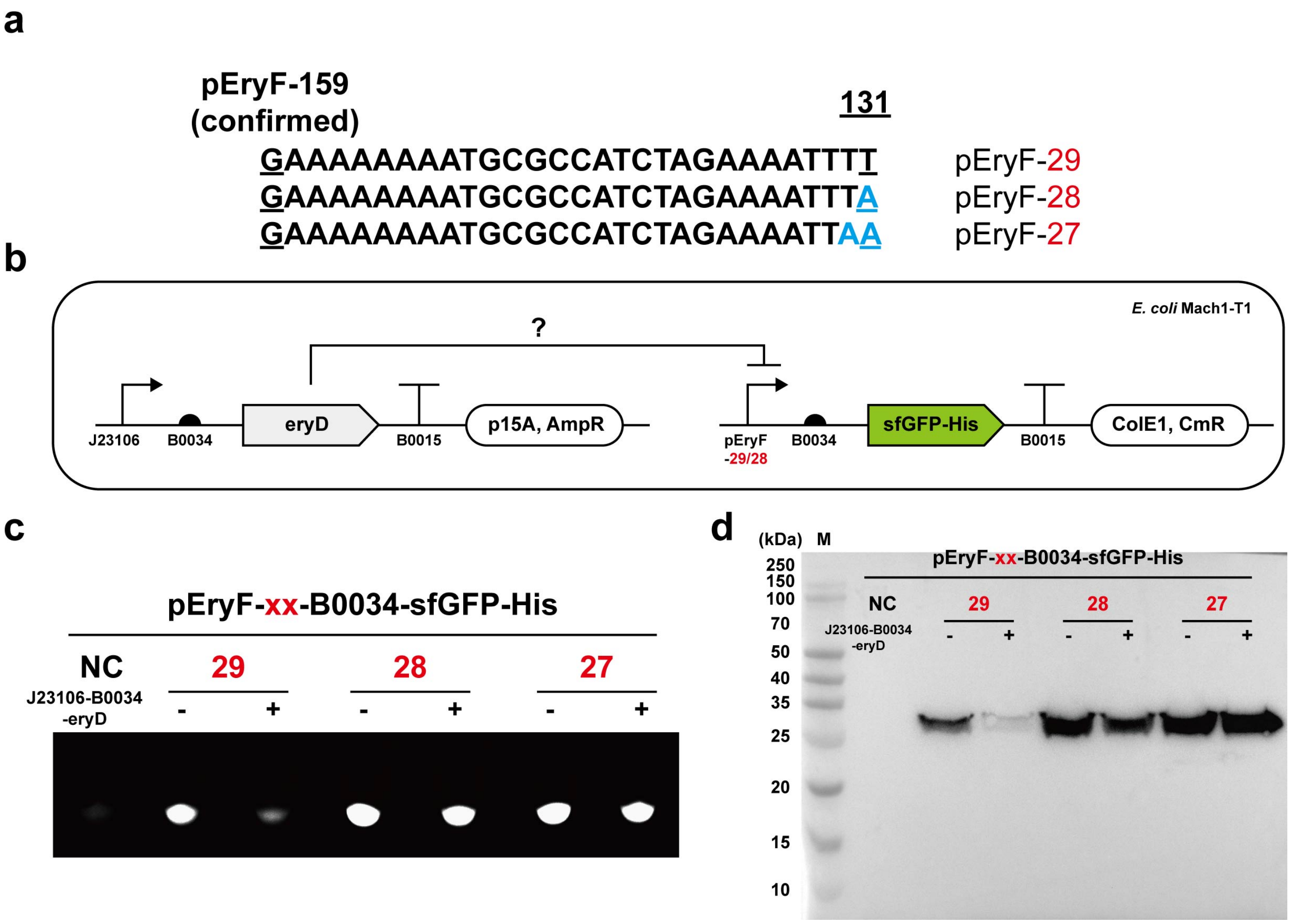

**Supplementary Figure 15. The third step for characterization of eryD binding site suffix.**

(**a**, **b**) Schematic characterization process of putative eryD binding site (eryO).

(**c**) *E. coli* Mach1-T1 pellets were collected from 1.5 mL overnight LB culture (37°C, 250 rpm for 16 h). Brighter pellets indicated more sfGFP expression. “NC” represents *E. coli* Mach1-T1 without plasmids, and the “-/+” means the *E. coli* strains without or with plasmid pFB261. The result showed that the suffix of eryO was located in pEryF-29 (5’-…AAAATTTT-3’), and the characterized eryO sequence was 5’-GAAAAAAAATGCGCCATCTAGAAAATTTT-3’. Fluorescence images were generated by UVP ChemStudio (analytikjena).

(**d**) Western-Blot analysis of sfGFP (6xHis) relative expression level, the samples were the same as (**c**), the cell pellets were resuspended (equal volume) with 1x phosphate-buffered saline (pH 7.4) and lysed by sonication, the total cell lysate was analyzed.

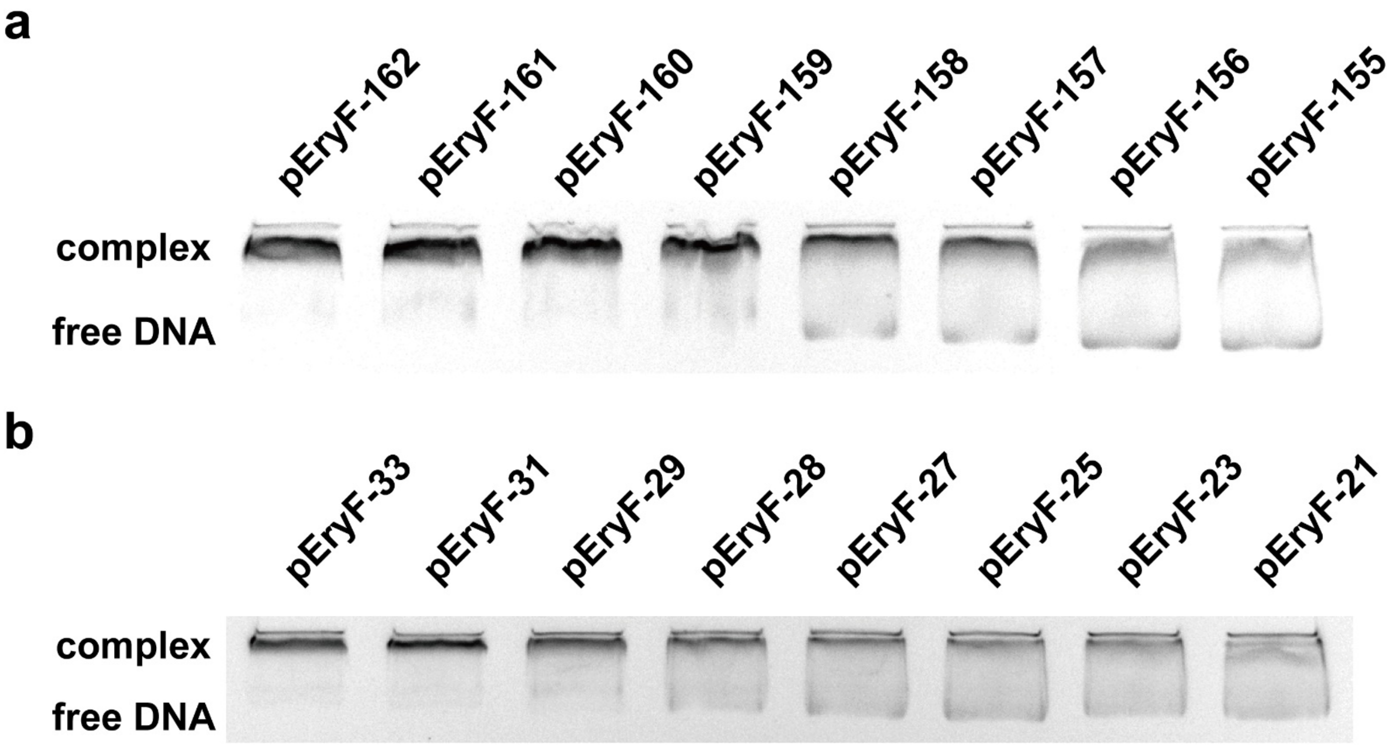

**Supplementary Figure 16. EMSA analysis of eryD-eryO interaction.**

(**a**) EMSA analysis demonstrates that the prefix of eryO is located in pEryF-159 (5'-GAAAAAAAAT…-3') as the same as *in vivo* experiment results. Each 20 µL reaction consists of 100 nM 5'FAM-labelled pEryF variants and 400 nM purified eryD.

(**b**) EMSA analysis demonstrates that the suffix of eryO is located in pEryF-29 (5'-…AAAATTTT-3') as the same as *in vivo* experiment results. Each 20 µL reaction consists of 100 nM 5'FAM-labelled pEryF variants and 400 nM purified eryD.

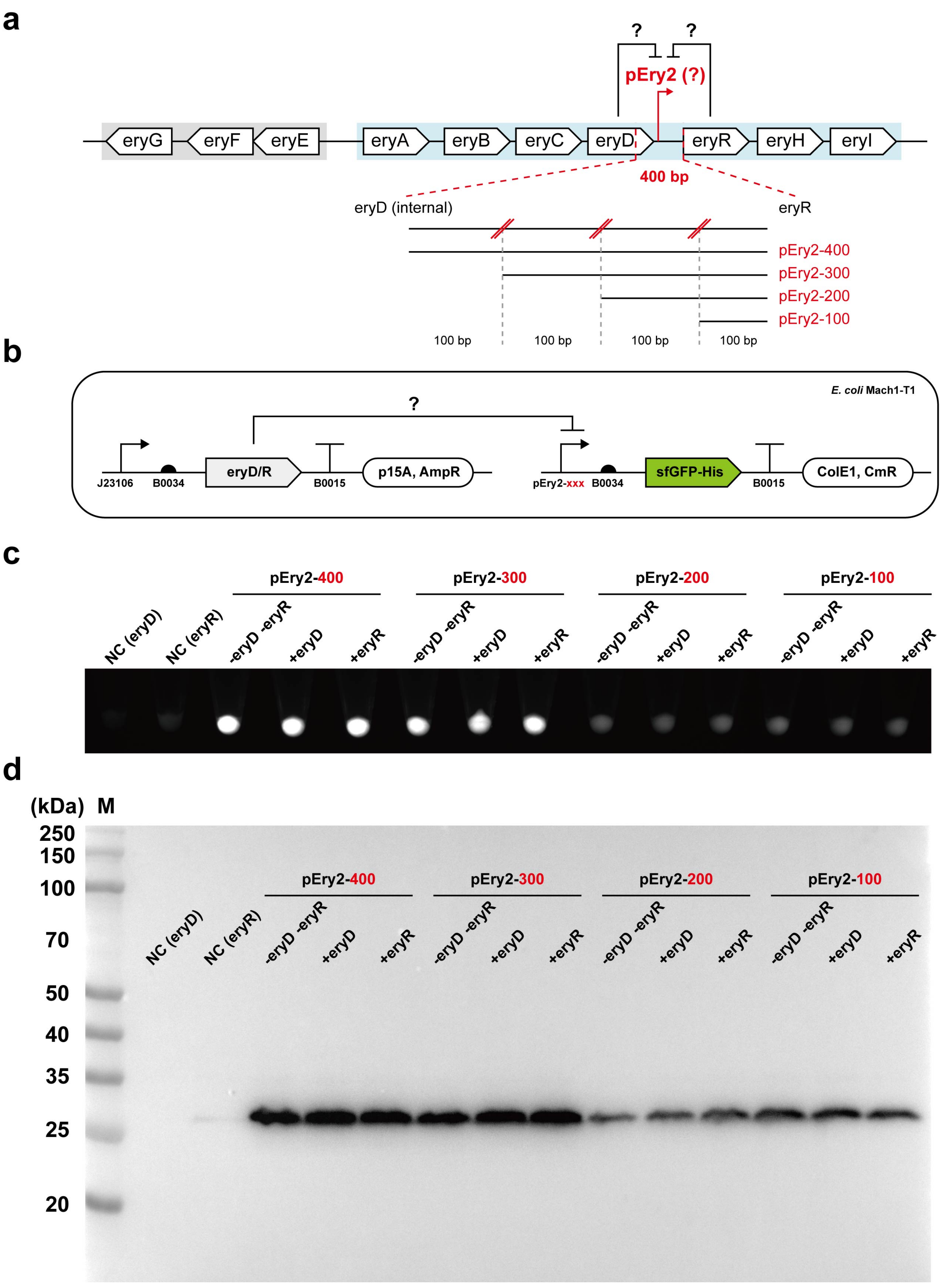
a
?
?
pEry2 (?)
eryG
eryF
eryE
eryA
eryB
eryC
eryD
eryR
eryH
eryI
400 bp
eryD (internal)
eryR
pEry2-400
pEry2-300
pEry2-200
pEry2-100
100 bp
100 bp
100 bp
100 bp
b
E. coli Mach1-T1
?
J23106
B0034
eryD/R
B0015
p15A, AmpR
pEry2-xxx
B0034
sfGFP-His
B0015
ColE1, CmR
c
pEry2-400
pEry2-300
pEry2-200
pEry2-100
NC (eryD)
NC (eryR)
-eryD -eryR
+eryD
+eryR
-eryD -eryR
+eryD
+eryR
-eryD -eryR
+eryD
+eryR
-eryD -eryR
+eryD
+eryR
d
(kDa)
M
250
150
100
70
50
40
35
25
20
pEry2-400
pEry2-300
pEry2-200
pEry2-100
NC (eryD)
NC (eryR)
-eryD -eryR
+eryD
+eryR
-eryD -eryR
+eryD
+eryR
-eryD -eryR
+eryD
+eryR
-eryD -eryR
+eryD
+eryR

**Supplementary Figure 17. pEry2 characterization.**
(**a**) Schematic characterization process of whether there were eryD/eryR binding sites in the region of pEry2 variants. We assumed that eryD/eryR might regulate the erythritol cluster, and the long DNA gap between eryD and eryR might locate eryD/eryR binding sites in both forward and reverse directions.
(**b**) Schematic compatible plasmids design in *E. coli* Mach1-T1. Plasmids that could strongly express eryD/eryR (pFB261 and pFB266) co-existed with the reporter plasmids, which contained 100 bp truncated pEry2 variants (pFB256 to pFB259).
(**c**) *E. coli* Mach1-T1 pellets were collected from 1.5 mL overnight LB culture (37°C, 250 rpm for 16 h). Brighter pellets indicated more sfGFP expression. Two "NC" are the same and represent *E. coli* Mach1-T1 without plasmids, and the "-/+" means the *E. coli* strains without or with plasmid pFB261 or pFB266. The result showed that there were not any eryD or eryR DNA binding sites in pEry2 region. Fluorescence images were generated by UVP ChemStudio (analytikjena).
(**d**) Western-Blot analysis of sfGFP (6xHis) relative expression level, the samples were the same as (**c**), the cell pellets were resuspended (equal volume) with 1x phosphate-buffered saline (pH 7.4) and lysed by sonication, the total cell lysate was analyzed.

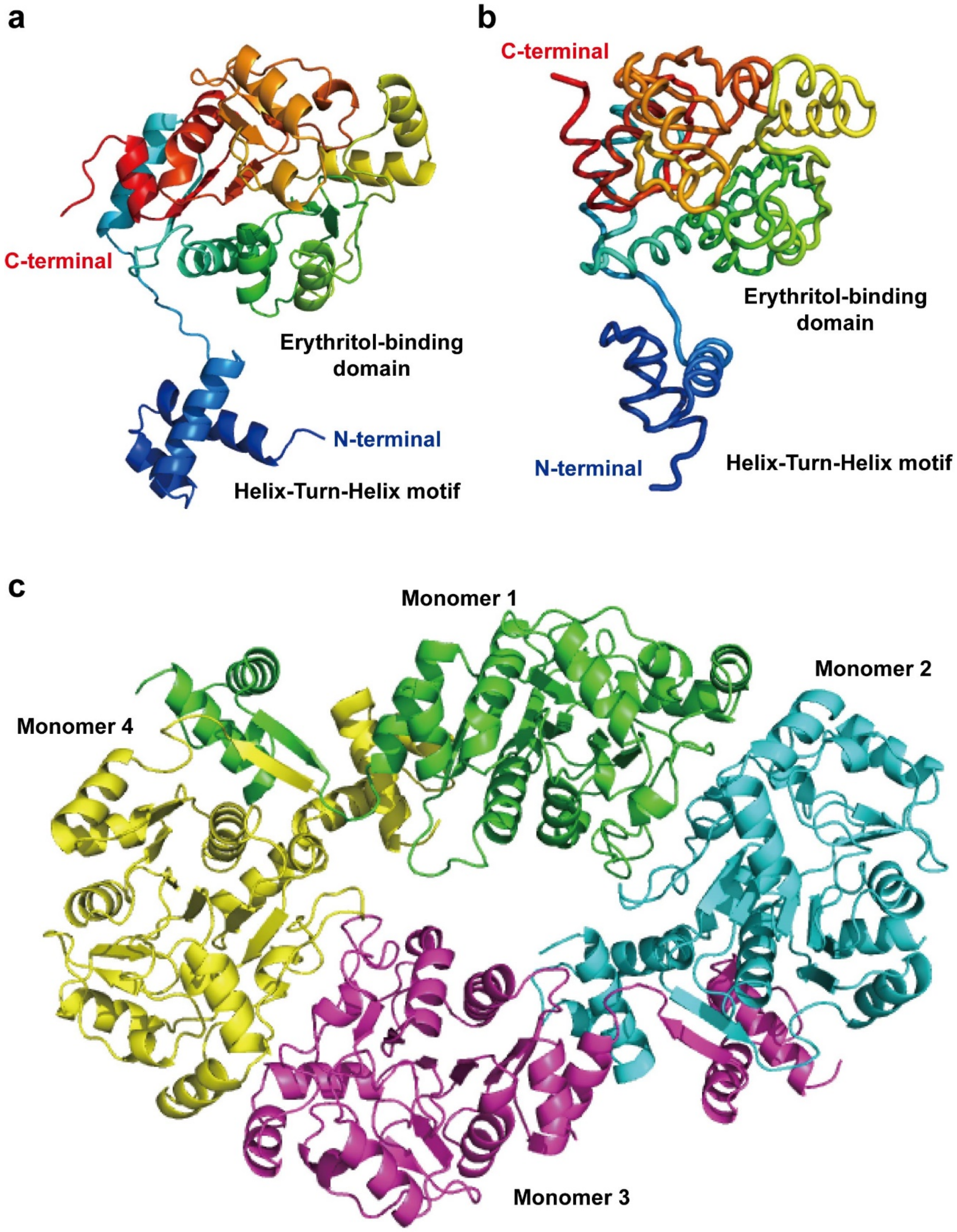

**Supplementary Figure 18. Structure models of eryD monomer and homotetramer.**
(**a**) AlphaFold[8,9] structure prediction of eryD monomer (Uniprot ID: Q2YIQ4). This predicted eryD is from *Brucella abortus* (strain 2308), and the amino acid sequence is 100% the same as the eryD we characterized in this work.
(**b**) Robetta[10] structure prediction of eryD that we characterized in this work.
(**c**) SWISS-MODEL[11,12] homotetramer structure prediction of eryD that we characterized in this work. This predicted model is built up with the PDB template 4l5i.1.A (Transcriptional regulator LsrR).

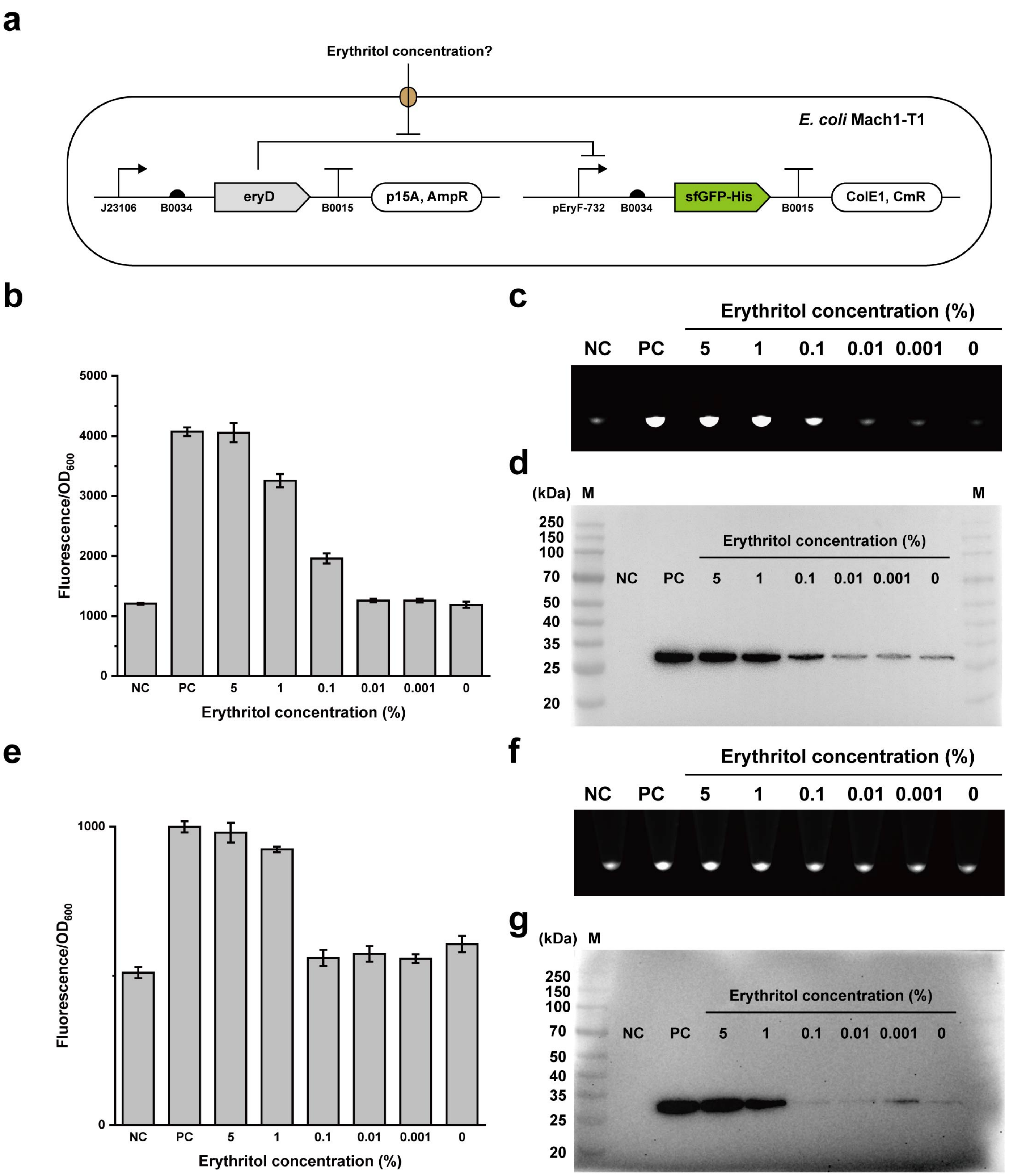

**Supplementary Figure 19. Erythritol induction of pEryF-732 at 30°C and 16°C.**
(**a**) Schematic diagram of used strain and plasmids. pFB186 and pFB261 co-existed in *E. coli* Mach1-T1.
(**b**) The mentioned strain in (**a**) was induced by different concentrations of erythritol during overnight incubation in LB medium at 30°C, 250 rpm for 16 h. “NC” represents *E. coli* Mach1-T1 without plasmids, “PC” represents *E. coli*

Mach1-T1 with plasmid pFB186 and without pFB261. All measurements were performed with three biological replicates.

(**c**) The collected *E. coli* Mach1-T1 pellets from (**b**), brighter pellets indicated more sfGFP expression.

(**d**) Western-Blot analysis of sfGFP (6xHis) relative expression level, the samples were the same as (**b**) and (**c**), the cell pellets were resuspended (equal volume) with 1x phosphate-buffered saline (pH 7.4) and lysed by sonication, the total cell lysate was analyzed.

(**e**) The mentioned strain in (**a**) was induced by different concentrations of erythritol during overnight incubation in LB medium at 16°C, 250 rpm for 24 h. “NC” represents *E. coli* Mach1-T1 without plasmids, “PC” represents *E. coli* Mach1-T1 with plasmid pFB186 and without pFB261. All measurements were performed with three biological replicates.

(**f**) The collected *E. coli* Mach1-T1 pellets from (**e**), brighter pellets indicated more sfGFP expression.

(**g**) Western-Blot analysis of sfGFP (6xHis) relative expression level, the samples were the same as (**e**) and (**f**), the cell pellets were resuspended (equal volume) with 1x phosphate-buffered saline (pH 7.4) and lysed by sonication, the total cell lysate was analyzed.

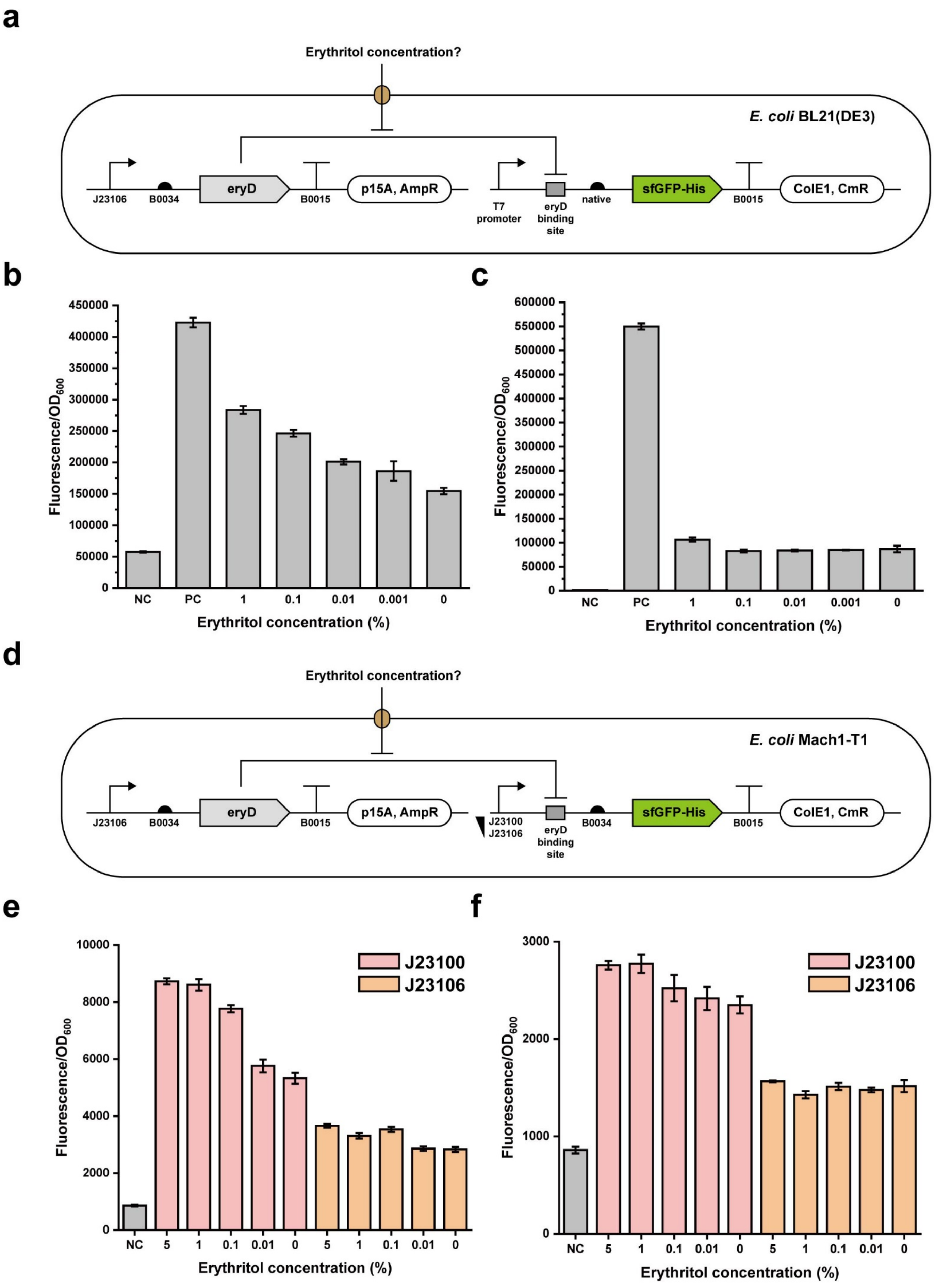
a
Erythritol concentration?
E. coli BL21(DE3)
J23106
B0034
eryD
B0015
p15A, AmpR
T7 promoter
eryD binding site
native
sfGFP-His
B0015
ColE1, CmR
b
Fluorescence/OD600
0
50000
100000
150000
200000
250000
300000
350000
400000
450000
NC
PC
1
0.1
0.01
0.001
0
Erythritol concentration (%)
c
Fluorescence/OD600
0
50000
100000
150000
200000
250000
300000
350000
400000
450000
500000
550000
600000
NC
PC
1
0.1
0.01
0.001
0
Erythritol concentration (%)
d
Erythritol concentration?
E. coli Mach1-T1
J23106
B0034
eryD
B0015
p15A, AmpR
J23100
J23106
eryD binding site
B0034
sfGFP-His
B0015
ColE1, CmR
e
J23100
J23106
Fluorescence/OD600
0
2000
4000
6000
8000
10000
NC
5
1
0.1
0.01
0
5
1
0.1
0.01
0
Erythritol concentration (%)
f
J23100
J23106
Fluorescence/OD600
0
1000
2000
3000
NC
5
1
0.1
0.01
0
5
1
0.1
0.01
0
Erythritol concentration (%)

**Supplementary Figure 20. Erythritol induction of synthetic operons at 30°C and 16°C.**

(**a**) Schematic diagram of used strain and plasmids for pT7-eryO operon characterization. pFB276 and pFB261 co-existed in *E. coli* BL21(DE3).

(**b**) The mentioned strain in (**a**) was induced by different concentrations of erythritol during overnight incubation in LB medium (with 0.5 mM IPTG for T7 RNA polymerase induction) at 30°C, 250 rpm for 16 h. “NC” represents the mentioned strain without IPTG and without erythritol induction, “PC” represents *E. coli* BL21(DE3) with plasmid pFB277 (with 0.5 mM IPTG for T7 RNA polymerase induction and without erythritol induction). All measurements were performed with three biological replicates.

(**c**) The mentioned strain in (**a**) was induced by different concentrations of erythritol during overnight incubation in LB medium (with 0.5 mM IPTG for T7 RNA polymerase induction) at 16°C, 250 rpm for 24 h. “NC” represents the mentioned strain without IPTG and without erythritol induction, “PC” represents *E. coli* BL21(DE3) with plasmid pFB277 (with 0.5 mM IPTG for T7 RNA polymerase induction and without erythritol induction). All measurements were performed with three biological replicates.

(**d**) Schematic diagram of used strain and plasmids for J23100-eryO/J23106-eryO operon characterization. pFB270 and pFB261/pFB272 and pFB261 co-existed in *E. coli* Mach1-T1.

(**e**) The mentioned strains in (**d**) were induced by different concentrations of erythritol during overnight incubation in LB medium at 30°C, 250 rpm for 16 h. “NC” represents *E. coli* Mach1-T1 without plasmids. All measurements were performed with three biological replicates.

(**f**) The mentioned strains in (**d**) were induced by different concentrations of erythritol during overnight incubation in LB medium at 16°C, 250 rpm for 24 h. “NC” represents *E. coli* Mach1-T1 without plasmids. All measurements were performed with three biological replicates.

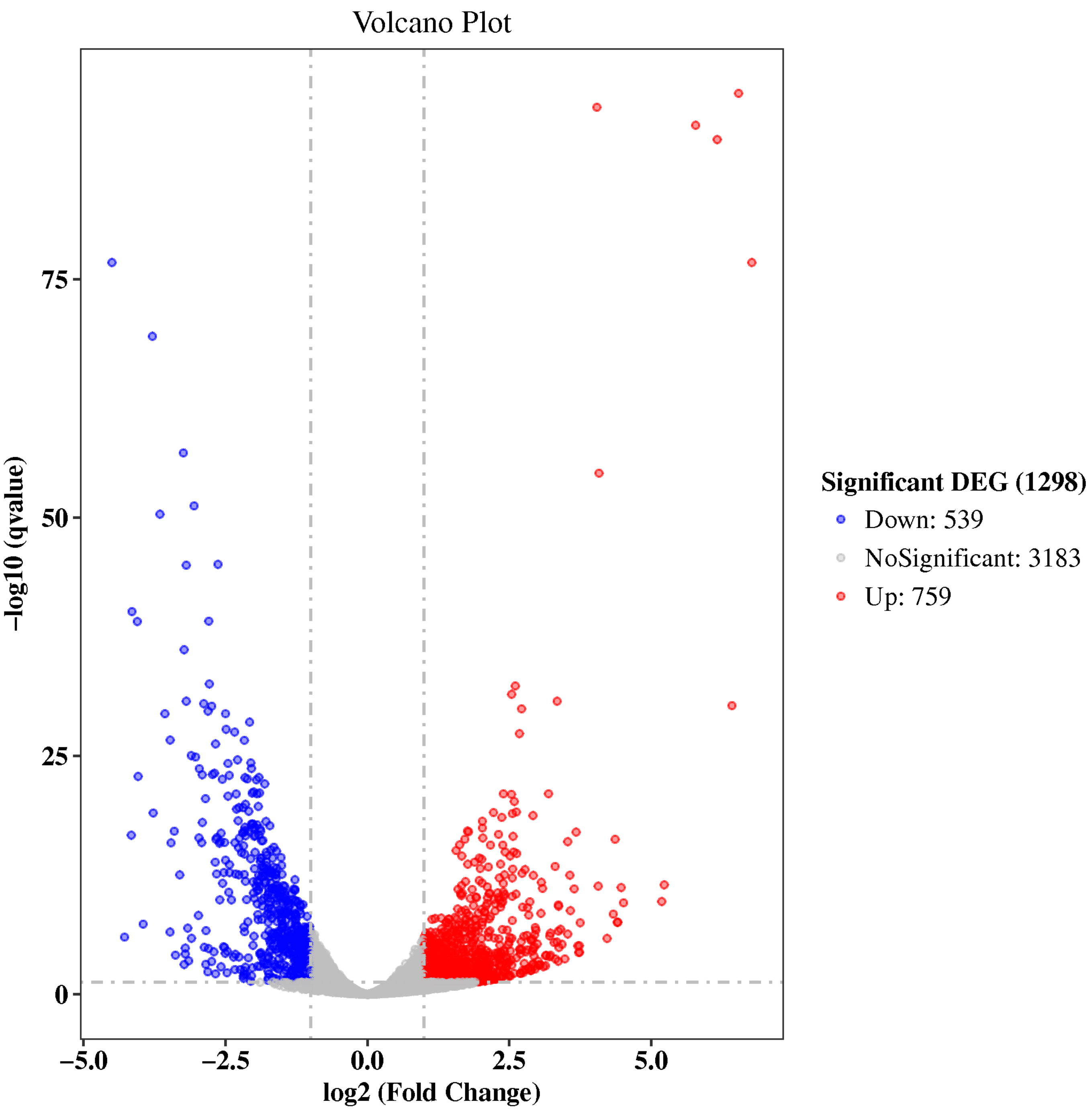

**Supplementary Figure 21. Volcano plot of differentially expressed genes from mRNA transcriptional analysis.** Volcano plot depicts differentially expressed genes between M9-glucose and M9-erythritol mediums (n=3 as independent biological replicates). The genes of more than two times or less than two times (q value ≤ 0.05) were recognized as differentially expressed.

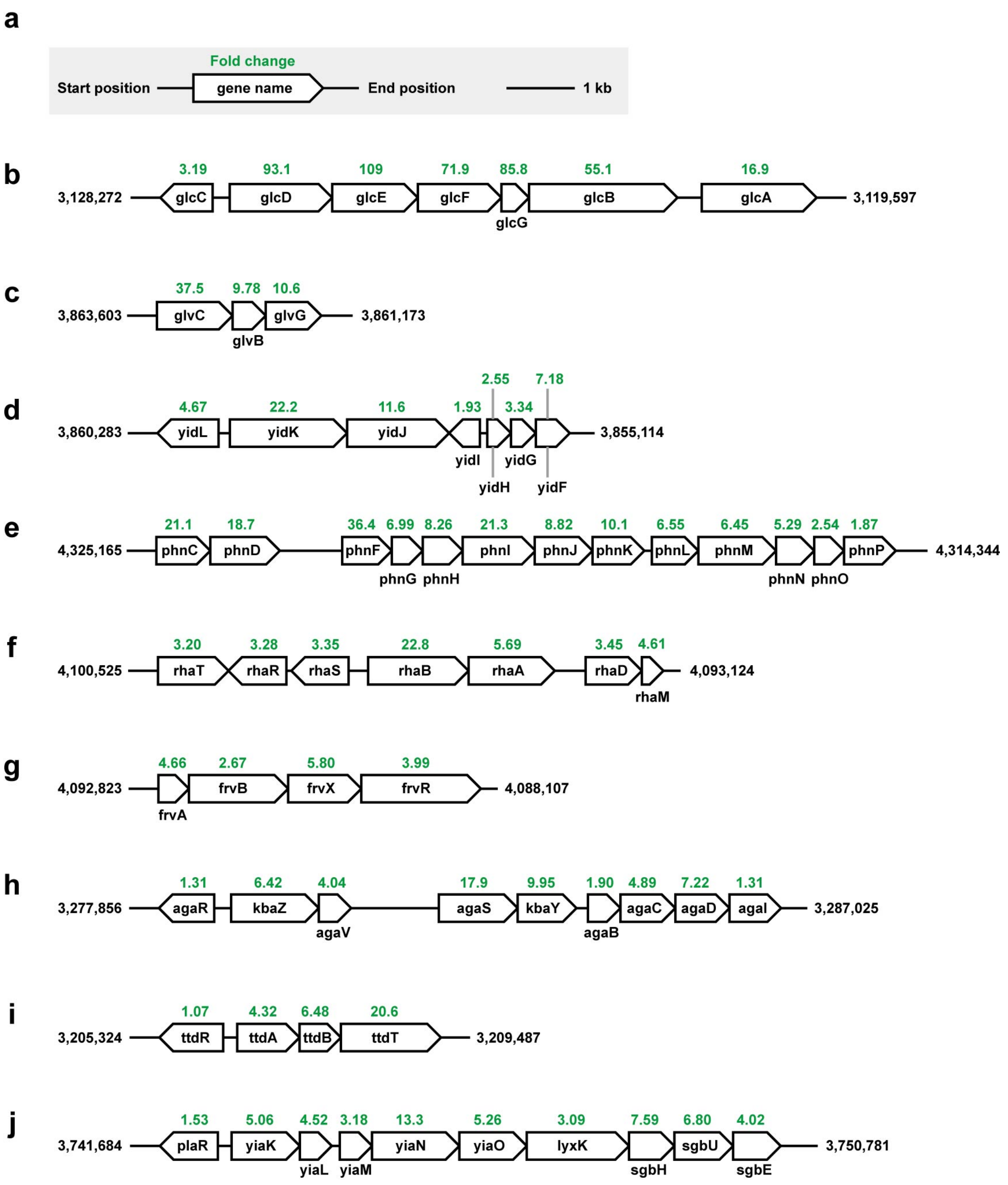

**Supplementary Figure 22. Some up-regulated gene clusters (RNA-seq) from M9-erythritol catabolic *E. coli* strain.** *E. coli* MG1655 genome sequence is used as a reference (GenBank: U00096.3)[13].

(**a**) Schematic diagram.

(**b**) Glycolate utilization cluster.

(**c**) *glv* operon as a putative PTS system.

(**d**) *yid* operon.

(**e**) *phn* operon as phosphonate uptake and utilization.

(**f**) *rha* operon as L-rhamnose uptake and utilization.

(**g**) *frv* operon as a putative PTS system.

(**h**) *aga* operon as N-acetylgalactosamine uptake and utilization.

(**i**) *ttd* operon as L-tartrate uptake and utilization.

(**j**) *yia-sgb* operon.

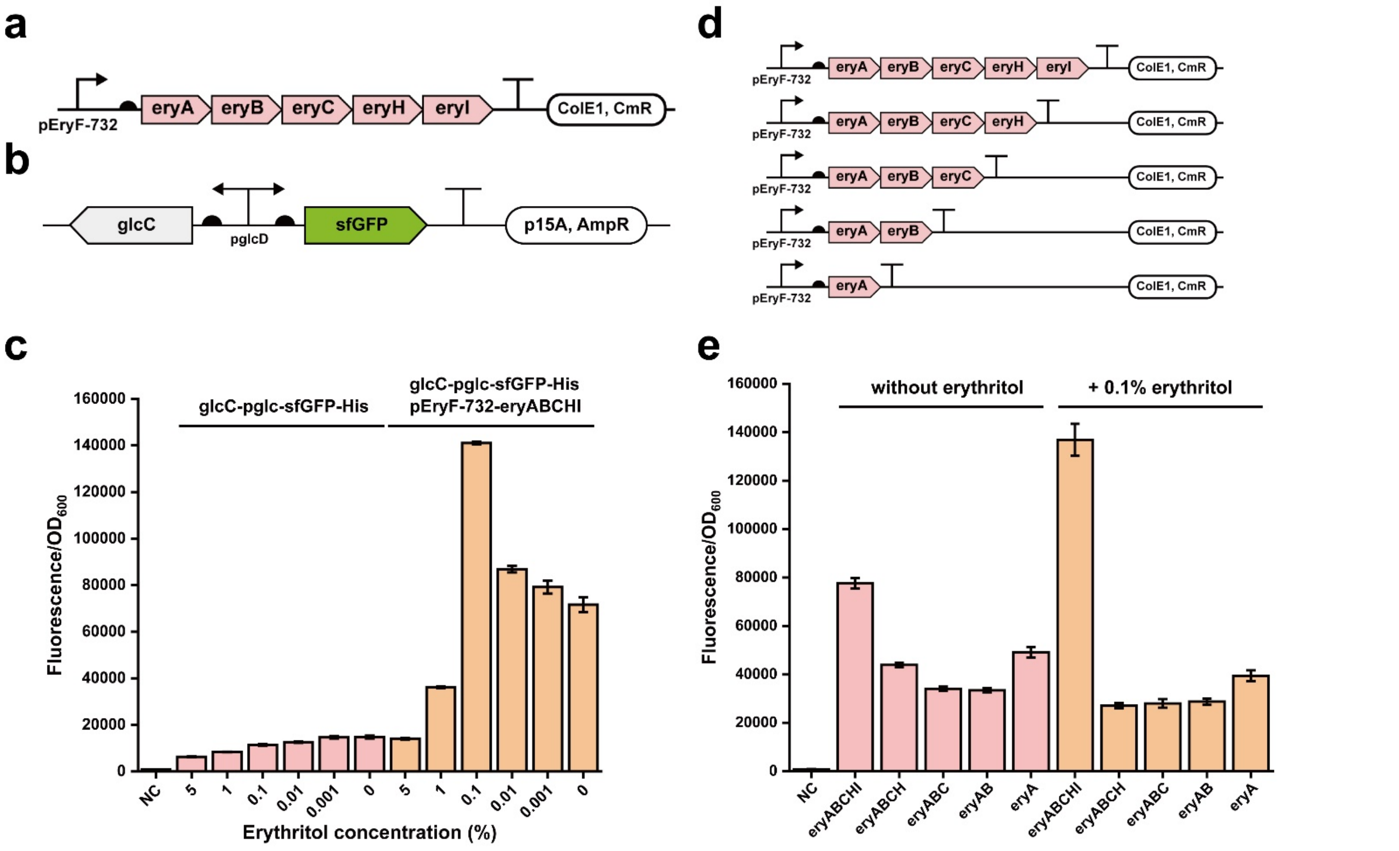

**Supplementary Figure 23. glcC can respond to erythritol catabolism.**

(**a**) Schematic of erythritol catabolism plasmid pFB147.

(**b**) glcC and putative glcC regulated promoter region pglcD were assembled with sfGFP as a reporter (plasmid pFB340).

(**c**) *E. coli* Mach1-T1 with plasmid pFB340 or pFB340 + pFB147 were incubated overnight with different concentrations of erythritol in 5 mL LB medium at 37°C and 250 rpm for 16 h. sfGFP fluorescence was measured with standardization. The results showed that with optimal 0.1% erythritol, glcC could well respond to erythritol catabolism-related molecules.

(**d**) A series of truncated erythritol catabolism plasmids (pFB147, pFB153 to pFB156) were constructed to determine whether glcC could respond to erythritol catabolism-related molecules.

(**e**) pFB340 co-existed with the truncated plasmids in *E. coli* Mach1-T1. Such strains were incubated overnight with or without 0.1% erythritol in 5 mL LB medium at 37°C and 250 rpm for 16 h. sfGFP fluorescence was measured with standardization. The results showed that glcC-responded substrate was not erythritol catabolism-related molecules.

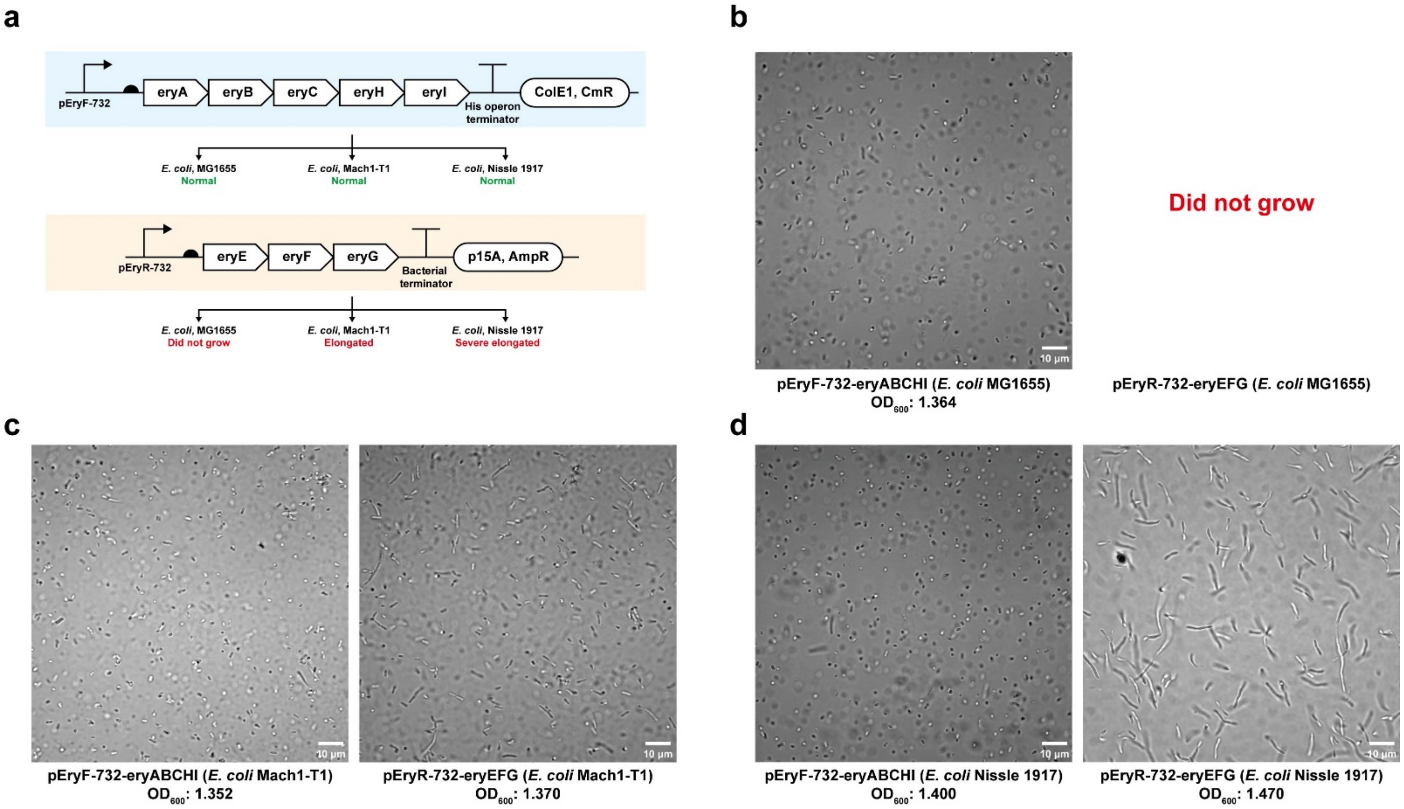

**Supplementary Figure 24. Erythritol ABC transporter cluster disturbs *E. coli* cell growth and division.**

(**a**) Schematic characterization process of whether erythritol catabolic cluster (pFB147) and erythritol ABC transporter cluster (pFB157) would influence cell division of different *E. coli* strains. All cell cultures were incubated in LB medium at 37°C and 250 rpm for 16 h.

(**b**) Cell morphology of *E. coli* MG1655. We could not pick up correct colonies with erythritol ABC transporter cluster, which means that it might cause severe cell toxicity.

(**c**) Cell morphology of *E. coli* Mach1-T1. The cells with erythritol ABC transporter cluster were elongated, which indicates that the ABC transporter could disturb cell division.

(**d**) Cell morphology of *E. coli* Nissle 1917. The cells with erythritol ABC transporter cluster were severe elongated, which indicates that the ABC transporter could severely disturb cell division.

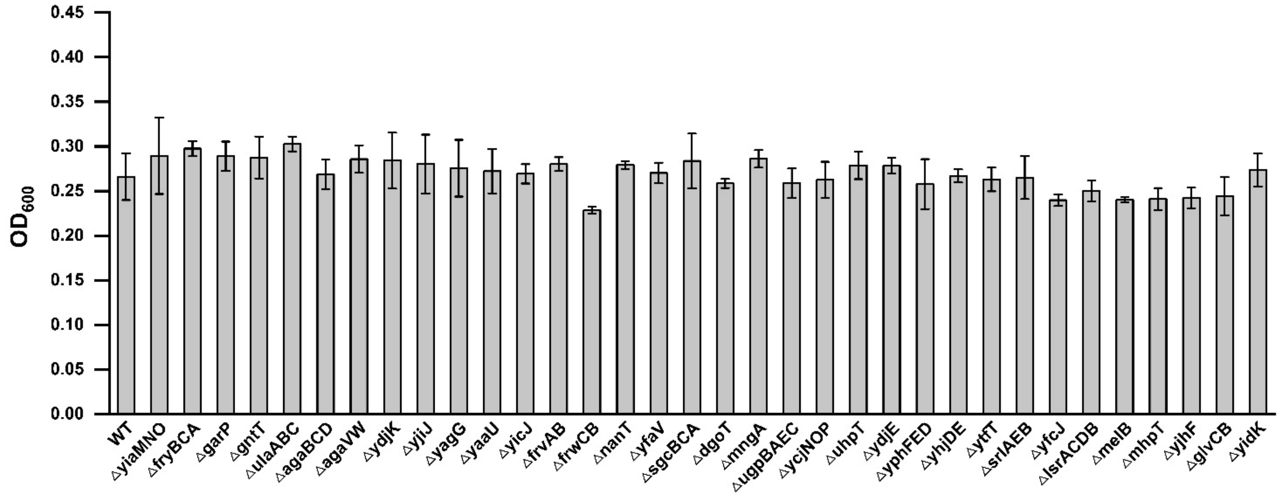

**Supplementary Figure 25. Up-regulated carbohydrate transporters were not essential for erythritol transport (from COG analysis of RNA-seq).**

COG analysis of RNA-seq indicated totally 34 up-regulated genes that might associate with "carbohydrate transport". All these genes (or gene clusters) were referred and we constructed 34 different knock-out *E. coli* MG1655 strains by Lambda-Red recombination[3]. All these 34 strains and WT (wild-type *E. coli* MG1655) strain were then transformed with pFB147 to utilize erythritol. After that, all strains were incubated (M9-erythritol medium, 37°C, no shaking) in 96-well plates for 108 h (stationary phase). $OD_{600}$ values were measured each 12 h. This graph showed the final $OD_{600}$ values at 108 h. All values were measured as three biological replicates. The error bars represent the standard deviation (s.d.). Student's *t*-tests are used for all the 34 genes' statistical analysis. Overall, there were no statistical significance ($p < 0.05$) between all the 34 measurements, which means that all the 34 genes were not essential for erythritol transport.

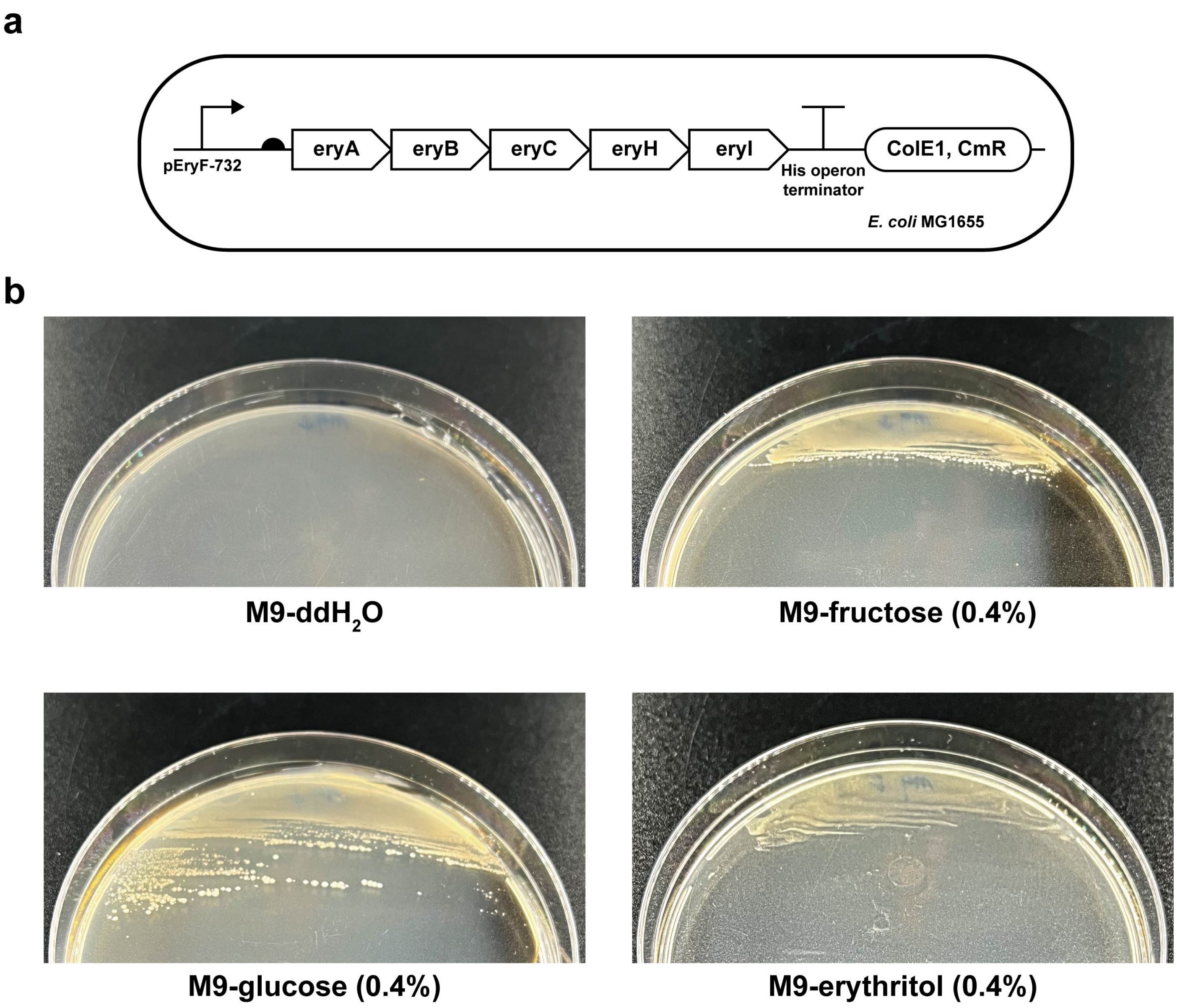

**Supplementary Figure 26. Erythritol catabolic *E. coli* strain utilizes different carbon sources.**
(**a**) Schematic diagram of *E. coli* MG1655 strain with the plasmid pFB147.
(**b**) The strain was incubated overnight in LB medium at 37°C and 250 rpm for 16 h. On the second day, cell pellets were washed with 1x phosphate-buffered saline (pH 7.4) for three times, then the suspension was spread onto different "M9-carbon source" agar plates and incubated at 37°C for 72 h. This experiment indicated that such "erythritol catabolic *E. coli* strain" could utilize fructose or glucose or erythritol as sole carbon source.

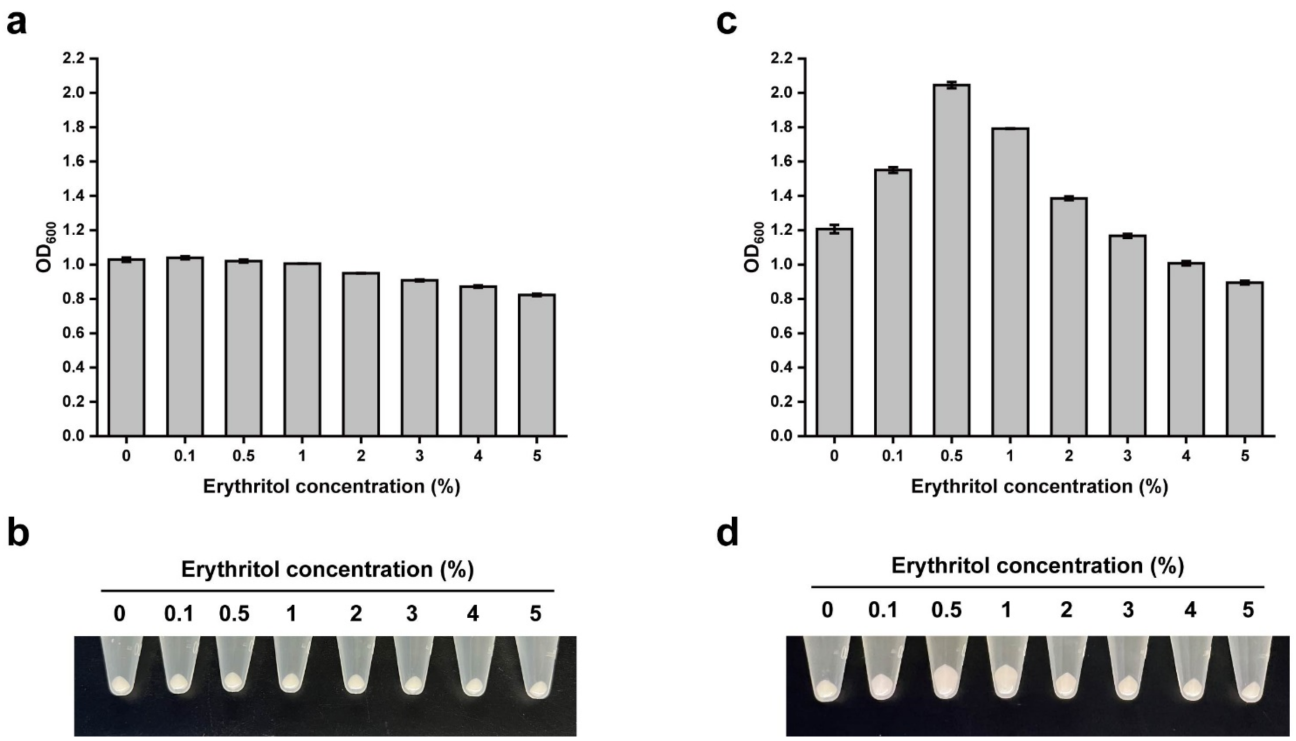

**Supplementary Figure 27. Additional erythritol catabolism promotes *E. coli* growth in LB medium.**
(**a**) Wild-type *E. coli* MG1655 without plasmids was incubated in 5 mL LB medium (without antibiotics). Additional erythritol was added to LB medium and then the cells were cultivated at 37°C, 250 rpm for 16 h. On the second day, the final $OD_{600}$ values were measured. The results showed that erythritol could not promote *E. coli* growth without the erythritol catabolic cluster.
(**b**) The same samples as described in (**a**). 1.5 mL different cell cultures were collected and centrifuged in 1.5 mL tubes. Cell pellets were accumulated at the tube bottom.
(**c**) *E. coli* MG1655 with plasmid pFB147 (erythritol catabolic cluster) was incubated in 5 mL LB medium (with 34 µg/mL chloramphenicol). Additional erythritol was added to LB medium and then the cells were cultivated at 37°C, 250 rpm for 16 h. On the second day, the final $OD_{600}$ values were measured. The results showed that erythritol could promote *E. coli* growth with the erythritol catabolic cluster, and the optimal erythritol concentration was 0.5% (w/v).
(**d**) The same samples as described in (**c**). 1.5 mL different cell cultures were collected and centrifuged in 1.5 mL tubes. Cell pellets were accumulated at the tube bottom.

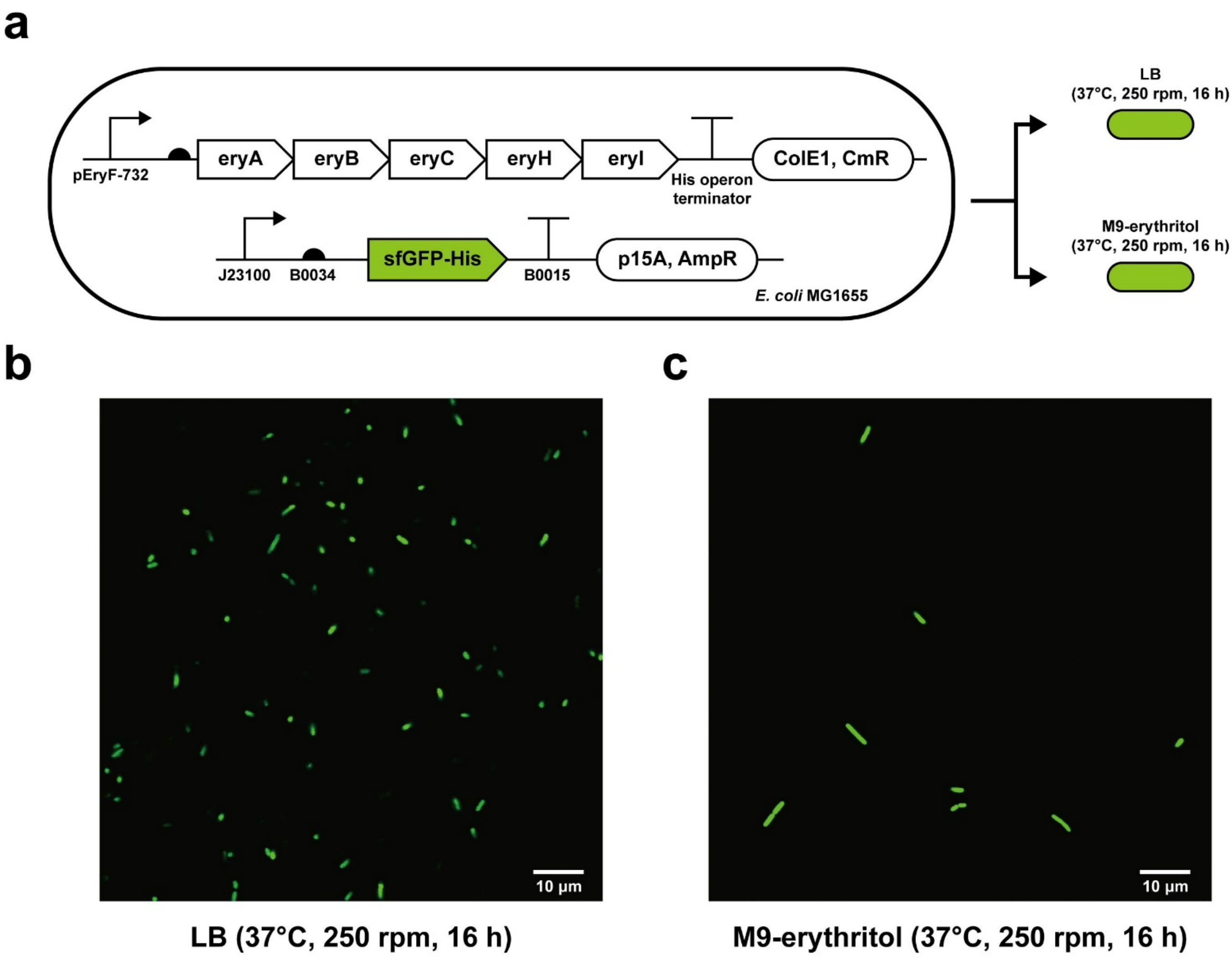

**Supplementary Figure 28.** ***E. coli*** **MG1655 with the erythritol catabolic cluster could express green fluorescent protein in M9-erythritol medium.**

(**a**) Schematic diagram of the engineered strain and plasmids (pFB147 and pFB285). This experiment aimed to test whether the *E. coli* strain with the erythritol catabolic cluster could express other heterologous proteins (e.g., sfGFP) in M9-erythritol medium. The same strain was incubated with both LB medium and M9-erythritol medium, respectively. After overnight cultivation, cell cultures were observed and imaged under confocal laser scanning microscopy.

(**b**) Confocal images of two cell cultures. The imaging parameters of the two samples were the same. We observed that in both cell cultures, sfGFP could be expressed and each cell showed a similar level of fluorescence intensity.

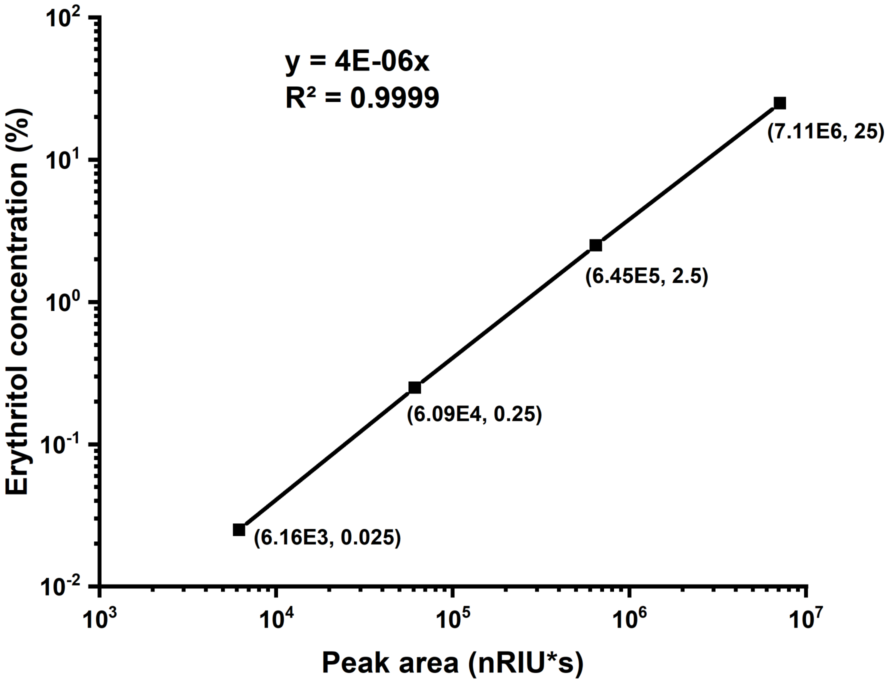

**Supplementary Figure 29. Standard curve of erythritol analyzed by HPLC (Refractive Index Detector, RID).** Each point was measured and calculated as an average value in triplicates.

III. Supplementary References

**References**

[1] *Registry of Standard Biological Parts*, <http://parts.igem.org/Main_Page?title=Main_Page> (2022).

[2] Ba, F., Liu, Y., Liu, W. Q., Tian, X., & Li, J. SYMBIOSIS: synthetic manipulable biobricks via orthogonal serine integrase systems. *Nucleic Acids Res.* **50**, 2973–2985 (2022).

[3] Datsenko, K. A. & Wanner, B. L. One-step inactivation of chromosomal genes in *Escherichia coli* K-12 using PCR products. *Proc. Natl. Acad. Sci. U.S.A.* **97**, 6640–6645 (2000).

[4] Altschul, S. F., Madden, T. L., Schäffer, A. A., Zhang, J., Zhang, Z., Miller, W., & Lipman, D. J. Gapped BLAST and PSI-BLAST: a new generation of protein database search programs. *Nucleic Acids Res*. **25**, 3389–3402 (1997).

[5] Johnson, M., Zaretskaya, I., Raytselis, Y., Merezhuk, Y., McGinnis, S., & Madden, T. L. NCBI BLAST: a better web interface. *Nucleic Acids Res*. **36**, W5–W9 (2008).

[6] Krogh, A., Larsson, B., von Heijne, G., & Sonnhammer, E. L. Predicting transmembrane protein topology with a hidden Markov model: application to complete genomes. *J. Mol. Biol*. **305**, 567–580 (2001).

[7] Reese M. G. Application of a time-delay neural network to promoter annotation in the *Drosophila melanogaster* genome. *Comput. Chem*. **26**, 51–56 (2001).

[8] Jumper, J. et al. Highly accurate protein structure prediction with AlphaFold. *Nature* **596**, 583–589 (2021).

[9] Varadi, M. et al. AlphaFold Protein Structure Database: massively expanding the structural coverage of protein-sequence space with high-accuracy models. *Nucleic Acids Res*. **50**, D439–D444 (2022).

[10] Kim, D. E., Chivian, D., & Baker, D. Protein structure prediction and analysis using the Robetta server. *Nucleic Acids Res*. **32**, W526–W531 (2004).

[11] Waterhouse, A. et al. SWISS-MODEL: homology modelling of protein structures and complexes. *Nucleic Acids Res*. **46**, W296-W303 (2018).

[12] Bienert, S., Waterhouse, A., de Beer, T. A., Tauriello, G., Studer, G., Bordoli, L., & Schwede, T. The SWISS-MODEL Repository-new features and functionality. *Nucleic Acids Res*. **45**, D313-D319 (2017).

[13] Blattner, F. R. et al. The complete genome sequence of *Escherichia coli* K-12. *Science* **277**, 1453–1462 (1997).
